# Characterization of bulk phosphatidylcholine compositions in human plasma using side-chain resolving lipidomics

**DOI:** 10.1101/626986

**Authors:** Jan D. Quell, Werner Römisch-Margl, Mark Haid, Jan Krumsiek, Thomas Skurk, Anna Halama, Nisha Stephan, Jerzy Adamski, Hans Hauner, Dennis Mook-Kanamori, Robert P. Mohney, Hannelore Daniel, Karsten Suhre, Gabi Kastenmüller

**Affiliations:** Institute of Bioinformatics and Systems Biology, Helmholtz Zentrum München - German Research Center for Environmental Health, Neuherberg, Germany; Experimental Bioinformatics, TUM School of Life Sciences Weihenstephan, Technical University of Munich, Freising-Weihenstephan, Germany; Research Unit Molecular Endocrinology and Metabolism, Helmholtz Zentrum München - German Research Center for Environmental Health, Neuherberg, Germany; Institute for Computational Biomedicine, Englander Institute for Precision Medicine, Department of Physiology and Biophysics, Weill Cornell Medicine, New York City, NY, USA; Institute of Computational Biology, Helmholtz Zentrum München - German Research Center for Environmental Health, Neuherberg, Germany; ZIEL Institute for Food and Health, Core Facility Human Studies, Technical University of Munich, Freising-Weihenstephan, Germany; Else Kroener-Fresenius-Center of Nutritional Medicine, Technical University of Munich, Freising-Weihenstephan, Germany; Weill Cornell Medical College in Qatar, Education City, Qatar Foundation, Doha, Qatar; Department of Biochemistry, Yong Loo Lin School of Medicine, National University of Singapore, Singapore; Institute for Nutritional Medicine, University Hospital Klinikum rechts der Isar, Technical University of Munich, Munich, Germany; Department of Clinical Epidemiology, Leiden University Medical Center, Leiden, The Netherlands; Department of Public Health and Primary Care, Leiden University Medical Center, Leiden, The Netherlands; Metabolon, Inc, Morrisville, NC, USA; Chair of Nutrition Physiology, TUM School of Life Sciences Weihenstephan, Technical University of Munich, Freising-Weihenstephan, Germany; German Center for Diabetes Research (DZD), Neuherberg, Germany

**Keywords:** metabolomics, lipidomics, phospholipids, isobaric phosphatidylcholines, lipid species, fatty acid composition, platform comparison, harmonization, imputation

## Abstract

Kit-based assays, such as Absolute*IDQ*™ p150, are widely used in large cohort studies and provide a standardized method to quantify blood concentrations of phosphatidylcholines (PCs). Many disease-relevant associations of PCs were reported using this method. However, their interpretation is hampered by lack of functionally relevant information on the detailed fatty acid side chain compositions as only the total number of carbon atoms and double bonds is identified by the kit. To enable more substantiated interpretations, we characterized these PC sums using the side chain resolving Lipidyzer™ platform, analyzing 223 samples in parallel to the Absolute*IDQ*™. Combining these datasets, we estimated the quantitative composition of PC sums and subsequently tested their replication in an independent cohort. We identified major constituents of 28 PC sums, revealing also various unexpected compositions. As an example, PC 16:0_22:5 accounted for more than 50% of the PC sum with in total 38 carbon atoms and 5 double bonds (PC aa 38:5). For 13 PC sums, we found relatively high abundances of odd-chain fatty acids. In conclusion, our study provides insights in PC compositions in human plasma, facilitating interpretation of existing epidemiological datasets and potentially enabling imputation of PC compositions for future meta-analyses of lipidomics data.

## Introduction

Phosphatidylcholines (PCs) are among the major constituents of eukaryotic cell membranes (1,2) and represent important components of lipoproteins (3). PCs consist of a polar phosphocholine head group, which is connected via a glycerol backbone to two fatty acid side chains of varying lengths and degree of saturation. The fatty acids are bound to the sn1 and sn2 positions of the glycerol backbone, either via two ester (acyl) bonds, or by one ester (acyl) and one ether (alkyl) bond. The abundance of PCs in human tissues and its relative abundance compared to other phospholipid classes, such as phosphatidylethanolamines (PEs), has been shown to play an important role in regulating energy metabolism (4). Furthermore, PC composition (i.e., fatty acid side chain lengths and desaturation levels) influences cell membrane properties, such as their fluidity. As an example, the fluidity of membranes that are made up mainly of PC (16:1/16:1) is higher than that of membranes containing mainly PC (16:0/16:1) or PC (16:0/16:0) (5). In case of PC (18:0/18:1), the fluidity is maximized when the double bond is located in the Δ9-position (6). Also for free fatty acids, which can be released from PCs by lipases (7), “minor” configurational molecular differences, such as in the position of the double bond in the fatty acid chain, can have a major functional impact. For instance, a balanced omega-6/omega-3 fatty acid-ratio of 4/1 associates with 70% lower mortality of patients with cardiovascular diseases (8).

Today, the availability of modern high throughput lipidomics approaches allows quantification of numerous PCs in thousands of blood samples from epidemiological cohorts. In various large-scale genome-wide and metabolome-wide association studies, many of these PCs have been reported to associate with common genetic variants and various diseases such as Alzheimer’s disease (9,10), type 2 diabetes (11–13), metabolic syndrome (14), coronary artery disease (15), and gastric cancer (16).

While the lipid side chain composition of PCs and the exact configuration of contained fatty acids induce major functional differences, most established lipidomics techniques cannot fully differentiate the specific compounds. Depending on the approach, phospholipids are measured to different degrees of structural resolution, starting at the coarsest level of lipid class/lipid species: For example, based on a precursor ion scan of the mass-to-charge ratio (m/z) 184 (= m/z of the phosphocholine head group), PCs and sphingomyelin (SM) species are selected and further characterized by their total m/z (e.g. PC (731)), which corresponds to the total numbers of carbon atoms and double bonds in the fatty acid side chains (17). This approach does not allow any differentiation of PC species at the bond-type level, i.e., PCs containing an ether bond (e.g. PC O-33:1) cannot be distinguished from isobaric diacyl-PC species (e.g. PC 32:1). Nonetheless, measures at the lipid level are often annotated with the assumption that even-numbered fatty acids are more common than odd-numbered fatty acids. For instance, PC (731) with m/z 731 covers PC 32:1 and the isobaric PC O-33:1 but is frequently labeled as PC 32:1. In contrast, MS-based lipidomics approaches that, for example, analyze fragmentation spectra can provide measures on the fatty acid level, i.e., they can differentiate between PC O-17:0_16:1 and PC 16:0_16:1. However, usually these fatty acid chain resolving techniques still do not provide any detailed information on the fatty acid position (e.g. PC O-17:0/16:1 versus PC O-16:1/17:0) or on the fatty acid structure (e.g. PC O-17:0/16:1(9Z)) (17,18).

The Absolute*IDQ*™ p150 kit (Biocrates Life Sciences AG, Innsbruck, Austria), which has been proven to provide robust, reproducible measurements (19), has generated large data sets of PCs for epidemiological studies (10,11,15,20,21). This method quantifies PCs on the lipid species level and annotates them under the assumption of even-numbered fatty acid chains (e.g. PC aa C32:1 for PC (731) and PC ae C34:1 for PC (745)) with aa indicating two acyl-bound and ae indicating one acyl-and one alkyl-bound fatty acids). As a consequence, only assumptions can be made regarding which specific PCs (i.e., which combination of fatty acid chain lengths and degrees of desaturation) dominate or significantly contribute to the concentration of the measured PC sum (e.g. PC 16:0_16:1, PC 14:1_18:0, and PC O-17:0_16:1 are all examples for possible constituents of PC aa C32:1). Due to the functional implications of fatty acid structures, mechanistic interpretation of observed genetic and disease associations with bulk PC measures is hampered.

In this study, we seek to reveal the fatty acid compositions of PCs in human plasma measured at the lipid species level to facilitate functional interpretation of existing association results. To this end, we combine quantitative data from the Absolute*IDQ*™ p150 and the fatty acid side chain resolving Lipidyzer™ platform powered by Metabolon® (Durham, USA). PC concentrations from both methods have been determined in 223 human plasma samples of healthy subjects. To test the transferability of identified PC compositions across different studies, we replicate our results in an independent cohort.

## Materials and methods

### Human plasma samples

Plasma samples were collected in the Human Metabolome (HuMet) study (22). The cohort consists of 15 healthy male subjects that have been selected based on following criteria: young age (22-33 y), no medication, and no abnormalities in standard clinical parameters. Participants were submitted to a series of metabolic challenges over four days: extended (36 h) fasting (FASTING) ingesting only 2.7 liters of mineral water, standard liquid mixed meal (SLD) in the form of a fiber-free formula drink supplying one third of their individual recommended daily energy, oral glucose tolerance test (OGTT) with 75 g glucose, physical exercise (PAT) with 30 min on a bicycle ergometer at their personal anaerobic power level, an oral lipid tolerance test (OLTT) with SLD with additional 35 g lipids per square meter of body surface area, and a stress test (STRESS) immersing one hand in ice water for a maximum of three minutes. Blood plasma samples were drawn at 56 different time points before, during and after challenges. All subjects gave their informed consent for inclusion before they participated in the HuMet study. The study protocol was approved by the ethical committee of the Technische Universität München (#2087/08) and corresponds with the Declaration of Helsinki.

In the present study, only 223 samples of four subjects (subject 5 – subject 8) were analyzed, as only for those data was available from both lipidomics methods.

The Qatar Metabolomics Study on Diabetes (QMDiab), in which we sought replication of our results, is a multi-ethnic diabetes case-control study, with participants aged between 17 and 82 years. Metabolite levels of 305 non-fasting blood plasma samples (152 male, 153 female) measured with both methods were available (23). All subjects gave their informed consent for inclusion before they participated in the QMDiab study. The study was conducted in accordance with the Declaration of Helsinki, and the protocol was approved by the Institutional Review Boards of HMC and Weill Cornell Medical College-Qatar (Research Protocol number 11131/11).

### Phosphatidylcholine quantification on the lipid species level (Absolute*IDQ*™ p150 kit)

For quantifying lipid sums, HuMet and QMDiab plasma samples were analyzed using the Absolute*IDQ*™ p150 kit (Biocrates Life Sciences AG, Innsbruck, Austria). Besides PCs (including lysophosphatidylcholines (lysoPCs)) and SMs, carnitines, amino acids, and hexoses are quantified by this targeted metabolomics approach. The corresponding analytical procedures have been described in detail before (19,22). In brief, 10 µl of human plasma were pipetted onto filter inserts (containing internal standards) in a 96 well plate. The filters were dried under a nitrogen stream, amino acids were derivatized by addition of a phenylisothiocyanate reagent (5%), and samples were dried again. After extraction with 5 mM ammonium acetate in methanol, the solution was centrifuged through a filter membrane and diluted with running solvent. Metabolites were detected by direct infusion to a 4000 QTRAP system (Sciex, Darmstadt, Germany) equipped with a Shimadzu Prominence LC20AD pump and SIL-20AC auto sampler. 76 PC, 14 lysoPC, and 15 SM species were measured in positive multiple reaction monitoring (MRM) scan mode selective for the common fragment ion of the phosphatidylcholine head group (m/z=184). The isobaric metabolites (within the mass resolution of the MS) PC *x*:*y* and PC O-*x*+1:*y* cannot be distinguished by this technique. Since odd chain lengths are considered rare for free fatty acids, measured PCs are principally labeled under the assumption of an even number of carbon atoms in the fatty acid chains, i.e., PC aa C*x*:*y* (for PC *x*:*y*) is chosen as label in case of even *x* and PC ae C*x*+1:*y* (corresponding to PC O-*x*+1:*y*) in case of odd *x*. Several internal standards were used for quantification of the phospholipid species. Concentrations were calculated by the Absolute*IDQ*™ kit software and reported in μmol/L. During quality control, metabolites with a coefficient of variation (CV) above 25% and metabolites that showed a significant correlation to the run day and had a CV above 20% (in samples of reference plasma measured along with the HuMet samples) were excluded, preserving 68 and 72 PC, 8 and 10 lysoPC, and 13 and 14 SM species from the HuMet and QMDiab samples for further analyses.

### Phosphatidylcholine quantification on the fatty acid level (Lipidyzer™)

For quantifying lipids resolved at the fatty acid level, plasma samples were analyzed on the Lipidyzer™ platform of AB Sciex Pte. Ltd., Framingham, USA, by Metabolon Inc., Durham, NC, USA. The method allows quantification of over 1100 lipid species from 14 lipid subclasses, including lysoPCs, PCs and SMs (24–28). Lipids were extracted from samples using dichloromethane and methanol in a modified Bligh-Dyer (29) extraction in the presence of internal standards with the lower, organic phase being used for analysis. The extracts were concentrated under nitrogen and reconstituted in 0.25 mL of dichloromethane:methanol (50:50) containing 10 mM ammonium acetate. The extracts were placed in vials for infusion-MS analyses, performed on a Sciex 5500 QTRAP equipped with the SelexION™ differential ion mobility spectrometry (DMS) cell, where ion separation is based on high-field asymmetric wave-form ion mobility mass spectrometry (FAIMS) (30,31). This additional device added to the electrospray ionization (ESI) source (applied in positive and negative mode) allows scanning or filtering molecules due to their ion mobility in alternating high and low electric fields before entering the mass spectrometer. With this technology a pre-separation of phospholipid classes (in PCs, lysoPCs, SMs, etc.), independent of their m/z ratio can be achieved. DMS-MS conditions were optimized for phospholipid classes. PCs were detected in negative MRM mode, with characteristic mass fragments for the fatty acid side chains, thus allowing a resolution of the fatty acid composition. In particular, several PC x:y with an odd chain fatty acid are reported by the Lipidyzer™ platform. These are identified by the characteristic fragment of the odd chain fatty acid that cannot be released from an isobaric PC-O x+1:y species with even-numbered side chains. Individual lipid species were quantified based on the ratio of signal intensity for target compounds to the signal intensity for an assigned internal standard of known concentration. For quantification of PCs, 10 stable isotope labeled compounds with a C16:0 fatty acid side chain in the *sn-1*-position and side chains in the range of C16:1 – C22:6 in the *sn-2*-position were used as internal standards (25,32).

### Qualitative description of phosphatidylcholine sums

For assigning PCs measured by Lipidyzer™ to Absolute*IDQ*™ p150 kit PC sums, we first collected all lipid isobars on the fatty acid precision level that can theoretically contribute to the PC *x*:*y* sum measure (annotated with PC aa C*x*:*y* in the Absolute*IDQ*™ p150 kit). To this end, we systematically distributed the total numbers of carbon atoms *x* and double bonds y to the two fatty acid chains, using the short hand notation PC *x*_*1*_:*y*_*1*__*x*_*2*_:*y*_*2*_ with *x=x*_*1*_*+x*_*2*_ and *y=y*_*1*_*+y*_*2*_ to indicate that we do not distinguish between the *sn-1* and *sn-2* positions of the side chains (17). Additionally, the isobaric compounds PC *x+1*:*y+7*, PC O-*x+1*:*y* and PC O-*x+2*:*y+7* were considered for PC *x*:*y*. Notably, we also included PCs containing odd chain fatty acids or fatty acids with very high desaturation to generate a comprehensive list of isobaric PCs. The software coming with the Absolute*IDQ*™ p150 kit, corrects concentrations of PC sums for the concentration of the (within the mass resolution) isobaric sphingomyelins [^13^C_1_]SM *x+4*:*y*, if the corresponding SM has been quantified. If the corresponding SM was not quantified by the kit (i.e., no isotope correction has been performed), we included [^13^C_1_]SM *x+4*:*y* in the list of possible lipid species measured under the PC *x:y* sum.

### Quantitative estimation of phosphatidylcholine sum composition

Both platforms report concentrations of lipid measures in μmol/L. To determine the proportion of a PC at the fatty acid level (measured on the Lipidyzer™ platform) within a PC sum (obtained by the Absolute*IDQ*™ p150 kit), we divided the concentration of the PC *j* (Lipidyzer™) by the concentration of the PC sum PC *i* (Absolute*IDQ*™ p150 kit) that includes this PC *j* according to our qualitative assignment (**Supplemental Table S1**): ratio 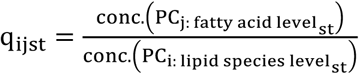. The characteristic composition of each PC sum is estimated by the mean q_ij_ of the respective ratios q_ijst_ over all subjects s and time points t. The variation was specified as 5 and 95% confidence interval, in which the 5% interval contains the lowest 5% and the 95% interval the largest 5% of the ratio values q_ijst_. Additionally, we determined fractions of all measured isobaric PCs of the fatty acid level, only using measures from the Lipidyzer™ platform: e.g. PC 16:0_20:2 / (PC 16:0_20:2 + PC 18:0_18:2 + PC 18:1_18:1).

### Replication of estimated quantitative phosphatidylcholine compositions in an independent cohort

Replication of estimated proportions q_ij_ (i.e., quotient of PCs of the fatty acid level (*j*) and the respective PC of the lipid species level (*i*)) was tested in the subset of 31 non-diabetic (control) male subjects aged between 17 and 40 years of the QMDiab cohort (to restrict the comparison to subjects more similar to the participants of the HuMet cohort). After running the cross-platform comparison of the selected QMDiab samples (as described above), we compared the identified main constituent for all PC sums, for which at least two PCs of the fatty acid level have been measured. In addition, we used a Kruskal-Wallis test (function ‘kruskal.test’ of R package ‘stats’ version 3.4.0 (33)) to check if the distributions of proportions q of HuMet-or QMDiab-subjects were significantly different (significance level α = 0.05/37 = 0.0014; with 37 testable PC sums, for which data was available in QMDiab).

### Variation of estimated phosphatidylcholine compositions between subjects and challenges

To assess the variation of PC composition within subjects, we determined the average standard deviation (SD) of the ratio q_ij_ observed for samples of the same individual at different timepoints for each PC sum *i* and PC constituent *j*. To this end, we considered “baseline” samples (drawn before the challenges), and “intervention” samples (at the timepoint with largest changes of the total metabolome compared to baseline) for each challenge with relation to lipid metabolism: FASTING: baseline at 8 am after 12 h overnight fasting; intervention at 36 h fasting; PAT: baseline at 4 pm (4 h after last meal); intervention after 30 min exercise; OLTT: baseline at 8 am after 12 h overnight fasting; intervention at noon. In addition, central trends of the ratio q_ij_ for baseline and intervention timepoints were substantiated by a Wilcoxon signed-rank test (function ‘wilcox.test’ with options ‘two sided’ and ‘paired’ of R package ‘stats’ version 3.4.0 (33)).

To assess the variation of PC composition between subjects, we determined the average SD of the ratio q_ij_ observed for samples at the same timepoint (baseline and intervention timepoints of each lipid-related challenge) in different subjects for each PC sum *i* and PC constituent *j*. We also compared the distributions of q_ij_ from all 56 timepoints of subjects using ANOVA (function ‘aov’ of R package ‘stats’ version 3.4.0 (33)) in case of normal distributions (tested by a Shapiro-Wilk test (function ‘shapiro.test’ of R package ‘stats’ version 3.4.0 (28)) and Kruskal-Wallis (function ‘kruskal.test’ of R package ‘stats’ version 3.4.0 (33)) otherwise. Finally, we compared the distributions of ratios q_*ij*_ obtained for male controls, female controls and all diabetics in QMDiab. A Kruskal-Wallis test was used to check if distributions were significantly different between those three groups.

### Estimation of unmapped part of phosphatidylcholine sums

To estimate to what extent the PC bulk measure (in μmol/L) is explained by the sum of all measured PC species (in μmol/L) at the fatty acid level, we applied linear models and Bland-Altman analyses. The formulas for linear models were constructed by PC_lipid species level_ ∼ *b* · ∑(PC_fatty acid level_) and evaluated by the ‘lm’ function of the R package ‘stats’, version 3.4.0 (33).

In the Bland-Altman plots, the difference of PC sums (lipid species level) and cumulated respective measured PCs of the fatty acid level (*conc*. (PC_lipid species level_) -∑*conc*. (PC_fatty acid level_)) were opposed to the mean of concentrations ((*conc*. (PC_lipid species level_) + ∑*conc*. (PC_fatty acid level_))/2) of both platforms.

## Results

In this study, we aimed at characterizing PC measures at the lipid class/species level by providing qualitative as well as quantitative compositions for these sum measures, which incorporate a variety of PCs at the fatty acid level. Our study is based on data from two targeted lipidomics approaches (Absolute*IDQ*™ p150 kit and Lipidyzer™ platform) for 223 human plasma samples of healthy human subjects. Using the Absolute*IDQ*™ p150 kit, 68 PCs have been quantified in our dataset. Concentrations of in total 109 PCs at the fatty acid level (i.e., providing the bond type as well as the lengths and degree of desaturation for the two fatty acid chains without differentiating between *sn-1* and *sn-2* positions) have been measured using the Lipidyzer™ platform. In a first step, we systematically collected all PCs of the fatty acid level that would theoretically fall under each of the 68 PC sums (lipid species level) in a list to provide a comprehensive description of their qualitative compositions. In a second step, we estimated the quantitative composition of 38 of the 68 PC sums for which at least one isobaric PC has been measured on the Lipidyzer™ platform with at least 25% coverage. Identification of the main PC constituent was replicated in an independent cohort for all 19 PC sums, to which more than one constituent on the fatty acid level was assigned. In a third step, we estimated the fraction of PC sums that could not be mapped to any measured PC of the fatty acid level.

### Qualitative composition of phosphatidylcholine sums

To map PCs of the fatty acid level to PCs of the lipid species level, we systematically distributed the total numbers of carbon atoms and double bonds to two fatty acid chains for all possible isobaric PCs (**Table 1**). The isobaric sphingomyelin [^13^C_1_]SM was also added as a possible compound, if it has not been quantified with the kit and, thus, has not been subtracted from the PC sum measure in the isotope correction step of the Absolute*IDQ*™ p150 kit processing (see Materials and Methods). To each of the 68 measured PC sums of the lipid species level, we thereby assigned between 150 and 456 theoretically possible isobaric compounds of the fatty acid level (**Supplemental Table S1**). In the supplemental table, we additionally tagged the PCs of the fatty acid level that were considered to be the most probable constituents of the PC sum (2 to 17 PCs per sum) according to Biocrates (34). In this table, we also highlighted those PCs that have been quantified on the Lipidyzer™ platform in our data set and that were assigned to one of the listed 68 PC sums (89 out of 109 PCs).

**Table 1:**
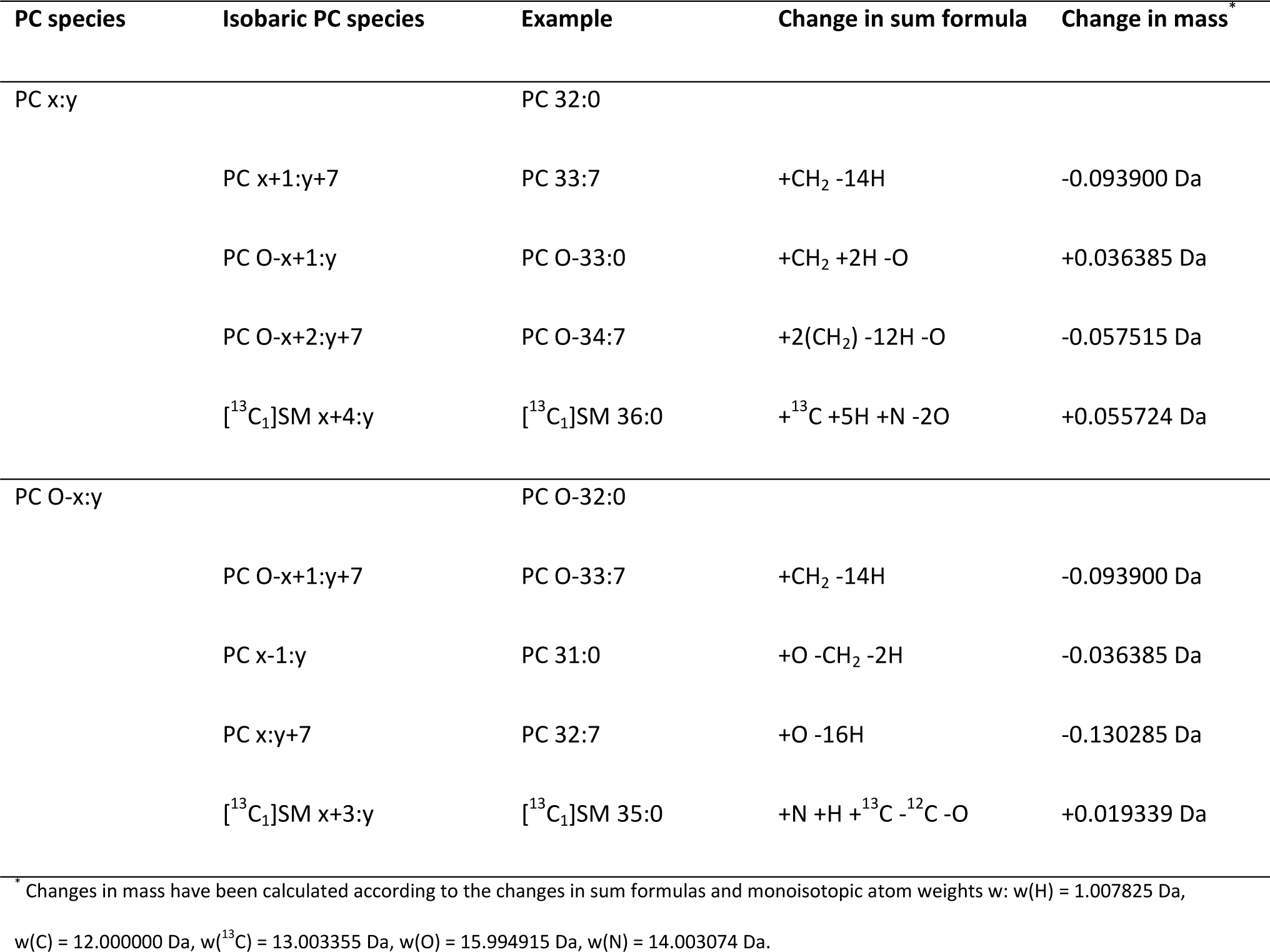
Isobaric phosphatidylcholine sums within mass resolution of the mass spectrometer.

| PC species | Isobaric PC species | Example | Change in sum formula | Change in mass<sup>*</sup> |
| --- | --- | --- | --- | --- |
| PC x:y |  | PC 32:0 |  |  |
|  | PC x+1:y+7 | PC 33:7 | +CH<sub>2</sub> -14H | -0.093900 Da |
|  | PC O-x+1:y | PC O-33:0 | +CH<sub>2</sub> +2H -O | +0.036385 Da |
|  | PC O-x+2:y+7 | PC O-34:7 | +2(CH<sub>2</sub>) -12H -O | -0.057515 Da |
|  | [<sup>13</sup>C<sub>1</sub>]SM x+4:y | [<sup>13</sup>C<sub>1</sub>]SM 36:0 | +<sup>13</sup>C +5H +N -2O | +0.055724 Da |
| PC O-x:y |  | PC O-32:0 |  |  |
|  | PC O-x+1:y+7 | PC O-33:7 | +CH<sub>2</sub> -14H | -0.093900 Da |
|  | PC x-1:y | PC 31:0 | +O -CH<sub>2</sub> -2H | -0.036385 Da |
|  | PC x:y+7 | PC 32:7 | +O -16H | -0.130285 Da |
|  | [<sup>13</sup>C<sub>1</sub>]SM x+3:y | [<sup>13</sup>C<sub>1</sub>]SM 35:0 | +N +H +<sup>13</sup>C -<sup>12</sup>C -O | +0.019339 Da |

<sup>\*</sup> Changes in mass have been calculated according to the changes in sum formulas and monoisotopic atom weights w: w(H) = 1.007825 Da, w(C) = 12.000000 Da, w(<sup>13</sup>C) = 13.003355 Da, w(O) = 15.994915 Da, w(N) = 14.003074 Da.

### Quantitative composition of phosphatidylcholine sums

For 38 (25 PC aa and 13 PC ae) out of the 68 PC sums quantified with the Absolute*IDQ*™ p150 kit, in total, 69 respective PCs of the fatty acid level (Lipidyzer™) were available at sufficient coverage (25%) in the HuMet Study (22) to estimate quantitative compositions. Each PC sum (lipid species level) consists of one to four measured PCs of the fatty acid level, e.g.: PC aa C36:2 = PC 16:0_20:2 + PC 18:0_18:2 + PC 18:1_18:1 + R, PC aa C34:3 = PC 14:0_20:3 + PC 16:0_18:3 + PC 18:2_16:1 + R, or PC aa C32:2 = PC 14:0_18:2 + R (**Supplemental Table S2**). Here, R corresponds to the sum of respective non-measured PCs of the fatty acid level that are specified in **Supplemental Table S1**. In the following, we use the fatty acid level of the short hand notation of lipids for PCs, e.g. PC *x*_*1*_:*y*_*1*__*x*_*2*_:*y*_*2*_ according to the resolution of the Lipidyzer™ technique (17). PCs of the lipid species level are labeled according to the Biocrates-specific notation, e.g. PC aa C*x*:*y* or PC ae C*x*:*y*, corresponding to PC *x*:*y* or PC O-*x*:*y* although the measurement did not allow differentiation of PCs on the bond type level; PC aa or PC ae labels were chosen under the assumption of even chain lengths of fatty acids.

In a first step, we calculated ratios q_ij_ to estimate the characteristic proportion of each measured PC of the fatty acid level *i* based on the concentration of the respective PC sum (lipid species level) *j* in the 223 HuMet samples (**Table 2**). As an example, three measured PCs of the fatty acid level, namely PC 16:0_20:2, PC 18:0_18:2, PC 18:1_18:1, are isobaric constituents of PC aa C36:2 measured at the lipid species level. The means of ratios PC 16:0_20:2/ PC aa C36:2, PC 18:0_18:2/ PC aa C36:2, and PC 18:1_18:1/ PC aa C36:2 over all subjects and time points [including their 5% and 95% confidence intervals] are 0.037 [0.023, 0.058], 0.967 [0.817, 1.14] and 0.099 [0.060, 0.147], respectively (**Figure 1A**), and, thus, represent estimates for the proportions of these three species in the PC sum. For 27 out of the 38 investigated PC sums, we identified at least one PC of the fatty acid level with a proportion greater than 20%. For three PC sums, we identified at least two PCs of the fatty acid level with q_ij_ > 0.2. As an example, for PC aa C34:3 the constituents PC 14:0_20:3 (0.048 [0.025, 0.073]), PC 16:0_18:3 (0.482 [0.354, 0.617]), and PC 18:2_16:1 (0.555 [0.381, 0.741]) were measured on the fatty acid level (**Figure 1B**). In ten cases, we found one major component with a proportion greater than 80%. Interestingly, for one PC sum labeled as the acyl-alkyl-PC (i.e., PC-O compound) PC ae C38:3, we found the diacyl-PC with an odd numbered fatty acid chain PC 17:0_20:3 measured on the fatty acid level as a major component of PC ae C38:3 (0.340 [0.225, 0.477]) (**Figure 1C**). For 11 PC sums, proportions for all measured constituents summed up to 80-120%. In four cases the sum of proportions was larger than 1.2.

**Table 2:**
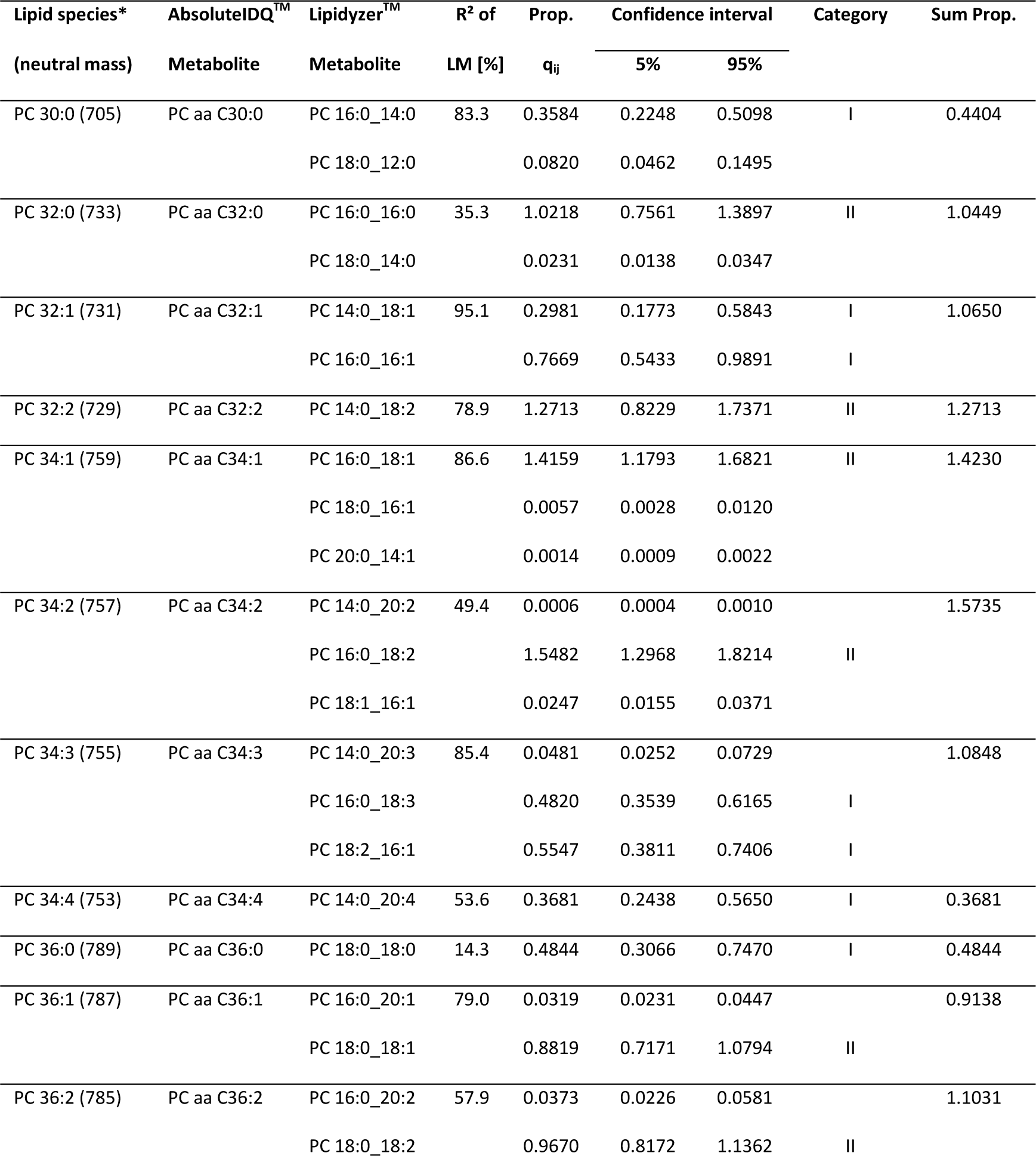
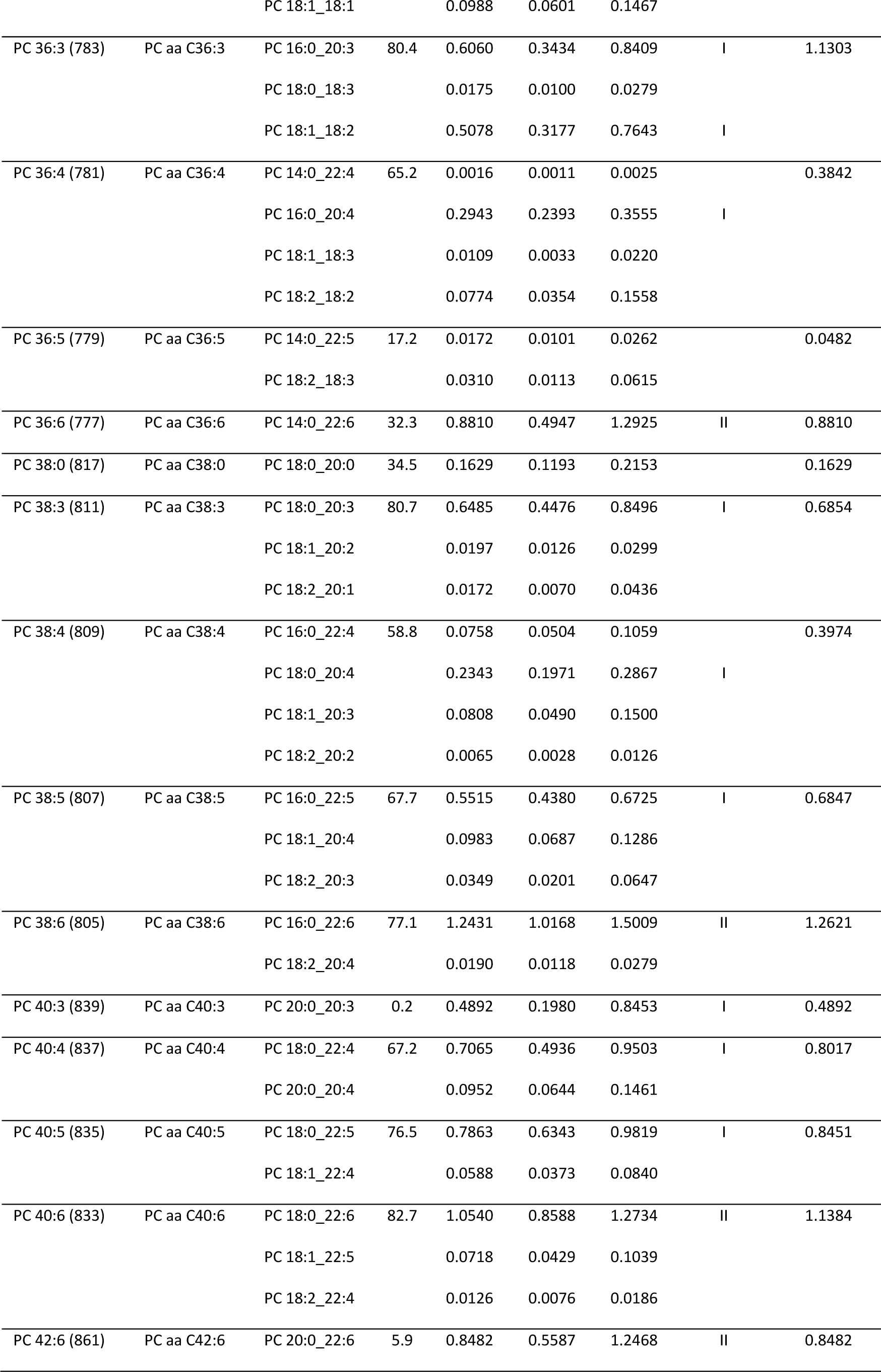
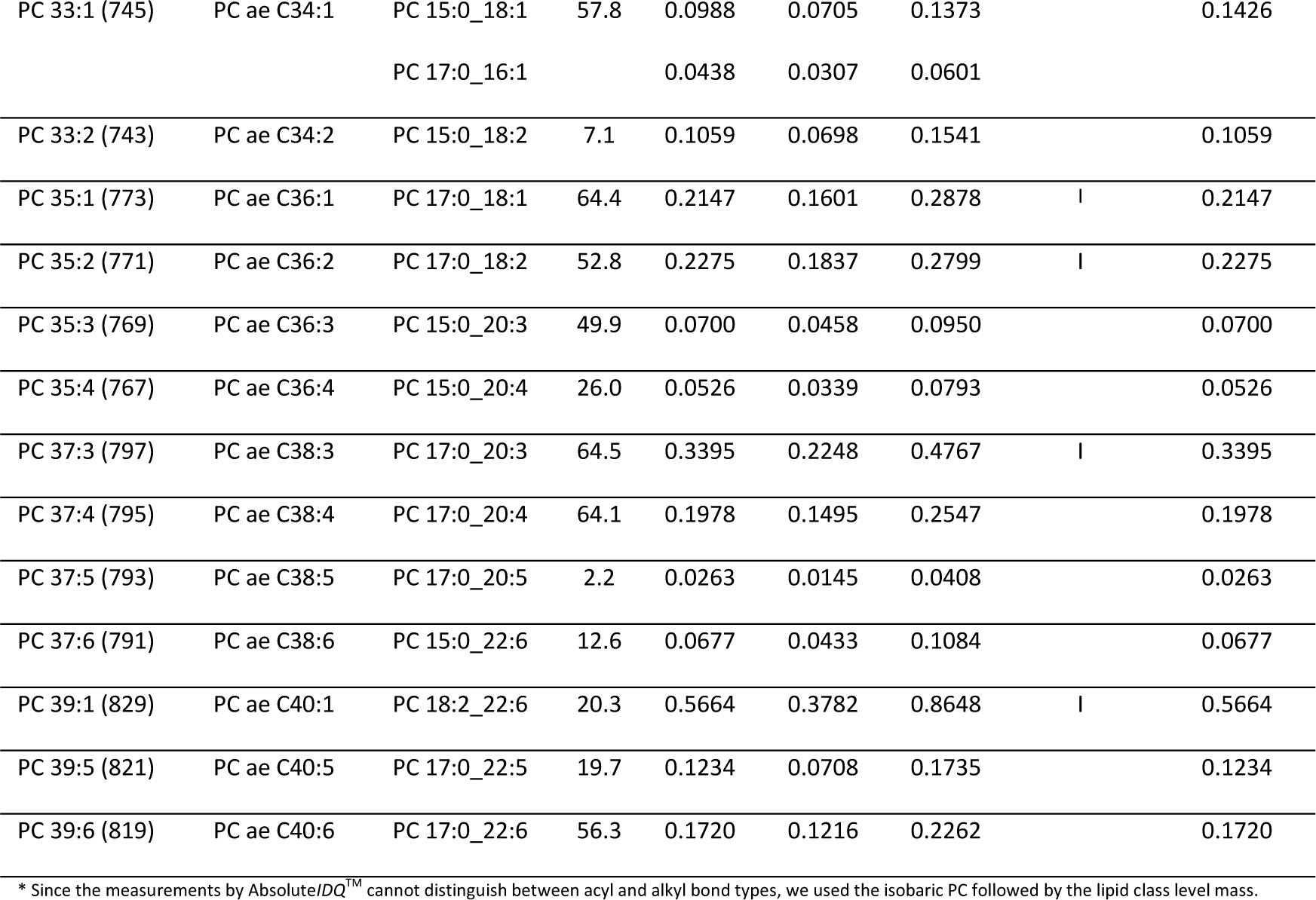
Quantitative composition of phosphatidylcholine sums.

**Figure 1:**
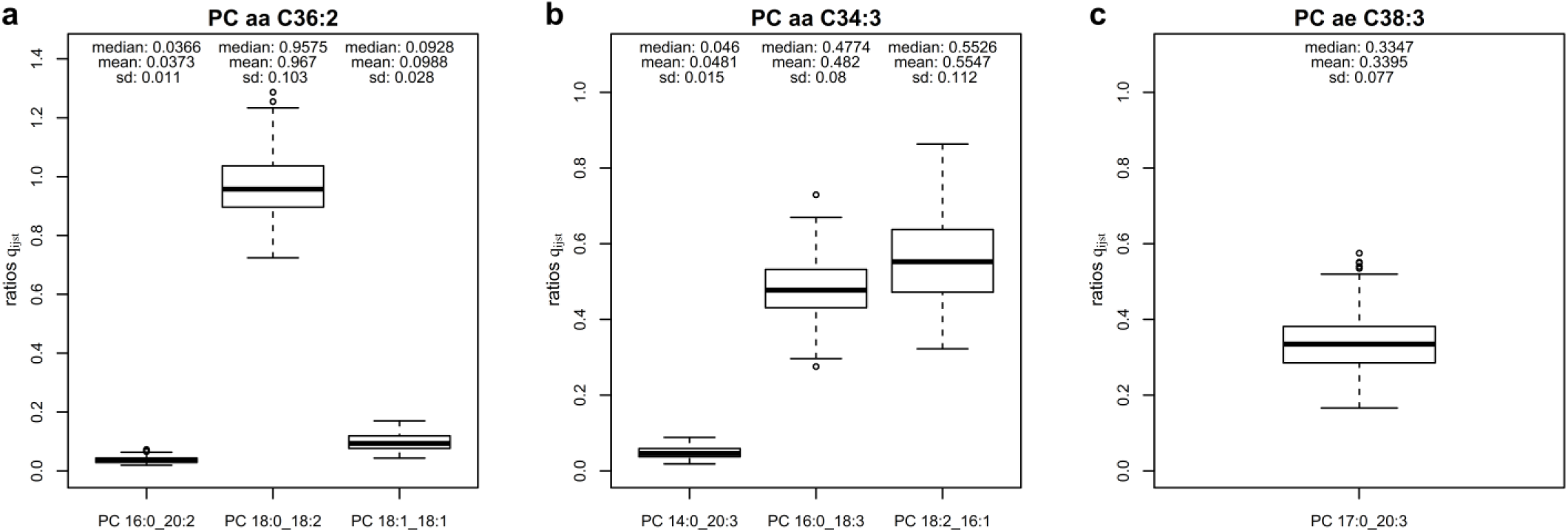
Examples of quantitative compositions of PC sums. The proportion q_ijst_ is defined as the ratio of the measured concentration (μmol/L) of the PC *j* resolved at the fatty acid level (e.g. PC 16:0_18:3) and the concentration (μmol/L) of the respective PC sum *i* at the lipid species level (e.g. PC aa C34:3) in a particular sample. (a) On average over all HuMet samples, PC aa C36:2 was mainly composed of PC 18:0_18:2 (96.7%). PC 18:1_18:1 and PC 16:0_20:2 only marginally contribute to the measured concentrations of PC aa C36:2 (9.88% and 3.73%, respectively. (b) In contrast to (a), PC aa C34:3, mainly consisted of two almost equally abundant PC species, PC 16:0_18:3 (48.2%) and PC 18:2_16:1 (55.5%). (c) As concentrations of acyl-alkyl-PCs were not reported for our samples, the diacyl-PC PC 17:0_20:3 was the only measured constituent of PC ae C38:3, for which we were able to calculate the proportions q_ijst_ in our data set. Interestingly, PC 17:0_20:3 with an odd-chain fatty acid accounted for a considerable fraction (33.95%) of the measured PC ae C38:3 sum.

Each PC sum resolved at the lipid species level (Absolute*IDQ*™ p150 kit) consists of a composition of PCs resolved at the fatty acid level (Lipidyzer™). A linear model estimated *R*^*2*^ as the percentage of variance of the PC sum that can be explained by the variances of the PCs of the fatty acid level. Proportions correspond to the mean of ratios of the PC at the fatty acid level (measured on the Lipidyzer™ platform) within the matching PC sum (obtained by the Absolute*IDQ*™ p150 kit) over all samples. The percentiles represent (5% and 95%) confidence intervals of proportions. The categories “I” and “II” correspond to proportions larger than 0.2 or larger than 0.8 respectively.

Since PC compositions may vary between individuals and depending on fasting state, we calculated and compared the standard deviation (SD) of the ratios q_ij_ of PC constituents separately for different time points (across subjects) and for the different subjects (across timepoints). For this analysis, we focused on baseline and intervention timepoints before and after lipid-related metabolic challenges, namely 36 h fasting (FASTING), physical activity test (PAT), and after ingestion of a lipid-enriched meal OLTT (see Methods), as we expected to observe the largest differences for them. As a result, we observed SD_challenges_ = 0.074 for the average intra-individual variation, and SD_subjects_ = 0.10 for the average inter-individual variation of q_ij_. No significant trends of the ratios q_ij_ between baseline and intervention timepoints were found for any challenge and any of the main PC constituents. However, the proportions of the main constituents of 13 PC sums were significantly different between subjects (**Supplemental Table S3**). When comparing the distributions of ratios q_ij_ considering all 56 timepoints, distributions were significantly different between subjects for 18 PC sums.

### Replication of quantitative compositions

To test the transferability of the proportions estimated from the HuMet study (healthy male subjects aged 22 – 33y of European ethnicity) to other populations and age ranges, we analyzed data from the two lipidomics methods for participants of the Qatar Metabolomics Study on Diabetes (QMDiab) with 151 healthy controls and 154 patients with diabetes (aged 17 – 82y). For 37 out of the 38 PC sums that we investigated, data were available in QMDiab. First, we sought replication of the main constituents (largest q_ij_) identified in HuMet in the 31 healthy male participants of QMDiab below 40y. For all 19 PC sums, for which at least two PC constituents were measured on the Lipidyzer™ platform, the same PCs of the fatty acid level were identified as the main constituents in QMDiab as in HuMet. However, for 23 PC sums, distributions of proportions of main PC constituents showed significant difference between the two cohorts (**Table 2, Supplemental Figure S2** and **Supplemental Table S4**). In general, estimated proportions (main and others) tended to be higher in samples of QMDiab, compared to HuMet (**Figure 2**). Largest differences between proportions in both cohorts were identified for PCs containing C20:4 with the exception of the proportions for PC 15:0_20:4 and PC 17:0_20:4.

**Figure 2:**
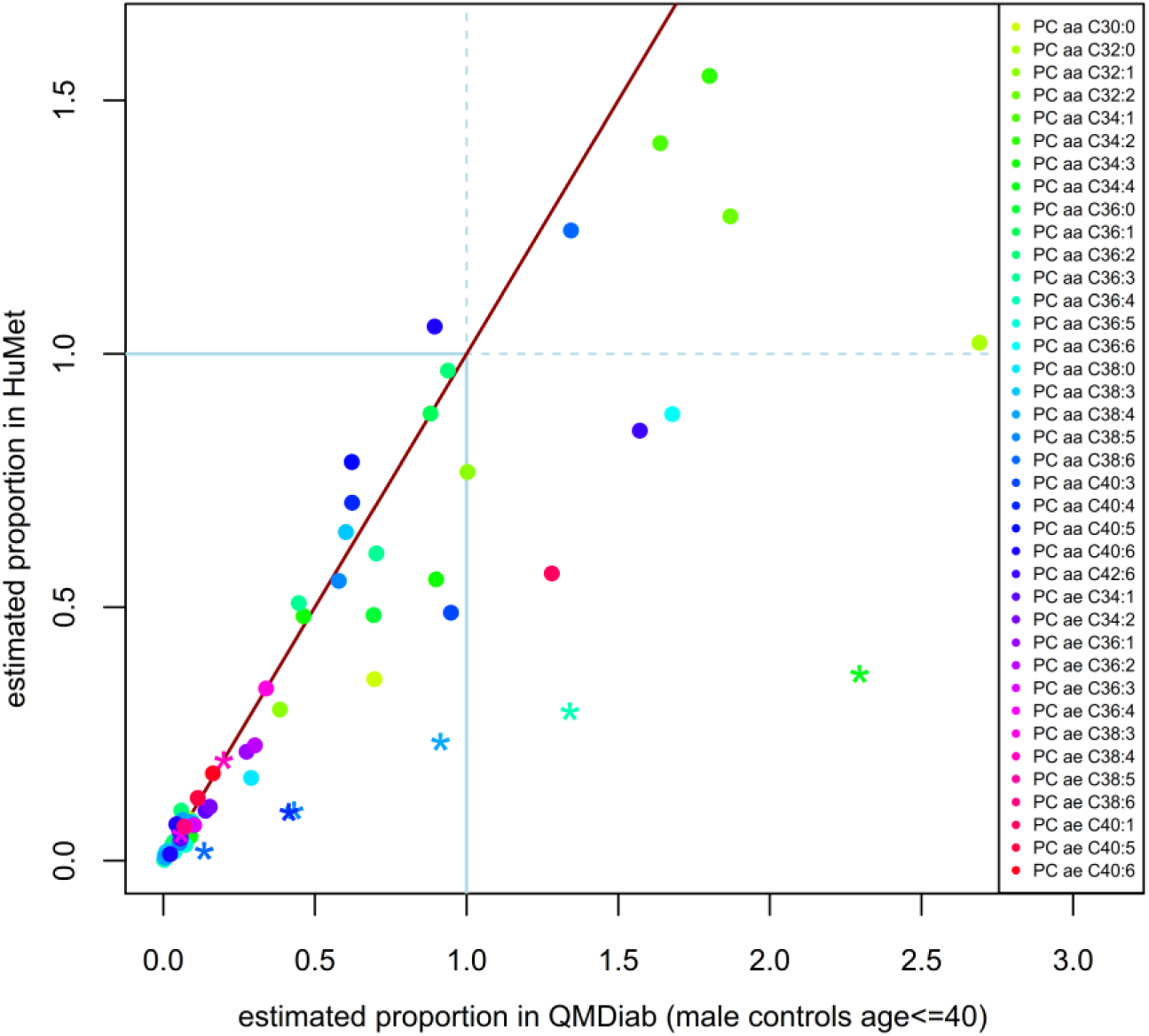
Comparison of characteristic proportions in HuMet and QMDiab. On average, proportions tend to be slightly higher in QMDiab than respective proportions in HuMet. The blue lines indicate the threshold where the concentration of the PC constituent measured on the Lipidyzer™ platform was higher than the respective containing PC sum measured with Absolute*IDQ*™ p150 kit. PCs of the fatty acid level that contain C20:4 as one of their lipid side chains are shown as asterisk. As QMDiab also includes samples from female participants and diabetes patients, we were able to compare proportions of PC constituents between these heterogeneous subgroups. Only for one PC sum, we observed a significant difference for the main (single) PC constituent PC 15:0_20:4 for PC ae C36:4.

### Estimation of the contribution of non-measured constituents in phosphatidylcholine sums

Out of 150 to 456 theoretically possible PCs of the fatty acid level per PC sum, the platform comparison only allowed mapping of one to four of these PCs to the respective sums to determine proportions (**Supplemental Table S1**). To estimate to which extent each PC sum could be explained by measured PCs of the fatty acid level in our study, we generated linear regression models (estimation of the explained variance; e.g. PC aa C36:2 ∼ b · (PC 16:0_20:2 + PC 18:0_18:2 + PC 18:1_18:1)) and Bland-Altman plots (comparison of absolute concentrations between both analytical methods) for the 38 PCs listed in **Table 2**.

In median, 58% (mean: 52%, SD: 28%) of the variance of PC sums could be explained by the corresponding sum of measured PCs of the fatty acid level (**Table 2** and **Supplemental Table S5**). In consequence, 42% of the variance of PC sums was related to further constituents or other factors such as experimental variation.

Since both methods reported metabolite levels in the same quantitative unit (micromolar), we also used Bland-Altman plots to investigate concordance of the PC sum measured on the lipid species level and the sums of constituents measured on the fatty acid level. Notably, here, concordance can only be expected for those PC sums for which all relevant constituents have been measured (**Supplemental Table S2**). In the plots, we related the differences between concentrations of the PC measure of the lipid species level and the sum of concentrations of mapped PCs of the fatty acid level (e.g. PC aa C36:2 – (PC 16:0_20:2 + PC 18:0_18:2 + PC 18:1_18:1)) to the average of the measure of the lipid species level and the sum of mapped measures on the fatty acid level. In general, small mean differences on the y-axis and small variation of these differences indicate concurrent measurements of both approaches. Relating these mean differences to the mean concentrations from both techniques on the x-axis of Bland-Altman plots enables identification of trends in discordance of measuring methods that depend on absolute values. In 19 of 25 PC aa compositions, the mean of this difference was within 2 SD of the PC aa measure, in 12 of those compositions (e.g., PC aa C40:4 = PC 18:0_22:4 + PC 20:0_20:4, **Figure 3A**)) mean difference was within 1 SD. In 10 cases, in which mean differences were negative, sum of concentrations of PCs of the fatty acid level were larger than those of PCs of the lipid species level (**Figure 3B**). In contrast to diacyl-PCs, PC ae sums could only be compared to the sum of isobaric diacyl-PCs containing an odd-numbered fatty acid chain, as phospholipid species with ether bonds have been reported on the Lipidyzer™ platform only for phosphoethanolamines in our data set. Those PCs with odd-numbered fatty acids are considered to be less abundant than the corresponding, unreported, alkyl-acyl-PCs. Supporting this expectation, all 13 PC ae sums showed positive mean deviations larger than 2 SD when compared to the sum of mapped measures on the fatty acid level. For several PCs, we observed systematic trends in the concordance between the compared sums depending on absolute concentration (e.g. PC aa C34:4, PC aa C36:4, PC aa C36:5, PC aa C38:0, **Supplemental Figure S1**).

**Figure 3:**
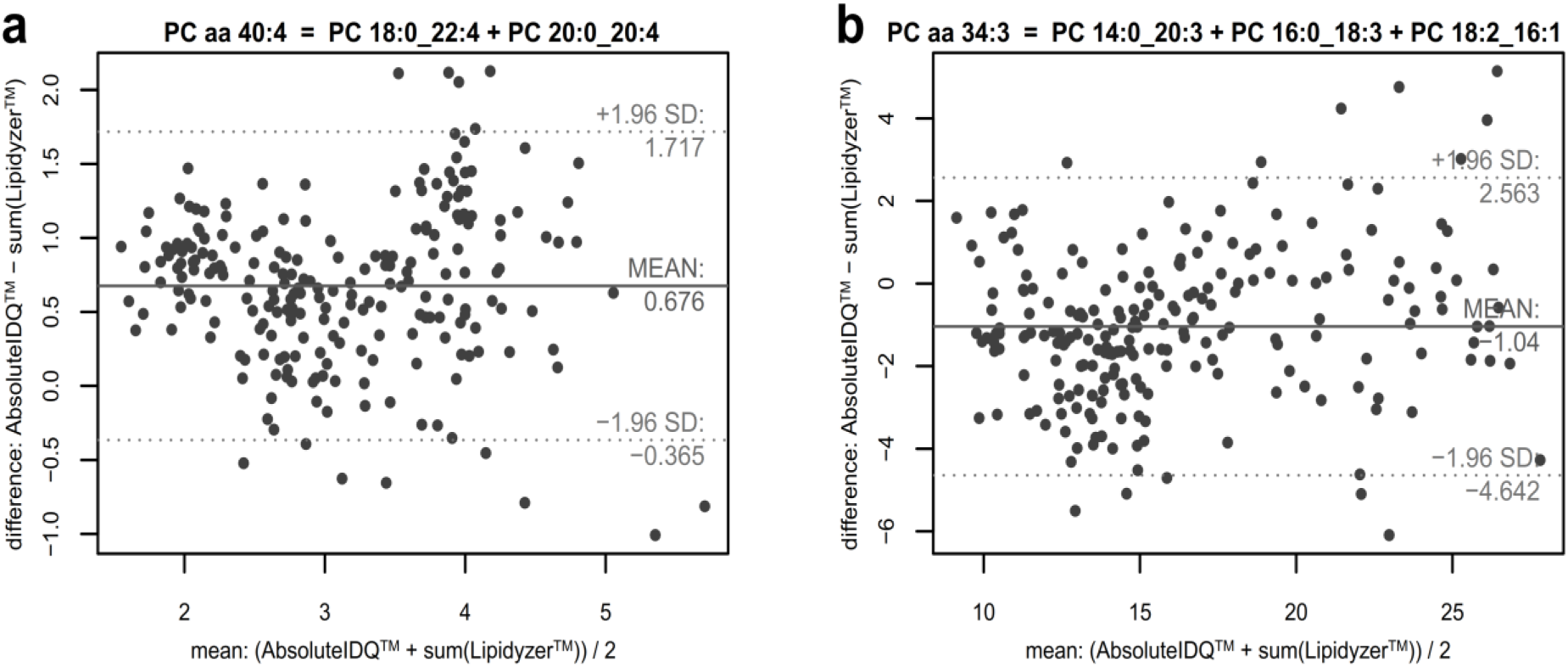
Deviation and mean of PCs of the lipid species level and sum of respective (measured) PCs of the fatty acid level. The Bland-Altman plots show the relative deviation of PC sums (measured on the lipid species level) and the sum of the respective PC constituents (measured on the fatty acid level) dependent on the mean of concentrations. In 19 of 25 cases the mean difference is within 2 SD of the concentration of the PC of the lipid species level.

## Discussion

Absolute*IDQ*™ p150 and p180 kits have emerged as established measurement kits for quantification of phospholipids, such as PCs, in blood from thousands of participants in large epidemiological cohorts (11,12,15,16,20,35–39) and in projects using a variety of human and non-human sample matrices (40–44). As the kits quantify PCs on the level of lipid species, each PC measure is composed of isobaric PCs (within mass resolution) with different combinations of fatty acid chain lengths and degrees of desaturation for the two residues. Depending on chain length and desaturation, biochemical properties such as membrane fluidity (5,6) or lipase activity (7) and, thus, the functions of these constituents, can vary significantly. To facilitate functional interpretation of identified associations between PC sum measures and the outcomes of interest, we here characterized PC compositions (i) by collecting all isobaric phosphocholine lipids on the level of fatty acids (with defined numbers of C atoms and double bonds) that, in theory, could be constituents of the measured PC sum, and (ii) by determining the contribution of constituents to the corresponding sum measure for those constituents that have been measured on the Lipidyzer™ platform in 223 human plasma samples with measurements from both analytical methods.

In a list provided by Biocrates, 2-16 PCs on the fatty acid level are mentioned as the most abundant variants for each of the 68 PC sums that we investigated in the present study (34). The selection of listed constituents is mainly based on knowledge from human plasma and serum. Some of the assumptions from human blood, such as higher abundance of fatty acids with even number of carbon atoms and specific limitations in the lengths of fatty acids, might not hold in other sample types. But also in human plasma interpretation of associations to PC sum measures might be hampered by focusing on the short-listed PC constituents. For reference, we therefore collected a comprehensive list of all arithmetically possible PC constituents on the fatty acid level for each of the 68 investigated PC sums quantified with the Absolute*IDQ*™ p150 kit. These lists also include PCs with fatty acids that presumably do not exist in human plasma such as very long chain fatty acids (e.g., C34) but might be present in plant extracts or other sample types. In this reference, we also included isobaric ^13^C isotopomers of sphingomyelins as in cases where the corresponding isobaric PC is of low abundance, the measured signal might be dominated by the first isotope peak of the not measured, “neighboring” sphingomyelin. While still ignoring the positions of double bonds and stereochemistry, our collection in total lists 150 to 456 constituents per PC sum measured with the Absolute*IDQ*™ p150 kit.

In human plasma, the majority of compounds in our reference list indeed are not expected to be present in considerable amounts. Based on the abundance of cellular free fatty acids, it can be assumed that PCs are mainly composed of the even numbered C16:0, C18:0, C18:1, C18:2 and C20:4 fatty acids as the most common fatty acids in human plasma (45–48). For 11 PC sums out of 23 PCs labeled as PC aa C32-C40, at least one PC containing two of those fatty acid chains was quantified on the Lipidyzer™ platform in the HuMet cohort. In 8 of the 11 cases, we could confirm the initial assumption as PCs composed of two highly abundant fatty acids were found at higher proportions than other combinations measured with Lipidyzer™. In 5 cases, those PCs alone accounted for more than 80% of the PC aa measure. However, for three cases, we also found notable exceptions in our data: (i) for PC aa C36:3, PC 16:0_20:3 was more abundant than PC 18:1_18:2; together they explained more than 80% of the PC sum; (ii) for PC aa C38:5, PC 16:0_22:5 was the major constituent (q_ij_ = 0.55), while PC 18:2_20:4 was only a minor constituent (q_ij_ = 0.098); (iii) for PC aa C38:6, PC 16:0_22:6 was by far the most abundant constituent and explained 77% of the PC sum; PC 18:2_20:4 contributed less to the measured PC sum (q_ij_ = 0.019). Although most proportions for constituents containing C20:4 were significantly larger in the independent QMDiab cohort, we identified the same constituents as major contributors to the corresponding PC sums in all three cases.

As no PCs with ether bonds were measured on the Lipidyzer™ platform in the HuMet cohort, we could not test directly whether the assumption that an alkyl-acyl-PC with even-numbered fatty acid chains were major components for the measures labeled as PC ae. The only constituents for 12 of the 13 PC ae sum measures that have been quantified in our data set were PCs containing an odd-numbered fatty acid chain. On average, proportions of these measured constituents were low (mean q_ij_ = 0.13), suggesting that indeed the not measured isobaric PC-O compounds with two even-numbered fatty acids might account for the major part of the PC sums labeled as PC ae. However, interestingly, we identified 3 PC ae sums containing notable amounts of odd-numbered constituents: PC ae C38:3 with q_ij_ = 0.34 for PC 17:0_20:3, PC ae C36:2 with q_ij_ = 0.23 for PC 17:0_18:2, and PC ae C36:1 with q_ij_ = 0.21 for PC 17:0_18:1.

In 1 of the 13 PC ae sums, namely PC ae C40:1, only PC 18:2_22:6 was measured as possible isobaric constituent (following the rule that PC-O x:y is isobaric with PC x:y+7; see **Table 1**). Notably, the estimated proportion for PC 18:2_22:6 was 0.57 and, thus, this diacyl-PC contributed considerably to the PC sum labeled as the acyl-alkyl-PC PC ae C40:1 in the Absolute*IDQ*™ p150 kit, posing the question whether this measure should rather be annotated with PC aa C40:8.

Due to availability of QMDiab as a replication cohort and of multi-timepoint data in the HuMet cohort, we were able to investigate variation of PC compositions depending on the individual and his/her fasting status. For various PCs, we observed significant differences in their compositions in terms of the estimated proportion of constituents. In particular, constituents that contained arachidonic acid (C20:4) showed higher proportions in QMDiab. We observed the same trend when analyzing QMDiab subsets (male non-diabetic controls, female non-diabetic controls, and diabetic patients) separately, excluding diabetes as the main reason for this observation. Further factors that could underlie the detected differences in C20:4 abundance between the two cohorts could be general health status (QMDiab diabetics and controls were patients in a dermatological clinic), nutrition, ethnicity, and pre-analytic processing among others. However, despite the observed differences in PC compositions, it has to be noted that the same PCs were identified as main constituents consistently in all subsets that we tested, demonstrating the transferability of our qualitative results to other studies in human plasma.

### Limitations

A major limitation in our study is that only for a small number of the possible PC constituents, measures are available on the fatty acid level, preventing a comprehensive overview of the actual compositions. For most PC sums with large cumulative proportions for the measured constituents, the sum of measured constituents also largely explained the variance in the data (R^2^). However, in some cases, we estimated large proportions, indicating that measured constituents are main parts of the PC sum, but very low R^2^ (e.g. PC aa C42:6 ∼ PC 20:0_22:6 with q_ij_ = 0.7 and R^2^ = 5.9%; PC aa C40:3 ∼ PC 20:0_20:3 with q_ij_ = 0.5 and R^2^ = 0.2%). The increased variation of proportions q_ij_ observed between subjects may indicate high inter-individual differences in compositions of the respective PC sums. Another reason for the discordance between proportions and explained variance could also result from overestimation of proportions due to differences in extraction and/or ionization efficiency between the two lipidomics approaches. Investigation of such differences in more detail would require measures for all possible (or all relevant) constituents. These differences in extraction and measurement procedures could also explain the estimation of proportions larger than 100% in our data. For four PCs, measured concentrations of constituents on the fatty acid level were much larger than the concentrations of the respective PC sums (lipid species level) with proportions q_ij_ = 1.2 to 1.5.

We used the fatty acid side chain-resolving Lipidyzer™ platform to characterize the molecular constituents of PC sums measured on the lipid species level with an unknown composition of chain lengths and double bonds. While characterizing PC sums by their constituents on the fatty acid level enables interpretations in a more precise biochemical context, PCs of the fatty acid level still denote classes of molecules rather than one compound with a defined chemical structure and function as the *sn-1* and *sn-2* positions are not distinguished and the positions of double bonds and their stereochemistry cannot be determined. Although we could not investigate these structural properties in our study, it has been shown previously that specific side chains prefer one of the *sn-1* or *sn-2* positions of PCs in human plasma (in 95% of cases 16:0 is bound to the *sn-1* position, 96% of 18:0 to position *sn-1*, 76% of 18:1 to position *sn-2*, 89% of 18:2 to position *sn-2* and almost 100% of 20:4 to position *sn-2*) (46). Notably, these previous findings can only be used for interpretation if compositions of the PCs are known at least on the fatty acid level.

Although PC compositions were qualitatively similar across cohorts and timepoints in human plasma samples from healthy and diabetic subjects in the two investigated studies, compositions might be different in other diseases. Also PC compositions identified in this study presumably cannot be readily transferred to other tissues or organisms (49).

### Concluding remarks

By systematically combining measurements from two lipidomics approaches, namely Absolute*IDQ*™ p150 kit and Lipidyzer™ platform, which quantify phosphatidylcholines on different levels of structural resolution, we here unmasked the composition of phosphatidylcholine sum measures in human plasma from healthy subjects. Knowing the composition of these sums facilitates the interpretation of associations that have been previously identified from the vast amount of data from the Absolute*IDQ*™ p150 kit in large epidemiological cohorts in a more precise biochemical context. In addition, the estimated proportions of constituents, which we determined in our study, might enable imputation of phosphatidylcholines at the fatty acid level based on concentrations of phosphatidylcholine sums to some extent. The possibility of imputation is key to combine studies with data from different lipidomics approaches for large meta-analyses in cases where re-measurement on a side chain-resolving lipidomics platform such as Lipidyzer™ is not an option.

## Abbreviations

CV: coefficient of variation
conc.: concentration
mGWAS: genome-wide association study with metabolic traits
MWAS: metabolome-wide association study
OGTT: oral glucose tolerance test
OLTT: oral lipid tolerance test
PC: phosphatidylcholine
pv: p-value
R: rest that consists of the totality of non-measured single PCs
SD: standard deviation

## Acknowledgements

Studies related to this research were funded in part by the Else Kroener-Fresenius-Foundation, Bad Homburg v.d.H., Germany. The QMDiab study was supported by the Biomedical Research Program at Weill Cornell Medicine – Qatar (WCM-Q), a program funded by the Qatar Foundation. We thank the staff of the HMC dermatology department and of WCM-Q for their contribution to QMDiab. GK is supported by National Institute on Aging (NIA) grants RF1 AG058942-01 and R01-AG057452-01, R01-AG059093-01, U01-AG061359-01, and the Qatar National Research Fund NPRP8-061-3-011. The statements made in this manuscript are solely the responsibility of the authors. Finally, we are grateful to all study participants of the HuMet and the QMDiab studies for their invaluable contributions to this study.

## Author contributions

Conceptualization: G.K.; Data curation: W.R.M, T.S., A.H., N.S., J.A., H.H., D.M.K., R.P.M., H.D. and K.S.; Formal analysis: J.D.Q.; Funding acquisition: K.S. and G.K.; Investigation: J.D.Q., W.R.M., M.H. and G.K.; Methodology: J.D.Q., W.R.M., J.K. and G.K.; Project administration: G.K.; Resources: W.R.M, M.H., J.K., T.S., A.H., N.S., J.A., H.H., D.M.K., R.P.M., H.D. and K.S.; Software: J.D.Q.; Supervision: G.K.; Validation: J.D.Q. and K.S.; Visualization: J.D.Q.; Writing – original draft: J.D.Q., W.R.M. and G.K.; Writing – review & editing: J.D.Q., W.R.M., M.H., J.K., T.S., A.H., N.S., J.A., H.H., D.M.K., R.P.M., H.D., K.S. and G.K.

## Conflicts of interest

Robert Mohney is employee of Metabolon Inc.

## Supplemental information

**Supplemental Table S1: Qualitative composition of PCs measured by Absolute*IDQ*™ p150 kit**

The chain length and double bonds of PCs of the lipid species level were systematically distributed to obtain isobaric molecules of the fatty acid level (150 to 456). If the isobaric sphingomyelin [^13^C_1_]SM x+4:y for the PC of the lipid species level PC x:y has not been measured and therefore has not been available for isotope correction, it was also included as possible constituent in the list.

**Supplemental Table S2: Isobaric measures from the Absolute*IDQ*™ p150 kit and the Lipidyzer™ platform with ranges of metabolite levels**

PCs of the lipid species level, measured by the Absolute*IDQ*™ p150 kit, are compared to the summed levels of related PCs of the fatty acid level measured by Lipidyzer™. Metabolites marked by an asterisk were excluded because of a missingness >75%.

**Supplemental Table S3: Variation of ratios q describing PC compositions**

The inter-and intra-individual variation of ratios q of PCs of the fatty acid level (Lipidyzer™) and PCs of the lipid species level (Absolute*IDQ*™ p150 kit) was assessed by comparing their distributions.

**Supplemental Table S4: Replication of findings in QMDiab**

Comparison of compositions of PCs between the HuMet and the QMDiab study and subsets thereof (male controls, female controls and cases) showed that the same PCs of the fatty acid level were identified as major compounds in the two studies.

**Supplemental Table S5: Explained variance of PCs of the lipid species level**

In median 67.2% (mean: 0.586, SD: 0.277) of the variance of the PCs of the lipid species level (Absolute*IDQ*™ p150 kit) can be explained by the sum of related PCs of the fatty acid level (Lipidyzer™).

**Supplemental Figure S1: Details of PC compositions**

This supplemental figure contains all plots and tabular information of the results part for each investigated PC composition.

**Supplemental Figure S2: Distributions of ratios q in HuMet and QMDiab**

Distributions of ratios q of PCs of the fatty acid level (Lipidyzer™) and PCs of the lipid species level (Absolute*IDQ*™ p150 kit) are shown separately for subjects of HuMet and subsets of QMDiab. The p-values of Kruskal-Wallis tests indicate if ratios q of HuMet and of male controls aged up to 40 of QMDiab are significantly different. Further p-values refer to comparisons between HuMet, complete QMDiab, and subsets of QMDiab (male controls, female controls, cases).

## References

1. van Meer G., D. R. Voelker, and G. W. Feigenson. 2008. Membrane lipids: where they are and how they behave. Nat. Rev. Mol. Cell Biol. 9: 112–24.

2. Leidl K., G. Liebisch, D. Richter, and G. Schmitz. 2008. Mass spectrometric analysis of lipid species of human circulating blood cells. Biochim. Biophys. Acta - Mol. Cell Biol. Lipids. 1781: 655–64.

3. Cole L. K., J. E. Vance, and D. E. Vance. 2012. Phosphatidylcholine biosynthesis and lipoprotein metabolism. Biochim. Biophys. Acta - Mol. Cell Biol. Lipids. 1821: 754–61.

4. van der Veen J. N., J. P. Kennelly, S. Wan, J. E. Vance, D. E. Vance, and R. L. Jacobs. 2017. The critical role of phosphatidylcholine and phosphatidylethanolamine metabolism in health and disease. Biochim. Biophys. Acta - Biomembr. 1859: 1558–72.

5. Abe M., Y. Sawada, S. Uno, S. Chigasaki, M. Oku, Y. Sakai, and H. Miyoshi. 2017. Role of Acyl Chain Composition of Phosphatidylcholine in Tafazzin-Mediated Remodeling of Cardiolipin in Liposomes. Biochemistry. 56: 6268–80.

6. Nagahama M., A. Otsuka, M. Oda, R. K. Singh, Z. M. Ziora, H. Imagawa, M. Nishizawa, and J. Sakurai. 2007. Effect of unsaturated bonds in the sn-2 acyl chain of phosphatidylcholine on the membrane-damaging action of Clostridium perfringens alpha-toxin toward liposomes. Biochim. Biophys. Acta - Biomembr. 1768: 2940–5.

7. Raclot T., C. Holm, and D. Langin. 2001. Fatty acid specificity of hormone-sensitive lipase: implication in the selective hydrolysis of triacylglycerols. J. Lipid Res. 42: 2049–57.

8. Simopoulos A. P. 2008. The Importance of the Omega-6/Omega-3 Fatty Acid Ratio in Cardiovascular Disease and Other Chronic Diseases. Exp. Biol. Med. 233: 674–88.

9. Whiley L., A. Sen, J. Heaton, P. Proitsi, D. García-Gómez, R. Leung, N. Smith, M. Thambisetty, I. Kloszewska, P. Mecocci, H. Soininen, M. Tsolaki, B. Vellas, S. Lovestone, and C. Legido-Quigley. 2014. Evidence of altered phosphatidylcholine metabolism in Alzheimer’s disease. Neurobiol. Aging. 35: 271–8.

10. Toledo J. B., M. Arnold, G. Kastenmüller, R. Chang, R. A. Baillie, X. Han, M. Thambisetty, J. D. Tenenbaum, K. Suhre, J. W. Thompson, L. St. John-Williams, S. MahmoudianDehkordi, D. M. Rotroff, J. R. Jack, A. Motsinger-Reif, S. L. Risacher, C. Blach, J. E. Lucas, T. Massaro, G. Louie, H. Zhu, G. Dallmann, K. Klavins, T. Koal, S. Kim, K. Nho, L. Shen, R. Casanova, S. Varma, C. Legido-Quigley, M. A. Moseley, K. Zhu, M. Y. R. Henrion, S. J. van der Lee, A. C. Harms, A. Demirkan, T. Hankemeier, C. M. van Duijn, J. Q. Trojanowski, L. M. Shaw, A. J. Saykin, M. W. Weiner, P. M. Doraiswamy, and R. Kaddurah-Daouk. 2017. Metabolic network failures in Alzheimer’s disease: A biochemical road map. Alzheimer’s Dement. 13: 965–84.

11. Floegel A., N. Stefan, Z. Yu, K. Mühlenbruch, D. Drogan, H.-G. Joost, A. Fritsche, H.-U. Häring, M. Hrabě de Angelis, A. Peters, M. Roden, C. Prehn, R. Wang-Sattler, T. Illig, M. B. Schulze, J. Adamski, H. Boeing, and T. Pischon. 2013. Identification of Serum Metabolites Associated With Risk of Type 2 Diabetes Using a Targeted Metabolomic Approach. Diabetes. 62: 639–48.

12. Suhre K., C. Meisinger, A. Döring, E. Altmaier, P. Belcredi, C. Gieger, D. Chang, M. V Milburn, W. E. Gall, K. M. Weinberger, H.-W. Mewes, M. Hrabé de Angelis, H.-E. Wichmann, F. Kronenberg, J. Adamski, and T. Illig. 2010. Metabolic Footprint of Diabetes: A Multiplatform Metabolomics Study in an Epidemiological Setting. PLoS One. 5: e13953.

13. Kulkarni H., M. Mamtani, G. Wong, J. M. Weir, C. K. Barlow, T. D. Dyer, L. Almasy, M. C. Mahaney, A. G. Comuzzie, R. Duggirala, P. J. Meikle, J. Blangero, and J. E. Curran. 2017. Genetic correlation of the plasma lipidome with type 2 diabetes, prediabetes and insulin resistance in Mexican American families. BMC Genet. 18: 48.

14. Meikle P. J., and M. J. Christopher. 2011. Lipidomics is providing new insight into the metabolic syndrome and its sequelae. Curr. Opin. Lipidol. 22: 210–5.

15. Draisma H. H. M., R. Pool, M. Kobl, R. Jansen, A.-K. Petersen, A. A. M. Vaarhorst, I. Yet, T. Haller, A. Demirkan, T. Esko, G. Zhu, S. Böhringer, M. Beekman, J. B. van Klinken, W. Römisch-Margl, C. Prehn, J. Adamski, A. J. M. de Craen, E. M. van Leeuwen, N. Amin, H. Dharuri, H.-J. Westra, L. Franke, E. J. C. de Geus, J. J. Hottenga, G. Willemsen, A. K. Henders, G. W. Montgomery, D. R. Nyholt, J. B. Whitfield, B. W. Penninx, T. D. Spector, A. Metspalu, P. Eline Slagboom, K. W. van Dijk, P. A. C. ‘t Hoen, K. Strauch, N. G. Martin, G.-J. B. van Ommen, T. Illig, J. T. Bell, M. Mangino, K. Suhre, M. I. McCarthy, C. Gieger, A. Isaacs, C. M. van Duijn, and D. I. Boomsma. 2015. Genome-wide association study identifies novel genetic variants contributing to variation in blood metabolite levels. Nat. Commun. 6: 7208.

16. Lario S., M. J. Ramírez-Lázaro, D. Sanjuan-Herráez, A. Brunet-Vega, C. Pericay, L. Gombau, F. Junquera, G. Quintás, and X. Calvet. 2017. Plasma sample based analysis of gastric cancer progression using targeted metabolomics. Sci. Rep. 7: 17774.

17. Liebisch G., J. A. Vizcaíno, H. Köfeler, M. Trötzmüller, W. J. Griffiths, G. Schmitz, F. Spener, and M. J. O. Wakelam. 2013. Shorthand notation for lipid structures derived from mass spectrometry. J. Lipid Res. 54: 1523–30.

18. Fahy E., S. Subramaniam, R. C. Murphy, M. Nishijima, C. R. H. Raetz, T. Shimizu, F. Spener, G. van Meer, M. J. O. Wakelam, and E. A. Dennis. 2009. Update of the LIPID MAPS comprehensive classification system for lipids. J. Lipid Res. 50: S9–14.

19. Siskos A. P., P. Jain, W. Römisch-Margl, M. Bennett, D. Achaintre, Y. Asad, L. Marney, L. Richardson, A. Koulman, J. L. Griffin, F. Raynaud, A. Scalbert, J. Adamski, C. Prehn, and H. C. Keun. 2017. Interlaboratory Reproducibility of a Targeted Metabolomics Platform for Analysis of Human Serum and Plasma. Anal. Chem. 89: 656–65.

20. Yet I., C. Menni, S.-Y. Shin, M. Mangino, N. Soranzo, J. Adamski, K. Suhre, T. D. Spector, G. Kastenmüller, and J. T. Bell. 2016. Genetic Influences on Metabolite Levels: A Comparison across Metabolomic Platforms. PLoS One. 11: e0153672.

21. Wang-Sattler R., Z. Yu, C. Herder, A. C. Messias, A. Floegel, Y. He, K. Heim, M. Campillos, C. Holzapfel, B. Thorand, H. Grallert, T. Xu, E. Bader, C. Huth, K. Mittelstrass, A. Döring, C. Meisinger, C. Gieger, C. Prehn, W. Roemisch-Margl, M. Carstensen, L. Xie, H. Yamanaka-Okumura, G. Xing, U. Ceglarek, J. Thiery, G. Giani, H. Lickert, X. Lin, Y. Li, H. Boeing, H. Joost, M. H. de Angelis, W. Rathmann, K. Suhre, H. Prokisch, A. Peters, T. Meitinger, M. Roden, H. Wichmann, T. Pischon, J. Adamski, and T. Illig. 2012. Novel biomarkers for pre-diabetes identified by metabolomics. Mol. Syst. Biol. 8.

22. Krug S., G. Kastenmüller, F. Stückler, M. J. Rist, T. Skurk, M. Sailer, J. Raffler, W. Römisch-Margl, J. Adamski, C. Prehn, T. Frank, K.-H. Engel, T. Hofmann, B. Luy, R. Zimmermann, F. Moritz, P. Schmitt-Kopplin, J. Krumsiek, W. Kremer, F. Huber, U. Oeh, F. J. Theis, W. Szymczak, H. Hauner, K. Suhre, and H. Daniel. 2012. The dynamic range of the human metabolome revealed by challenges. FASEB J. 26: 2607–19.

23. Mook-Kanamori D. O., M. M. E.-D. Selim, A. H. Takiddin, H. Al-Homsi, K. A. S. Al-Mahmoud, A. Al-Obaidli, M. A. Zirie, J. Rowe, N. A. Yousri, E. D. Karoly, T. Kocher, W. Sekkal Gherbi, O. M. Chidiac, M. J. Mook-Kanamori, S. Abdul Kader, W. A. Al Muftah, C. McKeon, and K. Suhre. 2014. 1,5-Anhydroglucitol in Saliva Is a Noninvasive Marker of Short-Term Glycemic Control. J. Clin. Endocrinol. Metab. 99: E479– 83.

24. Ubhi B. K., A. Conner, E. Duchoslav, A. Evans, R. Robinson, L. Wang, P. R. Baker, and S. Watkins. 2016. A Novel Lipid Screening Platform that Provides a Complete Solution for Lipidomics Research. AB SCIEX Technical Application Note, RUO-MKT-02-2871B.

25. Ubhi B. K. 2016. Novel Chemical Standards Kits Enable Facile Lipid Quantitation. AB SCIEX Technical Application Note, RUO-MKT-02-3879-A.

26. Baker P. R. S., A. M. Armando, J. L. Campbell, O. Quehenberger, and E. A. Dennis. 2014. Three-dimensional enhanced lipidomics analysis combining UPLC, differential ion mobility spectrometry, and mass spectrometric separation strategies. J. Lipid Res. 55: 2432–42.

27. Lintonen T. P. I., P. R. S. Baker, M. Suoniemi, B. K. Ubhi, K. M. Koistinen, E. Duchoslav, J. L. Campbell, and K. Ekroos. 2014. Differential Mobility Spectrometry-Driven Shotgun Lipidomics. Anal. Chem. 86: 9662–9.

28. Ubhi B. K. 2018. Direct infusion-tandem mass spectrometry (DI-MS/MS) analysis of complex lipids in human plasma and serum using the lipidyzer™ platform. In: Methods in Molecular Biology. Humana Press, New York, NY p. 227–36.

29. Bligh E. G., and W. J. Dyer. 1959. A RAPID METHOD OF TOTAL LIPID EXTRACTION AND PURIFICATION. Can. J. Biochem. Physiol. 37: 911–7.

30. LeBlanc Y., D. Caraiman, M. Aiello, and H. Ghobarah. 2015. SelexION Technology: A New Solution to Selectivity Challenges in Quantitative Bioanalysis. AB SCIEX Technical Note, RUO-MKT-02-3251-A.

31. Baker P. R., J. L. Campbell, E. Duchoslav, and C. Hunter. 2017. Differential Mobility Separation for Improving Lipidomic Analysis by Mass Spectrometry. AB SCIEX Technical Application Note, RUO-MKT-02-4802-A

32. Franko A., D. Merkel, M. Kovarova, M. Hoene, B. A. Jaghutriz, M. Heni, A. Königsrainer, C. Papan, S. Lehr, H. U. Häring, and A. Peter. 2018. Dissociation of fatty liver and insulin resistance in I148M PNPLA3 carriers: Differences in diacylglycerol (DAG) FA18:1 lipid species as a possible explanation. Nutrients. 10: 1314.

33. R Core Team. 2017. R: A Language and Environment for Statistical Computing. R Foundation for Statistical Computing. Vienna, Austria.

34. https://www.biocrates.com/images/List-of-Isobaric-and-Isomeric-Lipid-Species_v1_2018.pdf, Accessed: 01.12.2018.

35. Gieger C., L. Geistlinger, E. Altmaier, M. Hrabé de Angelis, F. Kronenberg, T. Meitinger, H.-W. Mewes, H.-E. Wichmann, K. M. Weinberger, J. Adamski, T. Illig, and K. Suhre. 2008. Genetics Meets Metabolomics: A Genome-Wide Association Study of Metabolite Profiles in Human Serum. PLOS Genet. 4: e1000282.

36. Illig T., C. Gieger, G. Zhai, W. Römisch-Margl, R. Wang-Sattler, C. Prehn, E. Altmaier, G. Kastenmüller, B. S. Kato, H.-W. Mewes, T. Meitinger, M. H. de Angelis, F. Kronenberg, N. Soranzo, H.-E. Wichmann, T. D. Spector, J. Adamski, and K. Suhre. 2009. A genome-wide perspective of genetic variation in human metabolism. Nat. Genet. 42: 137.

37. Ried J. S., S.-Y. Shin, J. Krumsiek, T. Illig, F. J. Theis, T. D. Spector, J. Adamski, H.-E. Wichmann, K. Strauch, N. Soranzo, K. Suhre, and C. Gieger. 2014. Novel genetic associations with serum level metabolites identified by phenotype set enrichment analyses. Hum. Mol. Genet. 23: 5847–57.

38. Trabado S., A. Al-Salameh, V. Croixmarie, P. Masson, E. Corruble, B. Fève, R. Colle, L. Ripoll, B. Walther, C. Boursier-Neyret, E. Werner, L. Becquemont, and P. Chanson. 2017. The human plasma-metabolome: Reference values in 800 French healthy volunteers; Impact of cholesterol, gender and age. Motta A, editor. PLoS One. 12: e0173615.

39. Lau C.-H. E., A. P. Siskos, and L. Maitre. 2018. Determinants of the urinary and serum metabolome in children from six European populations. Prep. 16: 202.

40. Zukunft S., C. Prehn, C. Röhring, G. Möller, M. Hrabě de Angelis, J. Adamski, and J. Tokarz. 2018. High-throughput extraction and quantification method for targeted metabolomics in murine tissues. Metabolomics. 14: 18.

41. Koal T., K. Klavins, D. Seppi, G. Kemmler, and C. Humpel. 2015. Sphingomyelin SM(d18:1/18:0) is significantly enhanced in cerebrospinal fluid samples dichotomized by pathological amyloid-β42, tau, and phospho-tau-181 levels. J. Alzheimers. Dis. 44: 1193–201.

42. Hagel J. M., R. Mandal, B. Han, J. Han, D. R. Dinsmore, C. H. Borchers, D. S. Wishart, and P. J. Facchini. 2015. Metabolome analysis of 20 taxonomically related benzylisoquinoline alkaloid-producing plants. BMC Plant Biol. 15: 220.

43. Bouatra S., F. Aziat, R. Mandal, A. C. Guo, M. R. Wilson, C. Knox, T. C. Bjorndahl, R. Krishnamurthy, F. Saleem, P. Liu, Z. T. Dame, J. Poelzer, J. Huynh, F. S. Yallou, N. Psychogios, E. Dong, R. Bogumil, C. Roehring, and D. S. Wishart. 2013. The Human Urine Metabolome. Dzeja P, editor. PLoS One. 8: e73076.

44. Turroni S., J. Fiori, S. Rampelli, S. L. Schnorr, C. Consolandi, M. Barone, E. Biagi, F. Fanelli, M. Mezzullo, A. N. Crittenden, A. G. Henry, P. Brigidi, and M. Candela. 2016. Fecal metabolome of the Hadza hunter-gatherers: A host-microbiome integrative view. Sci. Rep. 6: 32826.

45. Jenkins B., J. A. West, and A. Koulman. 2015. A review of odd-chain fatty acid metabolism and the role of pentadecanoic acid (C15:0) and heptadecanoic acid (C17:0) in health and disease. Molecules.

46. Marai L., and A. Kuksis. 1969. Molecular species of lecithins from erythrocytes and plasma of man. J. Lipid Res. 10: 141–52.

47. Bowden J. A., A. Heckert, C. Z. Ulmer, C. M. Jones, J. P. Koelmel, L. Abdullah, L. Ahonen, Y. Alnouti, A. M. Armando, J. M. Asara, T. Bamba, J. R. Barr, J. Bergquist, C. H. Borchers, J. Brandsma, S. B. Breitkopf, T. Cajka, A. Cazenave-Gassiot, A. Checa, M. A. Cinel, R. A. Colas, S. Cremers, E. A. Dennis, J. E. Evans, A. Fauland, O. Fiehn, M. S. Gardner, T. J. Garrett, K. H. Gotlinger, J. Han, Y. Huang, A. H. Neo, T. Hyötyläinen, Y. Izumi, H. Jiang, H. Jiang, J. Jiang, M. Kachman, R. Kiyonami, K. Klavins, C. Klose, H. C. Köfeler, J. Kolmert, T. Koal, G. Koster, Z. Kuklenyik, I. J. Kurland, M. Leadley, K. Lin, K. R. Maddipati, D. McDougall, P. J. Meikle, N. A. Mellett, C. Monnin, M. A. Moseley, R. Nandakumar, M. Oresic, R. Patterson, D. Peake, J. S. Pierce, M. Post, A. D. Postle, R. Pugh, Y. Qiu, O. Quehenberger, P. Ramrup, J. Rees, B. Rembiesa, D. Reynaud, M. R. Roth, S. Sales, K. Schuhmann, M. L. Schwartzman, C. N. Serhan, A. Shevchenko, S. E. Somerville, L. St. John-Williams, M. A. Surma, H. Takeda, R. Thakare, J. W. Thompson, F. Torta, A. Triebl, M. Trötzmüller, S. J. K. Ubhayasekera, D. Vuckovic, J. M. Weir, R. Welti, M. R. Wenk, C. E. Wheelock, L. Yao, M. Yuan, X. H. Zhao, and S. Zhou. 2017. Harmonizing lipidomics: NIST interlaboratory comparison exercise for lipidomics using SRM 1950–Metabolites in Frozen Human Plasma. J. Lipid Res. 58: 2275–88.

48. Quehenberger O., A. M. Armando, A. H. Brown, S. B. Milne, D. S. Myers, A. H. Merrill, S. Bandyopadhyay, K. N. Jones, S. Kelly, R. L. Shaner, C. M. Sullards, E. Wang, R. C. Murphy, R. M. Barkley, T. J. Leiker, C. R. H. Raetz, Z. Guan, G. M. Laird, D. A. Six, D. W. Russell, J. G. McDonald, S. Subramaniam, E. Fahy, and E. A. Dennis. 2010. Lipidomics reveals a remarkable diversity of lipids in human plasma,. J. Lipid Res. 51: 3299–305.

49. Harayama T., and H. Riezman. 2018. Understanding the diversity of membrane lipid composition. Vol. 19, Nature Reviews Molecular Cell Biology. Nature Publishing Group p. 281–96.

