## Supplemental Figure S1 for "Characterization of bulk phosphatidylcholine compositions in human plasma using side-chain resolving lipidomics"

# PC aa C28:1 = R

No independent Variable measured.

### Qualitative composition

PC aa C28:1 consists of:
PC 10:0_18:1, PC 12:0_16:1, PC 13:0_15:1, PC 14:0_14:1,
[13C1]SM 32:1
and further compounds

# PC aa C30:0 = PC 16:0_14:0 + PC 18:0_12:0 + R

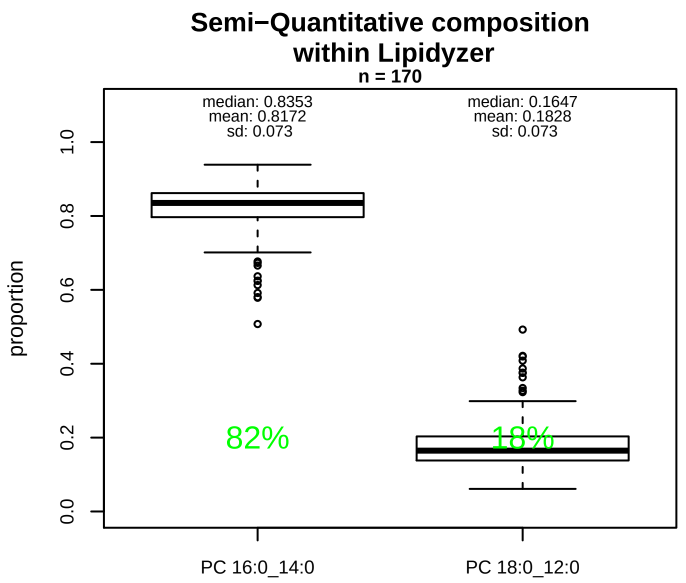


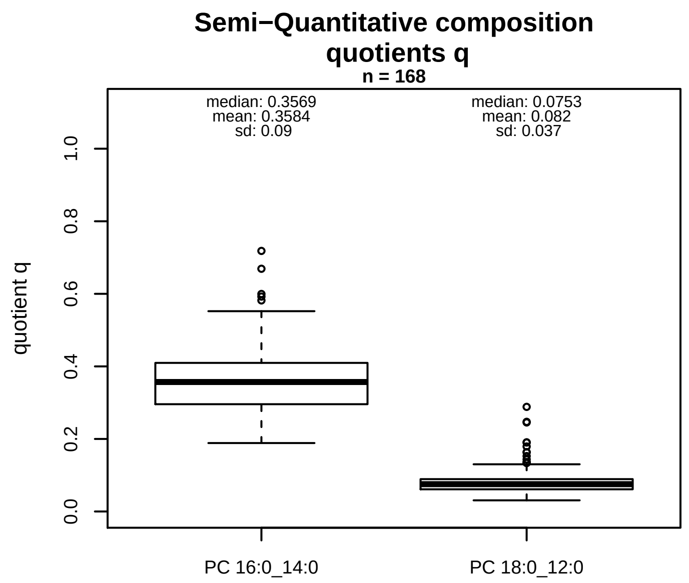


### Semi−Quantitative composition: Lipidyzer

PC aa C30:0 consists of:
PC 16:0_14:0 81.7%
PC 18:0_12:0 18.3%
and of (not quantified):
PC 10:0_20:0, PC 11:0_19:0, PC 13:0_17:0, PC 15:0_15:0,
PC 9:0_21:0, PC O−16:0_15:0, PC O−18:0_13:0, [13C1]SM 3
and further compounds

### Composition: mean of proportions q

conc(PC aa C30:0) * 0.3584 = conc(PC 16:0_14:0) [var(q)=0.3584]
Percentiles: 5%−>0.2248, 25%−>0.2958, 75%−>0.4096, 95%−>0.5098
conc(PC aa C30:0) * 0.082 = conc(PC 18:0_12:0) [var(q)=0.082]
Percentiles: 5%−>0.0462, 25%−>0.0612, 75%−>0.089, 95%−>0.1495

### Linear model

```
PC aa C30:0 ~ b * ( PC 16:0_14:0
   + PC 18:0_12:0 )
```

b = 1.30545

$R^2$ = 0.83347

### Ranges

| Measure | AbsoluteIDQ | sum(Lipidyzer) | delta |
|---|---|---|---|
| Min | 2.17 | 0.6 | 1.57 |
| Max | 7.97 | 4.8 | 3.17 |
| Mean | 3.77 | 1.85 | 1.91 |
| Median | 3.42 | 1.65 | 1.77 |
| SD | 1.25 | 0.89 | 0.36 |

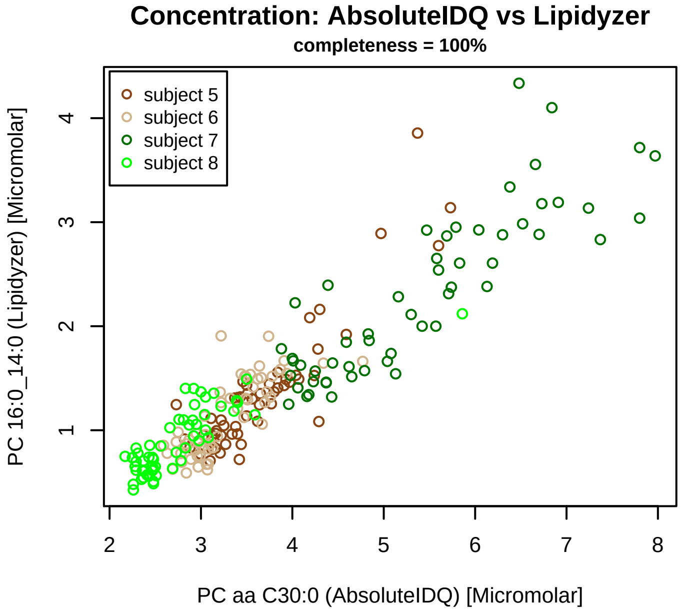


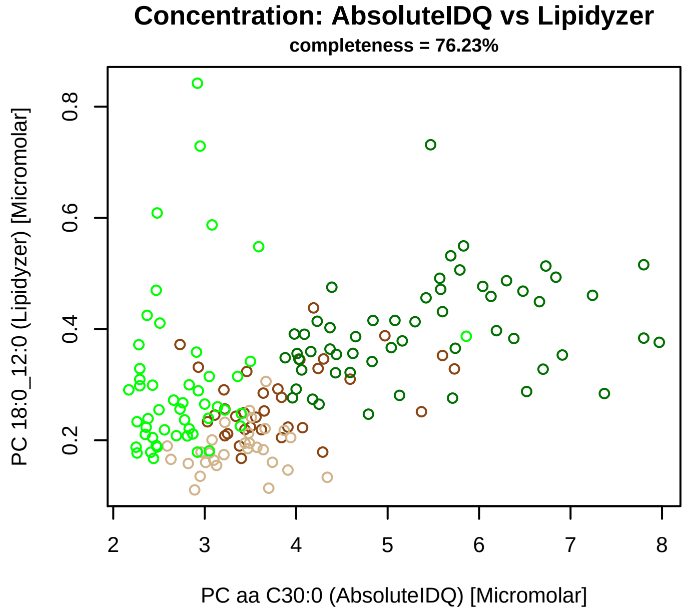


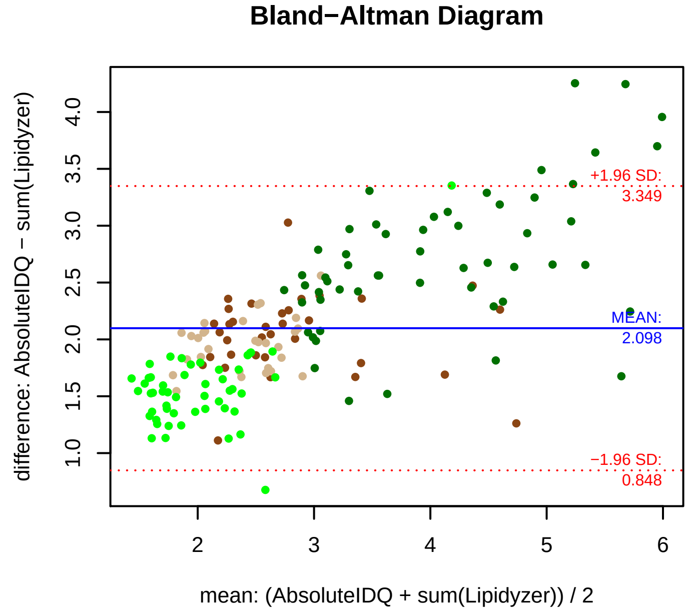


### Stability of composition

**(PC 16:0_14:0 + PC 18:0_12:0) / PC aa C30:0**
**Shapiro−Wilk Test of log quotients, pv: 0.42442; OK**
SW: ok; ANOVA: Comp. ~ Subject_ID −> pv = 0.70763; EQUAL
Wilcoxon: Challenge; fasting: 0.5; sport: 0.125; OLTT: 0.5

**PC 16:0_14:0 / PC aa C30:0**
**Shapiro−Wilk Test of log quotients, pv: 0.69141; OK**
SW: ok; ANOVA: Comp. ~ Subject_ID −> pv = 0.47349; EQUAL
Wilcoxon: Challenge; fasting: 0.125; sport: 0.125; OLTT: 0.625

**PC 18:0_12:0 / PC aa C30:0**
**Shapiro−Wilk Test of log quotients, pv: 3e−05; NO**
SW: no; Kruskal: Comp. ~ Subject_ID −> pv = 0; DIFFERENCE
Wilcoxon: Challenge; fasting: 0.5; sport: 0.125; OLTT: 0.5

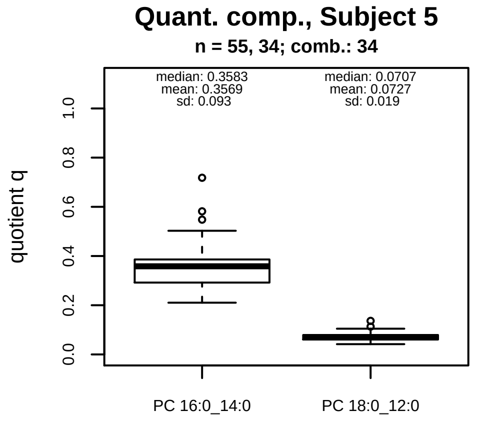


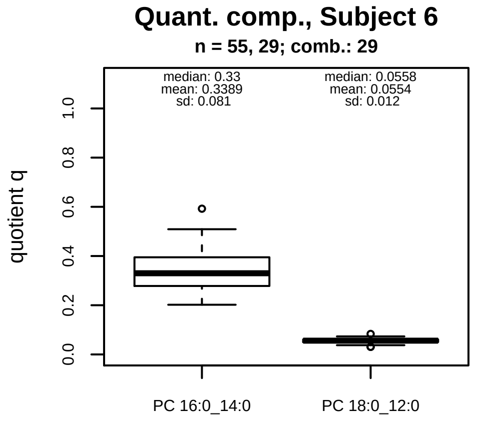


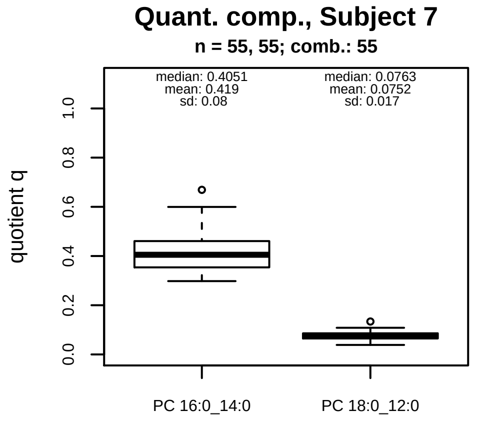


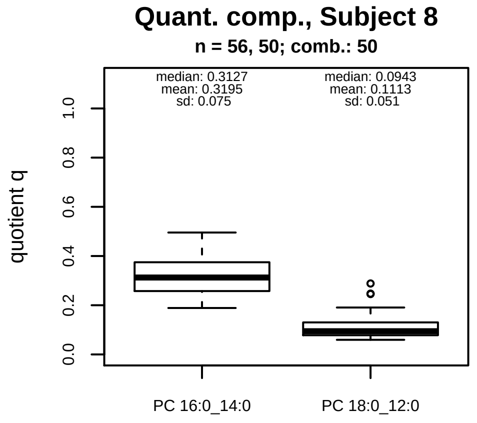


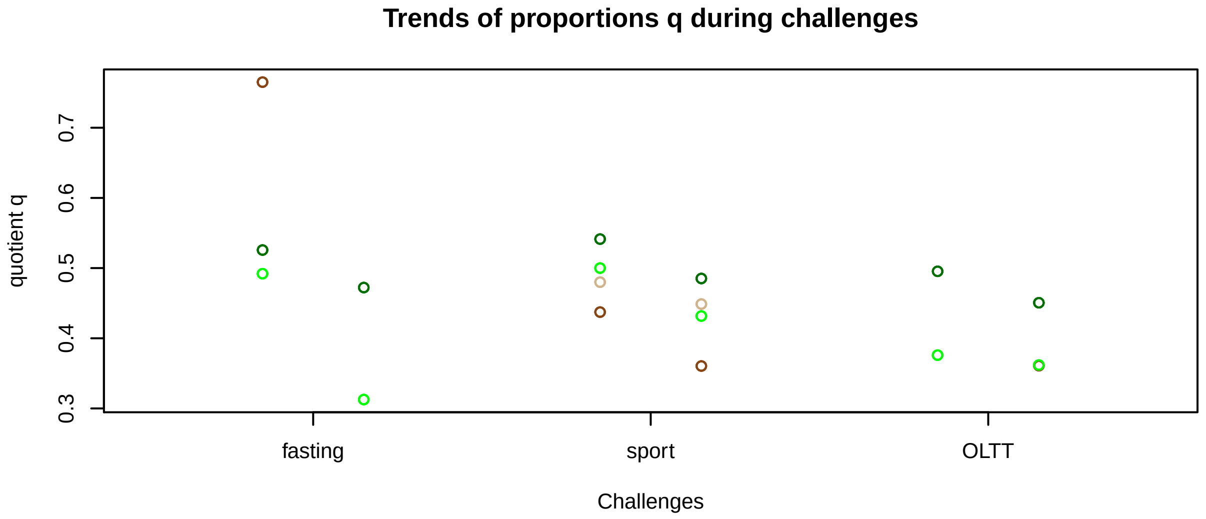


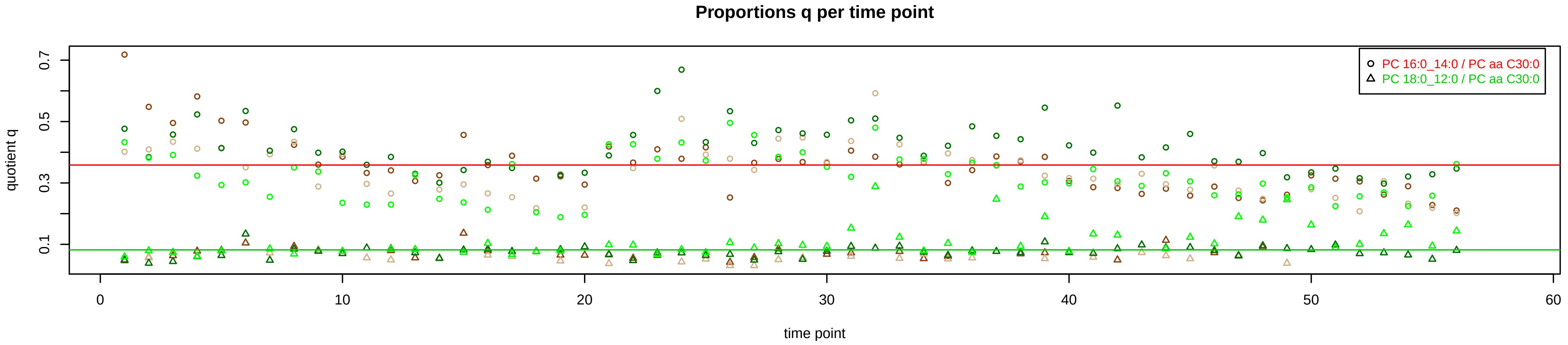

# PC aa C32:0 = PC 16:0_16:0 + PC 18:0_14:0 + R

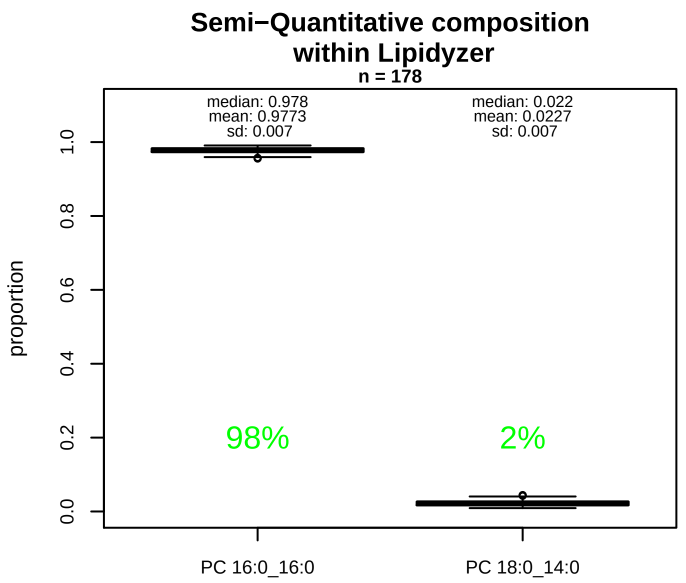


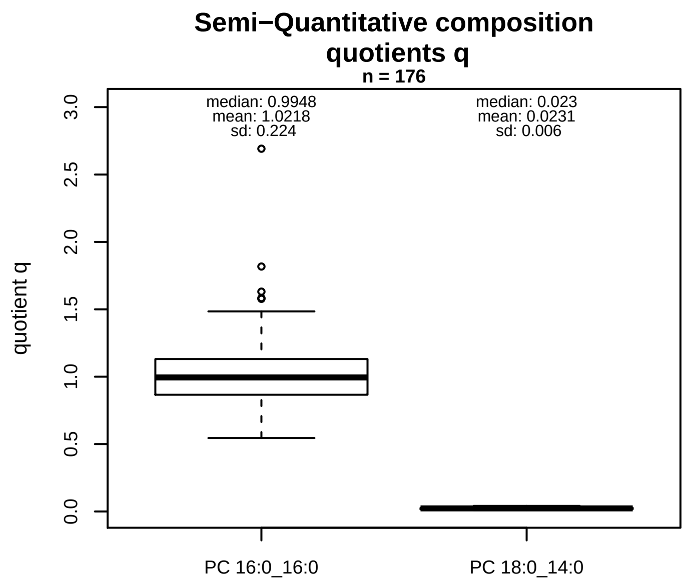


### Semi−Quantitative composition: Lipidyzer

PC aa C32:0 consists of:
PC 16:0_16:0 97.7%
PC 18:0_14:0 2.3%
and of (not quantified):
PC 10:0_22:0, PC 11:0_21:0, PC 12:0_20:0, PC 13:0_19:0,
PC 15:0_17:0, PC 9:0_23:0, PC O−16:0_17:0, PC O−18:0_1[illegible]
[13C1]SM 36:0
and further compounds

### Composition: mean of proportions q

conc(PC aa C32:0) * 1.0218 = conc(PC 16:0_16:0) [var(q)=1.0218]
Percentiles: 5%−>0.7561, 25%−>0.8661, 75%−>1.1308, 95%−>1.3897
conc(PC aa C32:0) * 0.0231 = conc(PC 18:0_14:0) [var(q)=0.0231]
Percentiles: 5%−>0.0138, 25%−>0.0173, 75%−>0.0274, 95%−>0.0347

### Linear model

```
PC aa C32:0 ~ b * ( PC 16:0_16:0
   + PC 18:0_14:0 )
```

b = 0.49186

$R^2$ = 0.35292

### Ranges

| Measure | AbsoluteIDQ | sum(Lipidyzer) | delta |
|---|---|---|---|
| Min | 8.95 | 7.19 | 1.76 |
| Max | 22.05 | 41.83 | 19.78 |
| Mean | 13.18 | 14 | 0.83 |
| Median | 12.22 | 13.74 | 1.52 |
| SD | 3.03 | 3.74 | 0.71 |

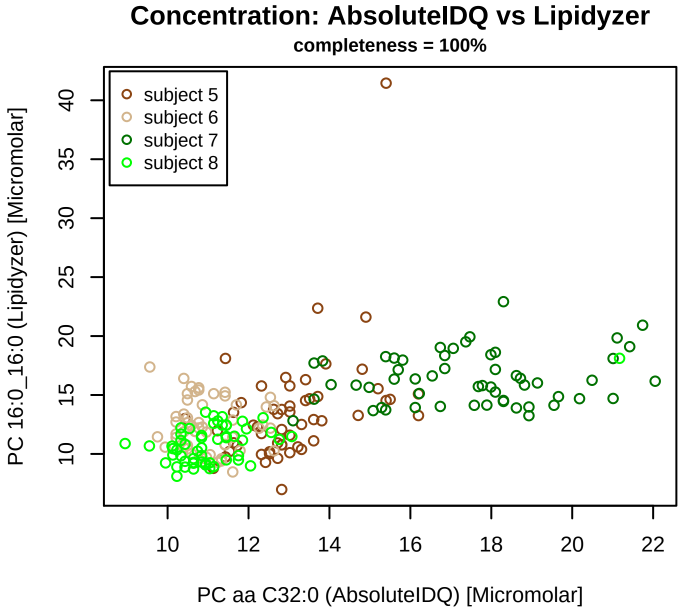


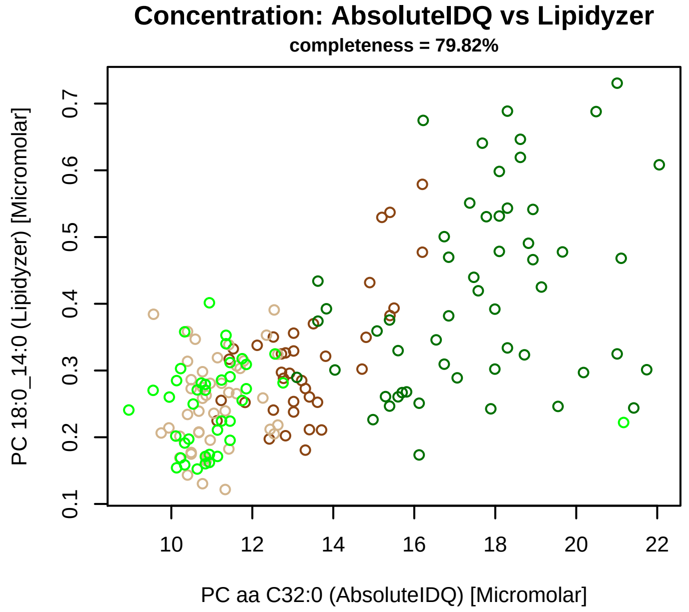


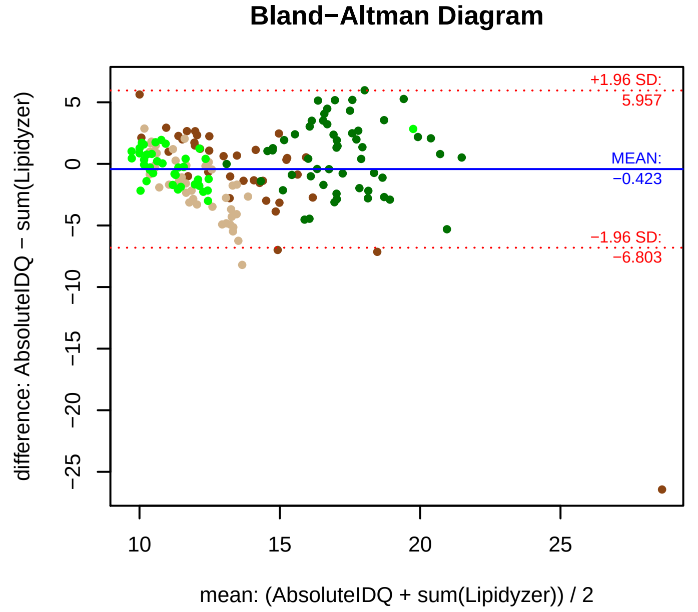


### Stability of composition

**(PC 16:0_16:0 + PC 18:0_14:0) / PC aa C32:0**
**Shapiro−Wilk Test of log quotients, pv: 0.00013; NO**
SW: no; Kruskal: Comp. ~ Subject_ID −> pv = 0.00011; DIFFERENCE
Wilcoxon: Challenge; fasting: 0.75; sport: 0.125; OLTT: 1

**PC 16:0_16:0 / PC aa C32:0**
**Shapiro−Wilk Test of log quotients, pv: 3e−05; NO**
SW: no; Kruskal: Comp. ~ Subject_ID −> pv = 2e−05; DIFFERENCE
Wilcoxon: Challenge; fasting: 0.875; sport: 0.125; OLTT: 0.875

**PC 18:0_14:0 / PC aa C32:0**
**Shapiro−Wilk Test of log quotients, pv: 0.02181; OK**
SW: ok; ANOVA: Comp. ~ Subject_ID −> pv = 0.53573; EQUAL
Wilcoxon: Challenge; fasting: 0.25; sport: 0.625; OLTT: 0.5

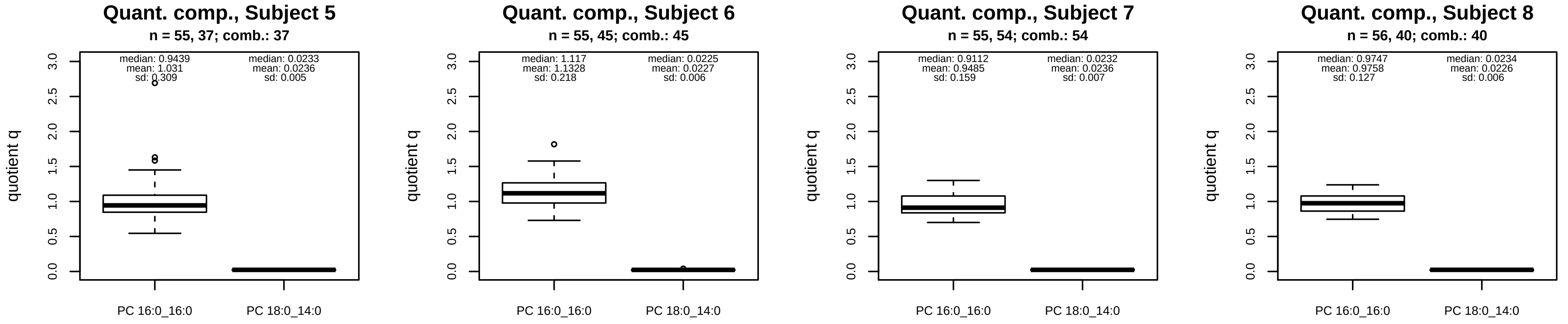


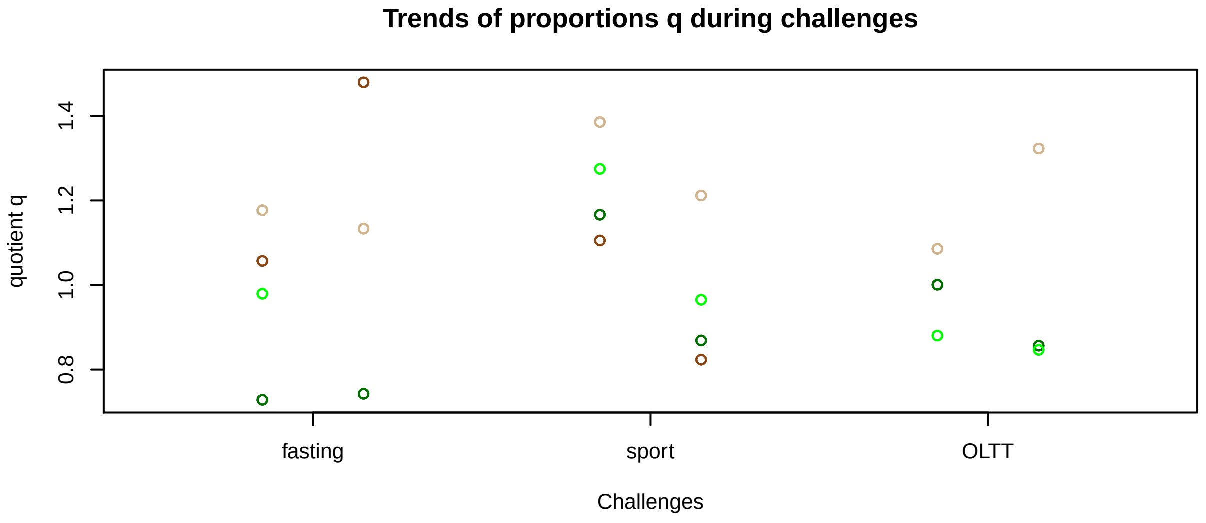


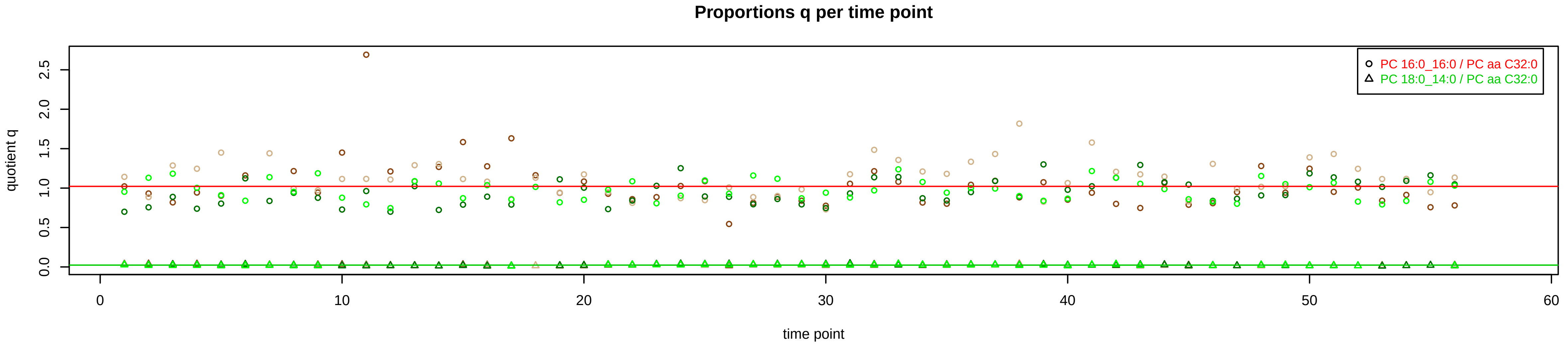

# PC aa C32:1 = PC 14:0_18:1 + PC 16:0_16:1 + R

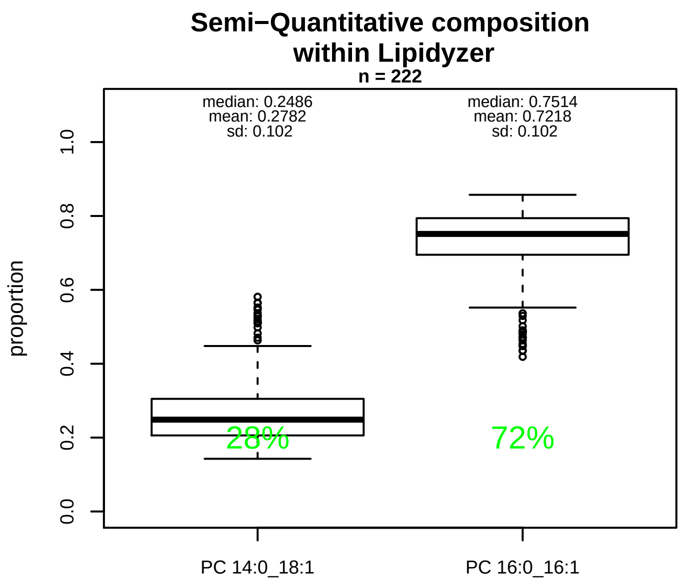


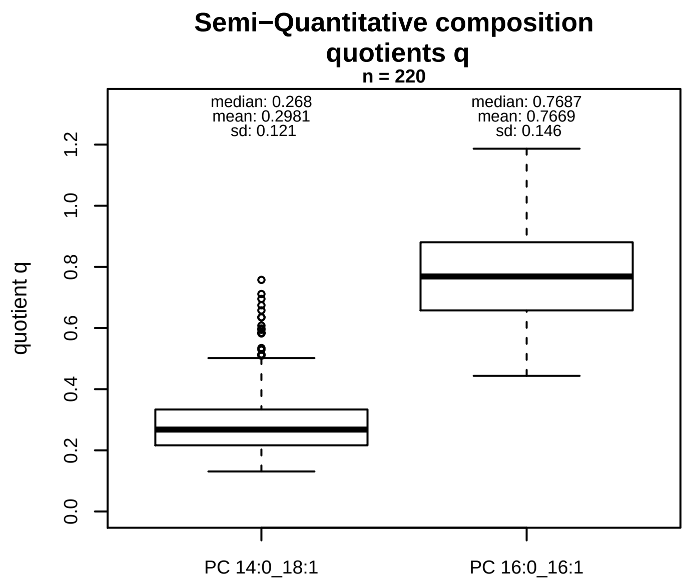


### Semi−Quantitative composition: Lipidyzer

PC aa C32:1 consists of:
PC 14:0_18:1 27.8%
PC 16:0_16:1 72.2%
and of (not quantified):
PC 12:0_20:1, PC 13:0_19:1, PC 14:1_18:0, PC 15:0_17:1,
PC 15:1_17:0, PC O−16:0_17:1, PC O−18:0_15:1, PC O−16:
PC O−20:1_13:0
and further compounds

### Composition: mean of proportions q

conc(PC aa C32:1) * 0.2981 = conc(PC 14:0_18:1) [var(q)=0.2981]
Percentiles: 5%−>0.1773, 25%−>0.2167, 75%−>0.3336, 95%−>0.5843
conc(PC aa C32:1) * 0.7669 = conc(PC 16:0_16:1) [var(q)=0.7669]
Percentiles: 5%−>0.5433, 25%−>0.6577, 75%−>0.8806, 95%−>0.9891

### Linear model

```
PC aa C32:1 ~ b * ( PC 14:0_18:1
   + PC 16:0_16:1 )
```

b = 0.97254

$R^2$ = 0.95086

### Ranges

| Measure | AbsoluteIDQ | sum(Lipidyzer) | delta |
|---|---|---|---|
| Min | 4.79 | 4.01 | 0.78 |
| Max | 38.24 | 41.4 | 3.16 |
| Mean | 13.42 | 14.16 | 0.74 |
| Median | 11.09 | 12.15 | 1.06 |
| SD | 8.15 | 8.14 | 0.01 |

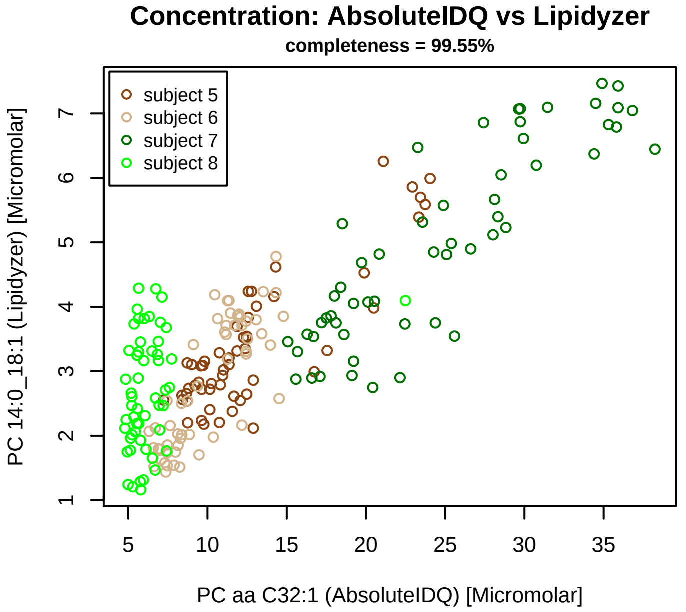


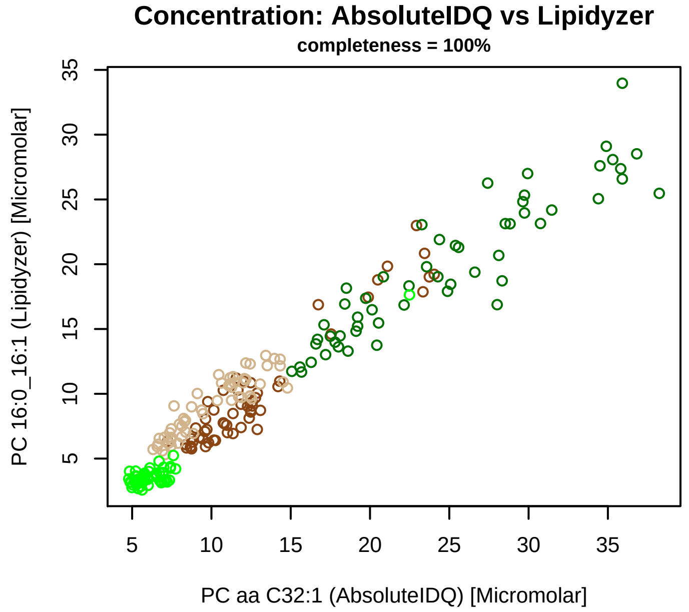


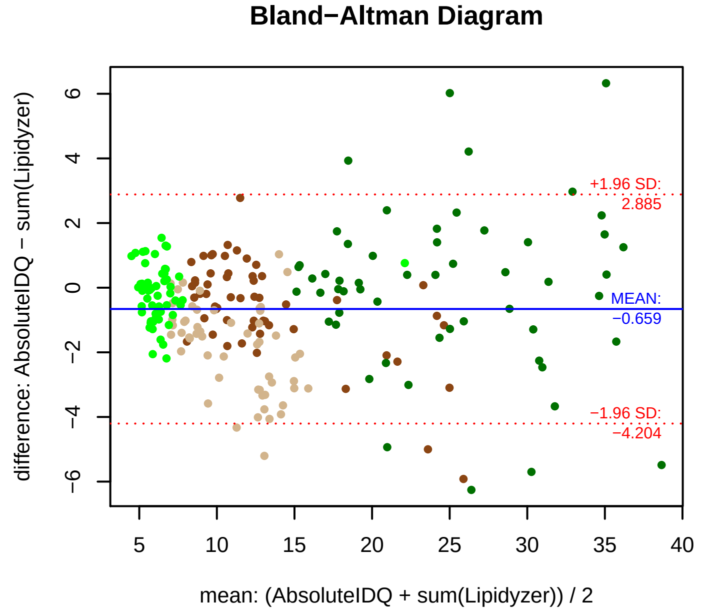


### Stability of composition

**(PC 14:0_18:1 + PC 16:0_16:1) / PC aa C32:1**
**Shapiro−Wilk Test of log quotients, pv: 0.23341; OK**
SW: ok; ANOVA: Comp. ~ Subject_ID −> pv = 0.02667; EQUAL
Wilcoxon: Challenge; fasting: 0.25; sport: 0.625; OLTT: 0.875

**PC 14:0_18:1 / PC aa C32:1**
**Shapiro−Wilk Test of log quotients, pv: 1e−05; NO**
SW: ok; ANOVA: Comp. ~ Subject_ID −> pv = 0; DIFFERENCE
Wilcoxon: Challenge; fasting: 0.125; sport: 0.625; OLTT: 0.25

**PC 16:0_16:1 / PC aa C32:1**
**Shapiro−Wilk Test of log quotients, pv: 0.00051; NO**
SW: ok; ANOVA: Comp. ~ Subject_ID −> pv = 0; DIFFERENCE
Wilcoxon: Challenge; fasting: 1; sport: 0.375; OLTT: 1

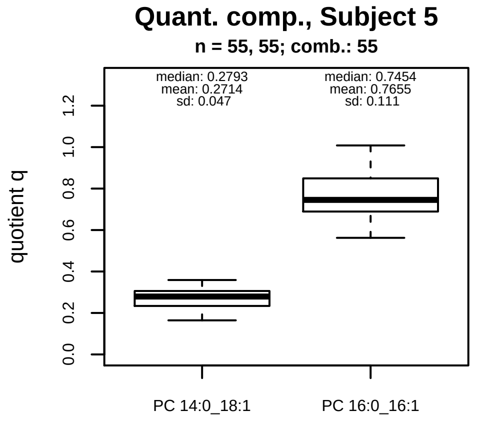


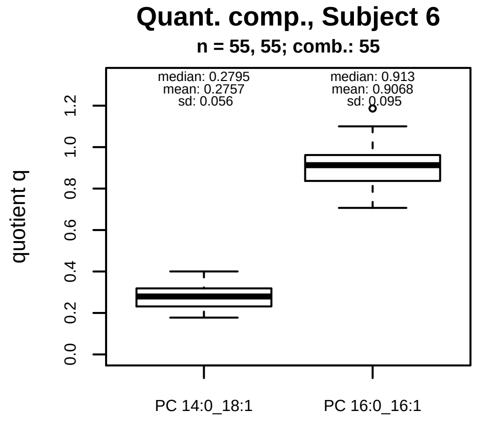


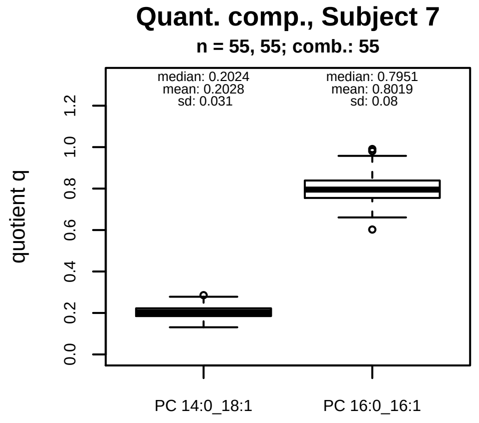


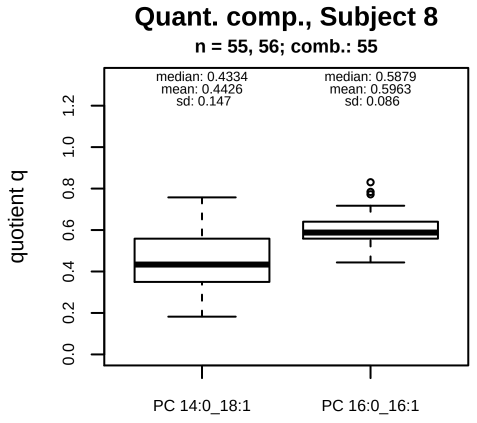


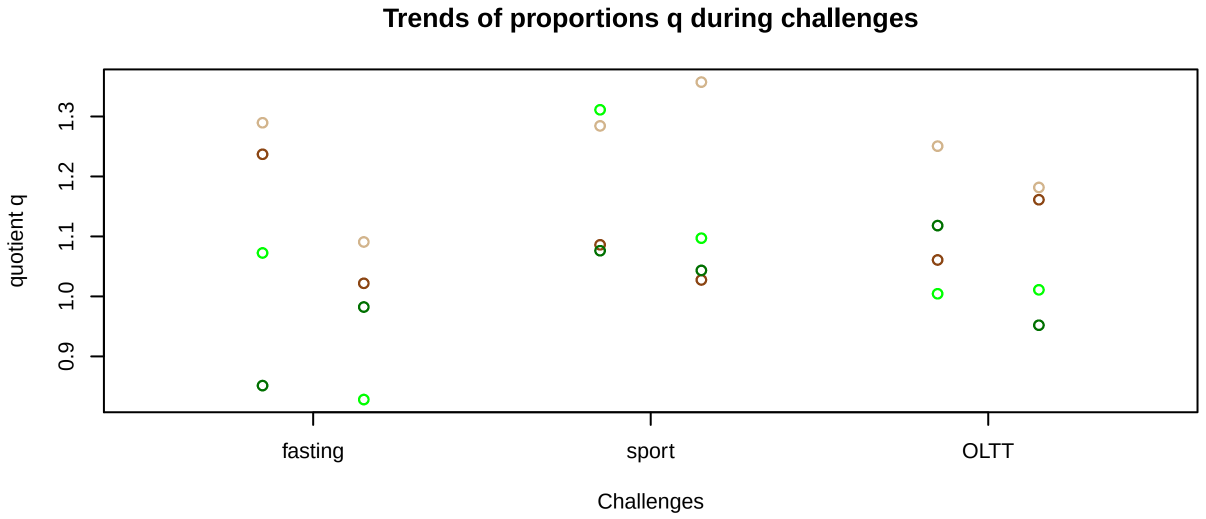


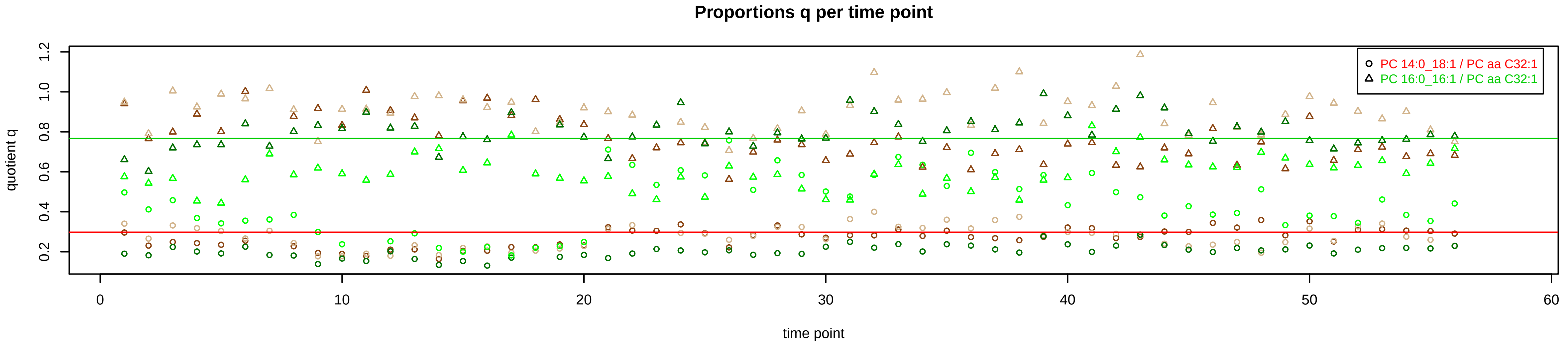

# PC aa C32:2 = PC 14:0_18:2 + R

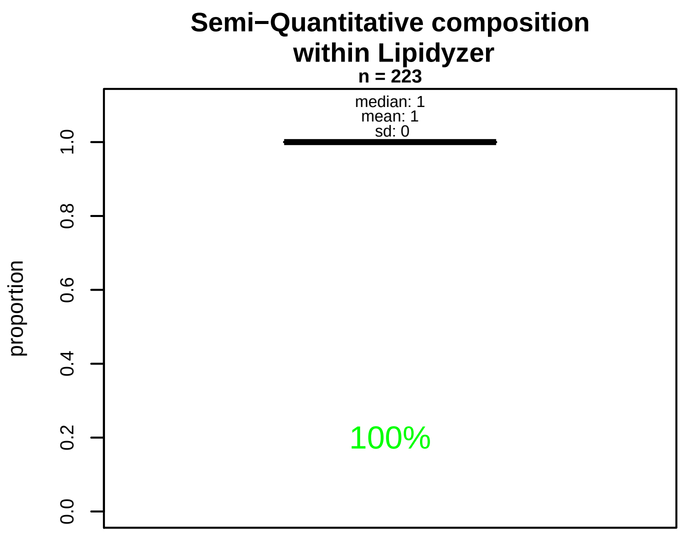


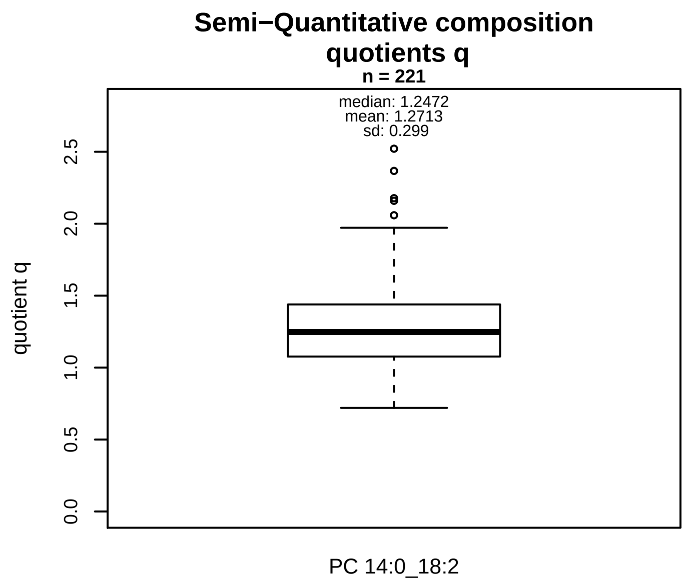


### Semi−Quantitative composition: Lipidyzer

PC aa C32:2 consists of:
PC 14:0_18:2 100%
and of (not quantified):
PC 12:0_20:2, PC 14:1_18:1, PC 15:0_17:2, PC 16:1_16:1, PC 15:1_17:1, PC O−16:0_17:2, PC O−16:1_17:1, PC O−18: and further compounds

### Composition: mean of proportions q

conc(PC aa C32:2) * 1.2713 = conc(PC 14:0_18:2) [var(q)=1.2713]
Percentiles: 5%−>0.8229, 25%−>1.0768, 75%−>1.439, 95%−>1.7371

### Linear model

```
PC aa C32:2 ~ b * ( PC 14:0_18:2 )
```

b = 1.03115

$R^2$ = 0.78893

### Ranges

| Measure | AbsoluteIDQ | sum(Lipidyzer) | delta |
|---|---|---|---|
| Min | 0.66 | 1.23 | 0.56 |
| Max | 5.99 | 6.04 | 0.05 |
| Mean | 2.55 | 2.99 | 0.43 |
| Median | 2.32 | 2.77 | 0.45 |
| SD | 1.27 | 1.09 | 0.18 |

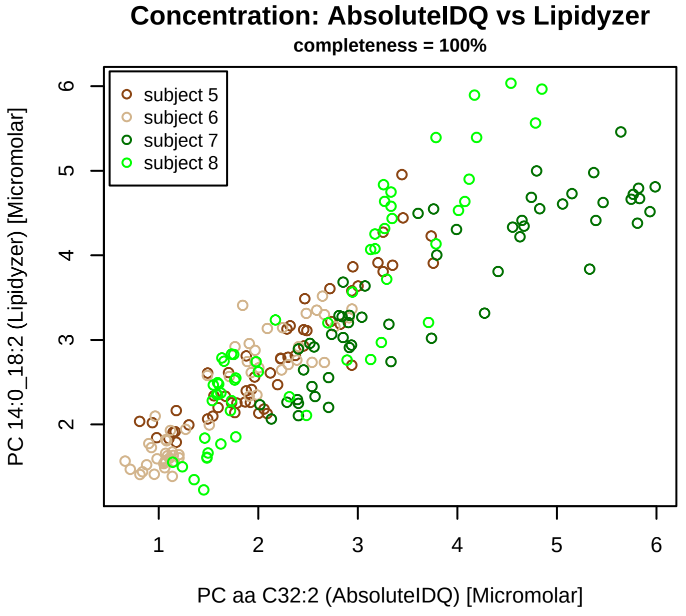


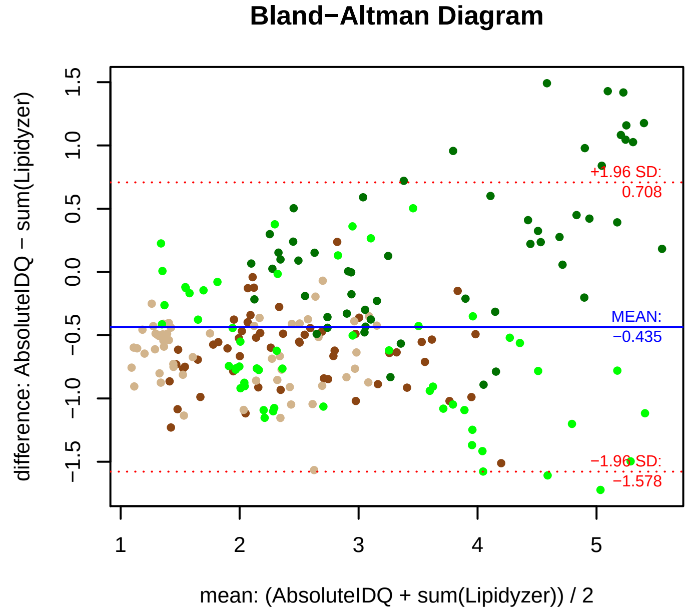


### Stability of composition

**PC 14:0_18:2 / PC aa C32:2**
**Shapiro−Wilk Test of log quotients, pv: 0.28199; OK**
SW: no; Kruskal: Comp. ~ Subject_ID −> pv = 0; DIFFERENCE
Wilcoxon: Challenge; fasting: 0.875; sport: 0.875; OLTT: 0.25

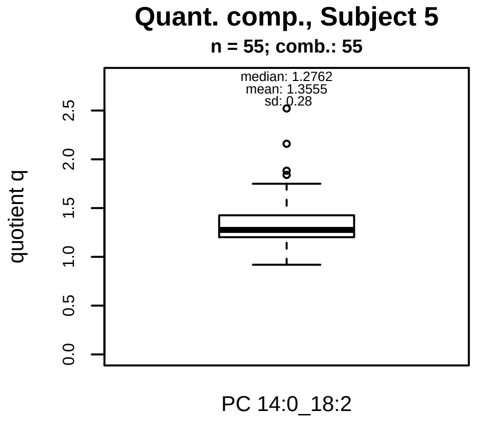


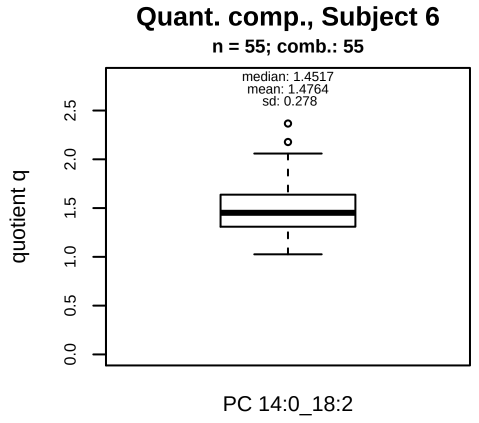


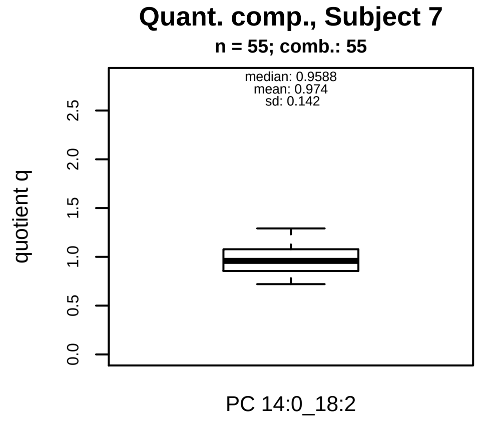


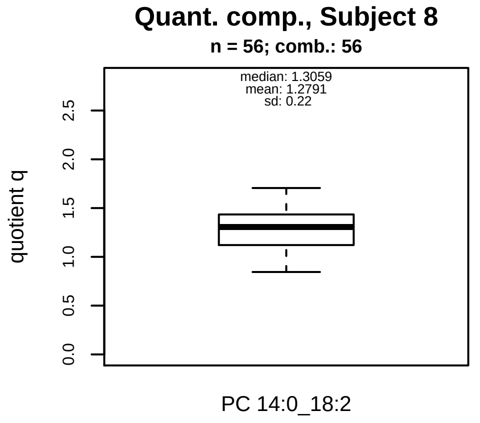


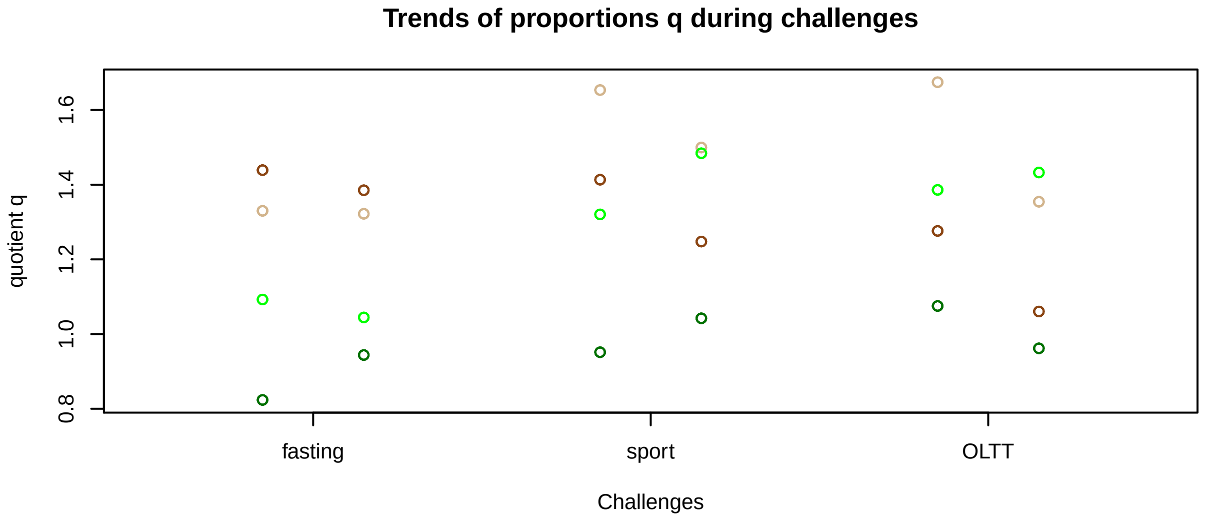


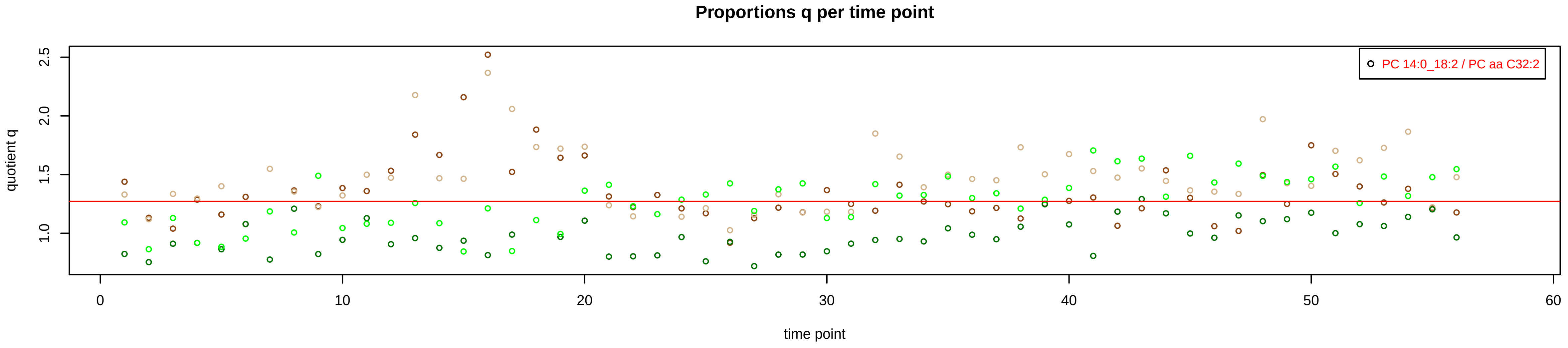

# PC aa C32:3 = PC 14:0_18:3 + R

PC 14:0_18:3 excluded because of missingness > 75%

**No independent Variable with coverage >0.25 out of PC 14:0_18:3**

### Qualitative composition

PC aa C32:3 consists of:
PC 14:0_18:3, PC 12:0_20:3, PC 14:1_18:2, PC 15:1_17:2,
[13C1]SM 36:3
and further compounds

# PC aa C34:1 = PC 14:0_20:1 + PC 16:0_18:1 + PC 18:0_16:1 + PC 20:0_14:1 + R

PC 14:0_20:1 excluded because of missingness > 75%

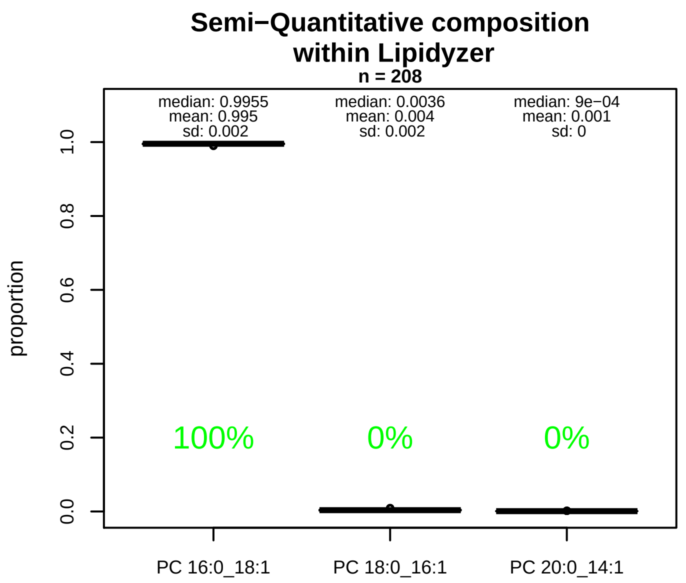


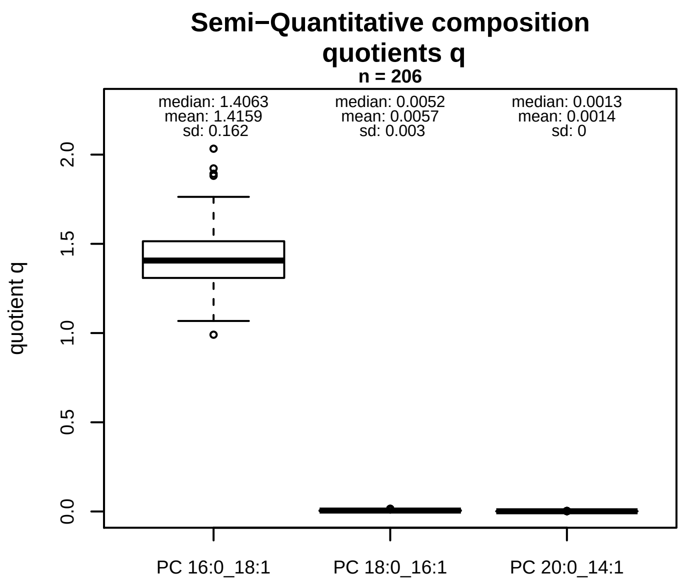


### Semi−Quantitative composition: Lipidyzer

PC aa C34:1 consists of:
PC 16:0_18:1 99.5%
PC 18:0_16:1 0.4%
PC 20:0_14:1 0.1%
and of (not quantified):
PC 14:0_20:1, PC 12:0_22:1, PC 14:1_20:0, PC 15:0_19:1,
PC 15:1_19:0, PC 17:0_17:1, PC O−17:0_18:1, PC O−16:0_1
PC O−18:0_17:1, PC O−20:0_15:1, PC O−16:1_19:0, PC O−
[13C1]SM 38:1
and further compounds

### Linear model

```
PC aa C34:1 ~ b * ( PC 16:0_18:1
   + PC 18:0_16:1
   + PC 20:0_14:1 )
```

b = 0.55441

$R^2$ = 0.86597

### Ranges

| Measure | AbsoluteIDQ | sum(Lipidyzer) | delta |
|---|---|---|---|
| Min | 89.8 | 127.65 | 37.85 |
| Max | 325.5 | 471.26 | 145.76 |
| Mean | 187.14 | 268.99 | 81.85 |
| Median | 183.6 | 261.66 | 78.06 |
| SD | 49.04 | 84.03 | 35 |

### Composition: mean of proportions q

conc(PC aa C34:1) * 1.4159 = conc(PC 16:0_18:1) [var(q)=1.4159]
Percentiles: 5%−>1.1793, 25%−>1.3089, 75%−>1.5146, 95%−>1.6821
conc(PC aa C34:1) * 0.0057 = conc(PC 18:0_16:1) [var(q)=0.0057]
Percentiles: 5%−>0.0028, 25%−>0.0036, 75%−>0.0067, 95%−>0.012
conc(PC aa C34:1) * 0.0014 = conc(PC 20:0_14:1) [var(q)=0.0014]
Percentiles: 5%−>9e−04, 25%−>0.0011, 75%−>0.0016, 95%−>0.0022

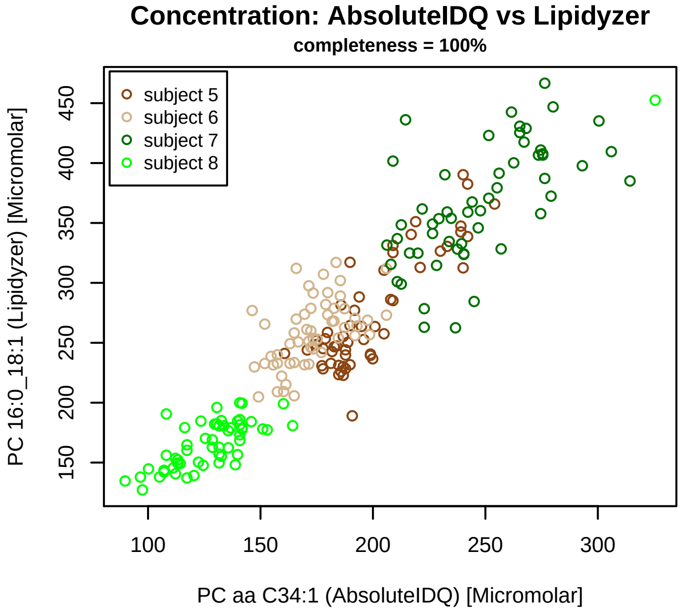


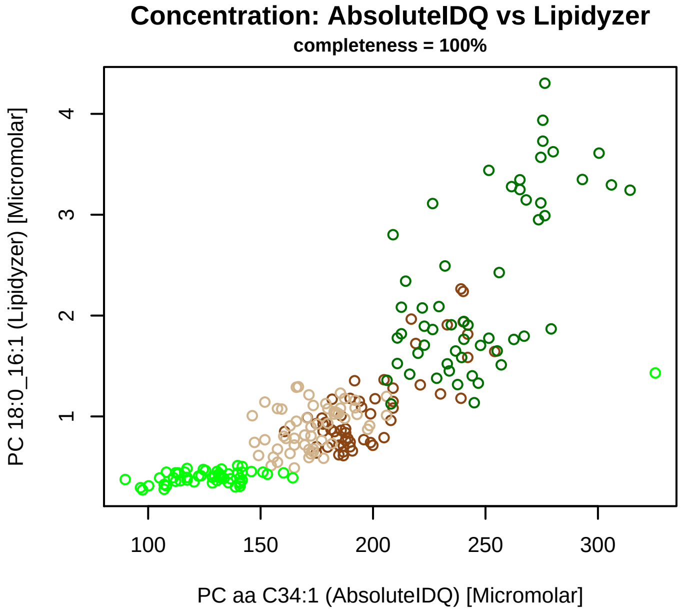


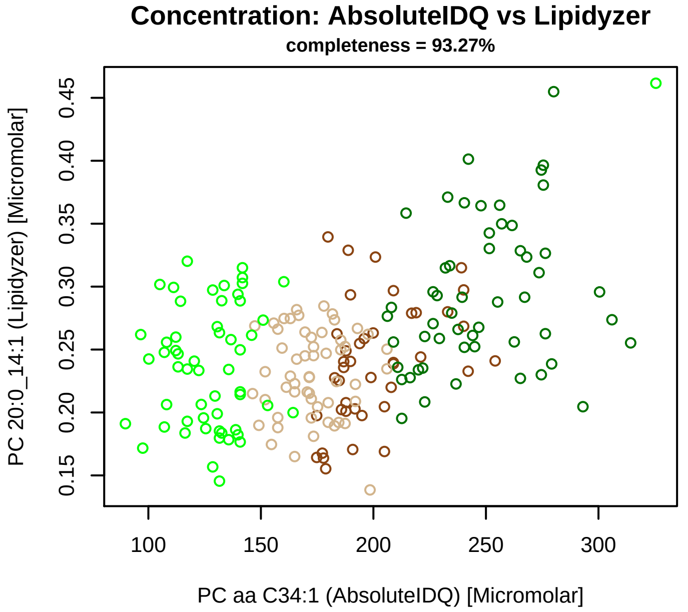


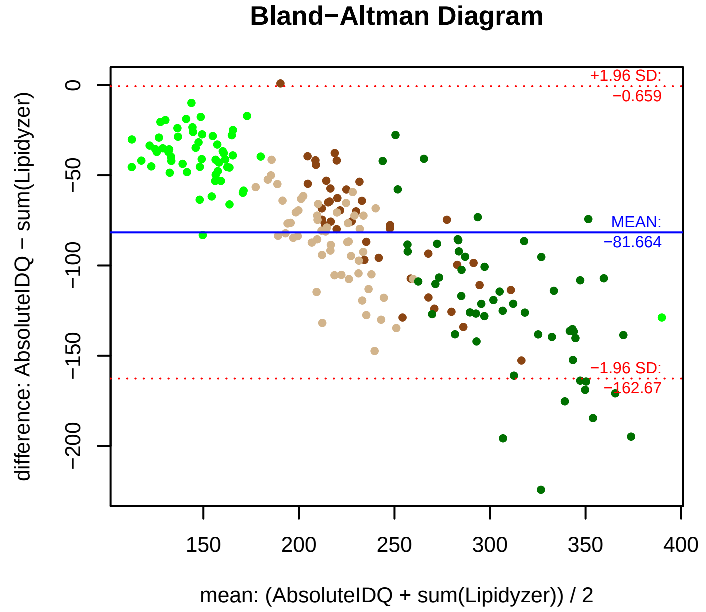


### Stability of composition

**(PC 16:0_18:1 + PC 18:0_16:1 + PC 20:0_14:1) / PC aa C34:1**
**Shapiro−Wilk Test of log quotients, pv: 0.82819; OK**
SW: ok; ANOVA: Comp. ~ Subject_ID −> pv = 0.00213; DIFFERENCE
Wilcoxon: Challenge; fasting: 0.75; sport: 0.125; OLTT: 1

**PC 16:0_18:1 / PC aa C34:1**
**Shapiro−Wilk Test of log quotients, pv: 0.85563; OK**
SW: ok; ANOVA: Comp. ~ Subject_ID −> pv = 0.02562; EQUAL
Wilcoxon: Challenge; fasting: 0.625; sport: 0.125; OLTT: 1

**PC 18:0_16:1 / PC aa C34:1**
**Shapiro−Wilk Test of log quotients, pv: 0.00069; NO**
SW: ok; ANOVA: Comp. ~ Subject_ID −> pv = 0.00029; DIFFERENCE
Wilcoxon: Challenge; fasting: 0.125; sport: 0.25; OLTT: 0.125

**PC 20:0_14:1 / PC aa C34:1**
**Shapiro−Wilk Test of log quotients, pv: 0.01446; OK**
SW: ok; ANOVA: Comp. ~ Subject_ID −> pv = 0; DIFFERENCE
Wilcoxon: Challenge; fasting: 0.75; sport: 0.625; OLTT: 1

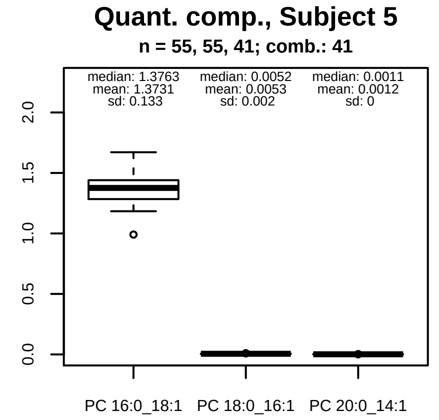


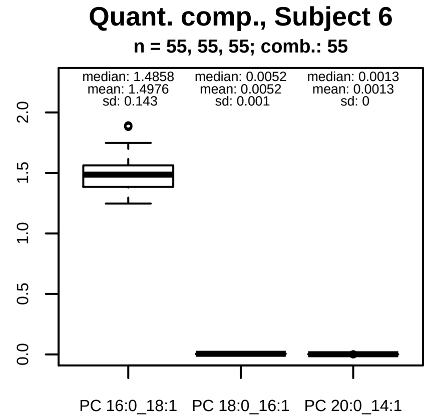


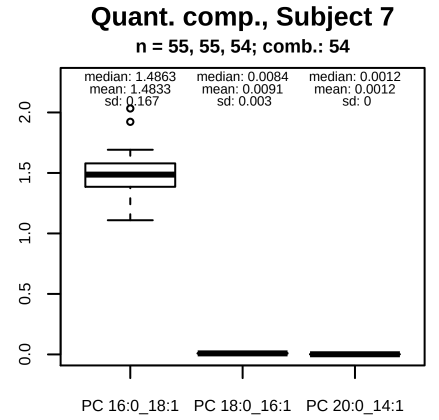


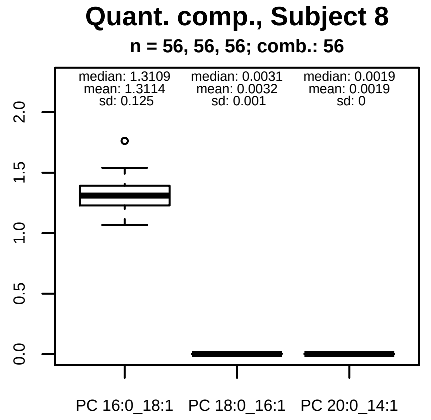


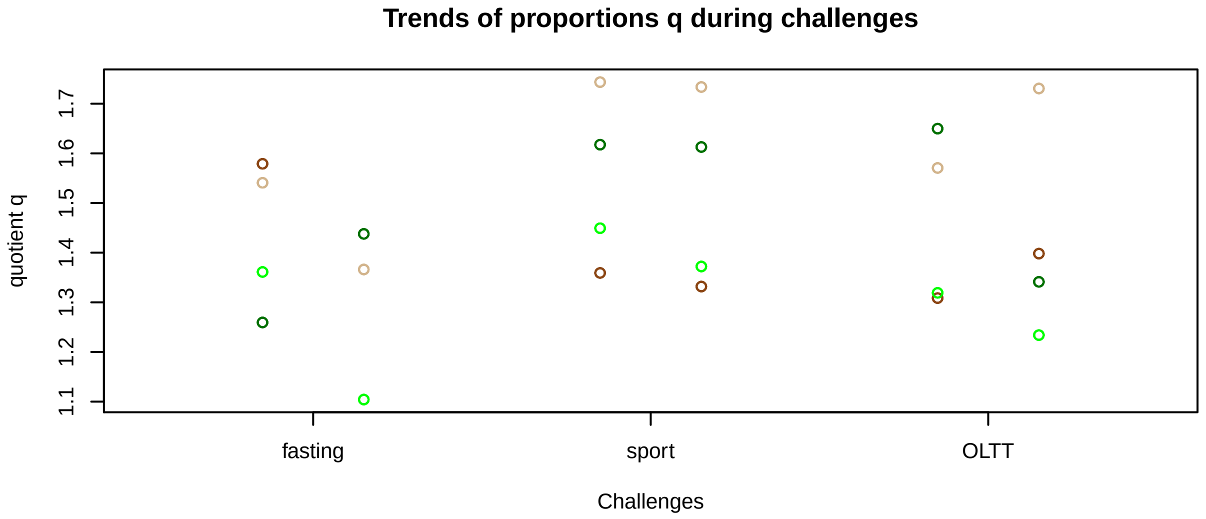


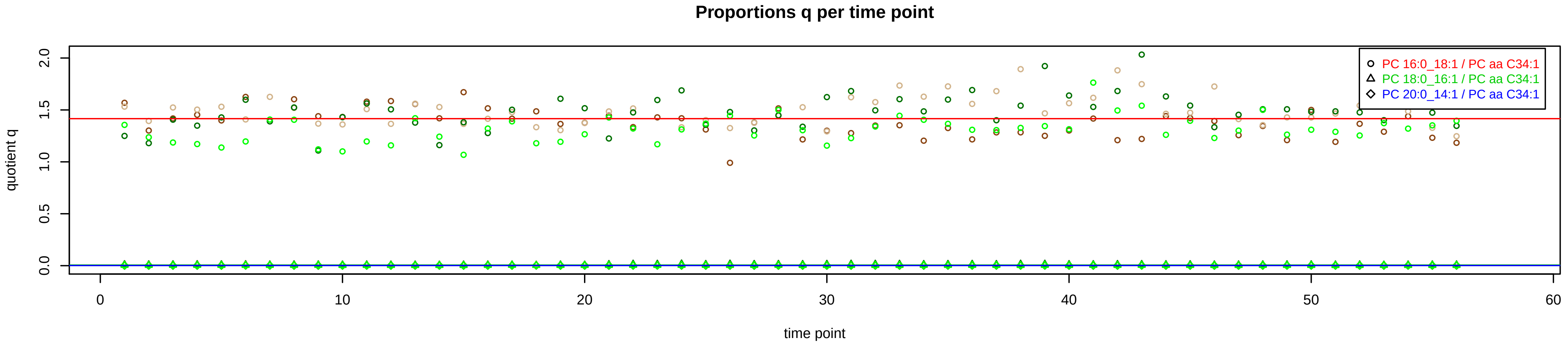

# PC aa C34:2 = PC 14:0_20:2 + PC 16:0_18:2 + PC 18:1_16:1 + R

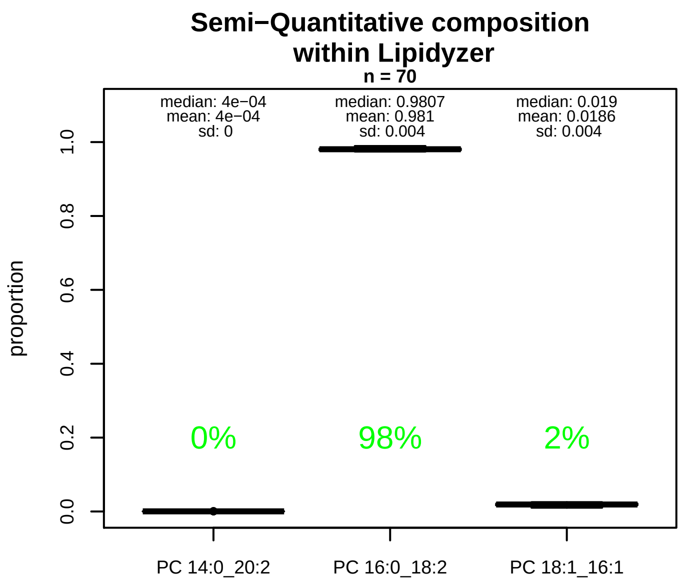


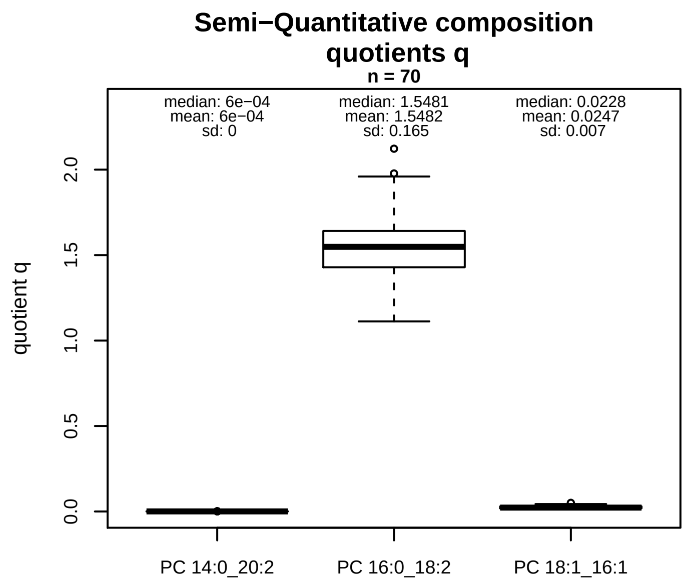


### Semi−Quantitative composition: Lipidyzer

PC aa C34:2 consists of:
- PC 14:0_20:2 <0.05%
- PC 16:0_18:2 98.1%
- PC 18:1_16:1 1.9%

and of (not quantified):
PC 12:0_22:2, PC 14:1_20:1, PC 15:1_19:1, PC 17:0_17:2, PC 17:1_17:1, PC 18:0_16:2, PC O−18:0_17:2, PC O−16:1_
[13C1]SM 38:2
and further compounds

### Composition: mean of proportions q

conc(PC aa C34:2) * 6e−04 = conc(PC 14:0_20:2) [var(q)=6e−04]
Percentiles: 5%−>4e−04, 25%−>5e−04, 75%−>7e−04, 95%−>0.001
conc(PC aa C34:2) * 1.5482 = conc(PC 16:0_18:2) [var(q)=1.5482]
Percentiles: 5%−>1.2968, 25%−>1.4291, 75%−>1.6414, 95%−>1.8214
conc(PC aa C34:2) * 0.0247 = conc(PC 18:1_16:1) [var(q)=0.0247]
Percentiles: 5%−>0.0155, 25%−>0.0188, 75%−>0.0299, 95%−>0.0371

### Linear model

```
PC aa C34:2 ~ b * ( PC 14:0_20:2
  + PC 16:0_18:2
  + PC 18:1_16:1 )
```

b = 0.39293

$R^2$ = 0.49433

### Ranges

| Measure | AbsoluteIDQ | sum(Lipidyzer) | delta |
|---|---|---|---|
| Min | 233.6 | 349.63 | 116.03 |
| Max | 465 | 708.04 | 243.04 |
| Mean | 320.57 | 543.8 | 223.23 |
| Median | 315.6 | 551.65 | 236.05 |
| SD | 40.33 | 79.51 | 39.18 |

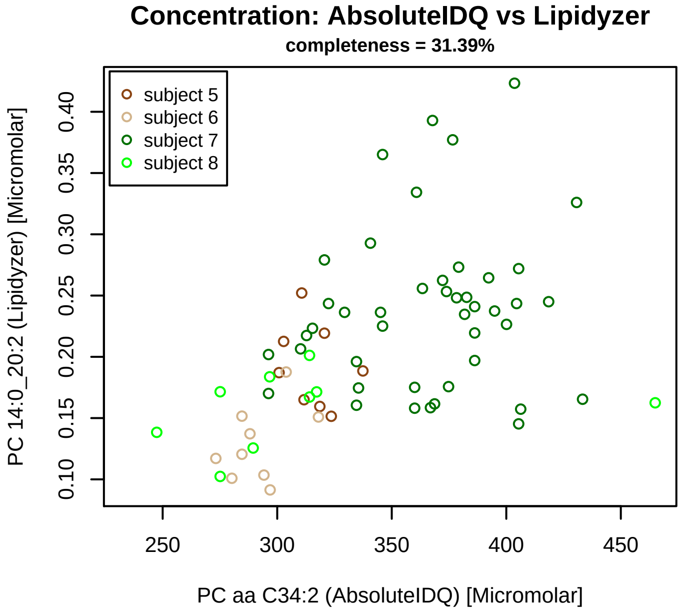


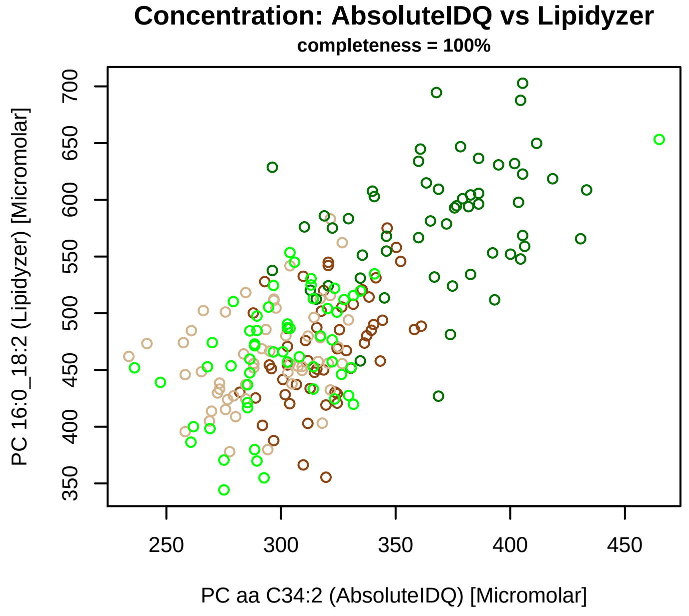


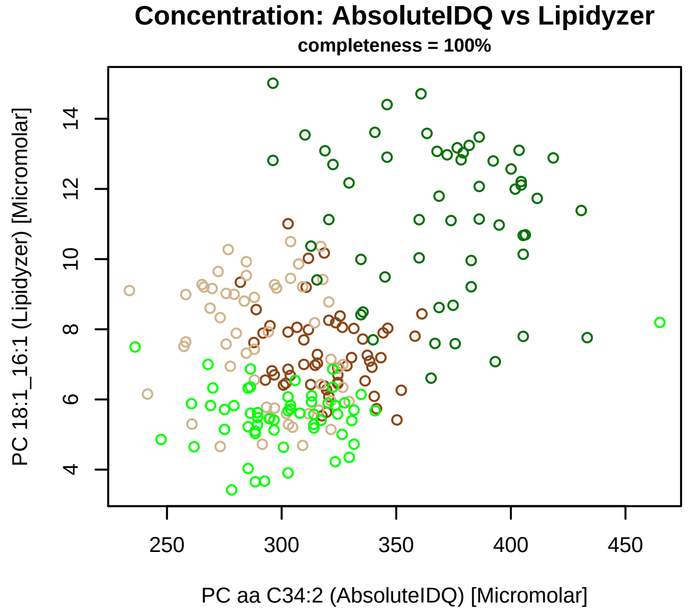


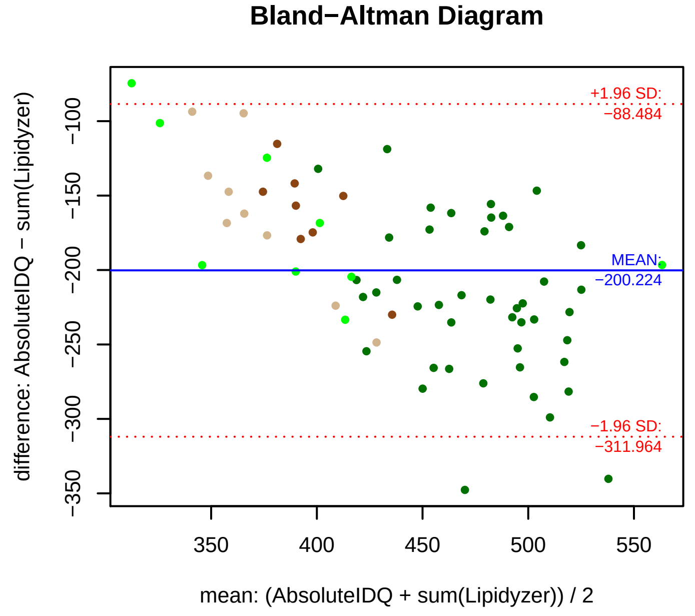


### Stability of composition

**(PC 14:0_20:2 + PC 16:0_18:2 + PC 18:1_16:1) / PC aa C34:2**
**Shapiro−Wilk Test of log quotients, pv: 0.55528; OK**
SW: ok; ANOVA: Comp. ~ Subject_ID −> pv = 0.38377; EQUAL
Wilcoxon: Challenge; fasting: 1; sport: 1; OLTT: 1

**PC 14:0_20:2 / PC aa C34:2**
**Shapiro−Wilk Test of log quotients, pv: 0.38507; OK**
SW: ok; ANOVA: Comp. ~ Subject_ID −> pv = 0.61799; EQUAL
Wilcoxon: Challenge; fasting: 1; sport: 1; OLTT: 1

**PC 16:0_18:2 / PC aa C34:2**
**Shapiro−Wilk Test of log quotients, pv: 0.95815; OK**
SW: ok; ANOVA: Comp. ~ Subject_ID −> pv = 0.021; EQUAL
Wilcoxon: Challenge; fasting: 0.375; sport: 0.875; OLTT: 0.625

**PC 18:1_16:1 / PC aa C34:2**
**Shapiro−Wilk Test of log quotients, pv: 0.00691; NO**
SW: no; Kruskal: Comp. ~ Subject_ID −> pv = 0; DIFFERENCE
Wilcoxon: Challenge; fasting: 0.25; sport: 0.375; OLTT: 0.125

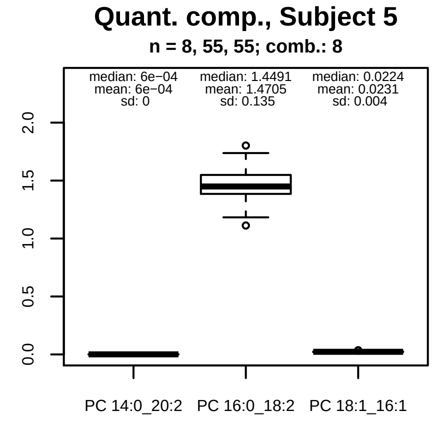


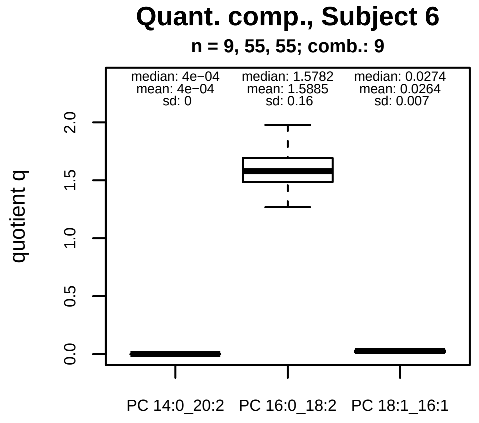


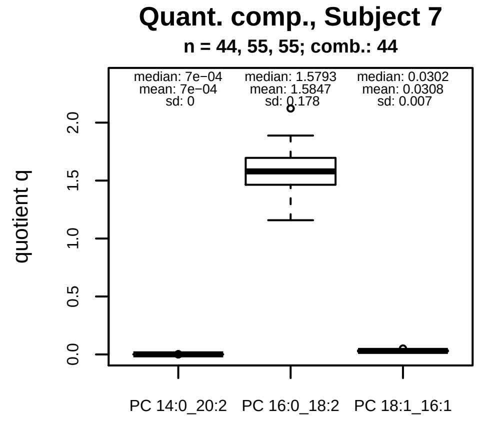


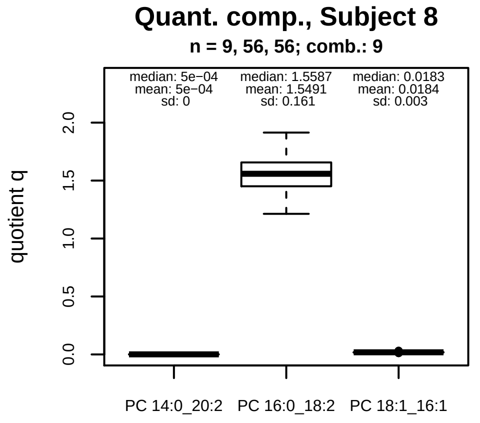


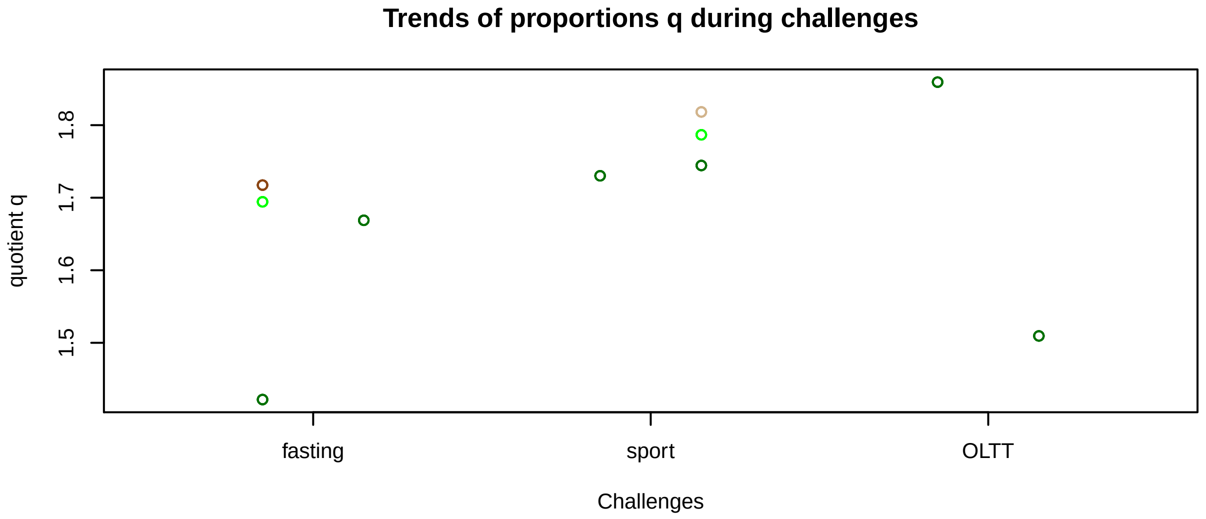


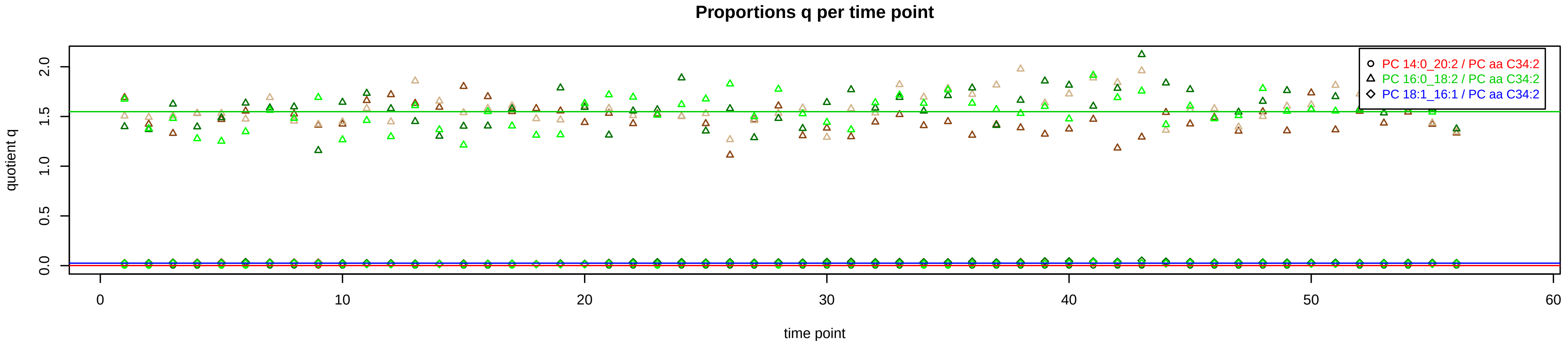

# PC aa C34:3 = PC 14:0_20:3 + PC 16:0_18:3 + PC 18:2_16:1 + R

#### Semi−Quantitative composition within Lipidyzer

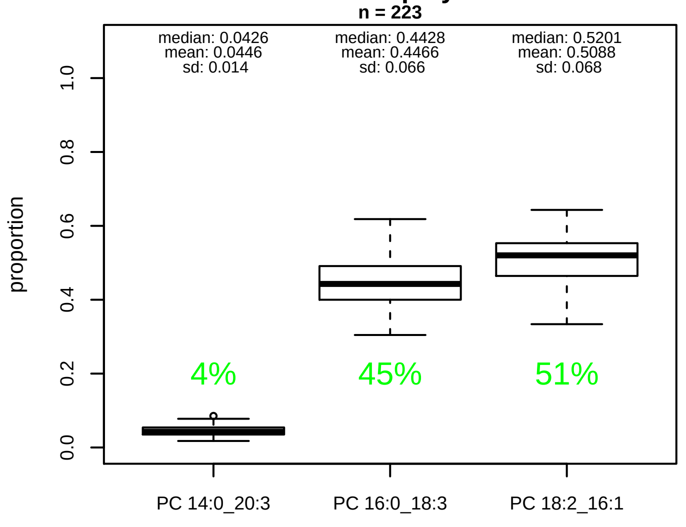


#### Semi−Quantitative composition quotients q

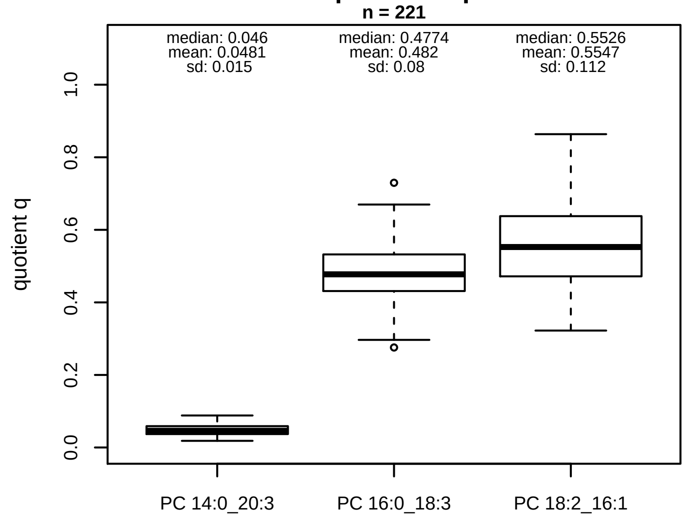


#### Semi−Quantitative composition: Lipidyzer

PC aa C34:3 consists of:
PC 14:0_20:3 4.5%
PC 16:0_18:3 44.7%
PC 18:2_16:1 50.9%
and of (not quantified):
PC 14:1_20:2, PC 16:1_18:2, PC 17:1_17:2, PC O−18:1_17:2
and further compounds

#### Linear model

```
PC aa C34:3 ~ b * ( PC 14:0_20:3
   + PC 16:0_18:3
   + PC 18:2_16:1 )
```

b = 1.0004

$R^2$ = 0.85431

#### Ranges

| Measure | AbsoluteIDQ | sum(Lipidyzer) | delta |
|---|---|---|---|
| Min | 8.23 | 8.34 | 0.11 |
| Max | 29 | 29.94 | 0.94 |
| Mean | 15.67 | 16.72 | 1.05 |
| Median | 13.98 | 15.66 | 1.68 |
| SD | 4.82 | 4.43 | 0.39 |

#### Composition: mean of proportions q

conc(PC aa C34:3) * 0.0481 = conc(PC 14:0_20:3) [var(q)=0.0481]
Percentiles: 5%−>0.0252, 25%−>0.0369, 75%−>0.0588, 95%−>0.0729
conc(PC aa C34:3) * 0.482 = conc(PC 16:0_18:3) [var(q)=0.482]
Percentiles: 5%−>0.3539, 25%−>0.4312, 75%−>0.5322, 95%−>0.6165
conc(PC aa C34:3) * 0.5547 = conc(PC 18:2_16:1) [var(q)=0.5547]
Percentiles: 5%−>0.3811, 25%−>0.4718, 75%−>0.6377, 95%−>0.7406

#### Concentration: AbsoluteIDQ vs Lipidyzer

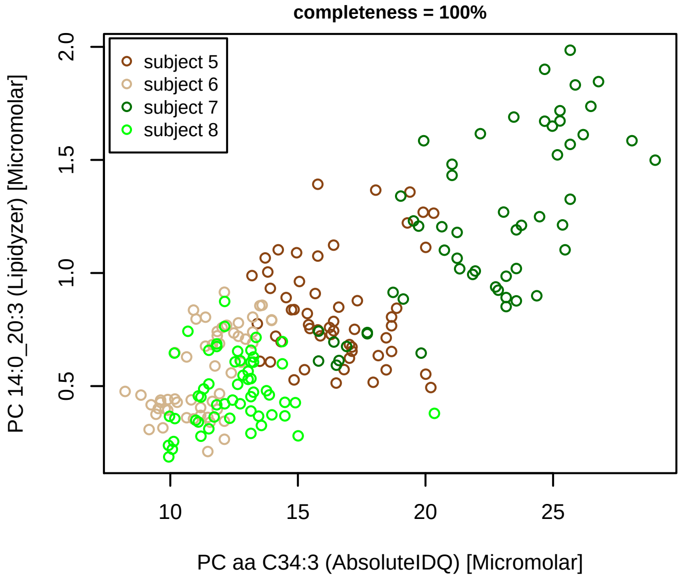


#### Concentration: AbsoluteIDQ vs Lipidyzer

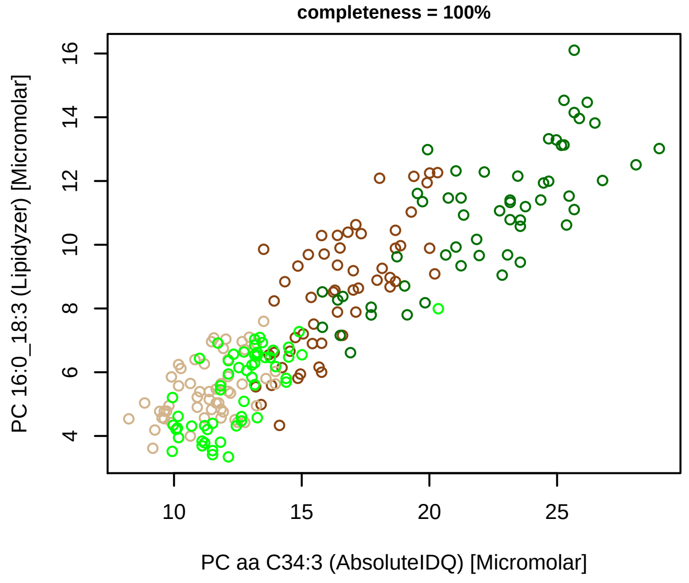


#### Concentration: AbsoluteIDQ vs Lipidyzer

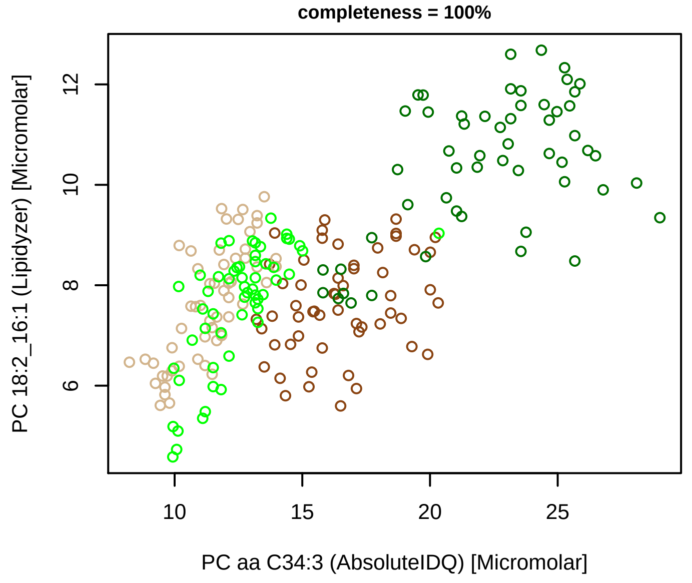


#### Bland−Altman Diagram

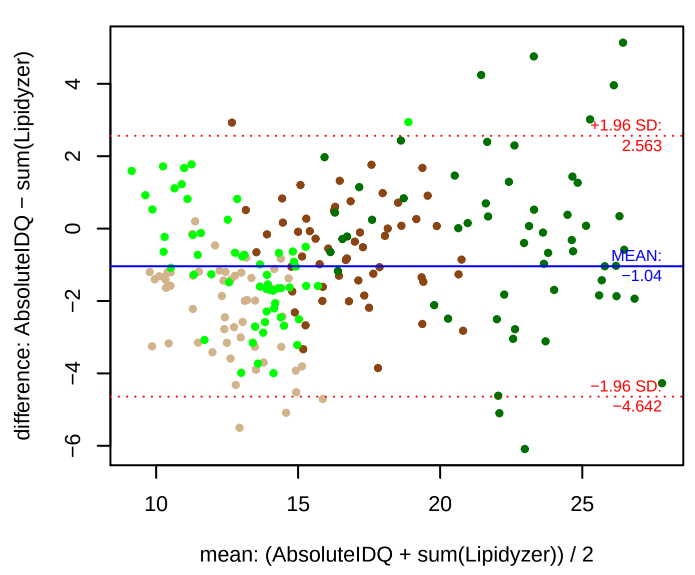


#### Stability of composition

**(PC 14:0_20:3 + PC 16:0_18:3 + PC 18:2_16:1) / PC aa C34:3**
**Shapiro−Wilk Test of log quotients, pv: 0.6127; OK**
SW: ok; ANOVA: Comp. ~ Subject_ID −> pv = 0.52902; EQUAL
Wilcoxon: Challenge; fasting: 0.375; sport: 0.25; OLTT: 0.375

**PC 14:0_20:3 / PC aa C34:3**
**Shapiro−Wilk Test of log quotients, pv: 0.00108; NO**
SW: ok; ANOVA: Comp. ~ Subject_ID −> pv = 4e−05; DIFFERENCE
Wilcoxon: Challenge; fasting: 0.125; sport: 0.25; OLTT: 0.25

**PC 16:0_18:3 / PC aa C34:3**
**Shapiro−Wilk Test of log quotients, pv: 0.13301; OK**
SW: ok; ANOVA: Comp. ~ Subject_ID −> pv = 0; DIFFERENCE
Wilcoxon: Challenge; fasting: 0.125; sport: 1; OLTT: 0.375

**PC 18:2_16:1 / PC aa C34:3**
**Shapiro−Wilk Test of log quotients, pv: 0.0115; OK**
SW: ok; ANOVA: Comp. ~ Subject_ID −> pv = 2e−04; DIFFERENCE
Wilcoxon: Challenge; fasting: 0.625; sport: 0.25; OLTT: 0.625

#### Quant. comp., Subject 5

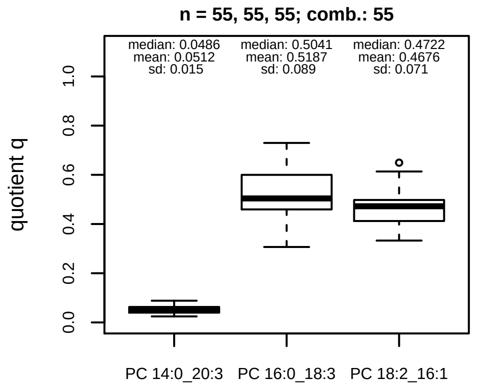


#### Quant. comp., Subject 6

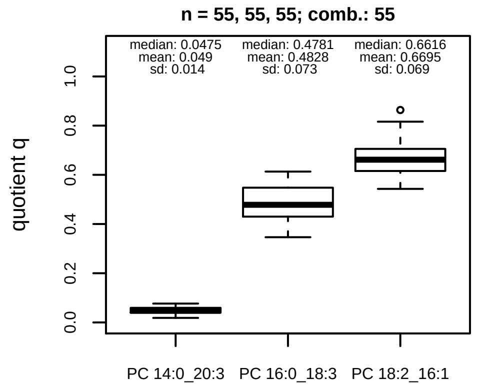


#### Quant. comp., Subject 7

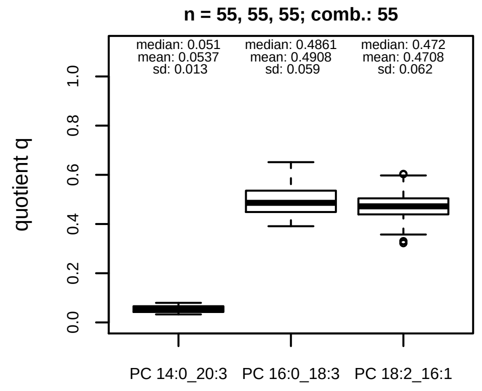


#### Quant. comp., Subject 8

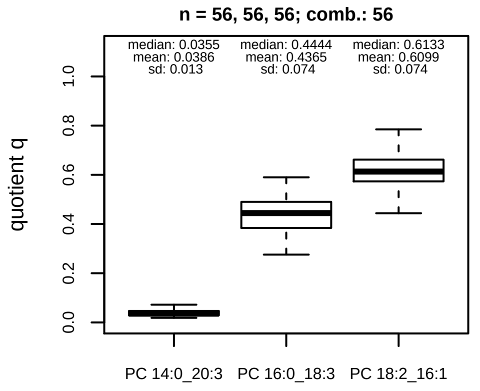


#### Trends of proportions q during challenges

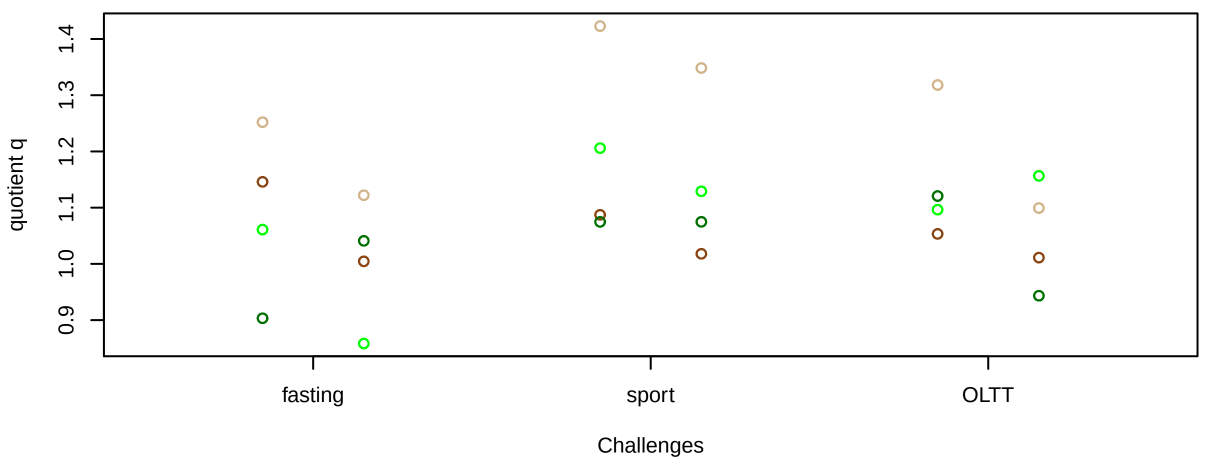


#### Proportions q per time point

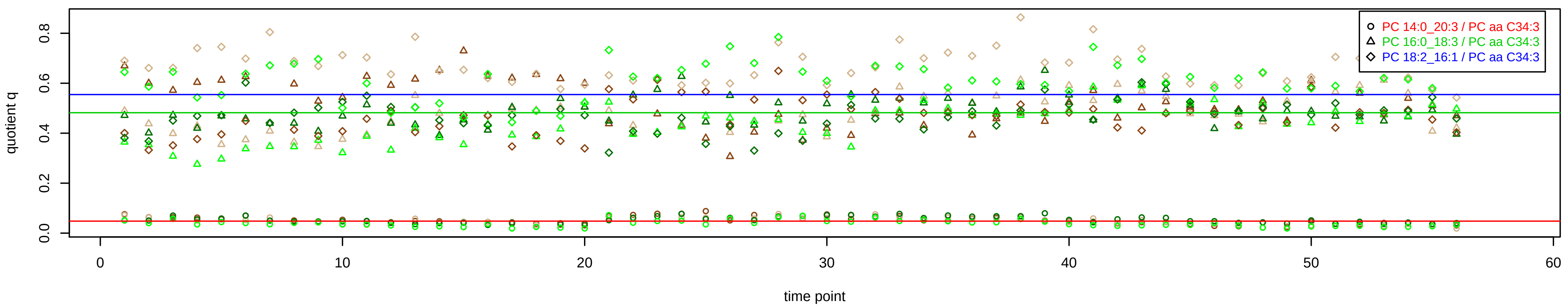

# PC aa C34:4 = PC 14:0_20:4 + PC 16:0_18:4 + R

PC 16:0_18:4 excluded because of missingness > 75%

#### Semi−Quantitative composition within Lipidyzer

n = 223

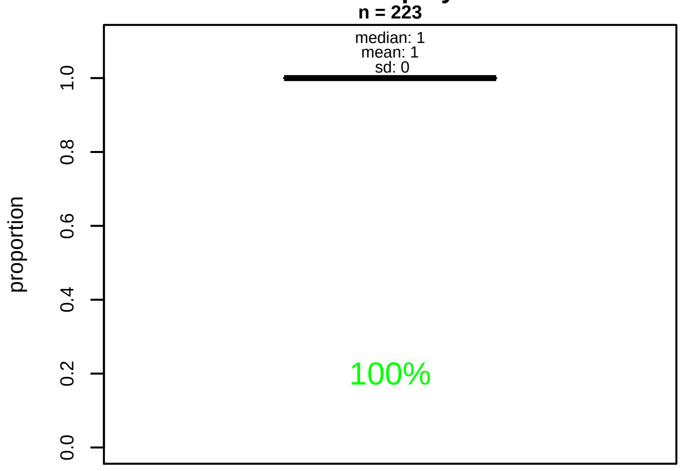


#### Semi−Quantitative composition quotients q

n = 221

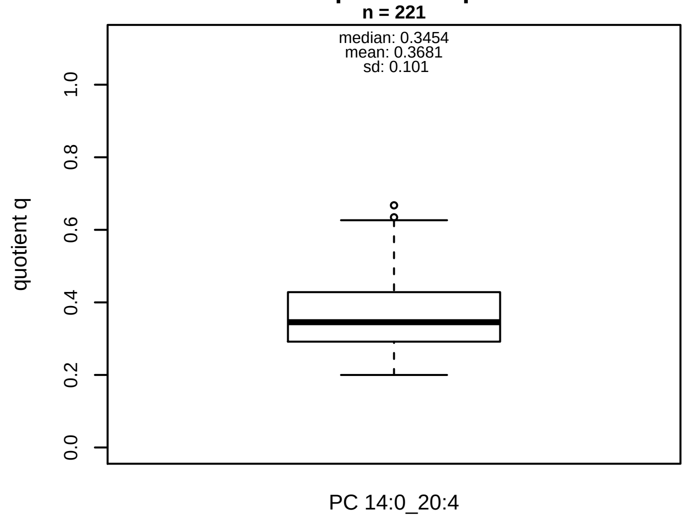


#### Semi−Quantitative composition: Lipidyzer

PC aa C34:4 consists of:
PC 14:0_20:4 100%
and of (not quantified):
PC 16:0_18:4, PC 12:0_22:4, PC 14:1_20:3, PC 16:1_18:3,
PC O−15:0_20:4
and further compounds

#### Composition: mean of proportions q

conc(PC aa C34:4) * 0.3681 = conc(PC 14:0_20:4) [var(q)=0.3681]
Percentiles: 5%−>0.2438, 25%−>0.2917, 75%−>0.4282, 95%−>0.565

#### Linear model

```
PC aa C34:4 ~ b * ( PC 14:0_20:4 )
```

b = 2.47409

$R^2$ = 0.53623

#### Ranges

| Measure | AbsoluteIDQ | sum(Lipidyzer) | delta |
|---|---|---|---|
| Min | 0.65 | 0.18 | 0.47 |
| Max | 2.72 | 0.96 | 1.77 |
| Mean | 1.41 | 0.49 | 0.92 |
| Median | 1.34 | 0.49 | 0.84 |
| SD | 0.52 | 0.16 | 0.37 |

#### Concentration: AbsoluteIDQ vs Lipidyzer

completeness = 100%

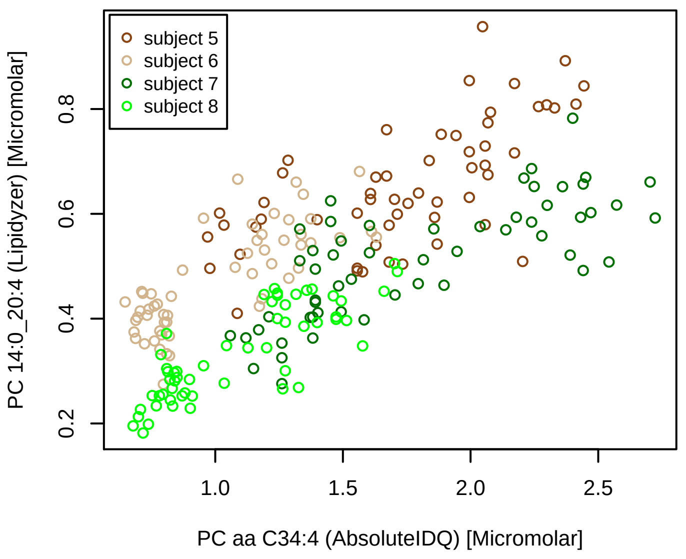


#### Bland−Altman Diagram

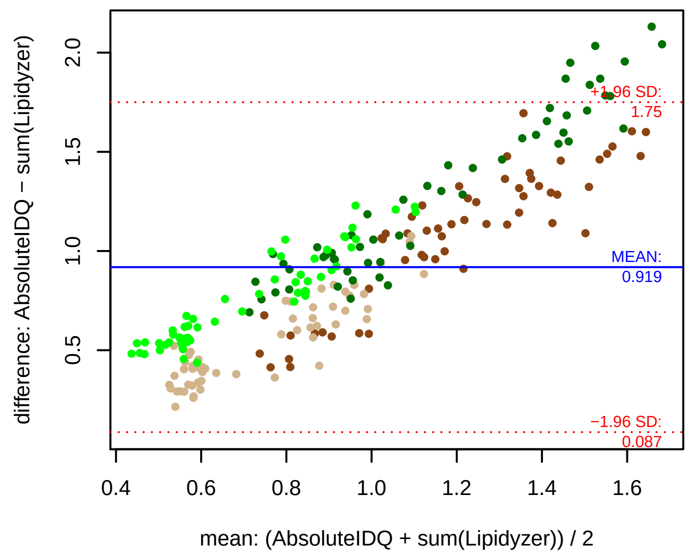


#### Stability of composition

**PC 14:0_20:4 / PC aa C34:4**
**Shapiro−Wilk Test of log quotients, pv: 0.01137; OK**
SW: ok; ANOVA: Comp. ~ Subject_ID −> pv = 0; DIFFERENCE
Wilcoxon: Challenge; fasting: 0.25; sport: 0.875; OLTT: 0.375

#### Quant. comp., Subject 5

n = 55; comb.: 55

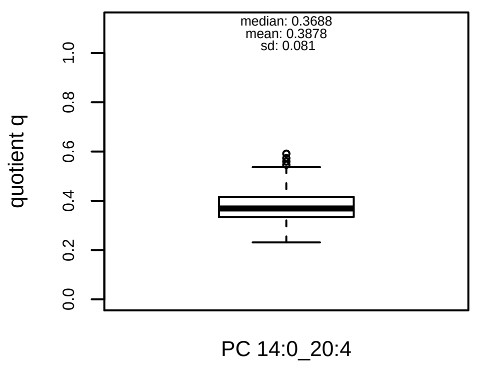


#### Quant. comp., Subject 6

n = 55; comb.: 55

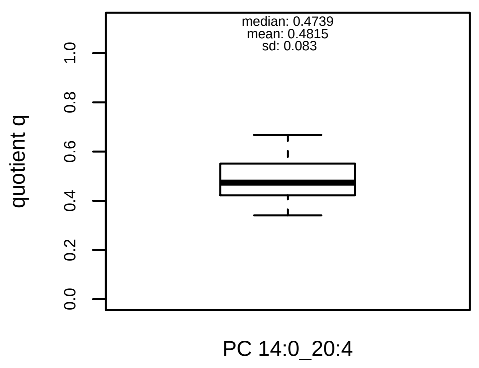


#### Quant. comp., Subject 7

n = 55; comb.: 55

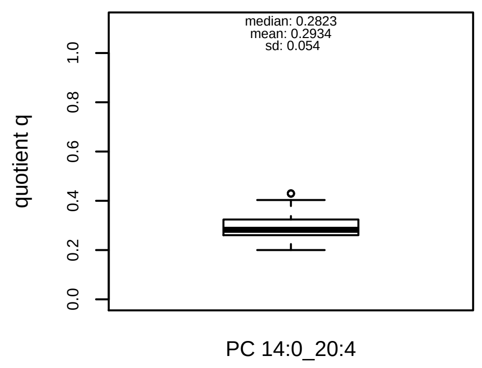


#### Quant. comp., Subject 8

n = 56; comb.: 56

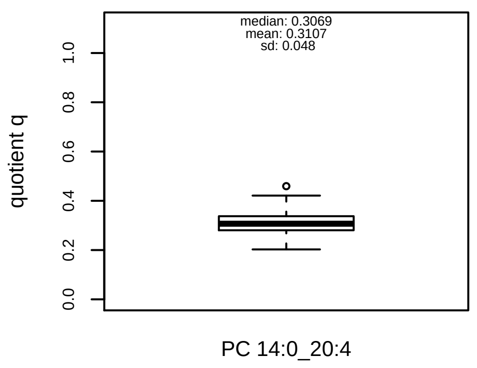


#### Trends of proportions q during challenges

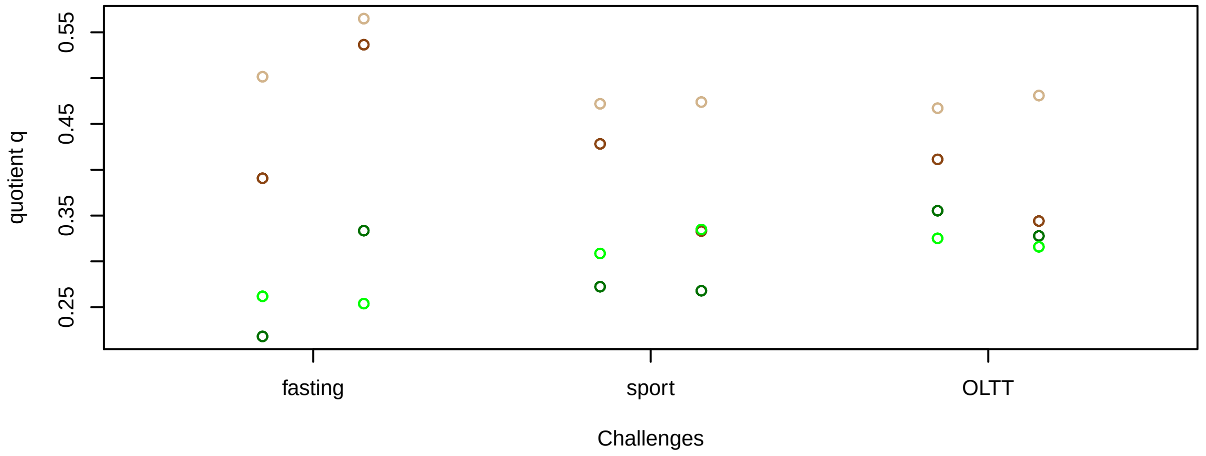


#### Proportions q per time point

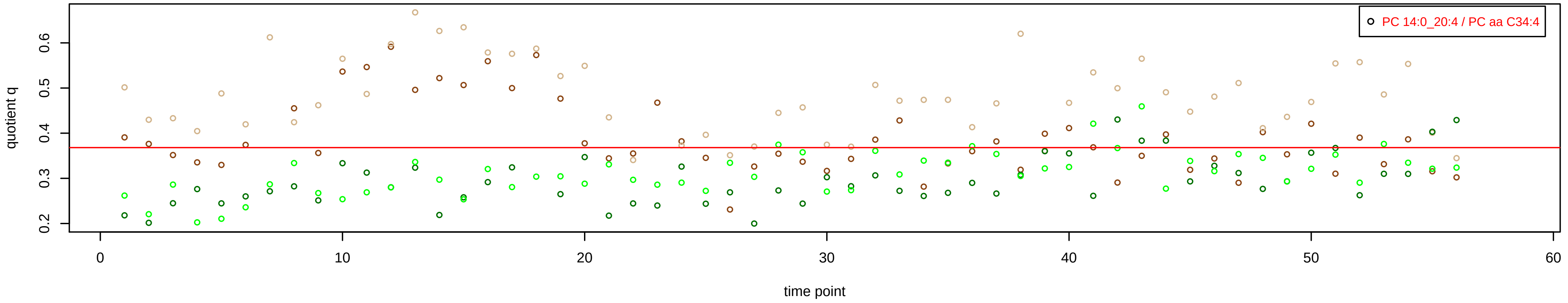

# PC aa C36:0 = PC 18:0_18:0 + R

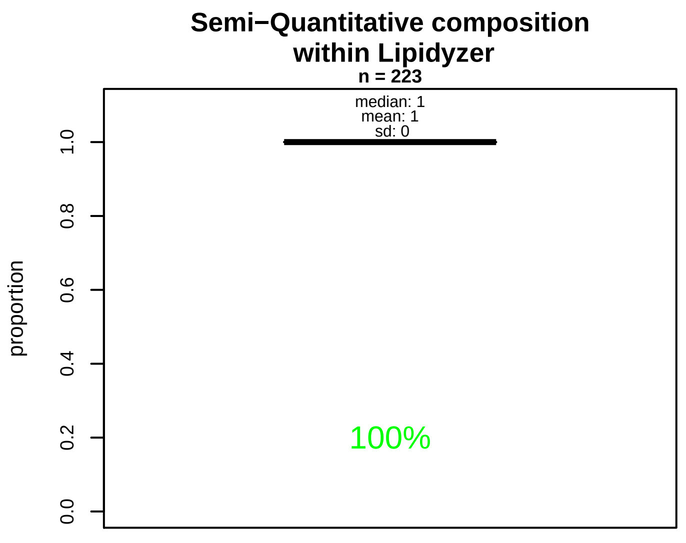


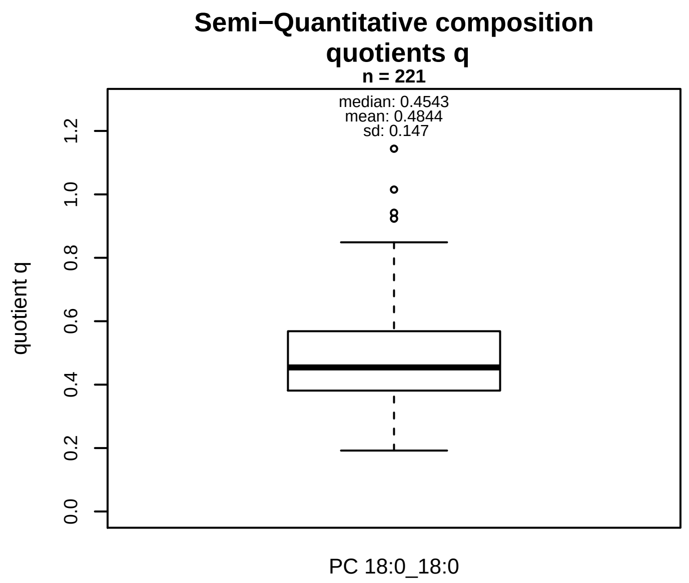


### Semi−Quantitative composition: Lipidyzer

PC aa C36:0 consists of:
PC 18:0_18:0 100%
and of (not quantified):
PC 11:0_25:0, PC 12:0_24:0, PC 13:0_23:0, PC 14:0_22:0,
PC 15:0_21:0, PC 16:0_20:0, PC 17:0_19:0, PC 15:1_22:6,
PC 17:2_20:5, PC O−17:0_20:0, PC O−16:0_21:0, PC O−18:
[13C1]SM 40:0
and further compounds

### Composition: mean of proportions q

conc(PC aa C36:0) * 0.4844 = conc(PC 18:0_18:0) [var(q)=0.4844]
Percentiles: 5%−>0.3066, 25%−>0.3812, 75%−>0.5685, 95%−>0.747

### Linear model

```
PC aa C36:0 ~ b * ( PC 18:0_18:0 )
```

b = 0.75569

$R^2$ = 0.14288

### Ranges

| Measure | AbsoluteIDQ | sum(Lipidyzer) | delta |
|---|---|---|---|
| Min | 1.69 | 0.56 | 1.12 |
| Max | 6.62 | 3.5 | 3.12 |
| Mean | 2.72 | 1.27 | 1.44 |
| Median | 2.48 | 1.24 | 1.24 |
| SD | 0.77 | 0.39 | 0.39 |

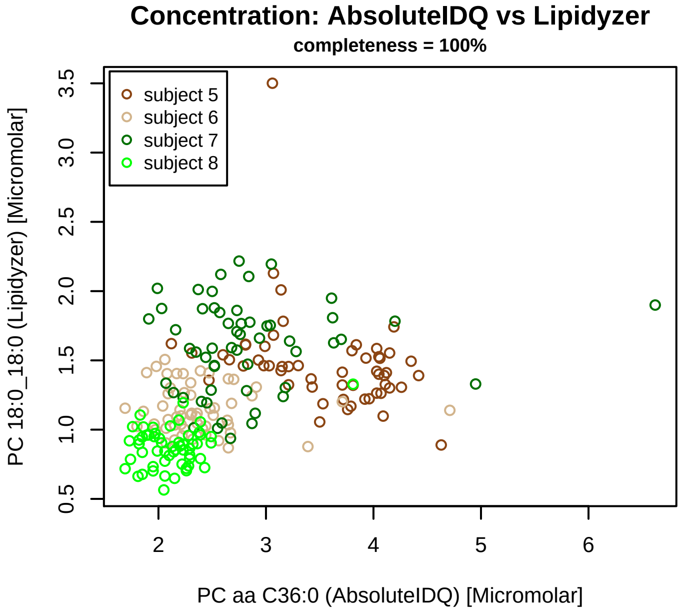


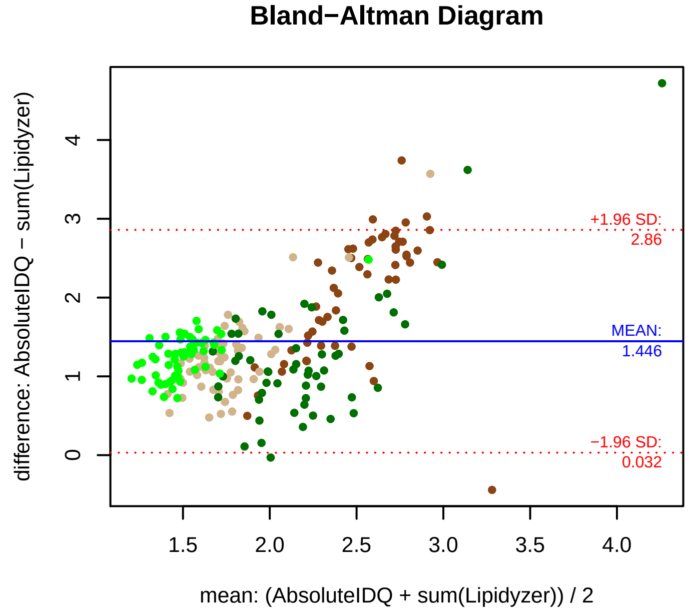


### Stability of composition

**PC 18:0_18:0 / PC aa C36:0**
**Shapiro−Wilk Test of log quotients, pv: 0.64583; OK**
SW: ok; ANOVA: Comp. ~ Subject_ID −> pv = 0.477; EQUAL
Wilcoxon: Challenge; fasting: 0.625; sport: 0.25; OLTT: 0.625

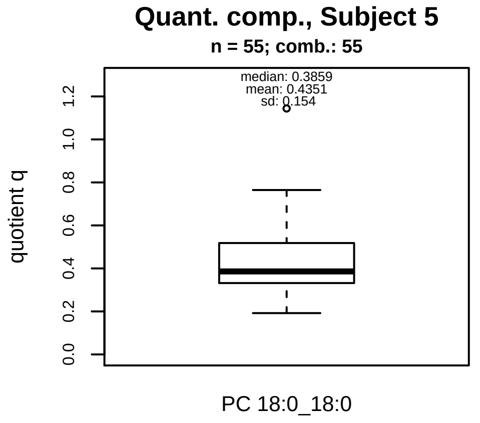


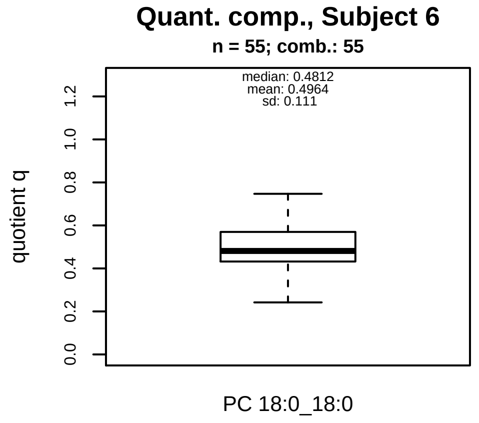


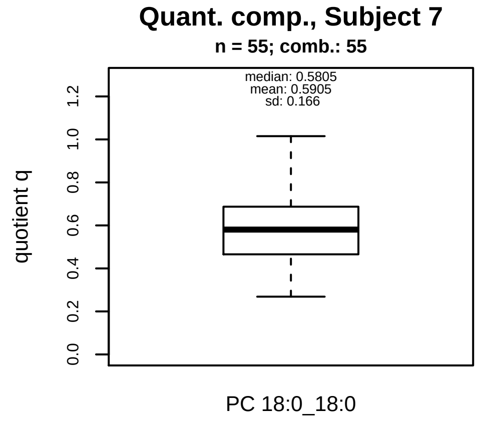


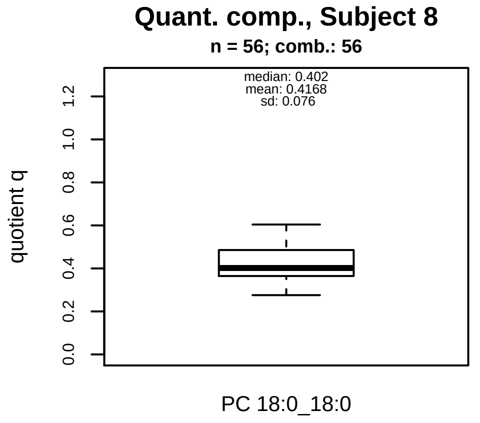


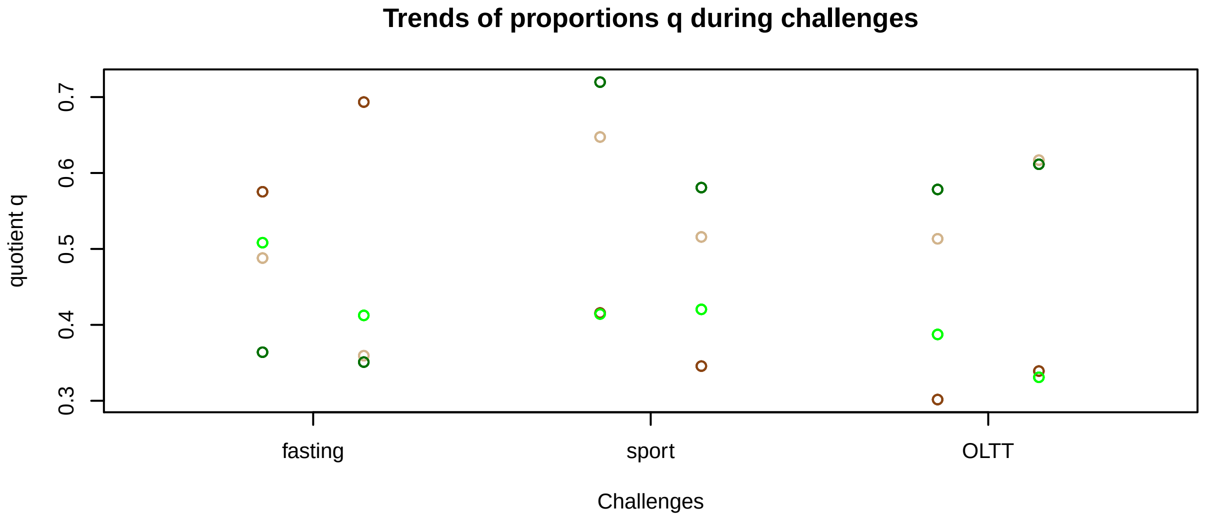


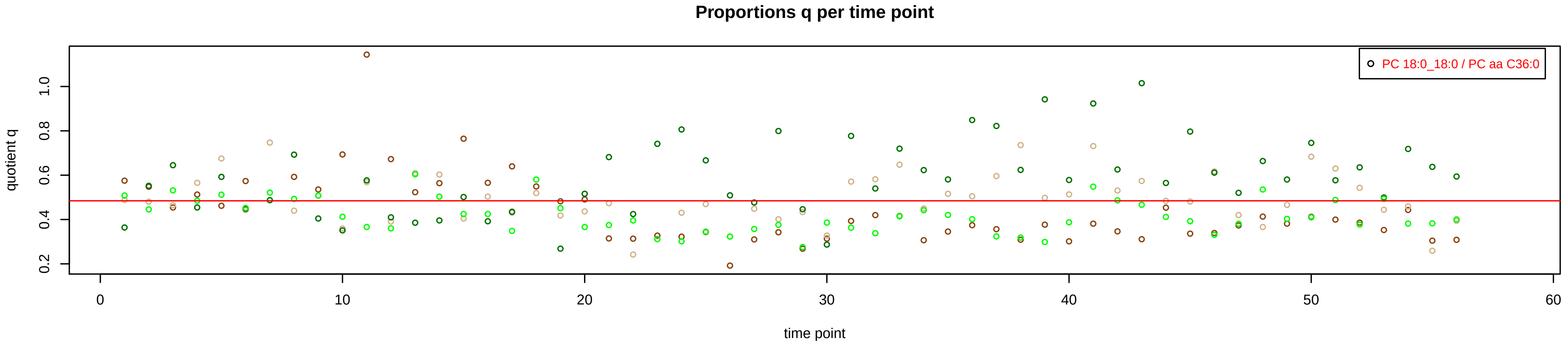

# PC aa C36:1 = PC 16:0_20:1 + PC 18:0_18:1 + PC 20:0_16:1 + R

PC 20:0_16:1 excluded because of missingness > 75%

#### Semi−Quantitative composition within Lipidyzer

**n = 223**

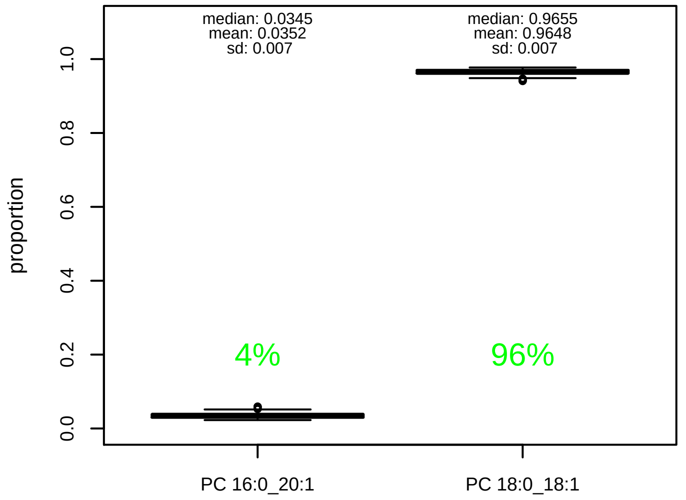


#### Semi−Quantitative composition quotients q

**n = 221**

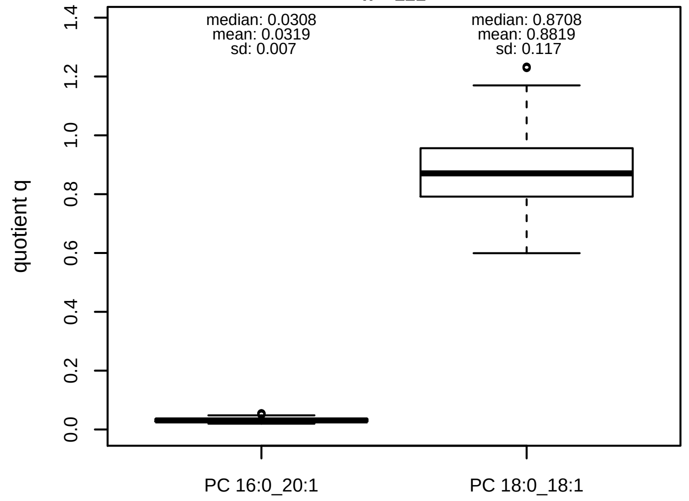


#### Semi−Quantitative composition: Lipidyzer

PC aa C36:1 consists of:
PC 16:0_20:1 3.5%
PC 18:0_18:1 96.5%
and of (not quantified):
PC 20:0_16:1, PC 14:0_22:1, PC 14:1_22:0, PC 15:1_21:0,
PC 16:1_20:0, PC 17:0_19:1, PC 17:1_19:0, PC O−18:0_19:[illegible]
PC O−20:0_17:1, PC O−16:1_21:0, PC O−18:1_19:0, PC O−[illegible]
[13C1]SM 40:1
and further compounds

#### Composition: mean of proportions q

conc(PC aa C36:1) * 0.0319 = conc(PC 16:0_20:1) [var(q)=0.0319]
Percentiles: 5%−>0.0231, 25%−>0.027, 75%−>0.0359, 95%−>0.0447
conc(PC aa C36:1) * 0.8819 = conc(PC 18:0_18:1) [var(q)=0.8819]
Percentiles: 5%−>0.7171, 25%−>0.7916, 75%−>0.9562, 95%−>1.0794

#### Linear model

```
PC aa C36:1 ~ b * ( PC 16:0_20:1
   + PC 18:0_18:1 )
```

b = 0.89548

$R^2$ = 0.7898

#### Ranges

| Measure | AbsoluteIDQ | sum(Lipidyzer) | delta |
|---|---|---|---|
| Min | 20.7 | 19.85 | 0.85 |
| Max | 68.9 | 68.55 | 0.35 |
| Mean | 41.15 | 37.5 | 3.66 |
| Median | 40.6 | 34.64 | 5.96 |
| SD | 11.29 | 11.16 | 0.13 |

#### Concentration: AbsoluteIDQ vs Lipidyzer

**completeness = 100%**

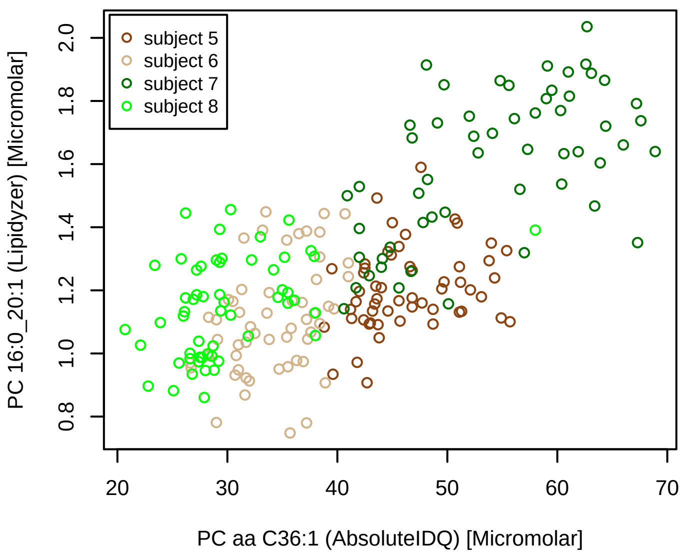


#### Concentration: AbsoluteIDQ vs Lipidyzer

**completeness = 100%**

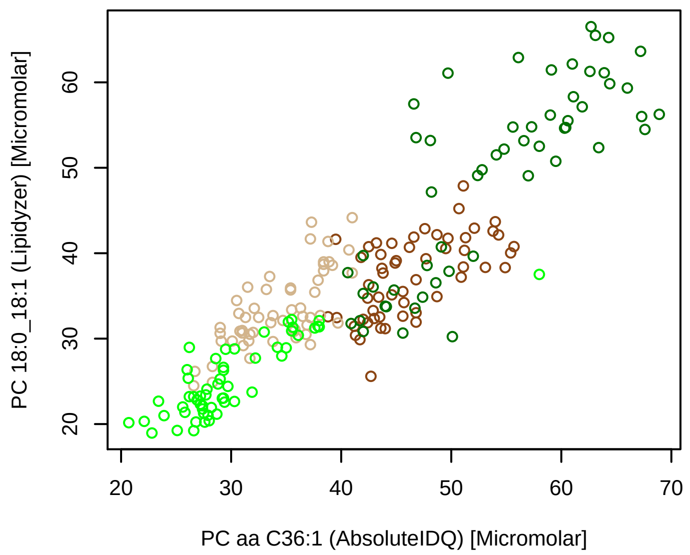


#### Bland−Altman Diagram

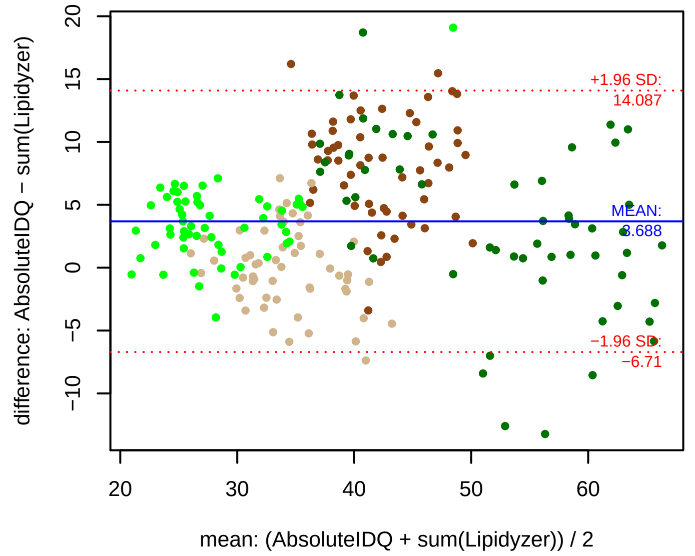


#### Stability of composition

**(PC 16:0_20:1 + PC 18:0_18:1) / PC aa C36:1**
**Shapiro−Wilk Test of log quotients, pv: 0.7629; OK**
SW: ok; ANOVA: Comp. ~ Subject_ID −> pv = 0.16842; EQUAL
Wilcoxon: Challenge; fasting: 0.25; sport: 0.125; OLTT: 0.875

**PC 16:0_20:1 / PC aa C36:1**
**Shapiro−Wilk Test of log quotients, pv: 0.27012; OK**
SW: ok; ANOVA: Comp. ~ Subject_ID −> pv = 0; DIFFERENCE
Wilcoxon: Challenge; fasting: 0.25; sport: 0.125; OLTT: 0.625

**PC 18:0_18:1 / PC aa C36:1**
**Shapiro−Wilk Test of log quotients, pv: 0.66034; OK**
SW: ok; ANOVA: Comp. ~ Subject_ID −> pv = 0.39144; EQUAL
Wilcoxon: Challenge; fasting: 0.25; sport: 0.125; OLTT: 0.875

#### Quant. comp., Subject 5

**n = 55, 55; comb.: 55**

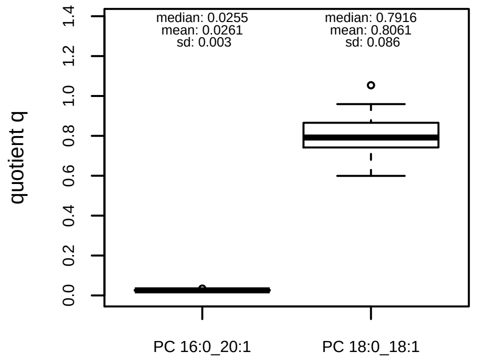


#### Quant. comp., Subject 6

**n = 55, 55; comb.: 55**

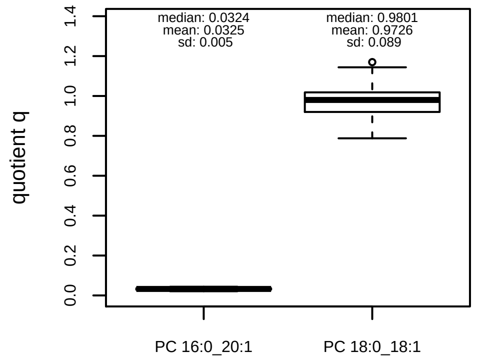


#### Quant. comp., Subject 7

**n = 55, 55; comb.: 55**

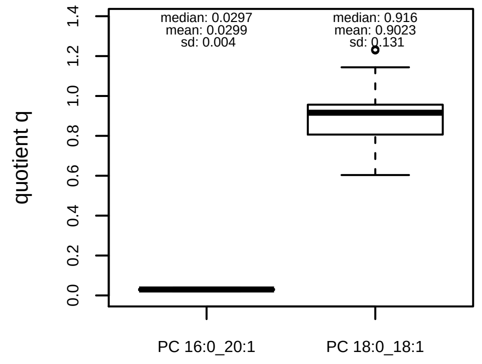


#### Quant. comp., Subject 8

**n = 56, 56; comb.: 56**

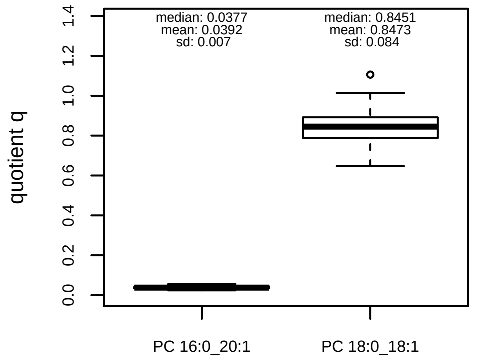


#### Trends of proportions q during challenges

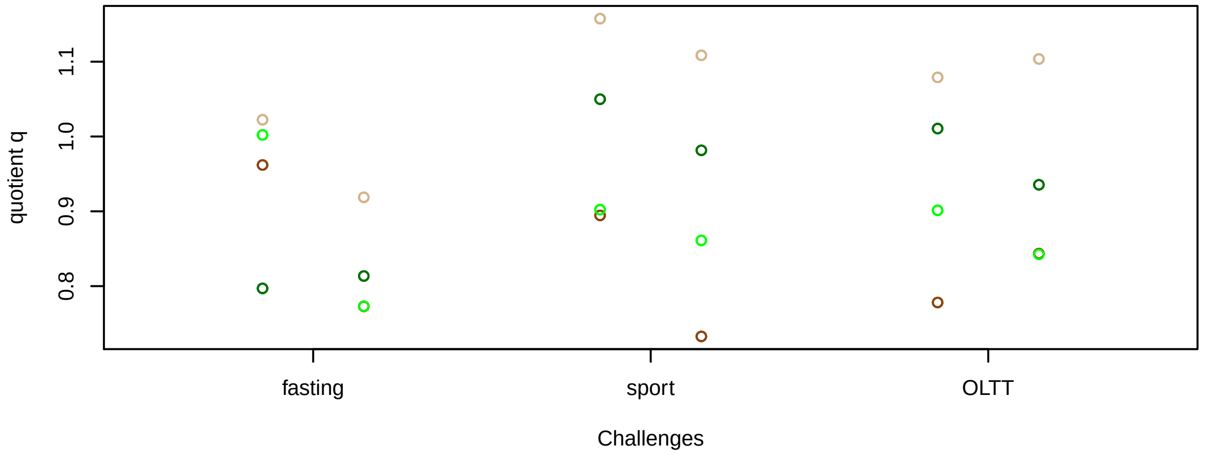


#### Proportions q per time point

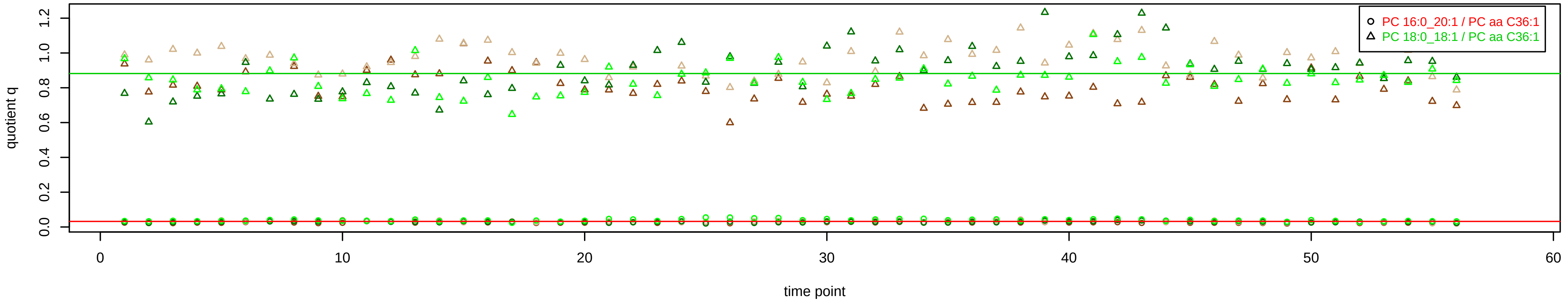

# PC aa C36:2 = PC 16:0_20:2 + PC 18:0_18:2 + PC 18:1_18:1 + R

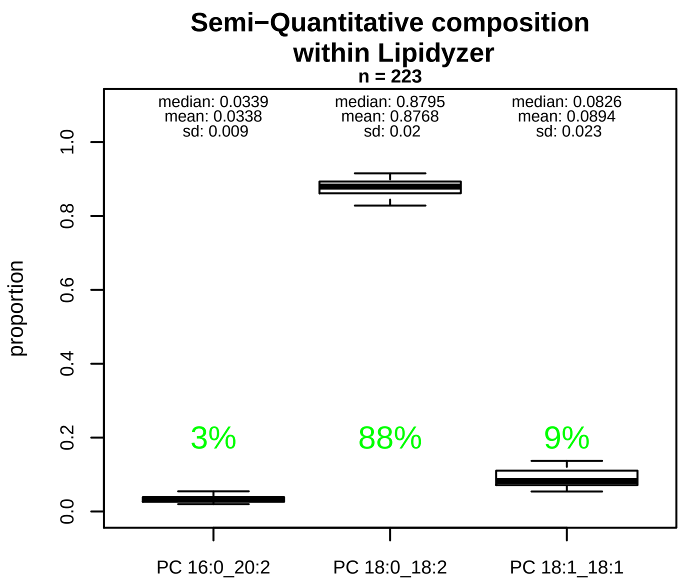


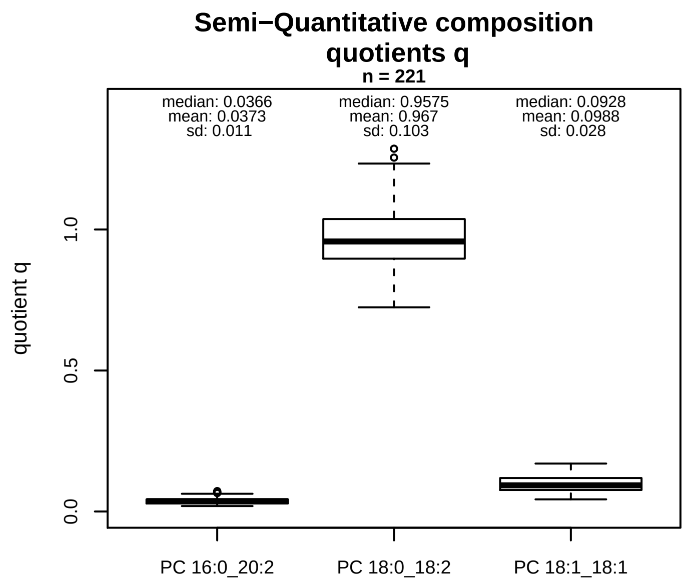


### Semi−Quantitative composition: Lipidyzer

PC aa C36:2 consists of:
PC 16:0_20:2 3.4%
PC 18:0_18:2 87.7%
PC 18:1_18:1 8.9%
and of (not quantified):
PC14 0 22:2, PC14 1 22:1, PC16 1 20:1, PC17 1 19:1,
PC17 2 19:0, PCO−20 0 17:2, PCO−18 1 19:1, PCO−20 1 17
[13C1]SM 40:2
and further compounds

### Composition: mean of proportions q

conc(PC aa C36:2) * 0.0373 = conc(PC 16:0_20:2) [var(q)=0.0373]
Percentiles: 5%−>0.0226, 25%−>0.0285, 75%−>0.0432, 95%−>0.0581
conc(PC aa C36:2) * 0.967 = conc(PC 18:0_18:2) [var(q)=0.967]
Percentiles: 5%−>0.8172, 25%−>0.8964, 75%−>1.037, 95%−>1.1362
conc(PC aa C36:2) * 0.0988 = conc(PC 18:1_18:1) [var(q)=0.0988]
Percentiles: 5%−>0.0601, 25%−>0.0762, 75%−>0.1186, 95%−>0.1467

### Linear model

```
PC aa C36:2 ~ b * ( PC 16:0_20:2
  + PC 18:0_18:2
  + PC 18:1_18:1 )
```

b = 0.6305

$R^2$ = 0.57943

### Ranges

| Measure | AbsoluteIDQ | sum(Lipidyzer) | delta |
|---|---|---|---|
| Min | 125.5 | 141.63 | 16.13 |
| Max | 256.9 | 302.37 | 45.47 |
| Mean | 190.68 | 209.51 | 18.83 |
| Median | 189.6 | 204.41 | 14.81 |
| SD | 27.84 | 33.51 | 5.67 |

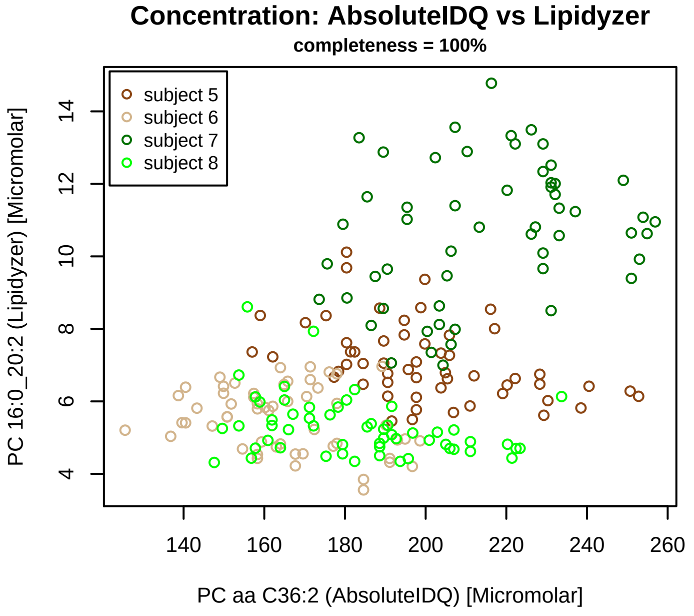


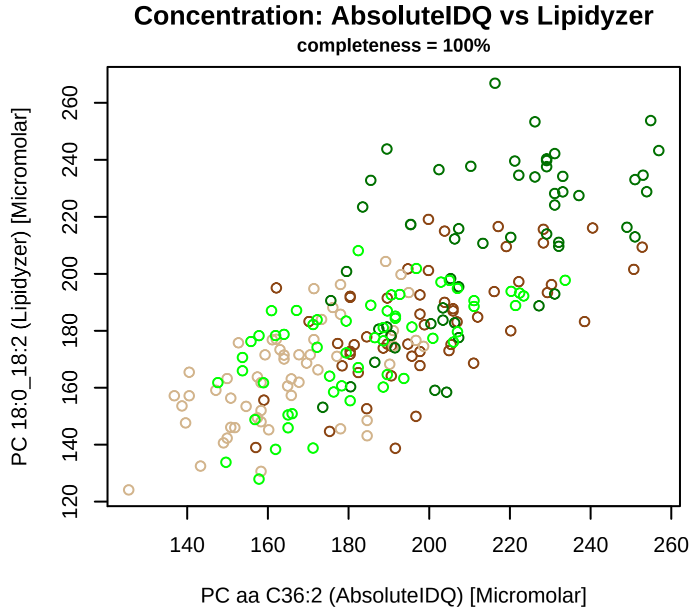


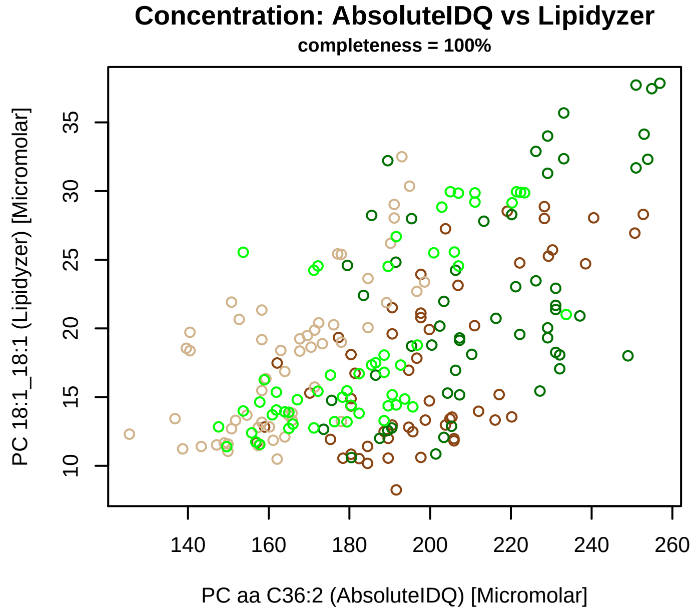


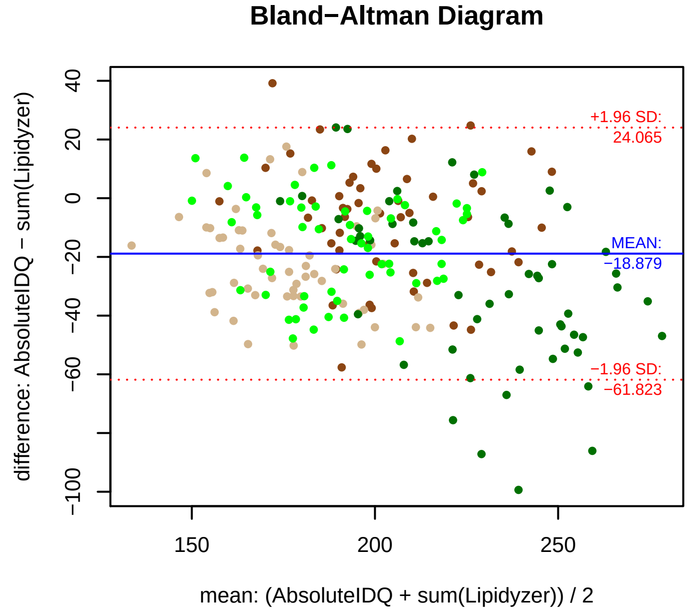


### Stability of composition

**(PC 16:0_20:2 + PC 18:0_18:2 + PC 18:1_18:1) / PC aa C36:2**
**Shapiro−Wilk Test of log quotients, pv: 0.68759; OK**
SW: ok; ANOVA: Comp. ~ Subject_ID −> pv = 0.03886; EQUAL
Wilcoxon: Challenge; fasting: 0.25; sport: 0.125; OLTT: 0.375

**PC 16:0_20:2 / PC aa C36:2**
**Shapiro−Wilk Test of log quotients, pv: 0.06915; OK**
SW: no; Kruskal: Comp. ~ Subject_ID −> pv = 0; DIFFERENCE
Wilcoxon: Challenge; fasting: 0.375; sport: 0.125; OLTT: 0.125

**PC 18:0_18:2 / PC aa C36:2**
**Shapiro−Wilk Test of log quotients, pv: 0.72516; OK**
SW: ok; ANOVA: Comp. ~ Subject_ID −> pv = 0.08316; EQUAL
Wilcoxon: Challenge; fasting: 0.125; sport: 0.125; OLTT: 0.375

**PC 18:1_18:1 / PC aa C36:2**
**Shapiro−Wilk Test of log quotients, pv: 0.00285; NO**
SW: no; Kruskal: Comp. ~ Subject_ID −> pv = 0.00031; DIFFERENCE
Wilcoxon: Challenge; fasting: 0.875; sport: 0.125; OLTT: 0.375

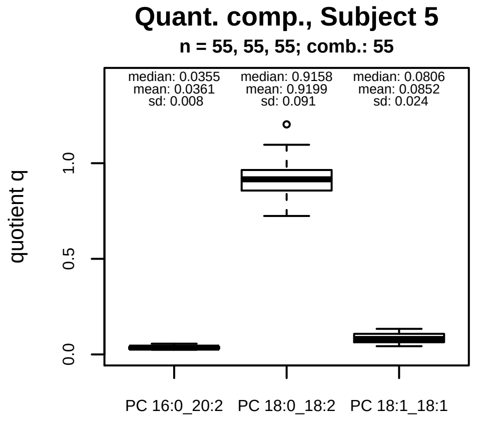


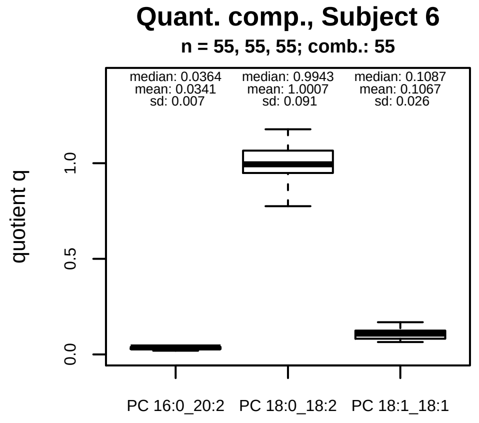


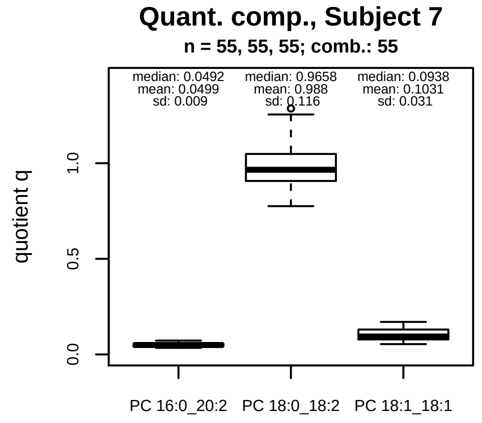


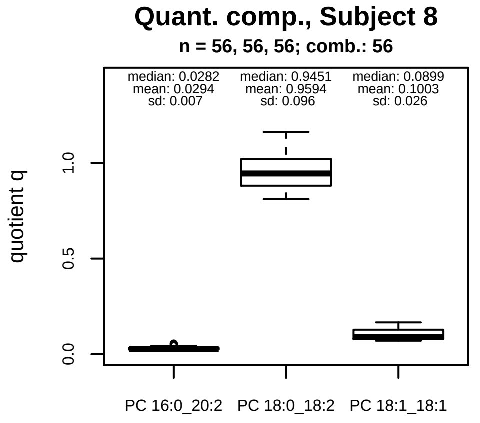


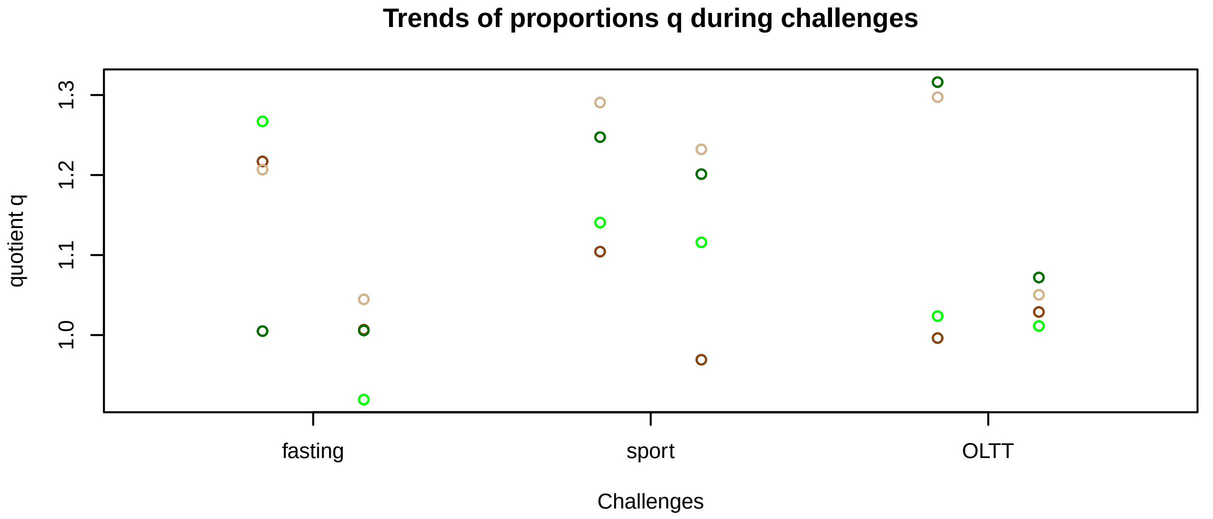


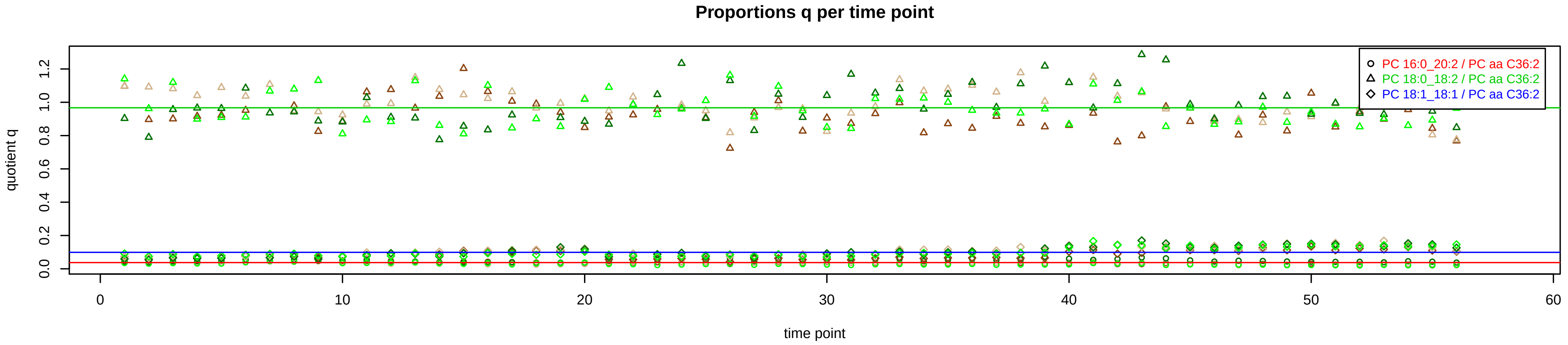

# PC aa C36:3 = PC 16:0_20:3 + PC 18:0_18:3 + PC 18:1_18:2 + R

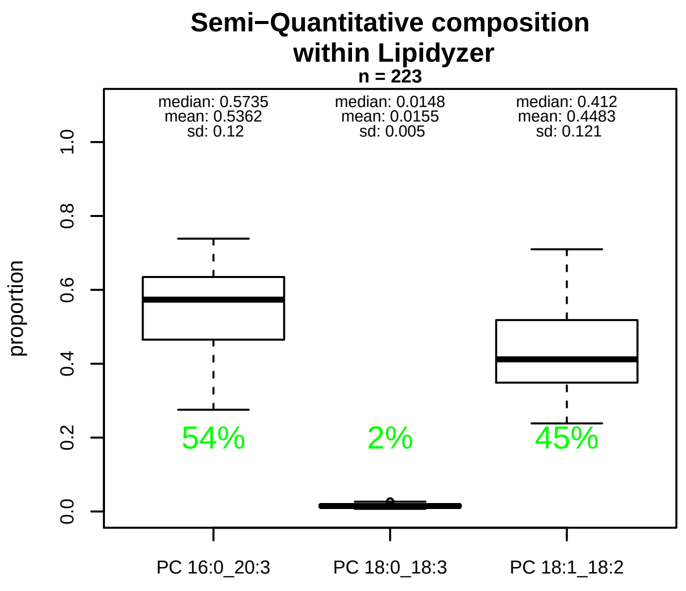


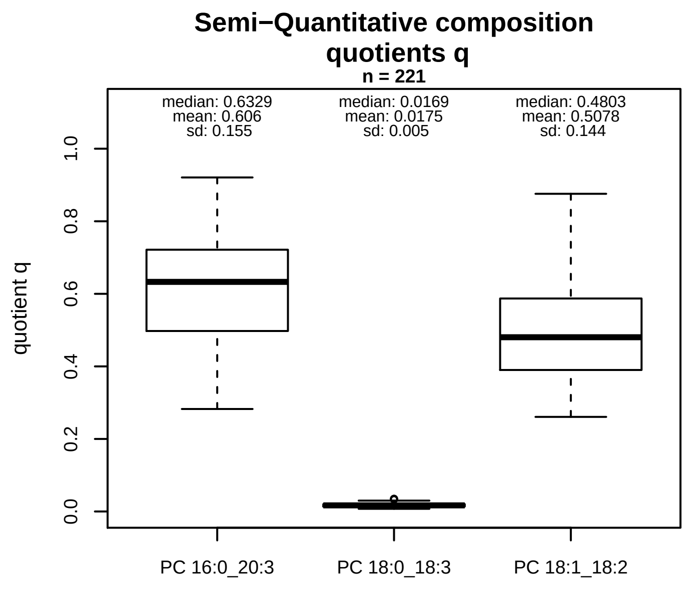


### Semi−Quantitative composition: Lipidyzer

PC aa C36:3 consists of:
PC 16:0_20:3 53.6%
PC 18:0_18:3 1.5%
PC 18:1_18:2 44.8%
and of (not quantified):
PC14 1 22:2, PC16 1 20:2, PC17 2 19:1, PCO−20 1 17:2, SM40.3
and further compounds

### Composition: mean of proportions q

conc(PC aa C36:3) * 0.606 = conc(PC 16:0_20:3) [var(q)=0.606]
Percentiles: 5%−>0.3434, 25%−>0.4975, 75%−>0.7216, 95%−>0.8409
conc(PC aa C36:3) * 0.0175 = conc(PC 18:0_18:3) [var(q)=0.0175]
Percentiles: 5%−>0.01, 25%−>0.0137, 75%−>0.0203, 95%−>0.0279
conc(PC aa C36:3) * 0.5078 = conc(PC 18:1_18:2) [var(q)=0.5078]
Percentiles: 5%−>0.3177, 25%−>0.3901, 75%−>0.5871, 95%−>0.7643

### Linear model

```
PC aa C36:3 ~ b * ( PC 16:0_20:3
  + PC 18:0_18:3
  + PC 18:1_18:2 )
```

b = 0.79714

$R^2$ = 0.80444

### Ranges

| Measure | AbsoluteIDQ | sum(Lipidyzer) | delta |
|---|---|---|---|
| Min | 84.1 | 86.21 | 2.11 |
| Max | 222 | 250.69 | 28.69 |
| Mean | 133.31 | 150 | 16.69 |
| Median | 126 | 141.27 | 15.27 |
| SD | 31.67 | 35.54 | 3.87 |

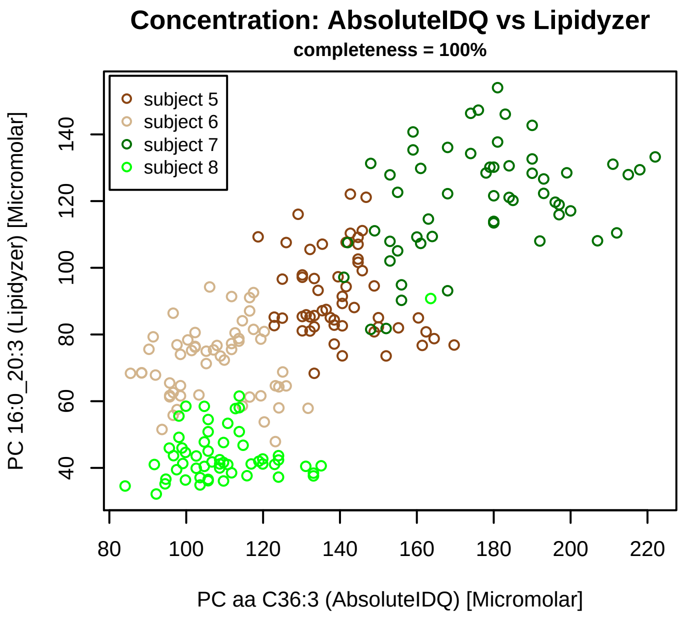


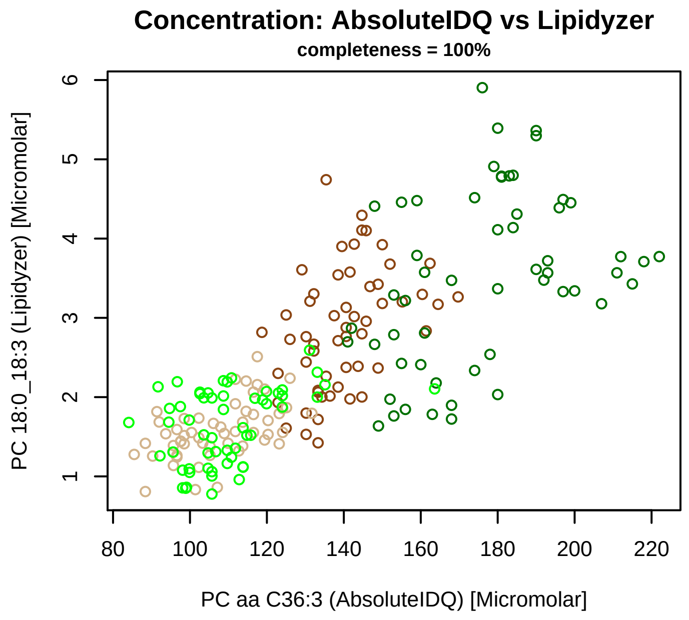


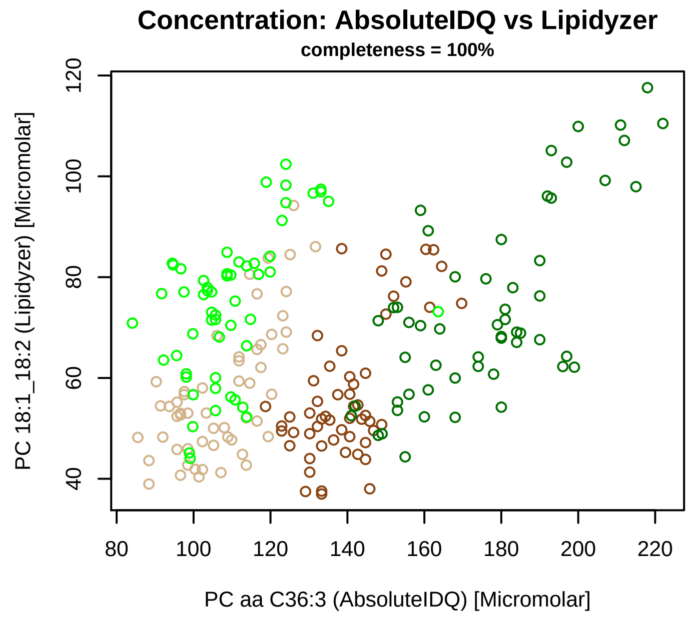


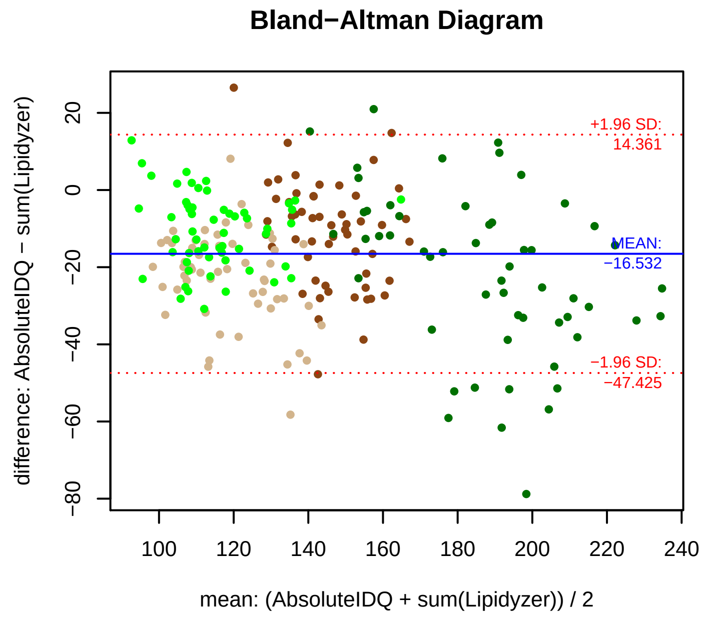


### Stability of composition

**(PC 16:0_20:3 + PC 18:0_18:3 + PC 18:1_18:2) / PC aa C36:3**
**Shapiro−Wilk Test of log quotients, pv: 0.43192; OK**
SW: ok; ANOVA: Comp. ~ Subject_ID −> pv = 0.6275; EQUAL
Wilcoxon: Challenge; fasting: 0.25; sport: 0.125; OLTT: 0.25

**PC 16:0_20:3 / PC aa C36:3**
**Shapiro−Wilk Test of log quotients, pv: 0; NO**
SW: no; Kruskal: Comp. ~ Subject_ID −> pv = 0; DIFFERENCE
Wilcoxon: Challenge; fasting: 0.375; sport: 0.125; OLTT: 0.125

**PC 18:0_18:3 / PC aa C36:3**
**Shapiro−Wilk Test of log quotients, pv: 0.6707; OK**
SW: ok; ANOVA: Comp. ~ Subject_ID −> pv = 0.00016; DIFFERENCE
Wilcoxon: Challenge; fasting: 0.125; sport: 0.625; OLTT: 0.625

**PC 18:1_18:2 / PC aa C36:3**
**Shapiro−Wilk Test of log quotients, pv: 0.00353; NO**
SW: no; Kruskal: Comp. ~ Subject_ID −> pv = 0; DIFFERENCE
Wilcoxon: Challenge; fasting: 0.125; sport: 0.125; OLTT: 1

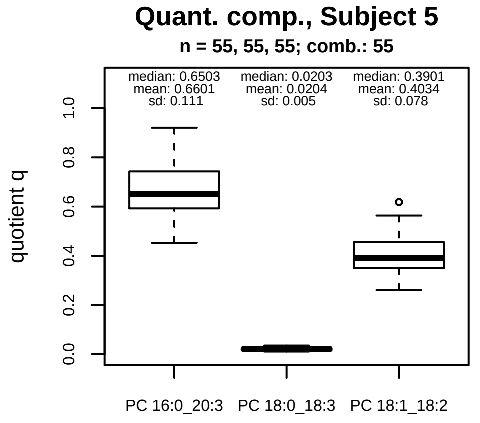


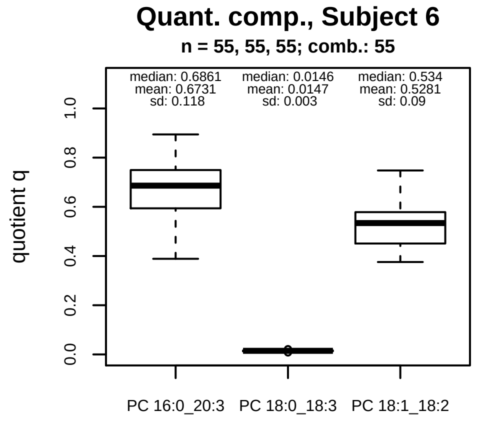


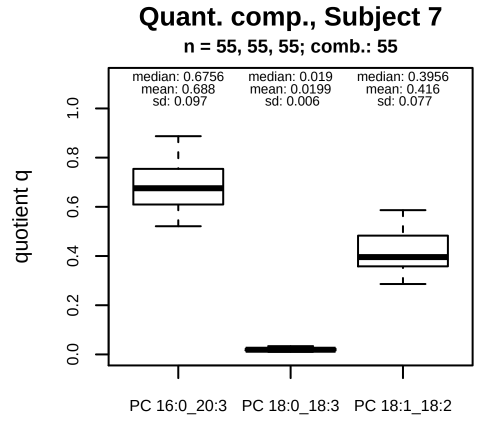


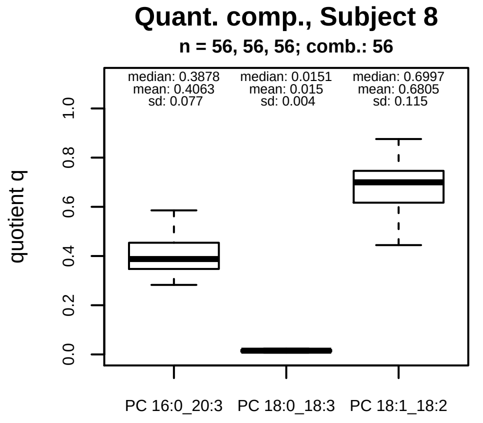


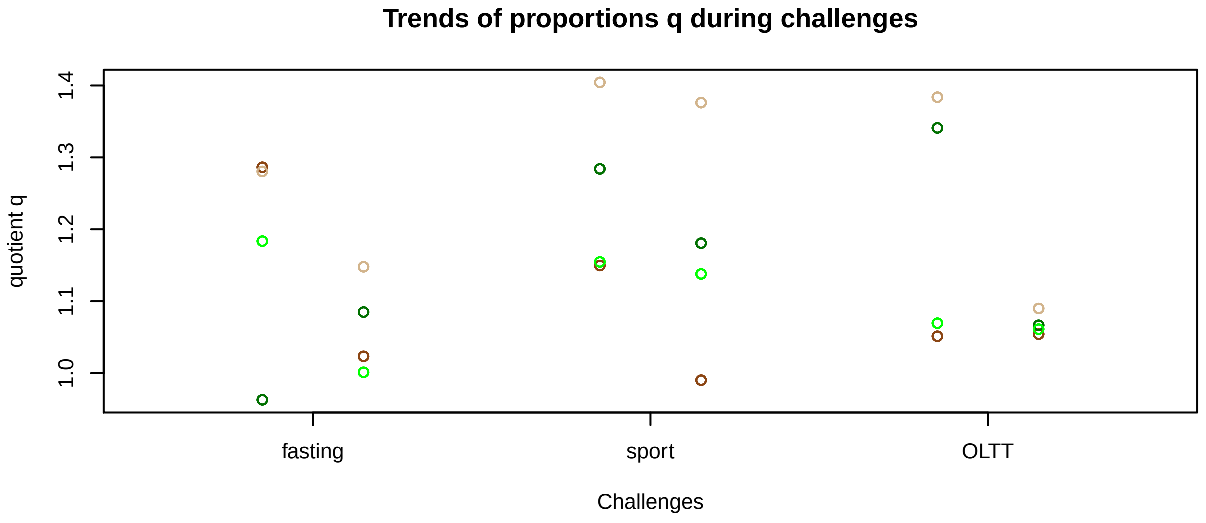


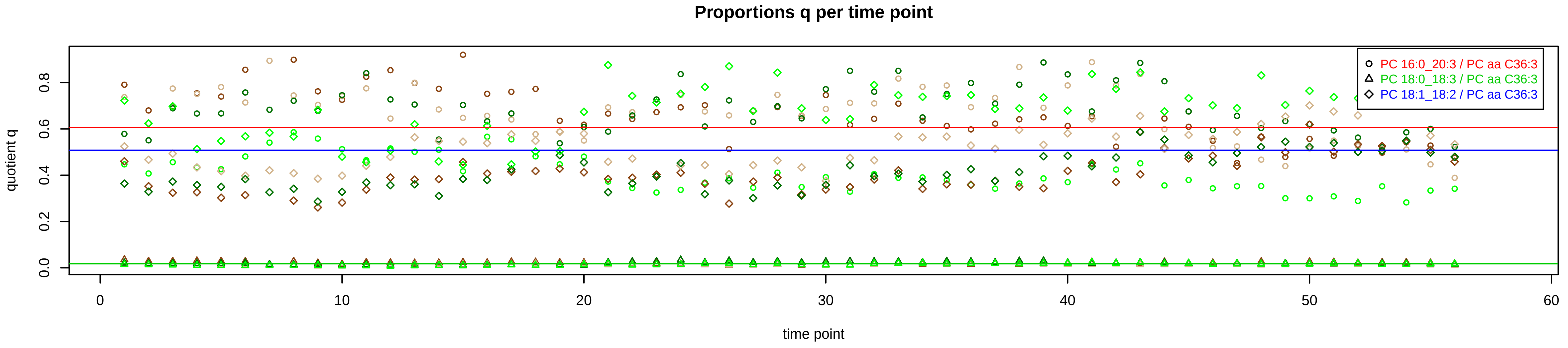

# PC aa C36:4 = PC 14:0_22:4 + PC 16:0_20:4 + PC 18:0_18:4 + PC 18:1_18:3 + PC 18:2_18:2 + R

PC 18:0_18:4 excluded because of missingness > 75%

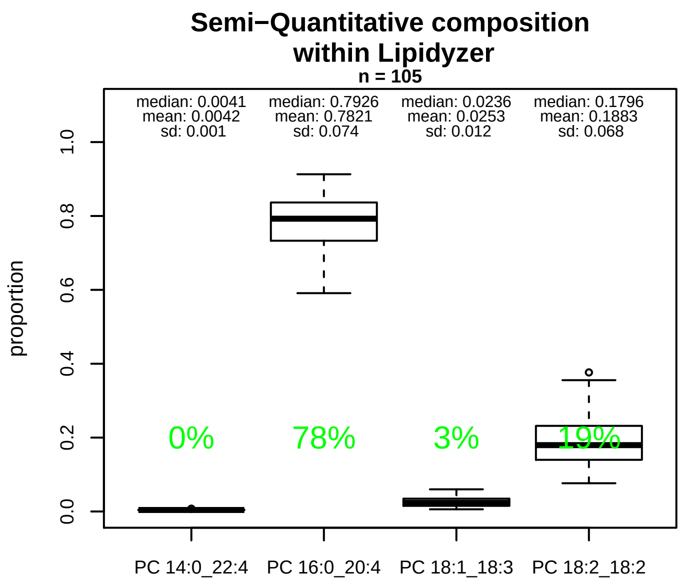


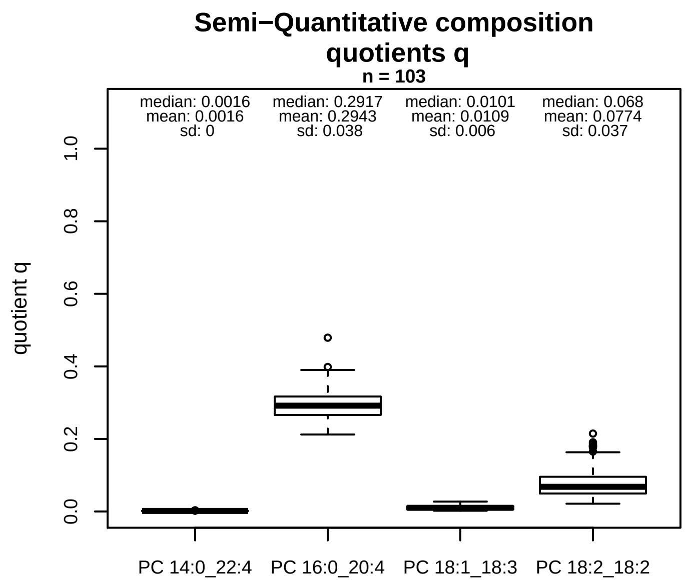


### Semi−Quantitative composition: Lipidyzer

PC aa C36:4 consists of:
- PC 14:0_22:4 0.4%
- PC 16:0_20:4 78.2%
- PC 18:1_18:3 2.5%
- PC 18:2_18:2 18.8%

and of (not quantified):
- PC O−17:0_20:4
- and further compounds

### Linear model

```
PC aa C36:4 ~ b * ( PC 14:0_22:4
   + PC 16:0_20:4
   + PC 18:1_18:3
   + PC 18:2_18:2 )
```

b = 2.55262

$R^2$ = 0.65155

### Ranges

| Measure | AbsoluteIDQ | sum(Lipidyzer) | delta |
|---|---|---|---|
| Min | 101.4 | 47.02 | 54.38 |
| Max | 290.2 | 98.75 | 191.45 |
| Mean | 180.31 | 70.22 | 110.09 |
| Median | 179.9 | 69.92 | 109.98 |
| SD | 35.18 | 10.41 | 24.76 |

### Composition: mean of proportions q

conc(PC aa C36:4) * 0.0016 = conc(PC 14:0_22:4) [var(q)=0.0016]
Percentiles: 5%−>0.0011, 25%−>0.0013, 75%−>0.0018, 95%−>0.0025
conc(PC aa C36:4) * 0.2943 = conc(PC 16:0_20:4) [var(q)=0.2943]
Percentiles: 5%−>0.2393, 25%−>0.2658, 75%−>0.317, 95%−>0.3555
conc(PC aa C36:4) * 0.0109 = conc(PC 18:1_18:3) [var(q)=0.0109]
Percentiles: 5%−>0.0033, 25%−>0.0056, 75%−>0.0147, 95%−>0.022
conc(PC aa C36:4) * 0.0774 = conc(PC 18:2_18:2) [var(q)=0.0774]
Percentiles: 5%−>0.0354, 25%−>0.0496, 75%−>0.0957, 95%−>0.1558

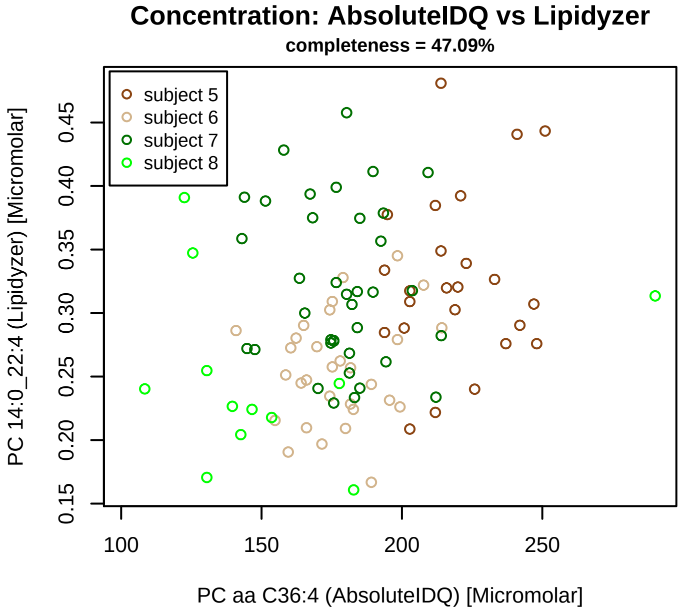


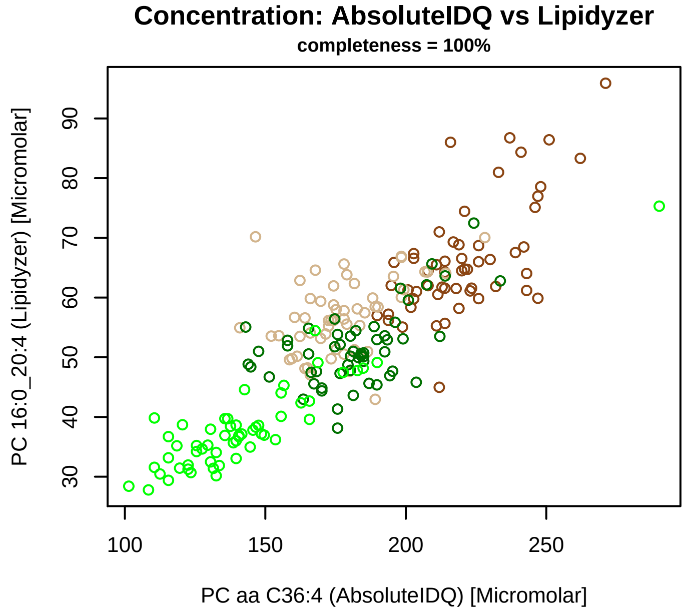


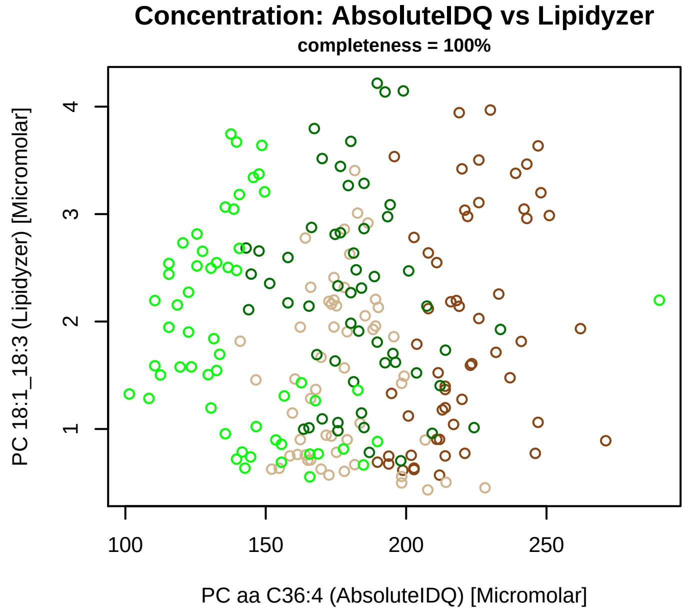


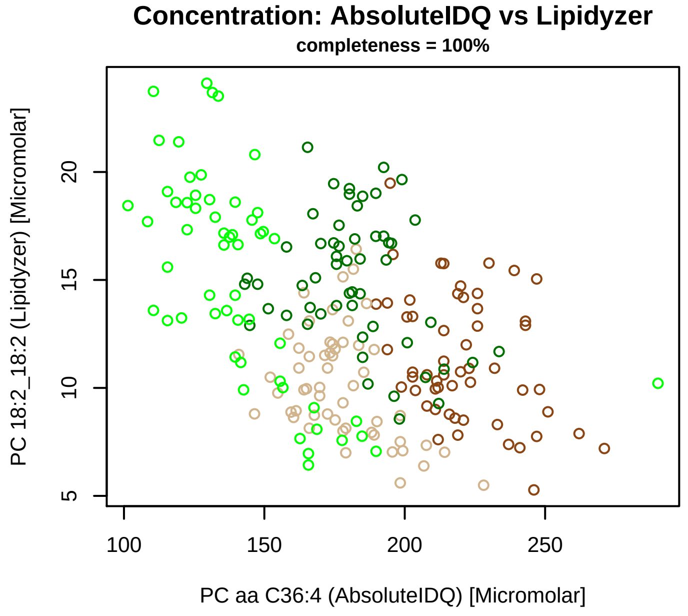


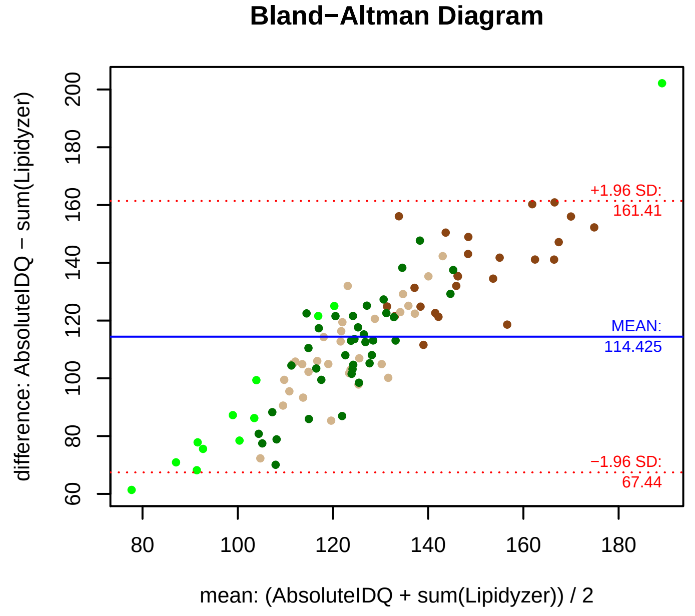


### Stability of composition

**(PC 14:0_22:4 + PC 16:0_20:4 + PC 18:1_18:3 + PC 18:2_18:2) / PC aa C36:4**
**Shapiro−Wilk Test of log quotients, pv: 0.91369; OK**
SW: ok; ANOVA: Comp. ~ Subject_ID −> pv = 0.21225; EQUAL
Wilcoxon: Challenge; sport: 0.5; OLTT: 1

**PC 14:0_22:4 / PC aa C36:4**
**Shapiro−Wilk Test of log quotients, pv: 0.48012; OK**
SW: ok; ANOVA: Comp. ~ Subject_ID −> pv = 0.00455; DIFFERENCE
Wilcoxon: Challenge; sport: 0.5; OLTT: 1

**PC 16:0_20:4 / PC aa C36:4**
**Shapiro−Wilk Test of log quotients, pv: 0.22093; OK**
SW: no; Kruskal: Comp. ~ Subject_ID −> pv = 0; DIFFERENCE
Wilcoxon: Challenge; fasting: 0.875; sport: 0.25; OLTT: 0.25

**PC 18:1_18:3 / PC aa C36:4**
**Shapiro−Wilk Test of log quotients, pv: 2e−05; NO**
SW: no; Kruskal: Comp. ~ Subject_ID −> pv = 0; DIFFERENCE
Wilcoxon: Challenge; fasting: 0.125; sport: 1; OLTT: 0.625

**PC 18:2_18:2 / PC aa C36:4**
**Shapiro−Wilk Test of log quotients, pv: 0.52175; OK**
SW: no; Kruskal: Comp. ~ Subject_ID −> pv = 0; DIFFERENCE
Wilcoxon: Challenge; fasting: 0.125; sport: 0.125; OLTT: 0.625

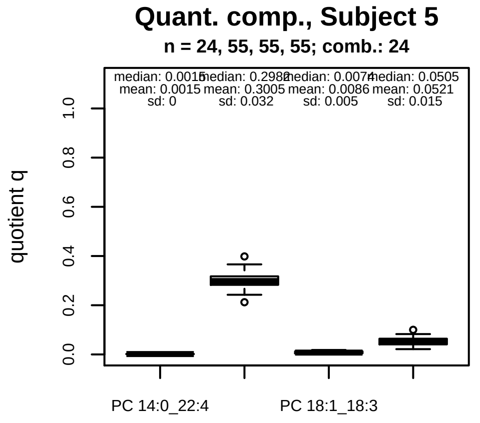


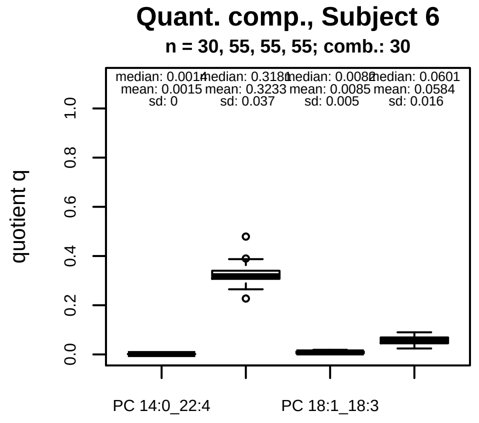


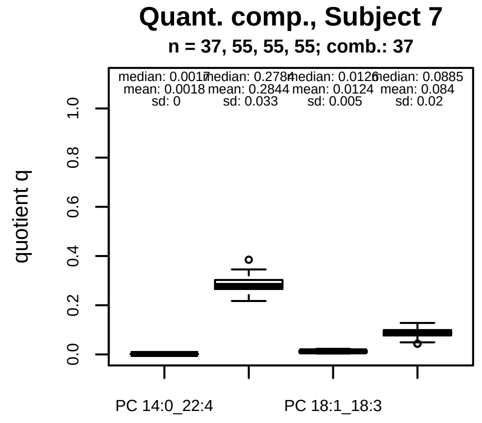


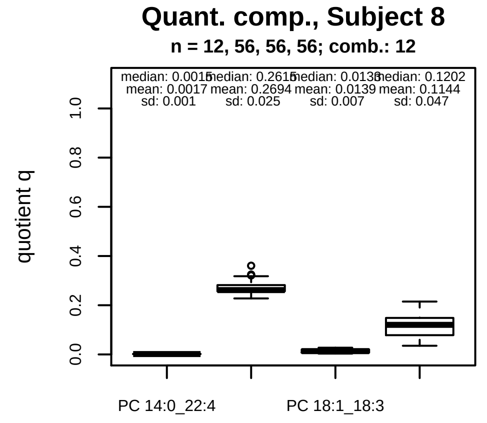


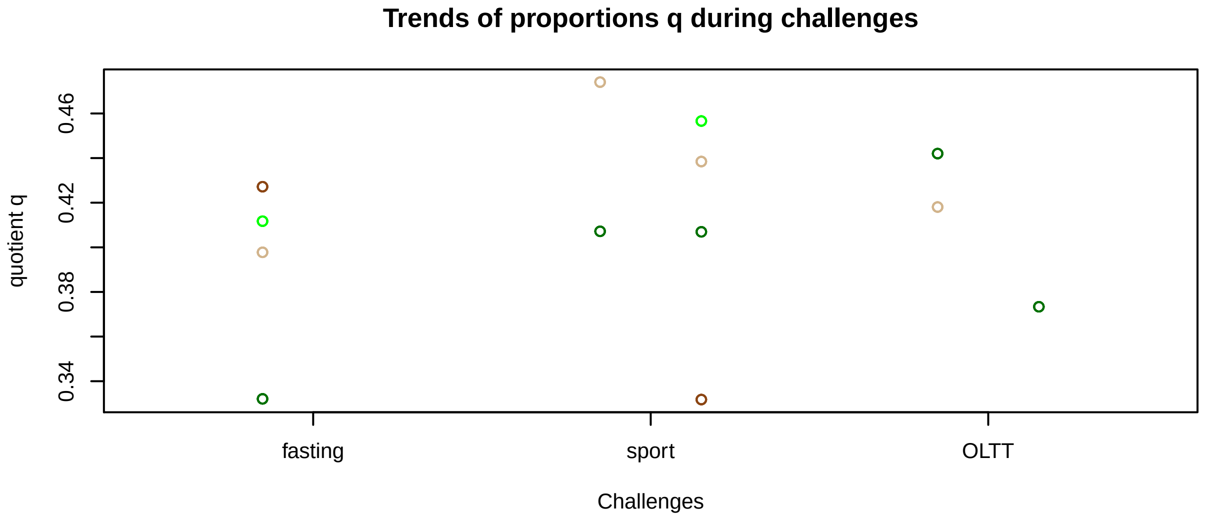


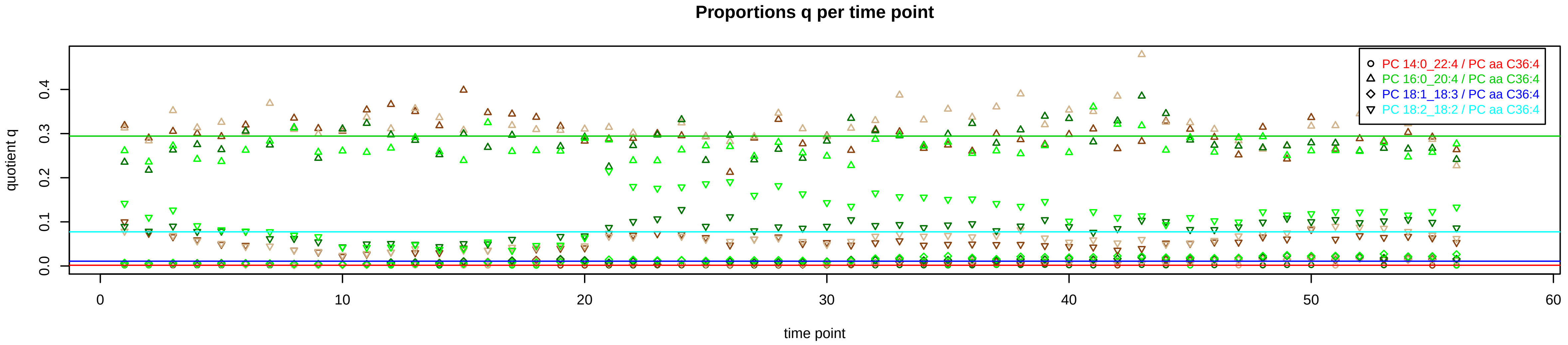

# PC aa C36:5 = PC 14:0_22:5 + PC 16:0_20:5 + PC 18:2_18:3 + R

PC 16:0_20:5 excluded because of missingness > 75%

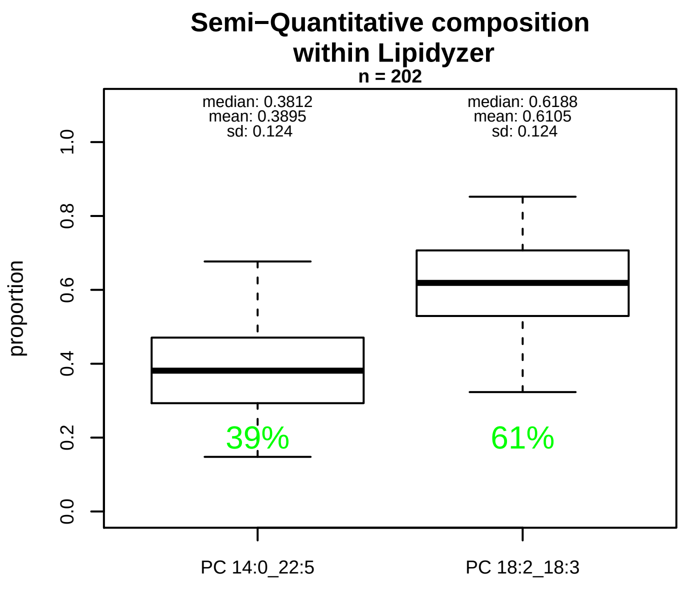


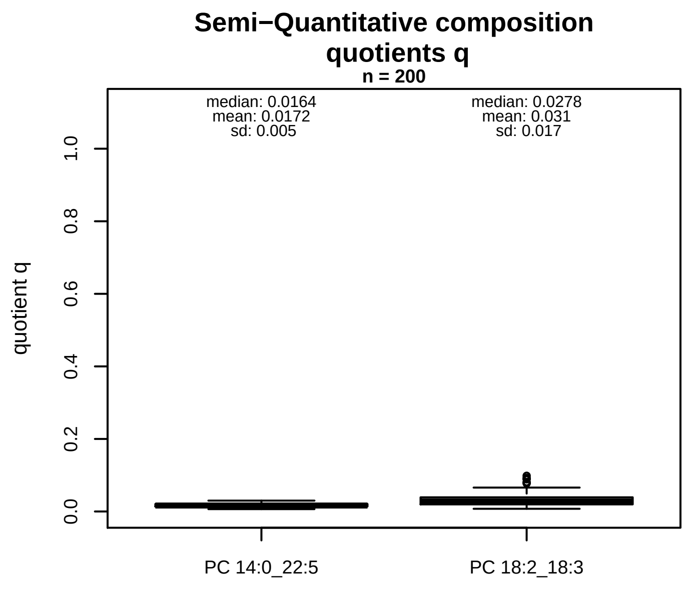


### Semi−Quantitative composition: Lipidyzer

PC aa C36:5 consists of:
PC 14:0_22:5 39%
PC 18:2_18:3 61%
and of (not quantified):
PC 16:0_20:5, PC 14 1_22:4, PC 16 1_20:4, PC 18 1_18:4,
and further compounds

### Linear model

```
PC aa C36:5 ~ b * ( PC 14:0_22:5
   + PC 18:2_18:3 )
```

b = 9.52537

$R^2$ = 0.17224

### Ranges

| Measure | AbsoluteIDQ | sum(Lipidyzer) | delta |
|---|---|---|---|
| Min | 12.6 | 0.49 | 12.11 |
| Max | 50.5 | 2.12 | 48.38 |
| Mean | 27.44 | 1.24 | 26.2 |
| Median | 28.7 | 1.24 | 27.46 |
| SD | 9.37 | 0.38 | 8.99 |

### Composition: mean of proportions q

conc(PC aa C36:5) * 0.0172 = conc(PC 14:0_22:5) [var(q)=0.0172]
Percentiles: 5%−>0.0101, 25%−>0.0138, 75%−>0.0206, 95%−>0.0262
conc(PC aa C36:5) * 0.031 = conc(PC 18:2_18:3) [var(q)=0.031]
Percentiles: 5%−>0.0113, 25%−>0.0196, 75%−>0.0391, 95%−>0.0615

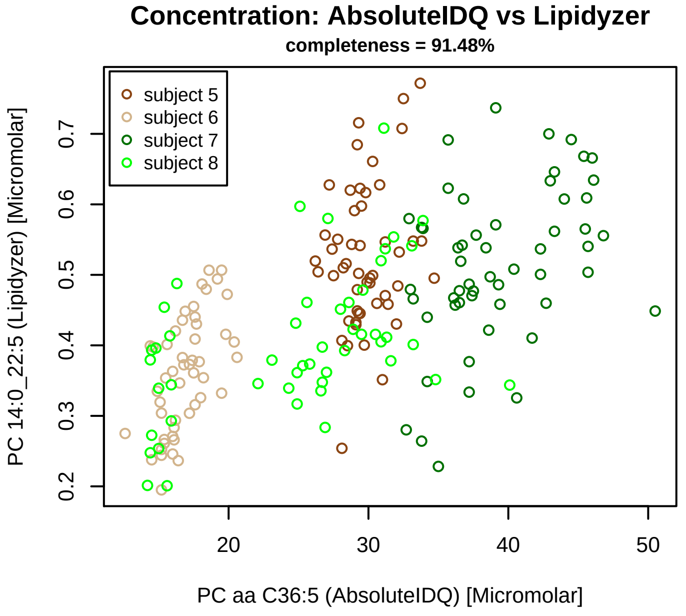


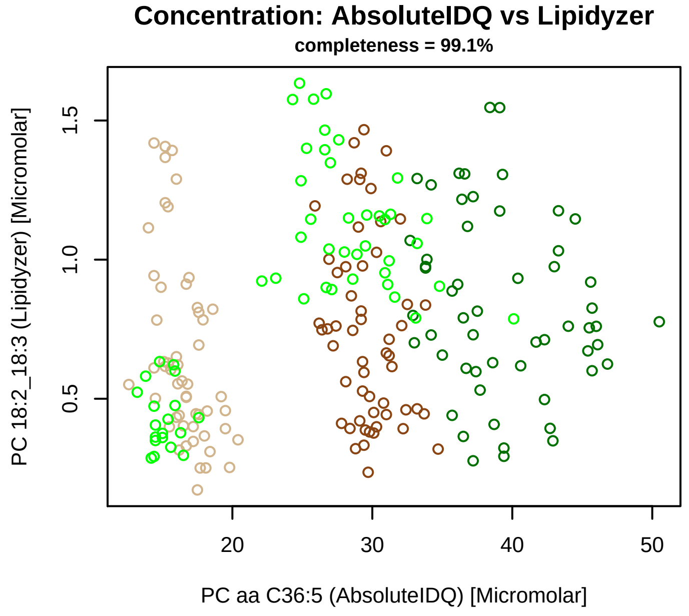


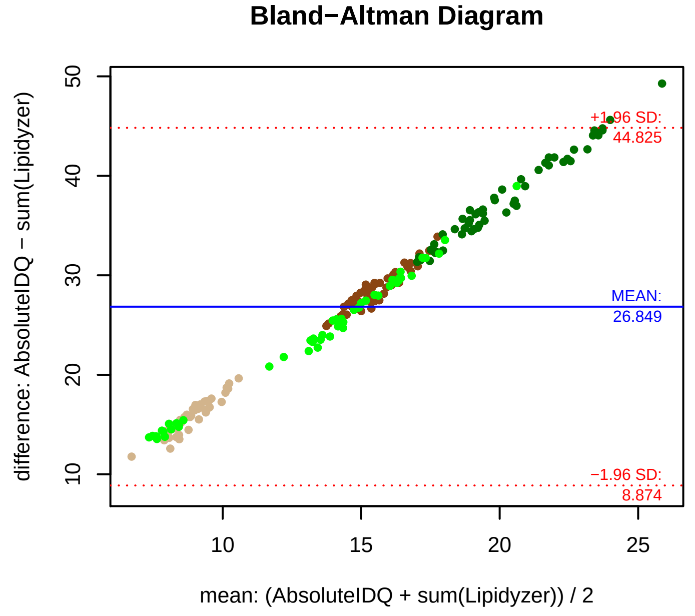


### Stability of composition

**(PC 14:0_22:5 + PC 18:2_18:3) / PC aa C36:5**
**Shapiro−Wilk Test of log quotients, pv: 0.49031; OK**
SW: ok; ANOVA: Comp. ~ Subject_ID −> pv = 0.43298; EQUAL
Wilcoxon: Challenge; fasting: 1; sport: 0.125; OLTT: 0.5

**PC 18:2_18:3 / PC aa C36:5**
**Shapiro−Wilk Test of log quotients, pv: 0.25945; OK**
SW: ok; ANOVA: Comp. ~ Subject_ID −> pv = 0.0212; EQUAL
Wilcoxon: Challenge; fasting: 0.25; sport: 0.125; OLTT: 0.125

**PC 14:0_22:5 / PC aa C36:5**
**Shapiro−Wilk Test of log quotients, pv: 0.03493; OK**
SW: ok; ANOVA: Comp. ~ Subject_ID −> pv = 2e−04; DIFFERENCE
Wilcoxon: Challenge; fasting: 1; sport: 0.625; OLTT: 1

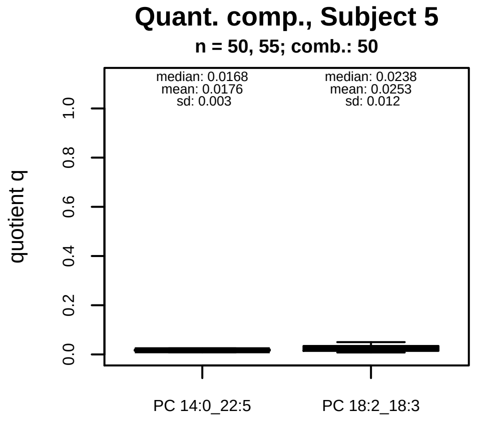


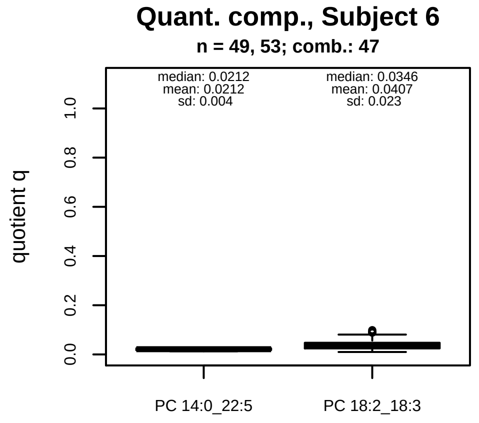


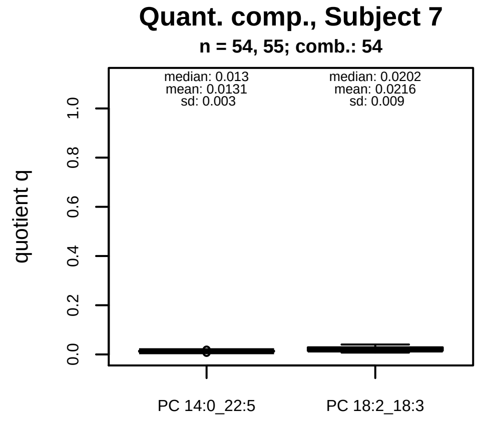


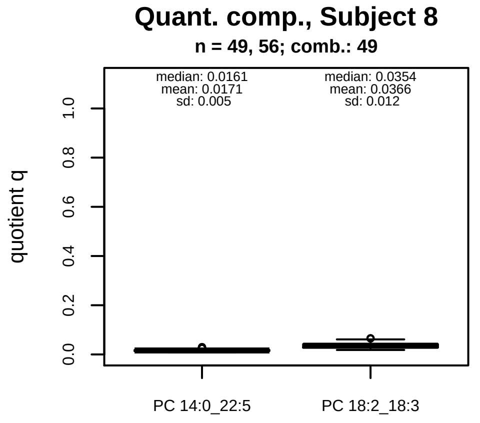


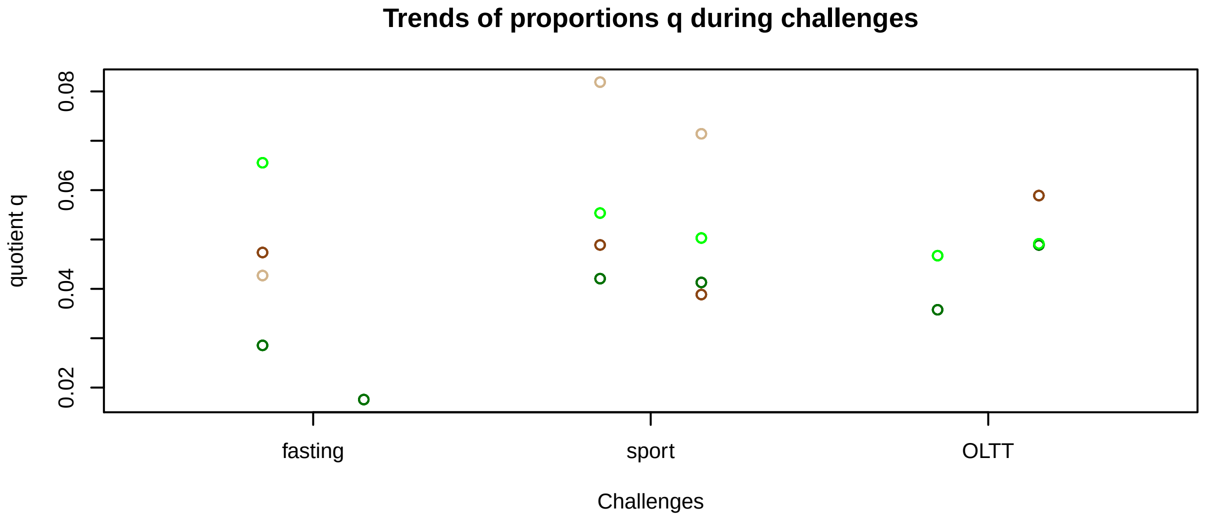


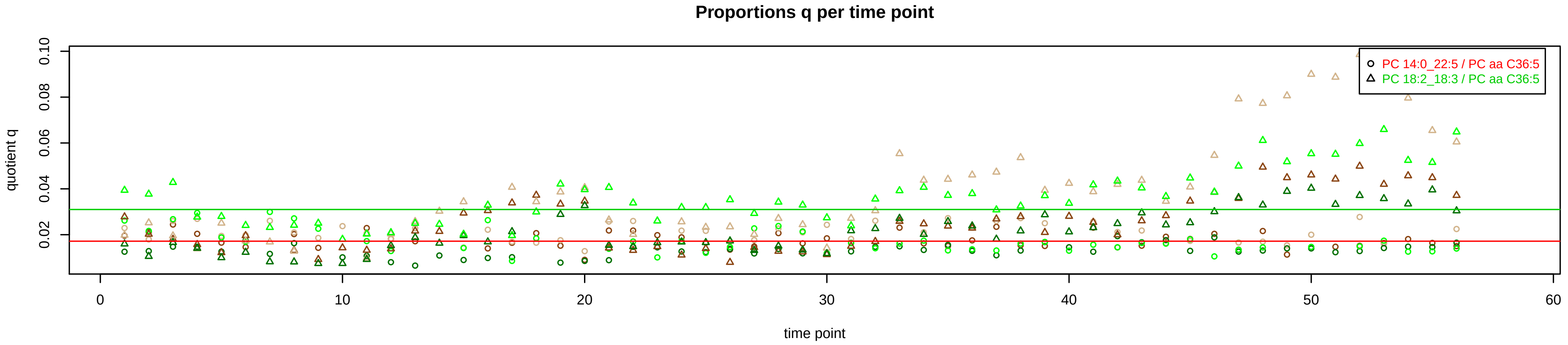

# PC aa C36:6 = PC 14:0_22:6 + R

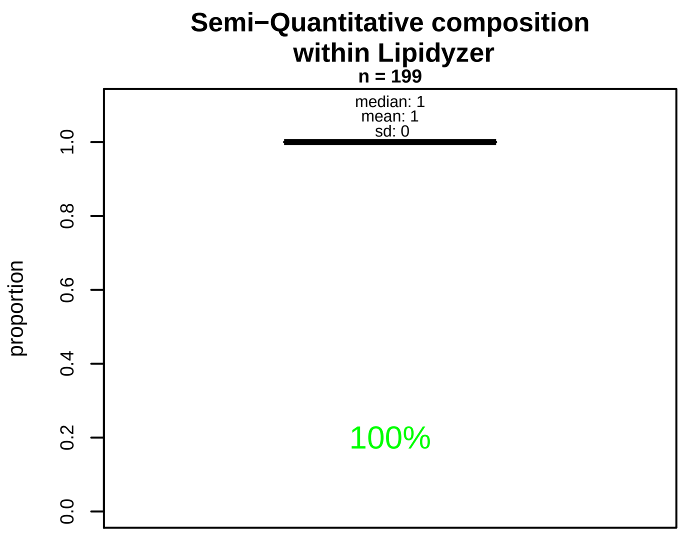


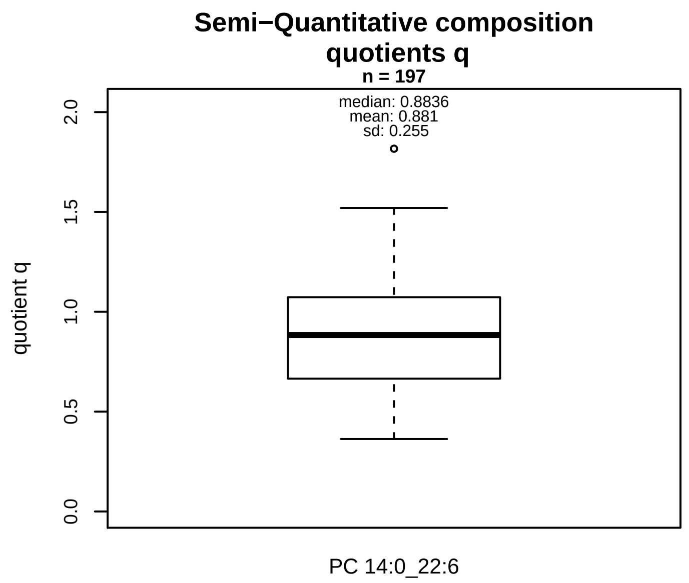


### Semi−Quantitative composition: Lipidyzer

PC aa C36:6 consists of:
PC 14:0_22:6 100%
and of (not quantified):
PC 14:1_22:5, PC 16:1_20:5, PC 18:3_18:3, PC 18:4_18:2,
and further compounds

### Linear model

```
PC aa C36:6 ~ b * ( PC 14:0_22:6 )
```

b = 0.74555

$R^2$ = 0.32274

### Ranges

| Measure | AbsoluteIDQ | sum(Lipidyzer) | delta |
|---|---|---|---|
| Min | 0.39 | 0.27 | 0.12 |
| Max | 1.69 | 1.67 | 0.02 |
| Mean | 0.91 | 0.79 | 0.13 |
| Median | 0.9 | 0.76 | 0.14 |
| SD | 0.33 | 0.26 | 0.08 |

### Composition: mean of proportions q

conc(PC aa C36:6) * 0.881 = conc(PC 14:0_22:6) [var(q)=0.881]
Percentiles: 5%−>0.4947, 25%−>0.6651, 75%−>1.0732, 95%−>1.2925

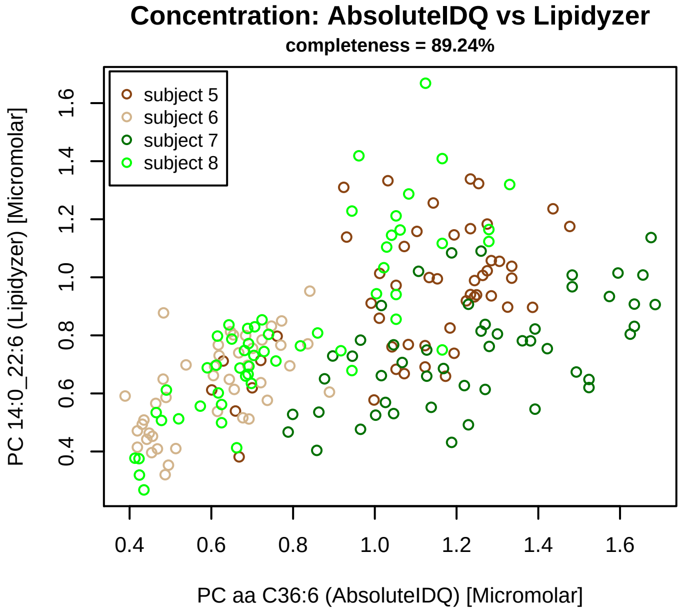


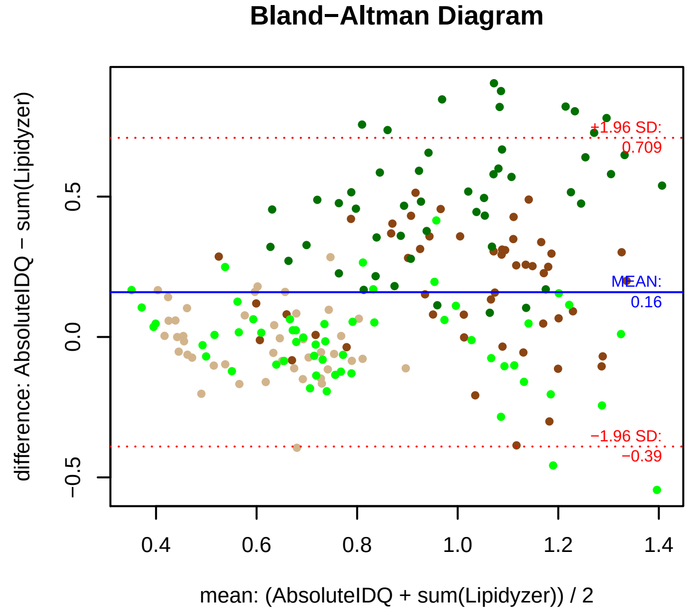


### Stability of composition

**PC 14:0_22:6 / PC aa C36:6**
**Shapiro−Wilk Test of log quotients, pv: 0.00996; NO**
SW: ok; ANOVA: Comp. ~ Subject_ID −> pv = 0.68751; EQUAL
Wilcoxon: Challenge; fasting: 0.25; sport: 0.625; OLTT: 0.25

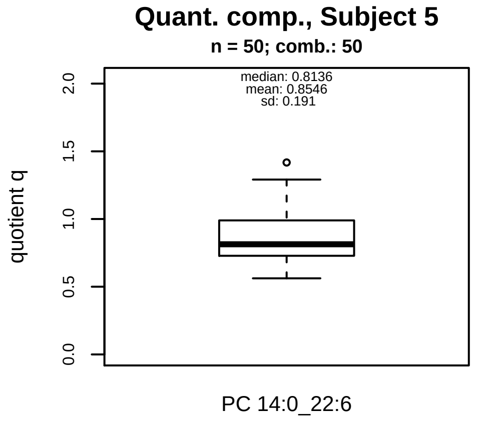


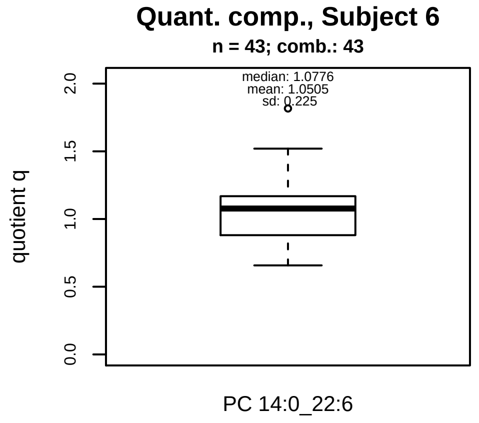


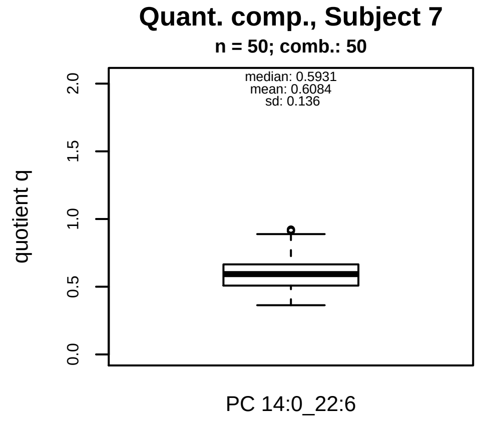


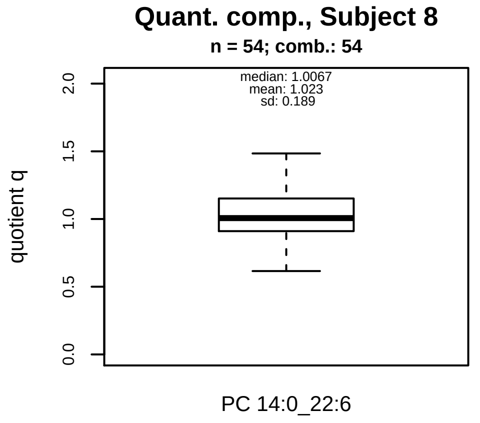


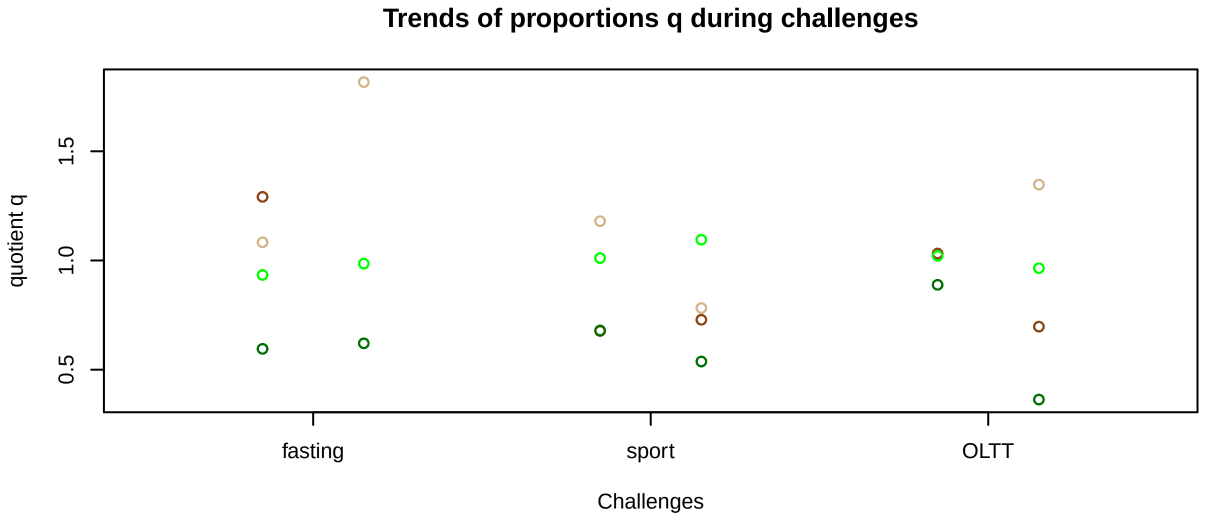


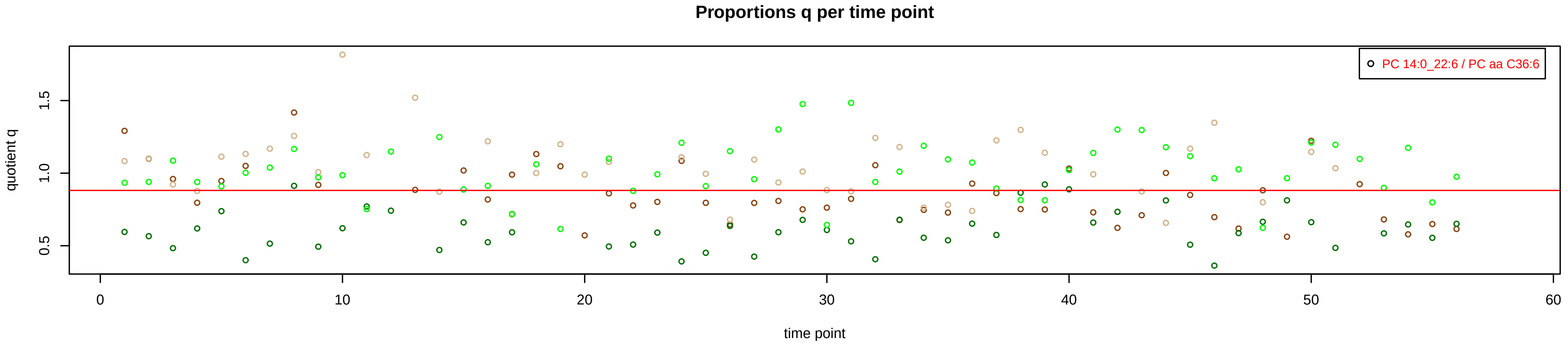

# PC aa C38:0 = PC 18:0_20:0 + R

#### Semi−Quantitative composition within Lipidyzer

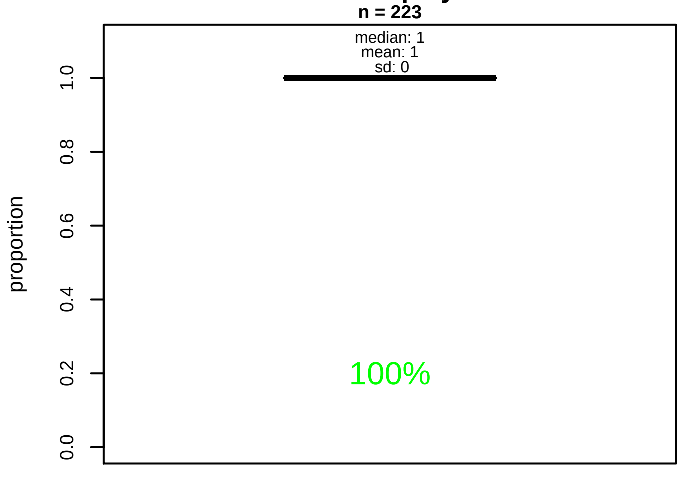


#### Semi−Quantitative composition quotients q

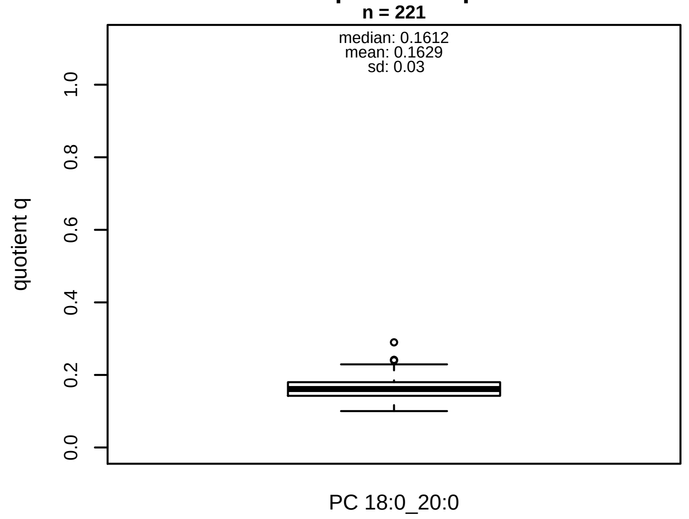


#### Semi−Quantitative composition: Lipidyzer

PC aa C38:0 consists of:
PC 18:0_20:0 100%
and of (not quantified):
PC 12:0_26:0, PC 13:0_25:0, PC 14:0_24:0, PC 16:0_22:0,
PC 17:0_21:0, PC 17:1_22:6, PC 19:0_19:0, PC O−17:0_22:(
PC O−18:0_21:0, PC O−20:0_19:0, PC O−18:1_22:6, SM42.(
and further compounds

#### Composition: mean of proportions q

conc(PC aa C38:0) * 0.1629 = conc(PC 18:0_20:0) [var(q)=0.1629]
Percentiles: 5%−>0.1193, 25%−>0.1424, 75%−>0.18, 95%−>0.2153

#### Linear model

```
PC aa C38:0 ~ b * ( PC 18:0_20:0 )
```

b = 3.58022

$R^2 = 0.34478$

#### Ranges

| Measure | AbsoluteIDQ | sum(Lipidyzer) | delta |
|---|---|---|---|
| Min | 2.33 | 0.35 | 1.98 |
| Max | 5.48 | 0.92 | 4.56 |
| Mean | 3.74 | 0.6 | 3.14 |
| Median | 3.66 | 0.59 | 3.07 |
| SD | 0.71 | 0.12 | 0.6 |

#### Concentration: AbsoluteIDQ vs Lipidyzer

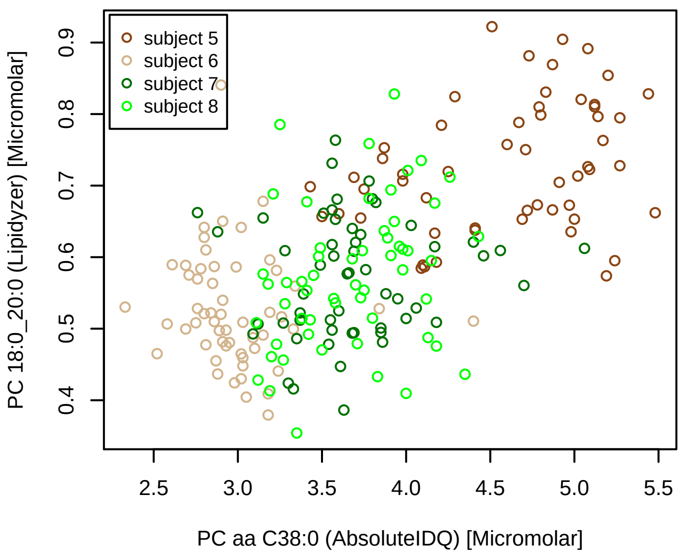


#### Bland−Altman Diagram

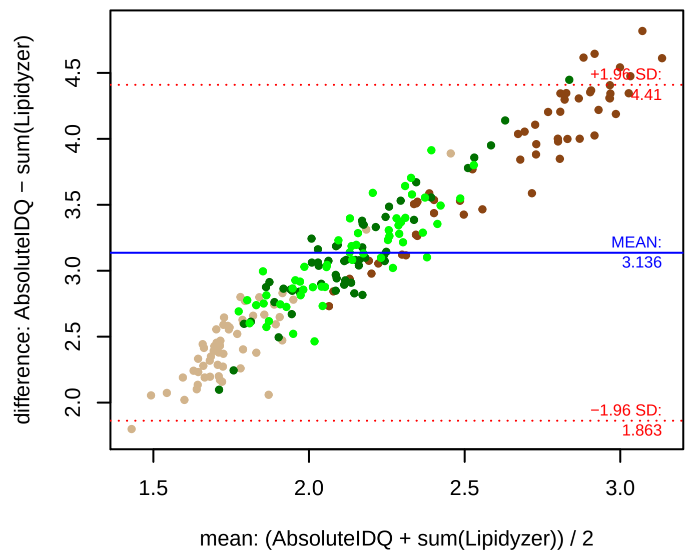


#### Stability of composition

**PC 18:0_20:0 / PC aa C38:0**
**Shapiro−Wilk Test of log quotients, pv: 0.94774; OK**
SW: ok; ANOVA: Comp. ~ Subject_ID −> pv = 0.05256; EQUAL
Wilcoxon: Challenge; fasting: 0.625; sport: 0.25; OLTT: 1

#### Quant. comp., Subject 5

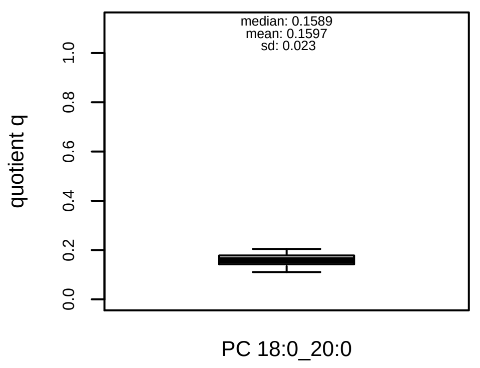


#### Quant. comp., Subject 6

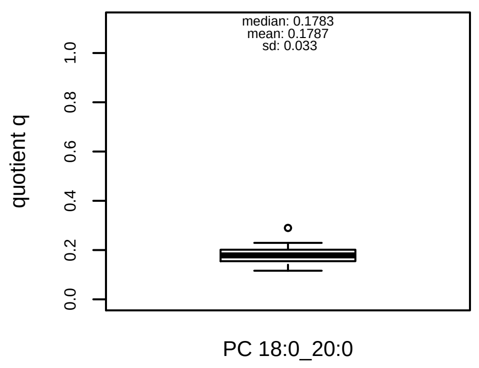


#### Quant. comp., Subject 7

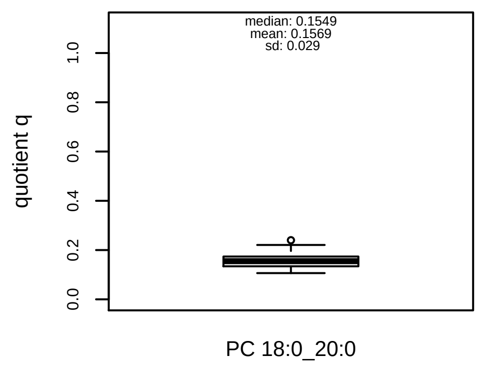


#### Quant. comp., Subject 8

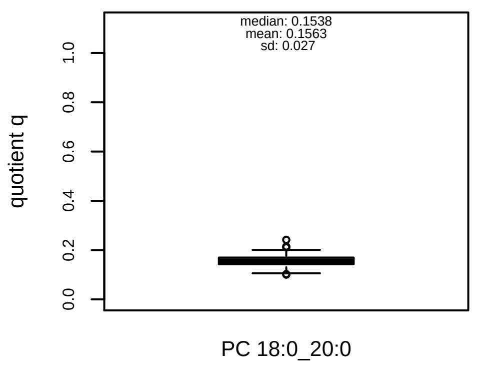


#### Trends of proportions q during challenges

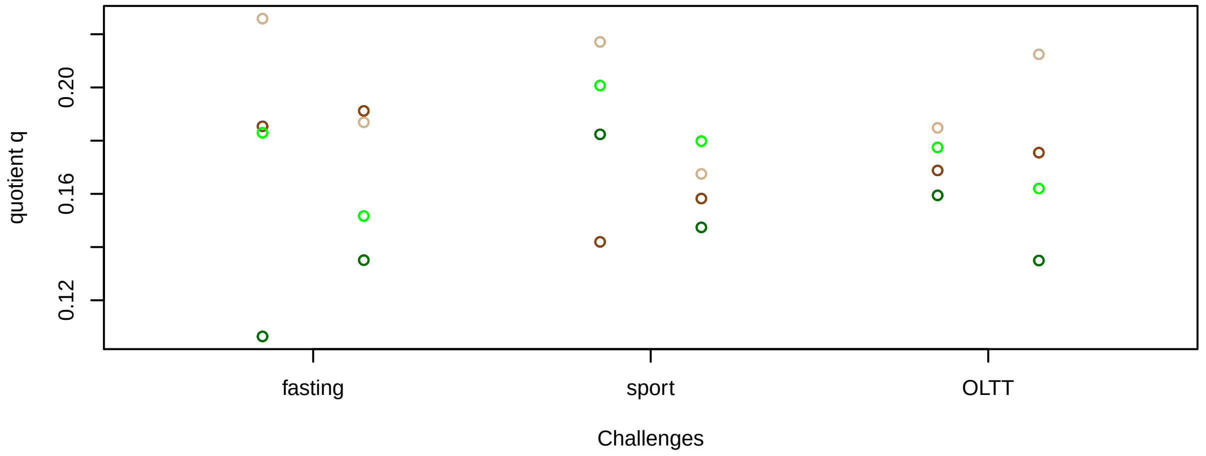


#### Proportions q per time point

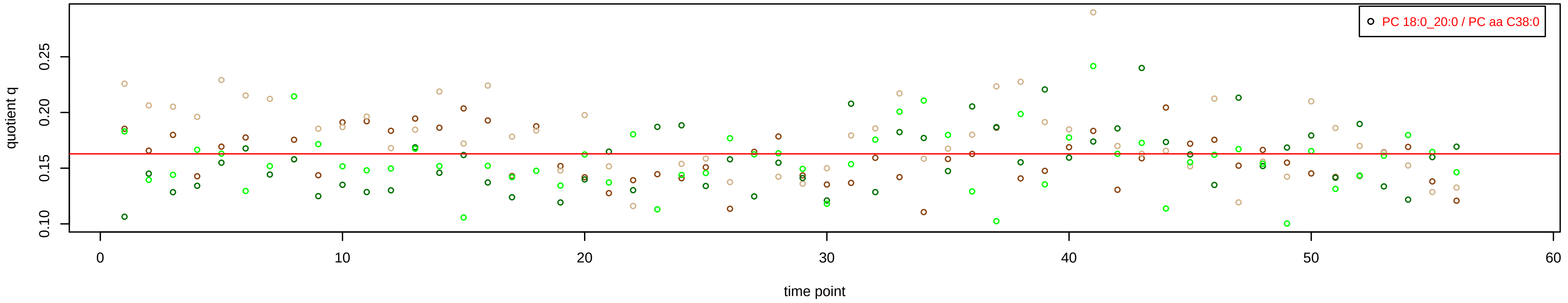

# PC aa C38:3 = PC 18:0_20:3 + PC 18:1_20:2 + PC 18:2_20:1 + PC 20:0_18:3 + R

PC 20:0_18:3 excluded because of missingness > 75%

**Semi−Quantitative composition within Lipidyzer**
n = 221

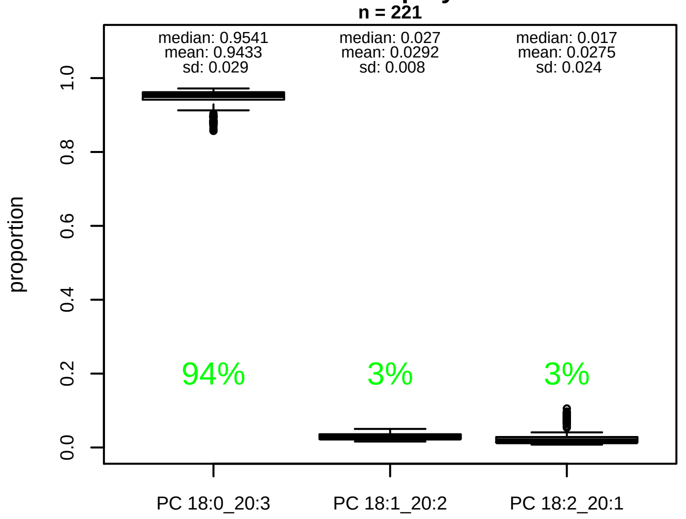


**Semi−Quantitative composition quotients q**
n = 219

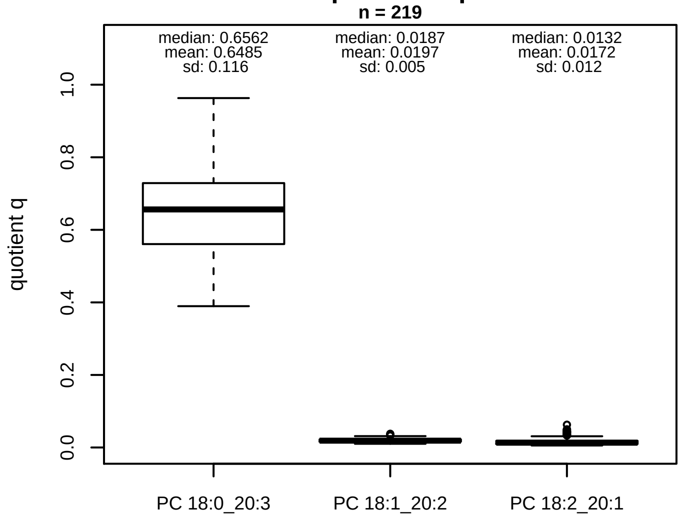


### Semi−Quantitative composition: Lipidyzer

PC aa C38:3 consists of:
- PC 18:0_20:3 94.3%
- PC 18:1_20:2 2.9%
- PC 18:2_20:1 2.7%

and of (not quantified):
PC 20:0_18:3, PC 16:1_22:2, PC 18:3_20:0, [13C1]SM 42:3, and further compounds

### Linear model

```
PC aa C38:3 ~ b * ( PC 18:0_20:3
   + PC 18:1_20:2
   + PC 18:2_20:1 )
```

b = 0.98751

$R^2$ = 0.80663

### Ranges

| Measure | AbsoluteIDQ | sum(Lipidyzer) | delta |
|---|---|---|---|
| Min | 25.7 | 13.72 | 11.98 |
| Max | 66.8 | 54.35 | 12.45 |
| Mean | 44.46 | 30.82 | 13.64 |
| Median | 42.4 | 30.43 | 11.97 |
| SD | 11.16 | 10.15 | 1.01 |

### Composition: mean of proportions q

conc(PC aa C38:3) * 0.6485 = conc(PC 18:0_20:3) [var(q)=0.6485]
Percentiles: 5%−>0.4476, 25%−>0.5606, 75%−>0.7287, 95%−>0.8496
conc(PC aa C38:3) * 0.0197 = conc(PC 18:1_20:2) [var(q)=0.0197]
Percentiles: 5%−>0.0126, 25%−>0.0161, 75%−>0.0224, 95%−>0.0299
conc(PC aa C38:3) * 0.0172 = conc(PC 18:2_20:1) [var(q)=0.0172]
Percentiles: 5%−>0.007, 25%−>0.0096, 75%−>0.0184, 95%−>0.0436

**Concentration: AbsoluteIDQ vs Lipidyzer**
completeness = 100%

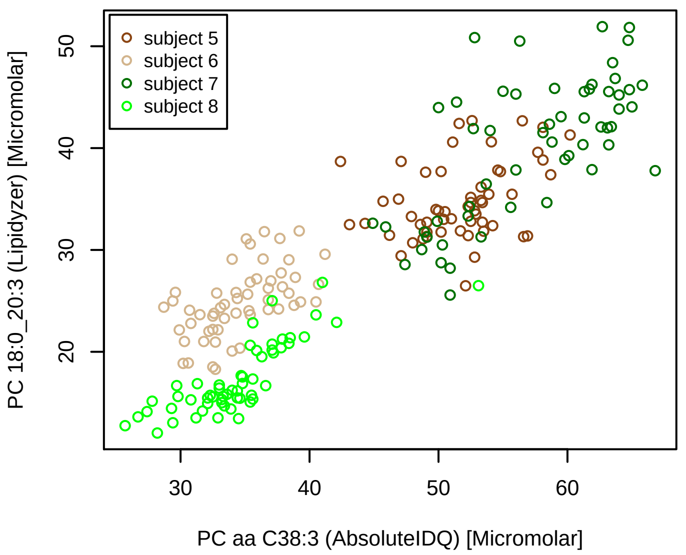


**Concentration: AbsoluteIDQ vs Lipidyzer**
completeness = 100%

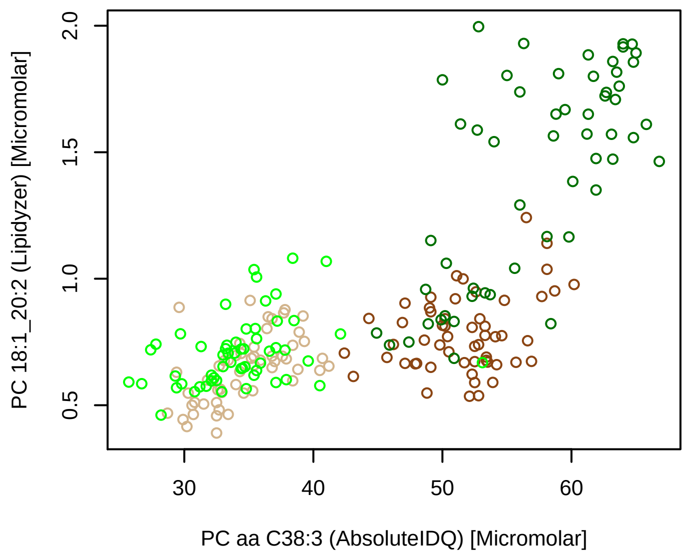


**Concentration: AbsoluteIDQ vs Lipidyzer**
completeness = 99.1%

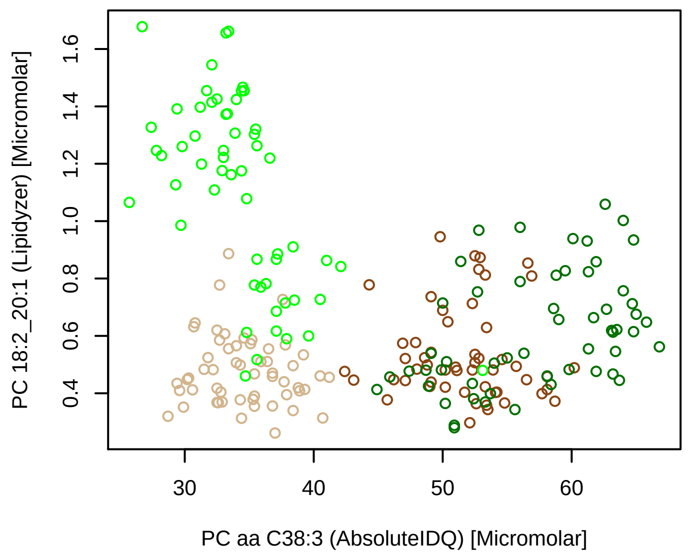


**Bland−Altman Diagram**

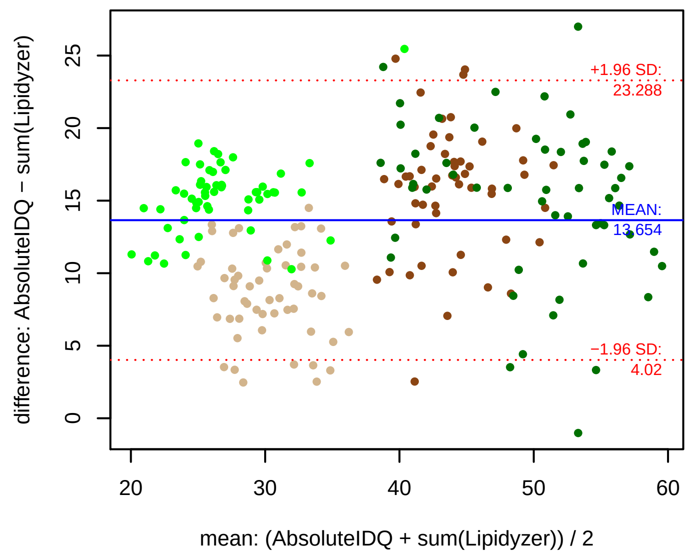


### Stability of composition

**(PC 18:0_20:3 + PC 18:1_20:2 + PC 18:2_20:1) / PC aa C38:3**
**Shapiro−Wilk Test of log quotients, pv: 0.06972; OK**
SW: ok; ANOVA: Comp. ~ Subject_ID −> pv = 0; DIFFERENCE
Wilcoxon: Challenge; fasting: 0.5; sport: 0.125; OLTT: 0.25

**PC 18:1_20:2 / PC aa C38:3**
**Shapiro−Wilk Test of log quotients, pv: 0.11436; OK**
SW: ok; ANOVA: Comp. ~ Subject_ID −> pv = 0; DIFFERENCE
Wilcoxon: Challenge; fasting: 0.125; sport: 1; OLTT: 0.375

**PC 18:0_20:3 / PC aa C38:3**
**Shapiro−Wilk Test of log quotients, pv: 2e−04; NO**
SW: ok; ANOVA: Comp. ~ Subject_ID −> pv = 0; DIFFERENCE
Wilcoxon: Challenge; fasting: 0.25; sport: 0.125; OLTT: 0.25

**PC 18:2_20:1 / PC aa C38:3**
**Shapiro−Wilk Test of log quotients, pv: 0; NO**
SW: no; Kruskal: Comp. ~ Subject_ID −> pv = 0; DIFFERENCE
Wilcoxon: Challenge; fasting: 0.5; sport: 0.25; OLTT: 0.875

**Quant. comp., Subject 5**
n = 55, 55, 53; comb.: 53

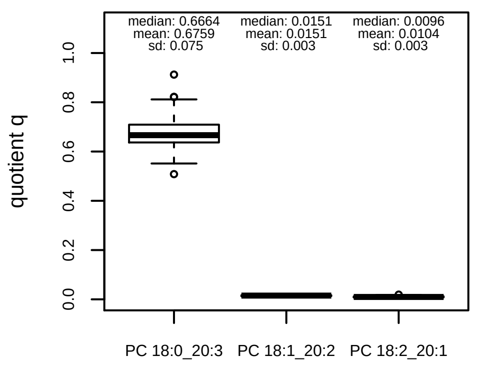


**Quant. comp., Subject 6**
n = 55, 55, 55; comb.: 55

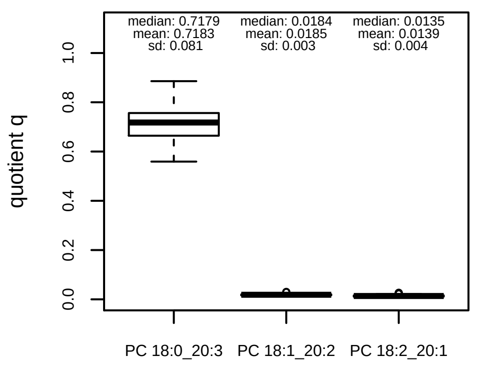


**Quant. comp., Subject 7**
n = 55, 55, 55; comb.: 55

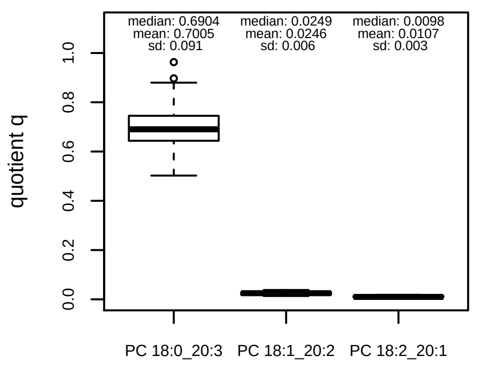


**Quant. comp., Subject 8**
n = 56, 56, 56; comb.: 56

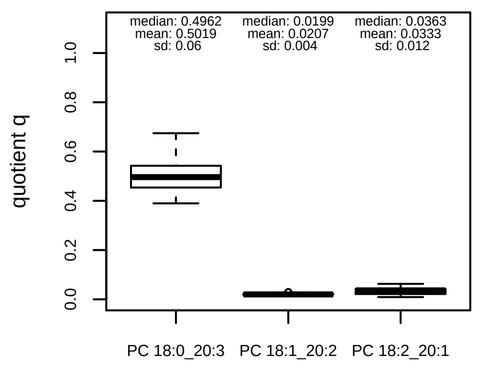


**Trends of proportions q during challenges**

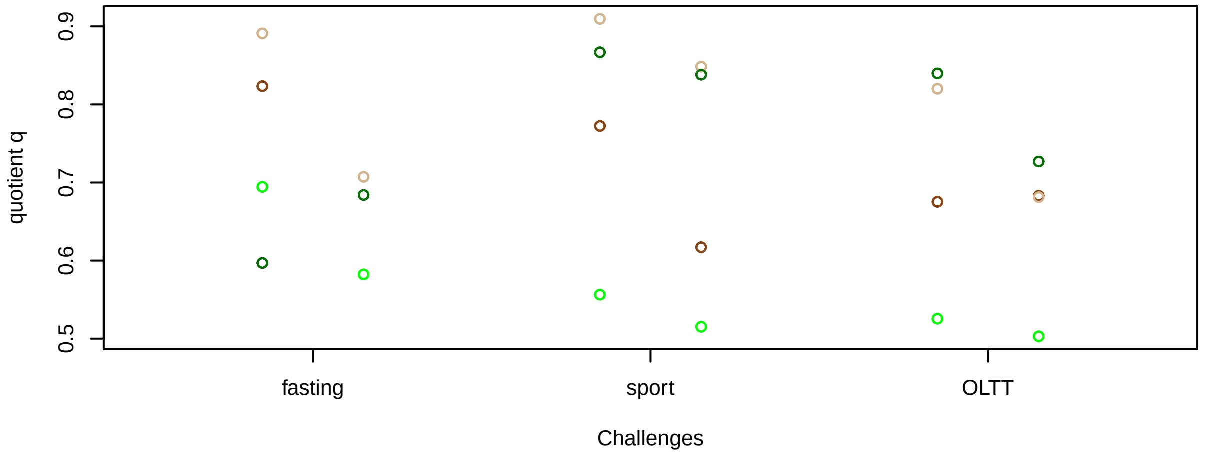


**Proportions q per time point**

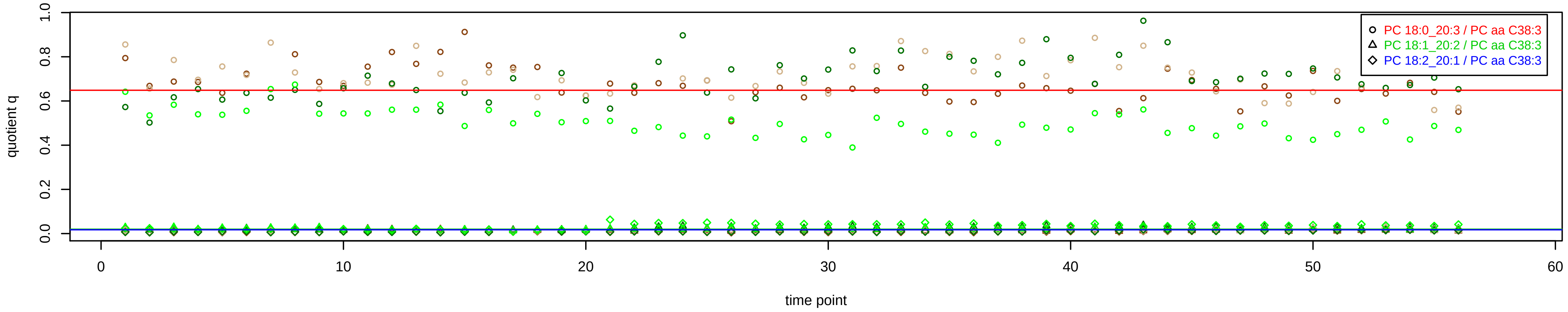

# PC aa C38:4 = PC 16:0_22:4 + PC 18:0_20:4 + PC 18:1_20:3 + PC 18:2_20:2 + R

#### Semi−Quantitative composition within Lipidyzer
n = 222

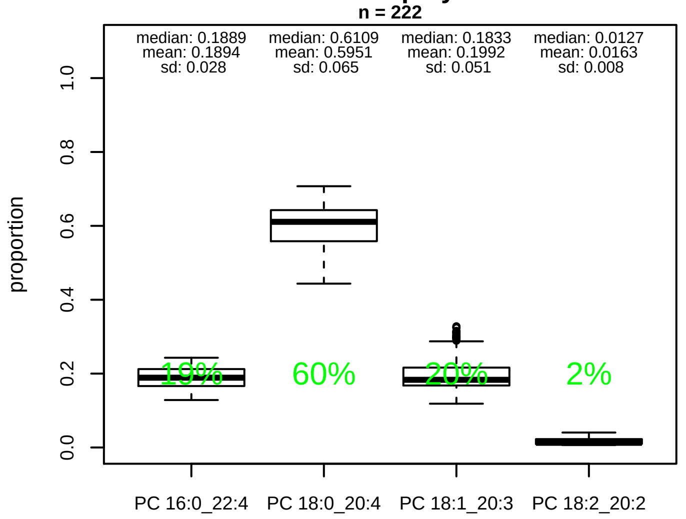


#### Semi−Quantitative composition quotients q
n = 220

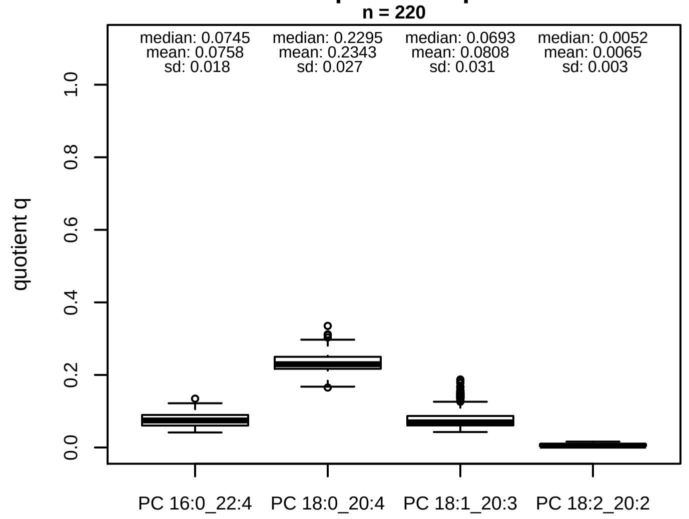


#### Semi−Quantitative composition: Lipidyzer

PC aa C38:4 consists of:
PC 16:0_22:4 18.9%
PC 18:0_20:4 59.5%
PC 18:1_20:3 19.9%
PC 18:2_20:2 1.6%
and of (not quantified):
PC 18:4_20:0
and further compounds

#### Linear model

```
PC aa C38:4 ~ b * ( PC 16:0_22:4
   + PC 18:0_20:4
   + PC 18:1_20:3
   + PC 18:2_20:2 )
```

b = 1.99845

$R^2$ = 0.58796

#### Ranges

| Measure | AbsoluteIDQ | sum(Lipidyzer) | delta |
|---|---|---|---|
| Min | 54 | 19.61 | 34.39 |
| Max | 153 | 61.31 | 91.69 |
| Mean | 99.43 | 39.14 | 60.29 |
| Median | 95.5 | 40.09 | 55.41 |
| SD | 21.26 | 8.1 | 13.16 |

#### Composition: mean of proportions q

conc(PC aa C38:4) * 0.0758 = conc(PC 16:0_22:4) [var(q)=0.0758]
Percentiles: 5%−>0.0504, 25%−>0.0605, 75%−>0.0897, 95%−>0.1059
conc(PC aa C38:4) * 0.2343 = conc(PC 18:0_20:4) [var(q)=0.2343]
Percentiles: 5%−>0.1971, 25%−>0.217, 75%−>0.2499, 95%−>0.2867
conc(PC aa C38:4) * 0.0808 = conc(PC 18:1_20:3) [var(q)=0.0808]
Percentiles: 5%−>0.049, 25%−>0.0608, 75%−>0.0869, 95%−>0.15
conc(PC aa C38:4) * 0.0065 = conc(PC 18:2_20:2) [var(q)=0.0065]
Percentiles: 5%−>0.0028, 25%−>0.0037, 75%−>0.0094, 95%−>0.0126

#### Concentration: AbsoluteIDQ vs Lipidyzer
completeness = 100%

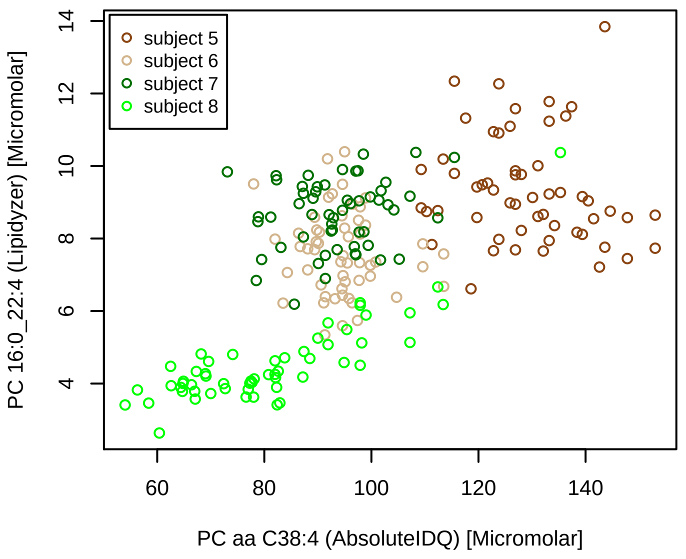


#### Concentration: AbsoluteIDQ vs Lipidyzer
completeness = 100%

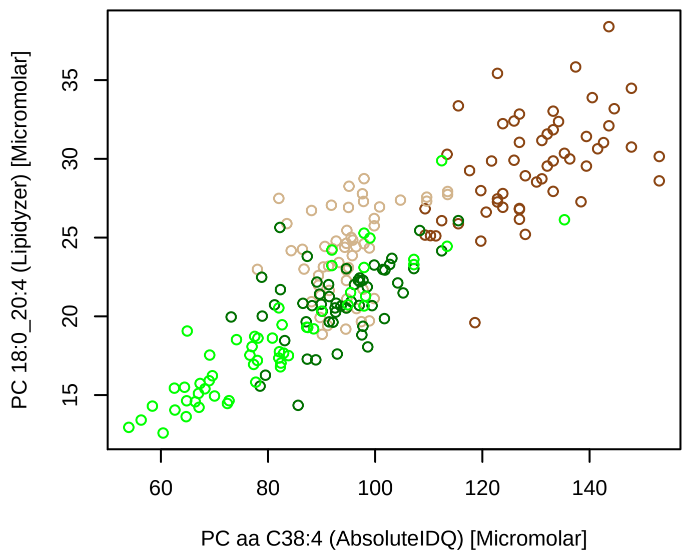


#### Concentration: AbsoluteIDQ vs Lipidyzer
completeness = 100%

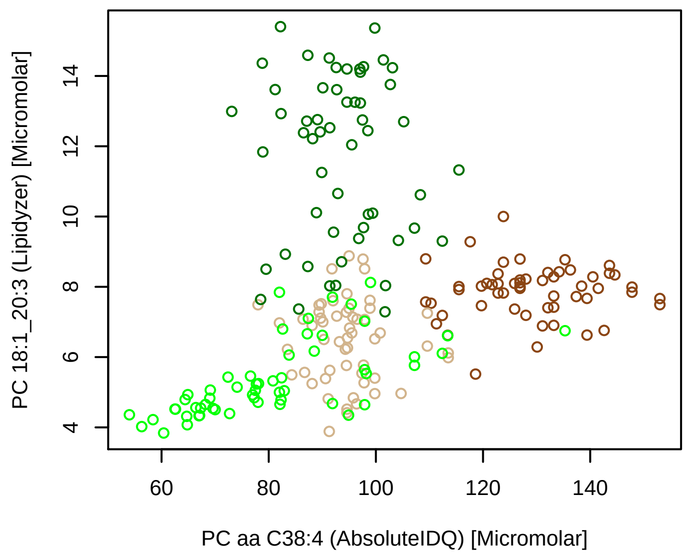


#### Concentration: AbsoluteIDQ vs Lipidyzer
completeness = 99.55%

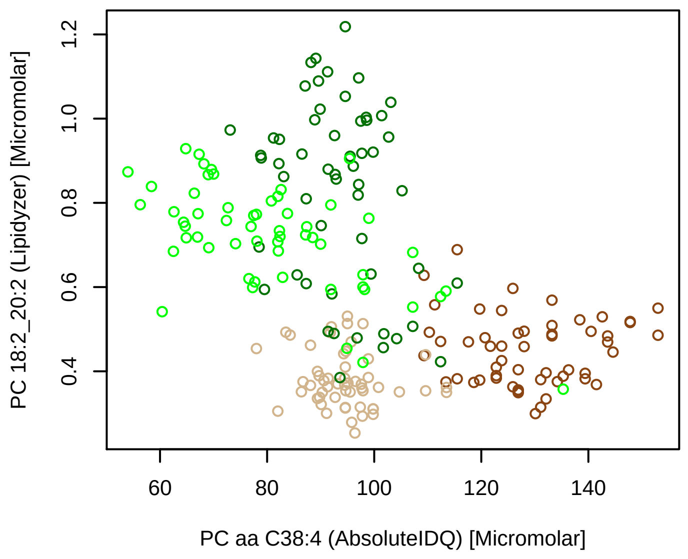


#### Bland−Altman Diagram

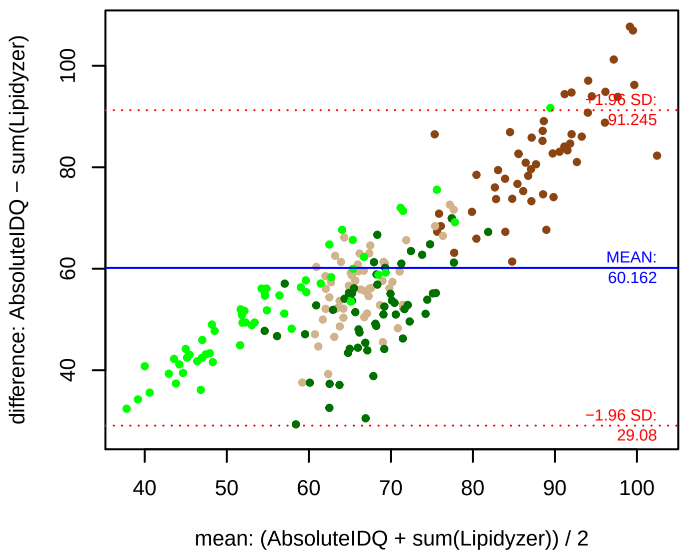


#### Stability of composition

**(PC 16:0_22:4 + PC 18:0_20:4 + PC 18:1_20:3 + PC 18:2_20:2) / PC aa C38:4**
**Shapiro−Wilk Test of log quotients, pv: 0.00246; NO**
SW: ok; ANOVA: Comp. ~ Subject_ID −> pv = 0.52376; EQUAL
Wilcoxon: Challenge; fasting: 0.25; sport: 0.125; OLTT: 0.625

**PC 16:0_22:4 / PC aa C38:4**
**Shapiro−Wilk Test of log quotients, pv: 0.01057; OK**
SW: ok; ANOVA: Comp. ~ Subject_ID −> pv = 2e−05; DIFFERENCE
Wilcoxon: Challenge; fasting: 0.625; sport: 0.875; OLTT: 0.375

**PC 18:0_20:4 / PC aa C38:4**
**Shapiro−Wilk Test of log quotients, pv: 0.04492; OK**
SW: ok; ANOVA: Comp. ~ Subject_ID −> pv = 0.05199; EQUAL
Wilcoxon: Challenge; fasting: 0.375; sport: 0.125; OLTT: 0.625

**PC 18:1_20:3 / PC aa C38:4**
**Shapiro−Wilk Test of log quotients, pv: 0; NO**
SW: ok; ANOVA: Comp. ~ Subject_ID −> pv = 2e−05; DIFFERENCE
Wilcoxon: Challenge; fasting: 0.125; sport: 0.25; OLTT: 0.125

**PC 18:2_20:2 / PC aa C38:4**
**Shapiro−Wilk Test of log quotients, pv: 0; NO**
SW: no; Kruskal: Comp. ~ Subject_ID −> pv = 0; DIFFERENCE
Wilcoxon: Challenge; fasting: 0.125; sport: 0.625; OLTT: 0.875

#### Quant. comp., Subject 5
n = 55, 55, 55, 54; comb.: 54

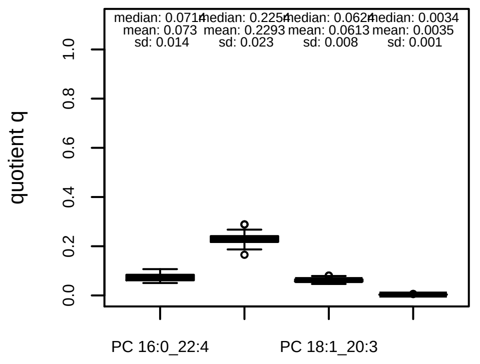

#### Quant. comp., Subject 6
n = 55, 55, 55, 55; comb.: 55

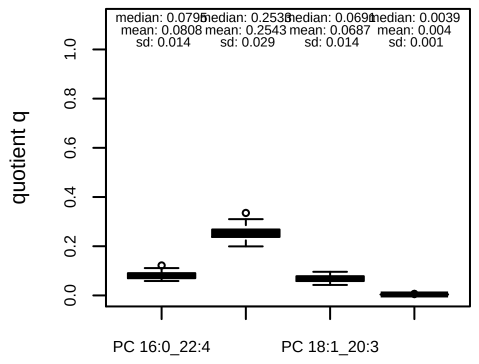

#### Quant. comp., Subject 7
n = 55, 55, 55, 55; comb.: 55

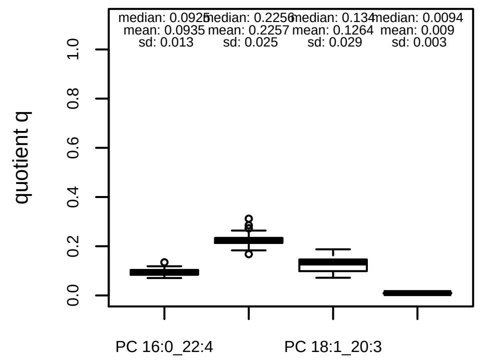

#### Quant. comp., Subject 8
n = 56, 56, 56, 56; comb.: 56

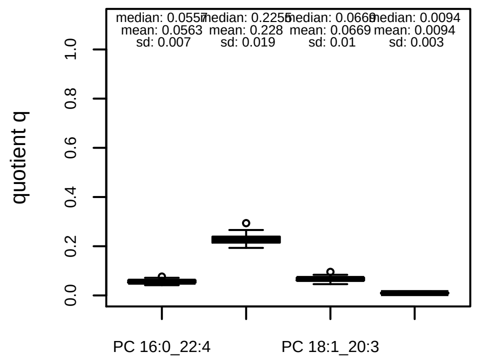

#### Trends of proportions q during challenges

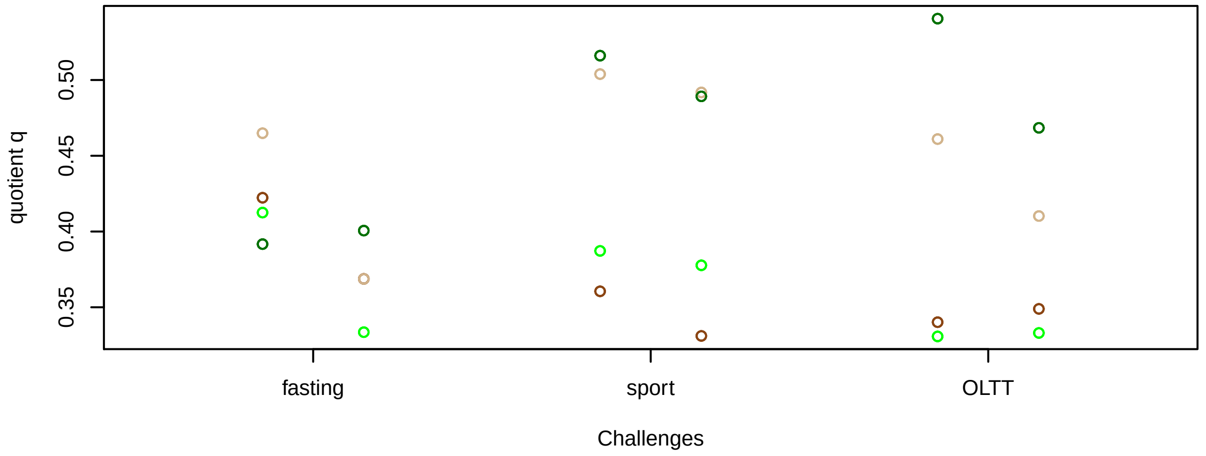


#### Proportions q per time point

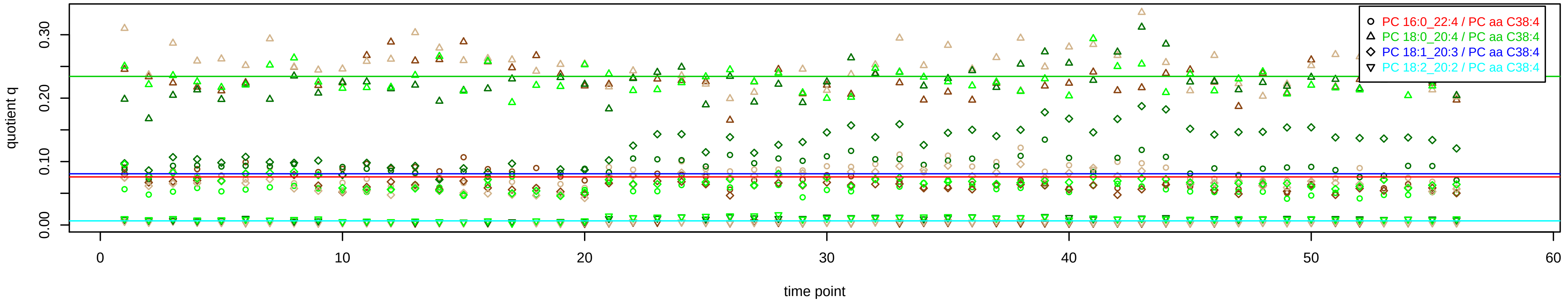

# PC aa C38:5 = PC 16:0_22:5 + PC 18:0_20:5 + PC 18:1_20:4 + PC 18:2_20:3 + R

PC 18:0_20:5 excluded because of missingness > 75%

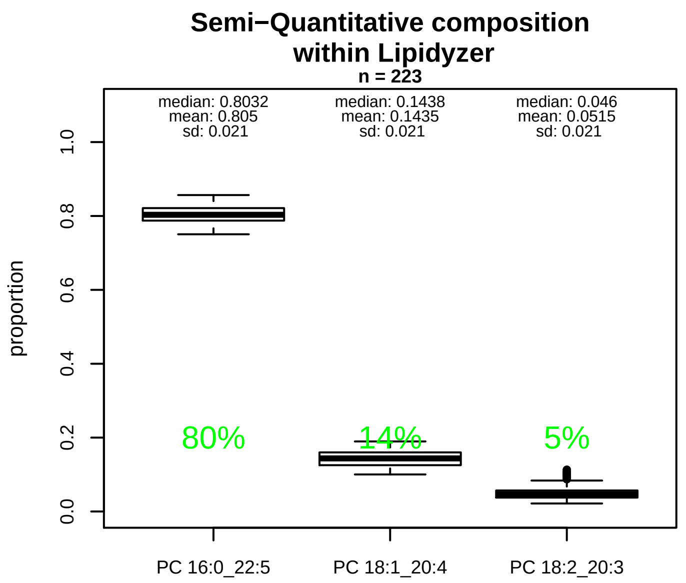


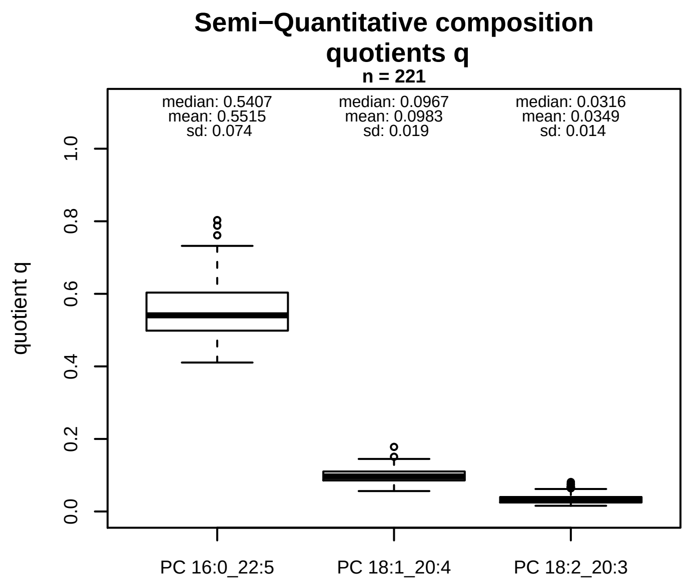


### Semi−Quantitative composition: Lipidyzer

PC aa C38:5 consists of:
PC 16:0_22:5 80.5%
PC 18:1_20:4 14.4%
PC 18:2_20:3 5.1%
and of (not quantified):
PC 18:0_20:5, PC 16:1_22:4, PC 18:3_20:2, PC 18:4_20:1,
and further compounds

### Composition: mean of proportions q

conc(PC aa C38:5) * 0.5515 = conc(PC 16:0_22:5) [var(q)=0.5515]
Percentiles: 5%−>0.438, 25%−>0.4985, 75%−>0.6034, 95%−>0.6725
conc(PC aa C38:5) * 0.0983 = conc(PC 18:1_20:4) [var(q)=0.0983]
Percentiles: 5%−>0.0687, 25%−>0.0856, 75%−>0.1103, 95%−>0.1286
conc(PC aa C38:5) * 0.0349 = conc(PC 18:2_20:3) [var(q)=0.0349]
Percentiles: 5%−>0.0201, 25%−>0.0248, 75%−>0.0399, 95%−>0.0647

### Linear model

```
PC aa C38:5 ~ b * ( PC 16:0_22:5
   + PC 18:1_20:4
   + PC 18:2_20:3 )
```

b = 1.21475

$R^2$ = 0.677

### Ranges

| Measure | AbsoluteIDQ | sum(Lipidyzer) | delta |
|---|---|---|---|
| Min | 34.9 | 22.92 | 11.98 |
| Max | 84.4 | 63.92 | 20.48 |
| Mean | 57.86 | 39.37 | 18.49 |
| Median | 58 | 40.77 | 17.23 |
| SD | 11.88 | 8.02 | 3.85 |

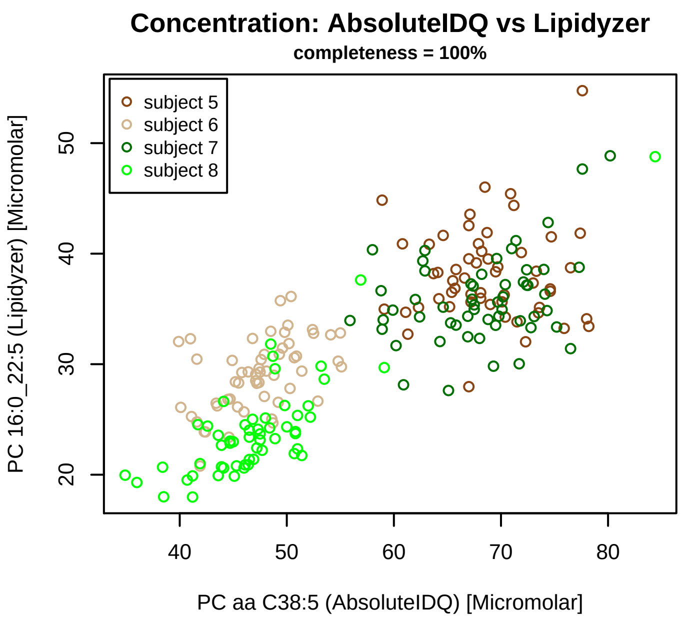


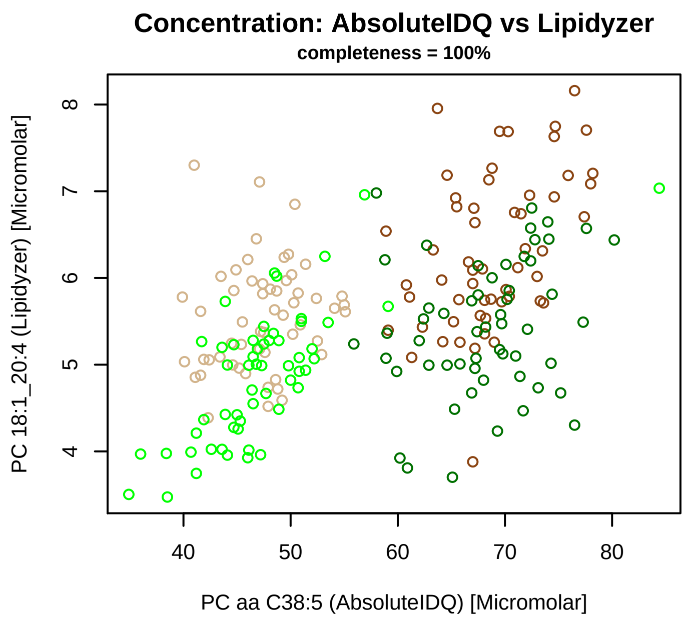


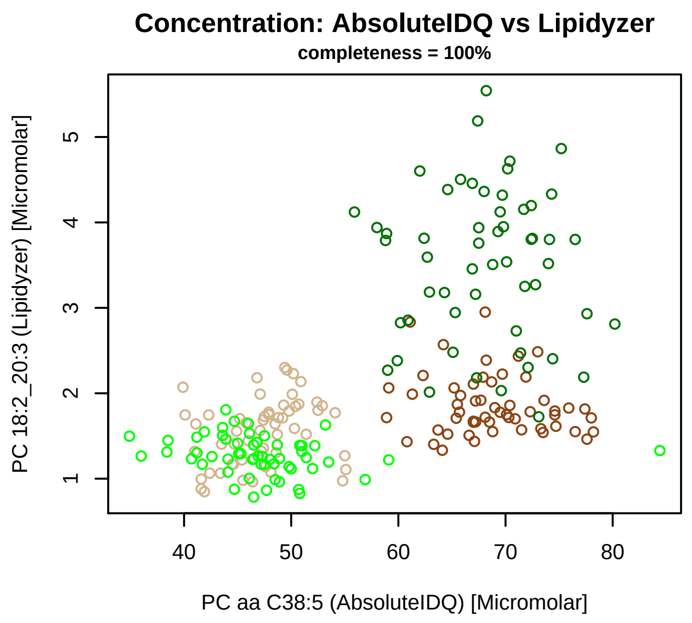


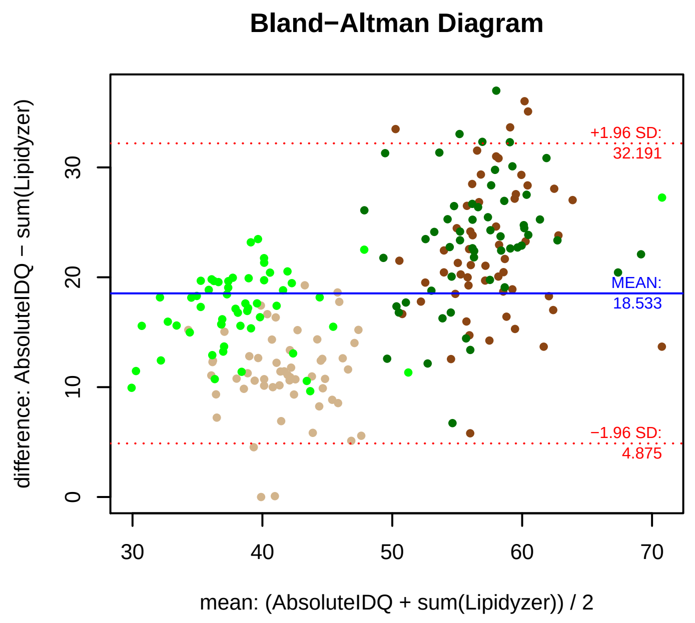


### Stability of composition

**(PC 16:0_22:5 + PC 18:1_20:4 + PC 18:2_20:3) / PC aa C38:5**
**Shapiro−Wilk Test of log quotients, pv: 0.37049; OK**
SW: ok; ANOVA: Comp. ~ Subject_ID −> pv = 2e−05; DIFFERENCE
Wilcoxon: Challenge; fasting: 0.625; sport: 0.25; OLTT: 0.25

**PC 16:0_22:5 / PC aa C38:5**
**Shapiro−Wilk Test of log quotients, pv: 0.26347; OK**
SW: ok; ANOVA: Comp. ~ Subject_ID −> pv = 0; DIFFERENCE
Wilcoxon: Challenge; fasting: 0.875; sport: 0.25; OLTT: 0.25

**PC 18:1_20:4 / PC aa C38:5**
**Shapiro−Wilk Test of log quotients, pv: 0.41624; OK**
SW: ok; ANOVA: Comp. ~ Subject_ID −> pv = 0.82498; EQUAL
Wilcoxon: Challenge; fasting: 0.375; sport: 0.25; OLTT: 0.375

**PC 18:2_20:3 / PC aa C38:5**
**Shapiro−Wilk Test of log quotients, pv: 0.00026; NO**
SW: ok; ANOVA: Comp. ~ Subject_ID −> pv = 0.03257; EQUAL
Wilcoxon: Challenge; fasting: 0.125; sport: 0.125; OLTT: 0.125

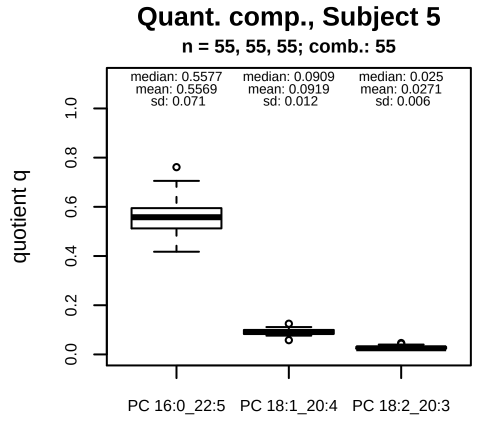


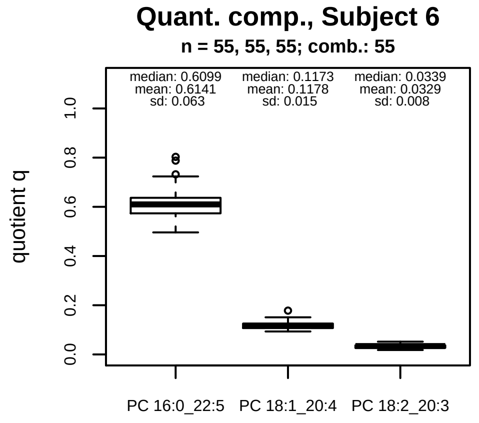


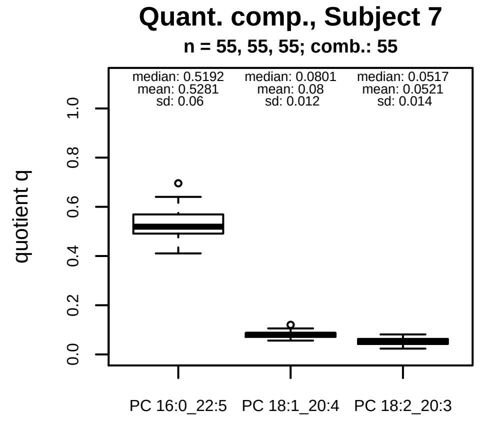


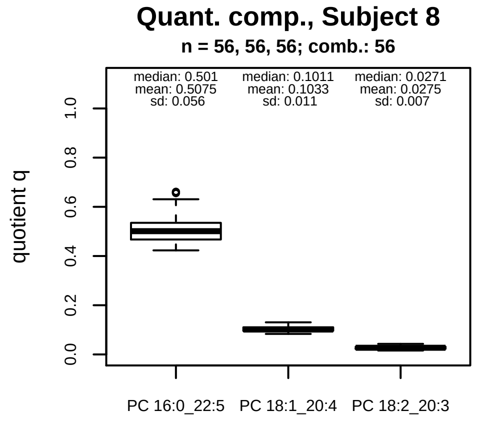


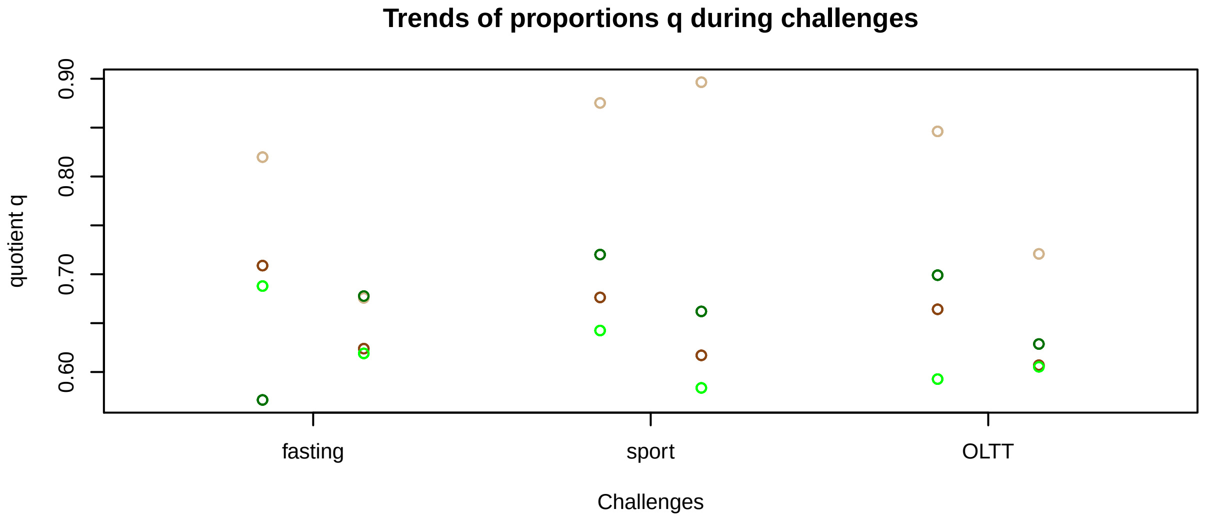


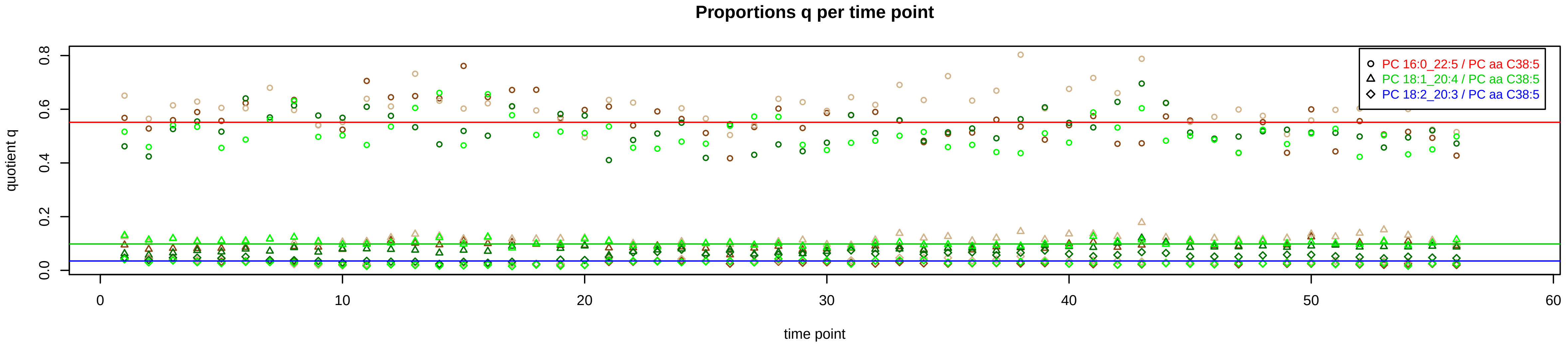

# PC aa C38:6 = PC 16:0_22:6 + PC 18:1_20:5 + PC 18:2_20:4 + R

PC 18:1_20:5 excluded because of missingness > 75%

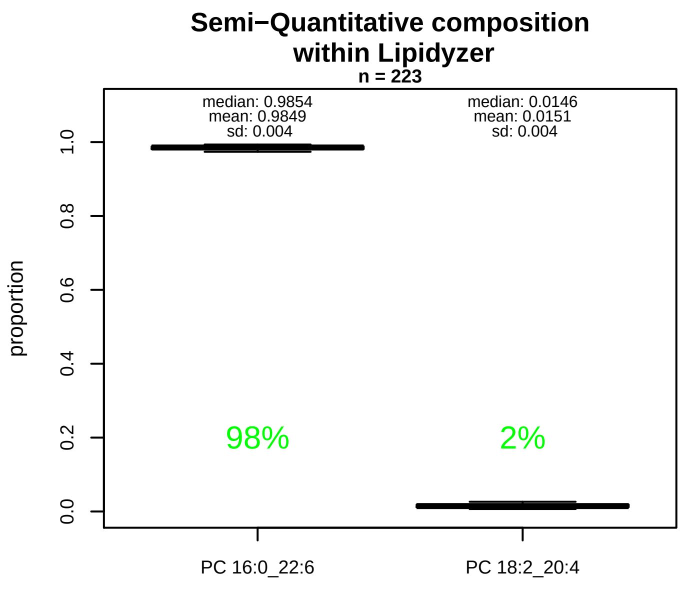


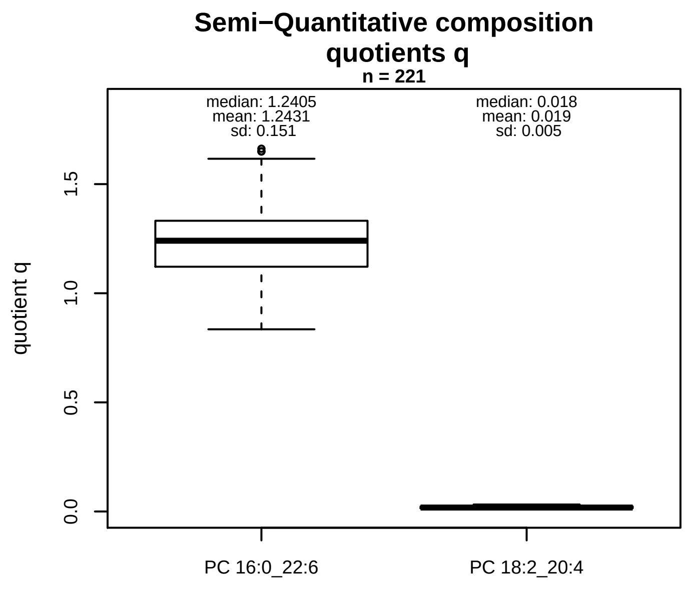


### Semi−Quantitative composition: Lipidyzer

PC aa C38:6 consists of:
PC 16:0_22:6 98.5%
PC 18:2_20:4 1.5%
and of (not quantified):
PC 18:1_20:5, PC 16:1_22:5, PC 18:3_20:3, PC 18:4_20:2,
and further compounds

### Linear model

```
PC aa C38:6 ~ b * ( PC 16:0_22:6
   + PC 18:2_20:4 )
```

b = 0.71118

$R^2$ = 0.77057

### Ranges

| Measure | AbsoluteIDQ | sum(Lipidyzer) | delta |
|---|---|---|---|
| Min | 50.4 | 67.49 | 17.09 |
| Max | 141.6 | 188.25 | 46.65 |
| Mean | 87.19 | 109.18 | 21.99 |
| Median | 82.9 | 100.57 | 17.67 |
| SD | 21.12 | 26.01 | 4.89 |

### Composition: mean of proportions q

conc(PC aa C38:6) * 1.2431 = conc(PC 16:0_22:6) [var(q)=1.2431]
Percentiles: 5%−>1.0168, 25%−>1.1214, 75%−>1.3321, 95%−>1.5009
conc(PC aa C38:6) * 0.019 = conc(PC 18:2_20:4) [var(q)=0.019]
Percentiles: 5%−>0.0118, 25%−>0.0148, 75%−>0.0234, 95%−>0.0279

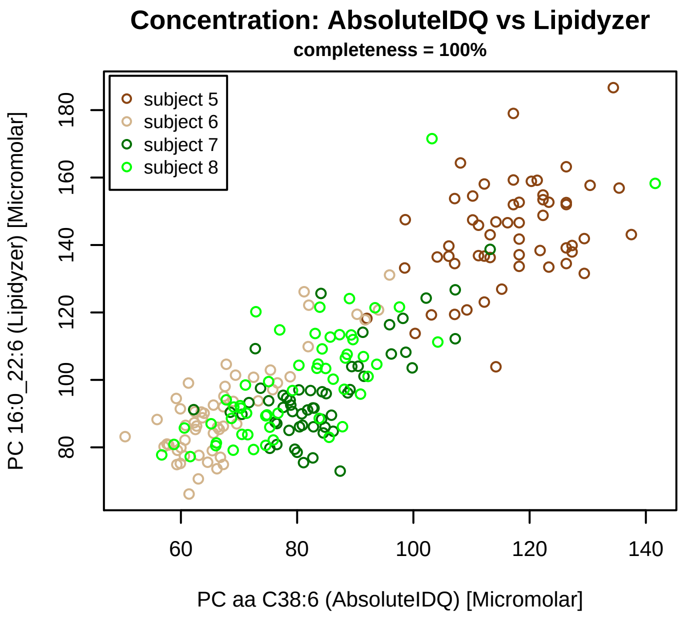


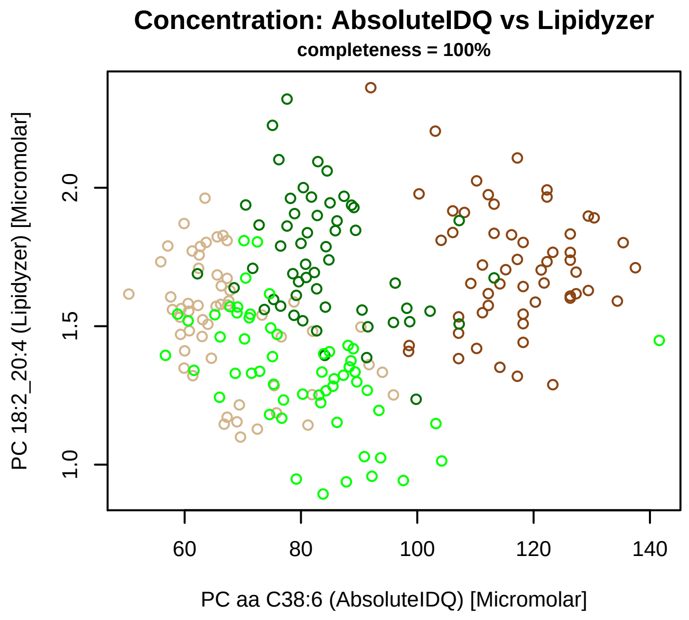


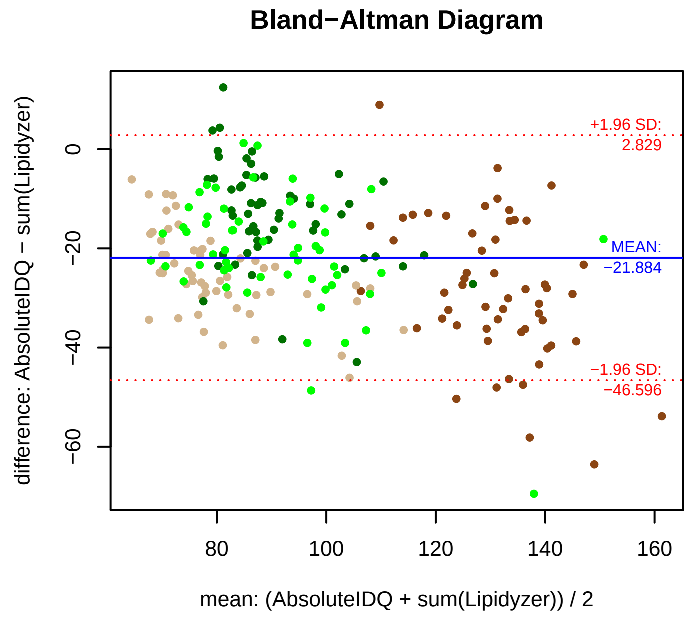


### Stability of composition

**(PC 16:0_22:6 + PC 18:2_20:4) / PC aa C38:6**
**Shapiro−Wilk Test of log quotients, pv: 0.86063; OK**
SW: ok; ANOVA: Comp. ~ Subject_ID −> pv = 0.06938; EQUAL
Wilcoxon: Challenge; fasting: 0.625; sport: 0.625; OLTT: 0.25

**PC 16:0_22:6 / PC aa C38:6**
**Shapiro−Wilk Test of log quotients, pv: 0.79949; OK**
SW: ok; ANOVA: Comp. ~ Subject_ID −> pv = 0.05839; EQUAL
Wilcoxon: Challenge; fasting: 0.625; sport: 0.625; OLTT: 0.25

**PC 18:2_20:4 / PC aa C38:6**
**Shapiro−Wilk Test of log quotients, pv: 0.00068; NO**
SW: no; Kruskal: Comp. ~ Subject_ID −> pv = 0; DIFFERENCE
Wilcoxon: Challenge; fasting: 0.125; sport: 0.125; OLTT: 0.625

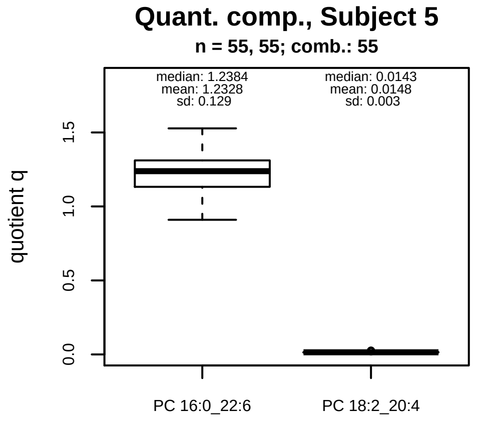


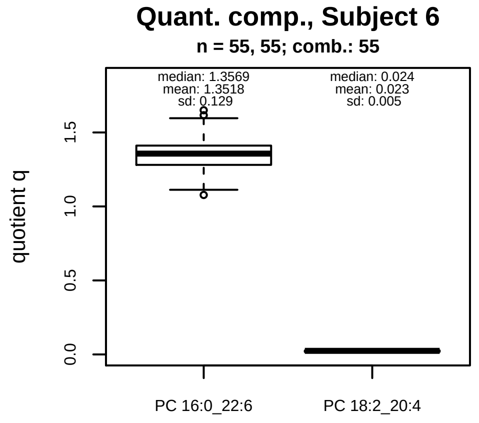


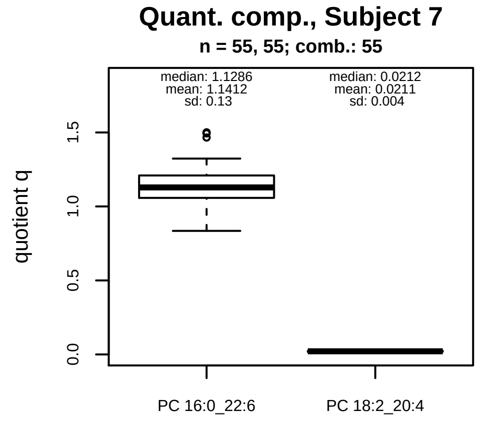


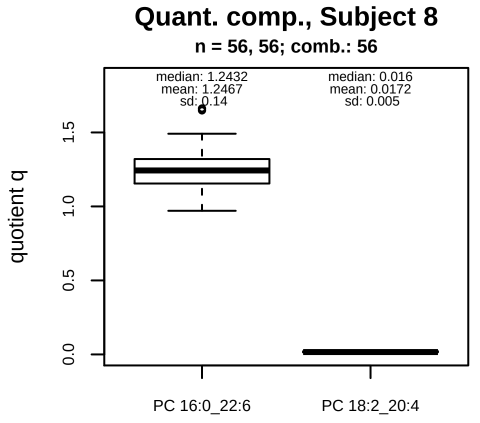


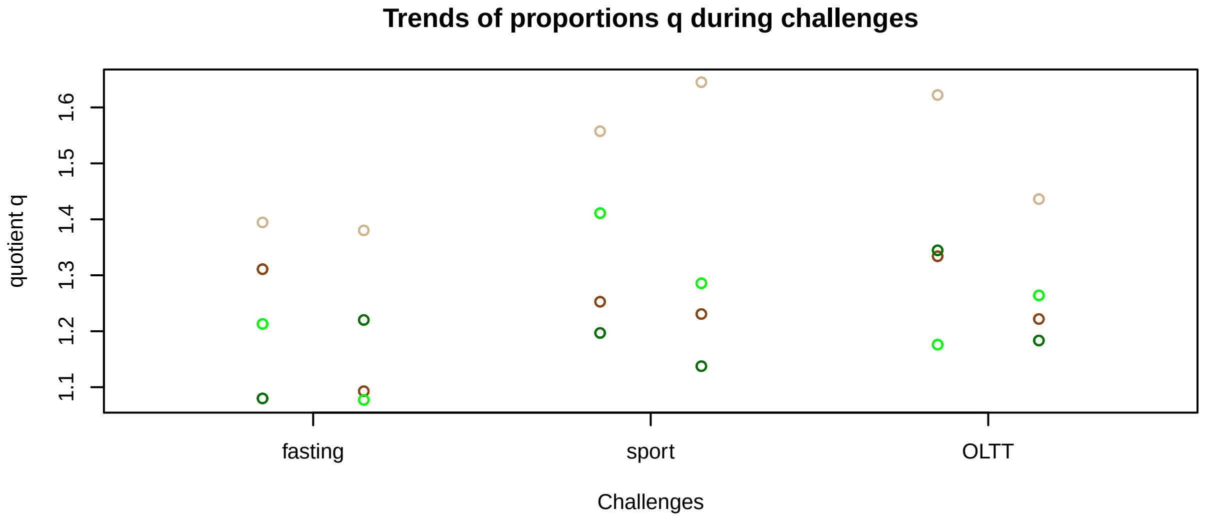


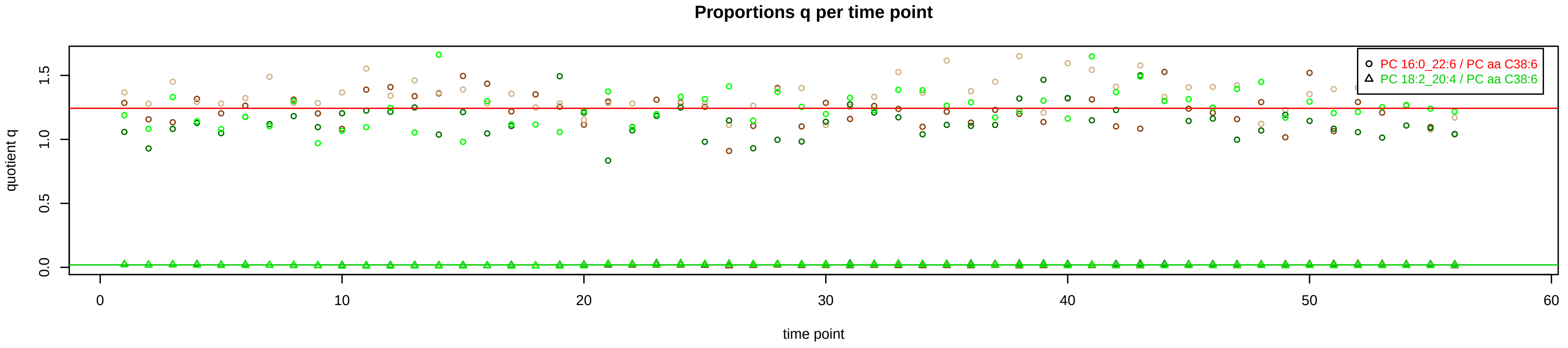

# PC aa C40:1 = R

No independent Variable measured.

### Qualitative composition

PC aa C40:1 consists of:
PC 16:0_24:1, PC 18:0_22:1, PC 18:1_22:0, PC 19:1_21:0,
PC O−20:1_21:0
and further compounds

# PC aa C40:2 = PC 18:0_22:2 + PC 18:1_22:1 + R

PC 18:0_22:2, PC 18:1_22:1 excluded because of missingness > 75%

**No independent Variable with coverage >0.25 out of PC 18:0_22:2, PC 18:1_22:1**

### Qualitative composition

PC aa C40:2 consists of:
PC 18:0_22:2, PC 18:1_22:1, PC 16:1_24:1, PC 18:2_22:0,
PC 20:1_20:1
and further compounds

# PC aa C40:3 = PC 18:2_22:1 + PC 20:0_20:3 + R

PC 18:2_22:1 excluded because of missingness > 75%

#### Semi−Quantitative composition within Lipidyzer

n = 216

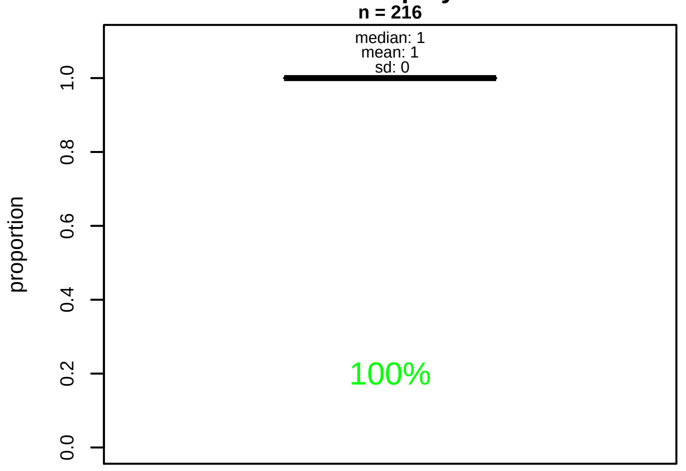


#### Semi−Quantitative composition quotients q

n = 214

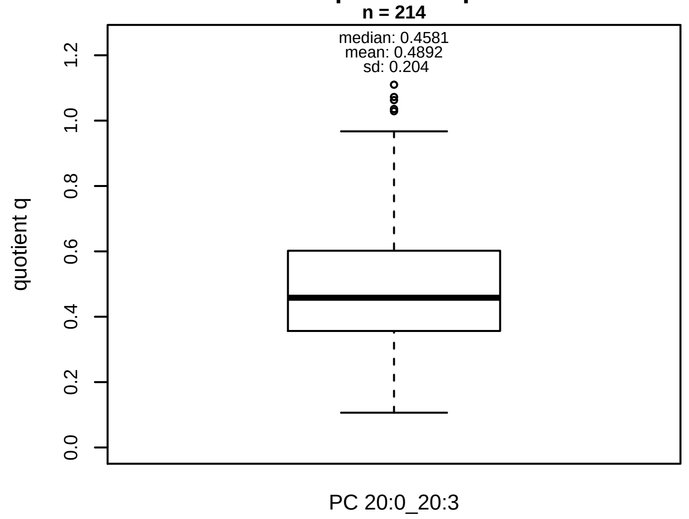


#### Semi−Quantitative composition: Lipidyzer

PC aa C40:3 consists of:
PC 20:0_20:3 100%
and of (not quantified):
PC 18:2_22:1, PC 18:0_22:3, PC 18:1_22:2, PC 18:3_22:0, PC 20:1_20:2
and further compounds

#### Linear model

```
PC aa C40:3 ~ b * ( PC 20:0_20:3 )
```

b = −0.06122

$R^2$ = 0.00209

#### Ranges

| Measure | AbsoluteIDQ | sum(Lipidyzer) | delta |
|---|---|---|---|
| Min | 0.42 | 0.11 | 0.31 |
| Max | 1.31 | 0.96 | 0.35 |
| Mean | 0.77 | 0.36 | 0.41 |
| Median | 0.73 | 0.32 | 0.41 |
| SD | 0.19 | 0.14 | 0.05 |

#### Composition: mean of proportions q

conc(PC aa C40:3) * 0.4892 = conc(PC 20:0_20:3) [var(q)=0.4892]
Percentiles: 5%−>0.198, 25%−>0.3568, 75%−>0.5993, 95%−>0.8453

#### Concentration: AbsoluteIDQ vs Lipidyzer

completeness = 96.86%

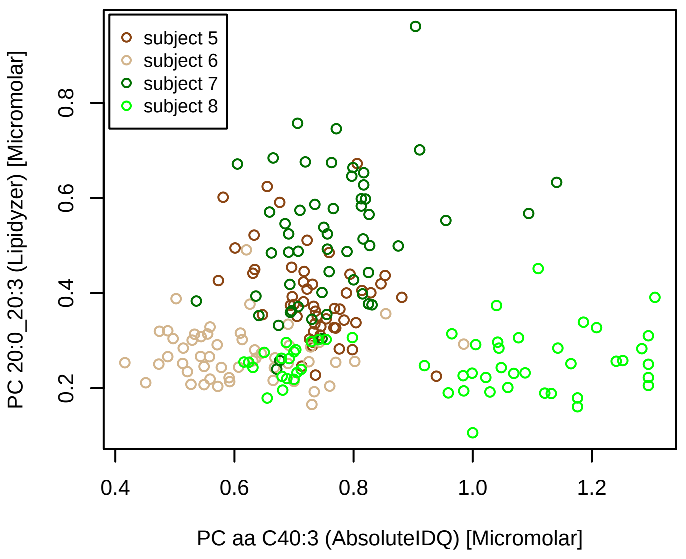


#### Bland−Altman Diagram

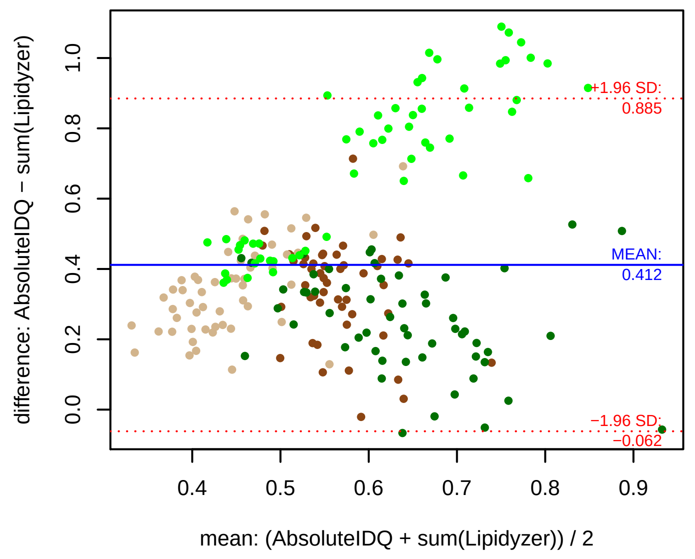


#### Stability of composition

**PC 20:0_20:3 / PC aa C40:3**
**Shapiro−Wilk Test of log quotients, pv: 0.0089; NO**
SW: ok; ANOVA: Comp. ~ Subject_ID −> pv = 0; DIFFERENCE
Wilcoxon: Challenge; fasting: 1; sport: 0.375; OLTT: 0.125

#### Quant. comp., Subject 5

n = 51; comb.: 51

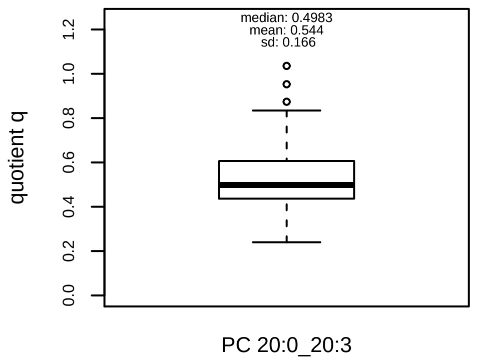


#### Quant. comp., Subject 6

n = 53; comb.: 53

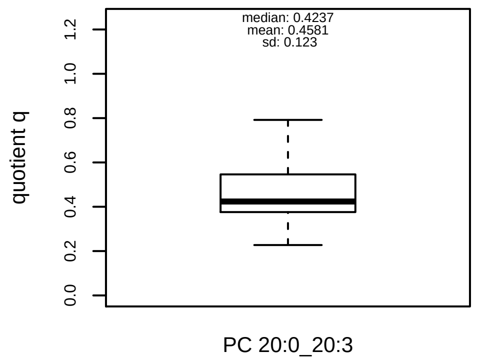


#### Quant. comp., Subject 7

n = 55; comb.: 55

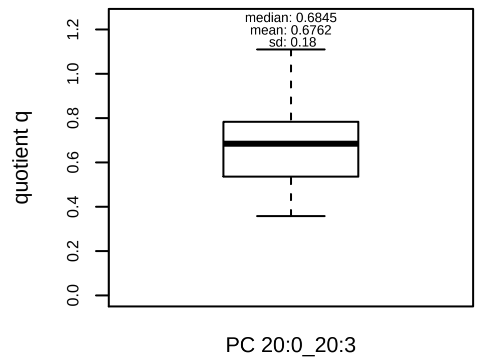


#### Quant. comp., Subject 8

n = 55; comb.: 55

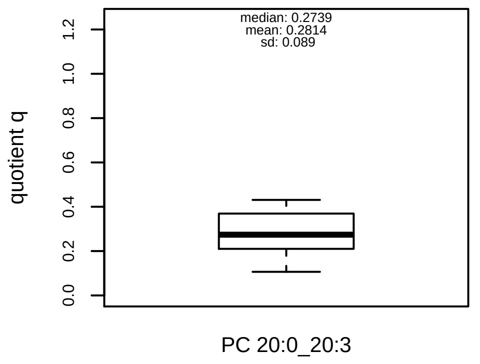


#### Trends of proportions q during challenges

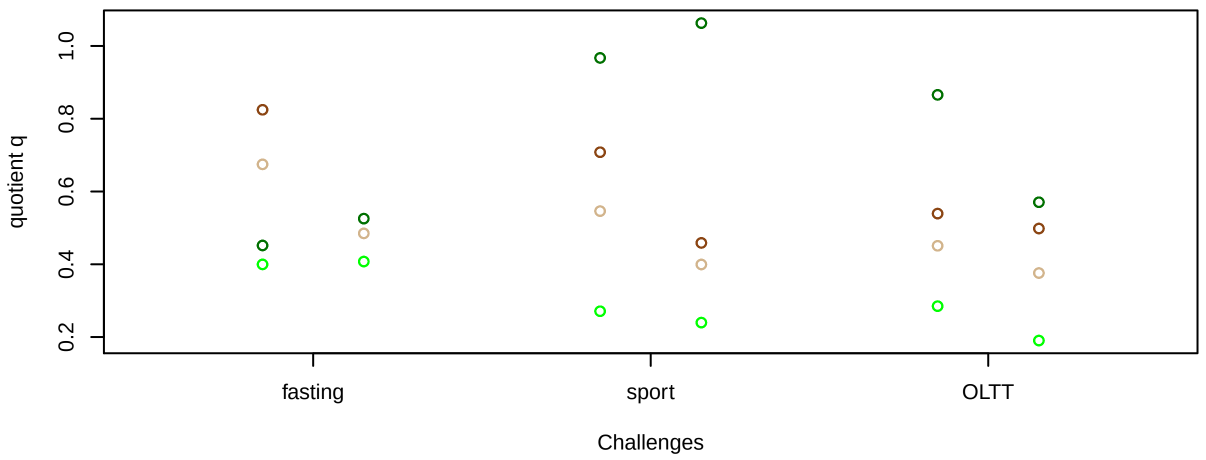


#### Proportions q per time point

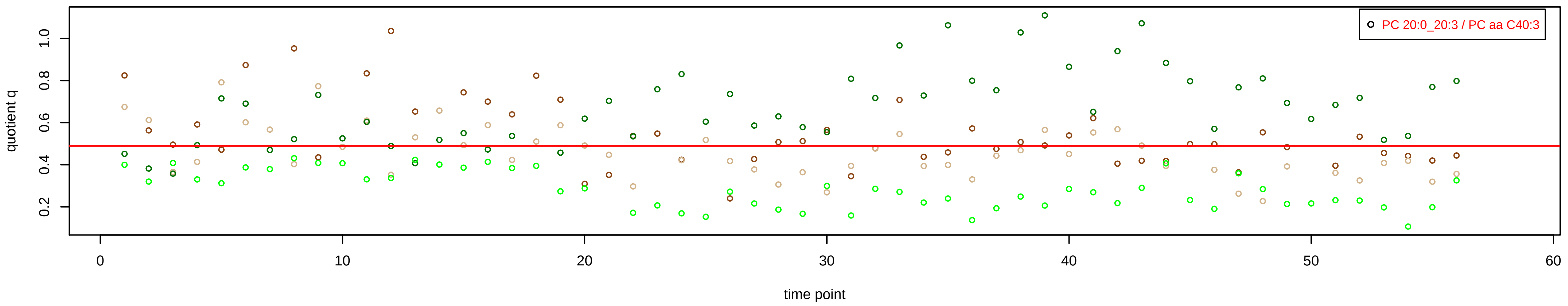

# PC aa C40:4 = PC 18:0_22:4 + PC 20:0_20:4 + R

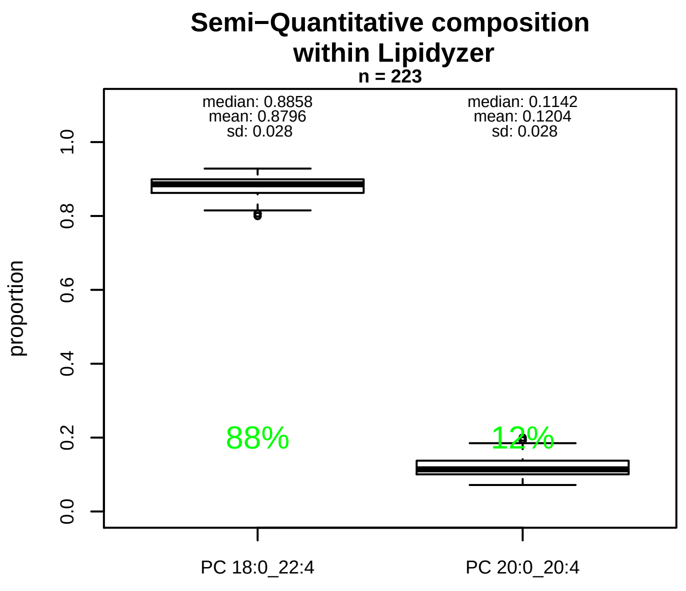


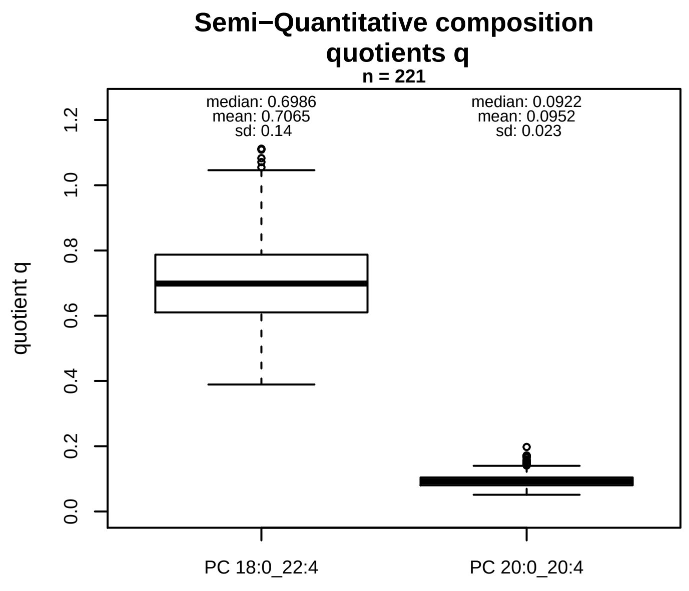


### Semi−Quantitative composition: Lipidyzer

PC aa C40:4 consists of:
PC 18:0_22:4 88%
PC 20:0_20:4 12%
and of (not quantified):
PC 18:2_22:2, PC 18:3_22:1, PC 18:4_22:0, PC 20:1_20:3, PC 20:2_20:2
and further compounds

### Linear model

```
PC aa C40:4 ~ b * ( PC 18:0_22:4
   + PC 20:0_20:4 )
```

b = 0.82589

$R^2$ = 0.67225

### Ranges

| Measure | AbsoluteIDQ | sum(Lipidyzer) | delta |
|---|---|---|---|
| Min | 1.84 | 1.08 | 0.76 |
| Max | 5.56 | 6.11 | 0.55 |
| Mean | 3.46 | 2.79 | 0.67 |
| Median | 3.25 | 2.85 | 0.4 |
| SD | 0.89 | 0.88 | 0 |

### Composition: mean of proportions q

conc(PC aa C40:4) * 0.7065 = conc(PC 18:0_22:4) [var(q)=0.7065]
Percentiles: 5%−>0.4936, 25%−>0.6103, 75%−>0.7872, 95%−>0.9503
conc(PC aa C40:4) * 0.0952 = conc(PC 20:0_20:4) [var(q)=0.0952]
Percentiles: 5%−>0.0644, 25%−>0.0803, 75%−>0.1044, 95%−>0.1461

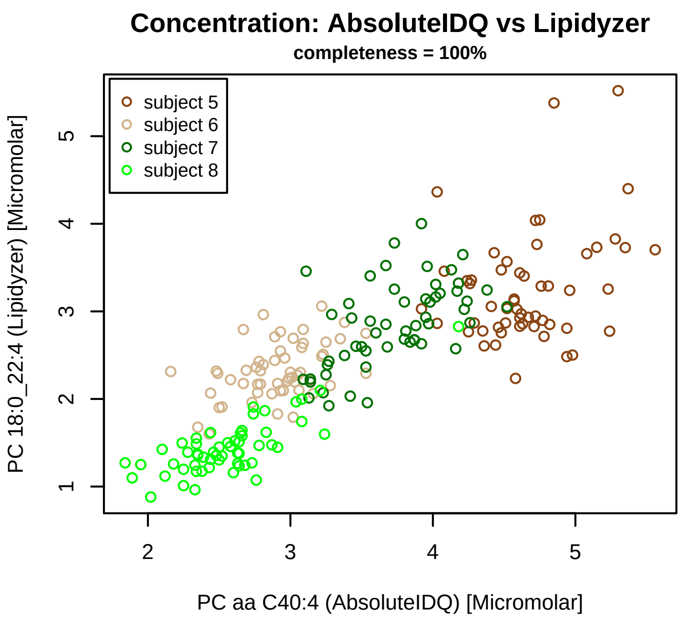


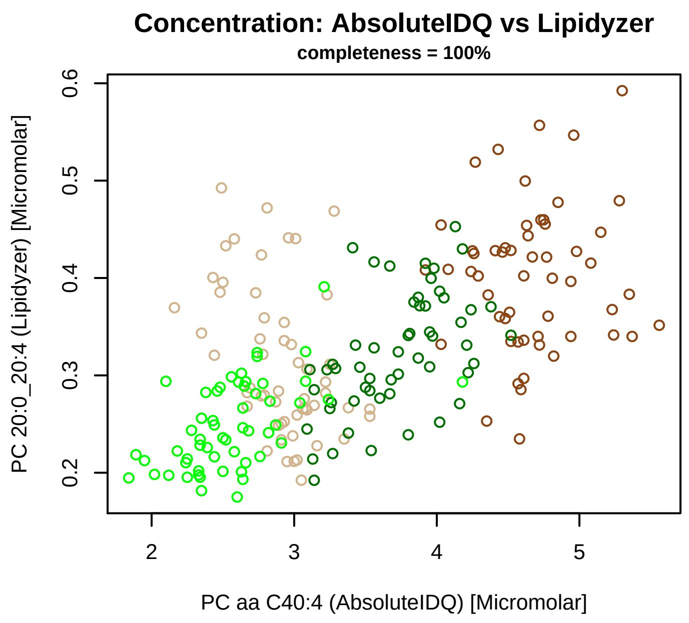


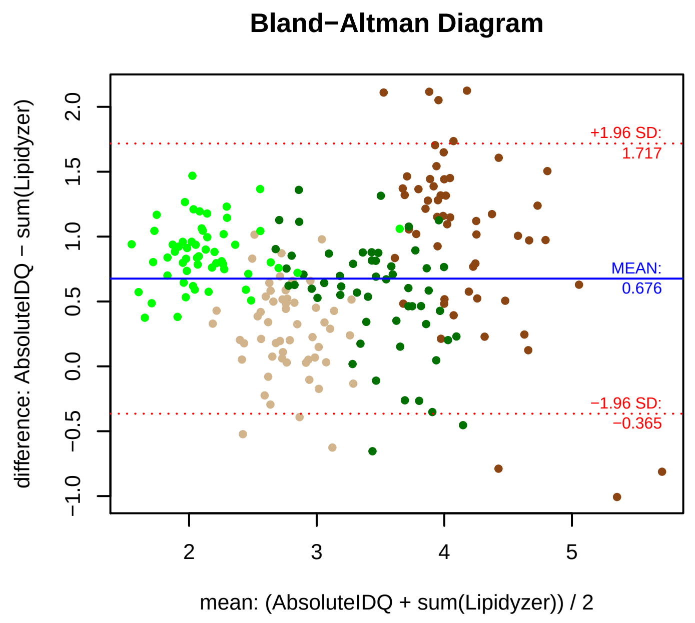


### Stability of composition

**(PC 18:0_22:4 + PC 20:0_20:4) / PC aa C40:4**
**Shapiro−Wilk Test of log quotients, pv: 0.92811; OK**
SW: ok; ANOVA: Comp. ~ Subject_ID −> pv = 0; DIFFERENCE
Wilcoxon: Challenge; fasting: 0.25; sport: 0.875; OLTT: 0.125

**PC 18:0_22:4 / PC aa C40:4**
**Shapiro−Wilk Test of log quotients, pv: 0.64098; OK**
SW: ok; ANOVA: Comp. ~ Subject_ID −> pv = 0; DIFFERENCE
Wilcoxon: Challenge; fasting: 0.25; sport: 0.875; OLTT: 0.125

**PC 20:0_20:4 / PC aa C40:4**
**Shapiro−Wilk Test of log quotients, pv: 0.00125; NO**
SW: ok; ANOVA: Comp. ~ Subject_ID −> pv = 0.19594; EQUAL
Wilcoxon: Challenge; fasting: 0.375; sport: 1; OLTT: 0.25

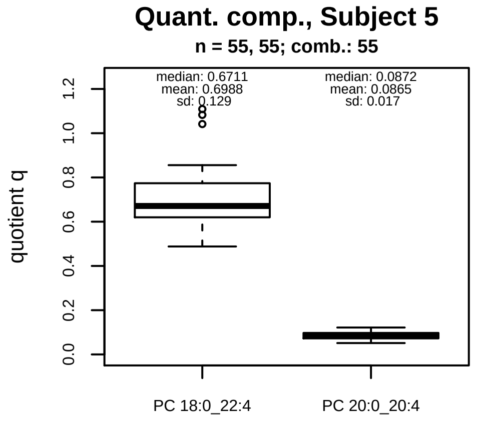


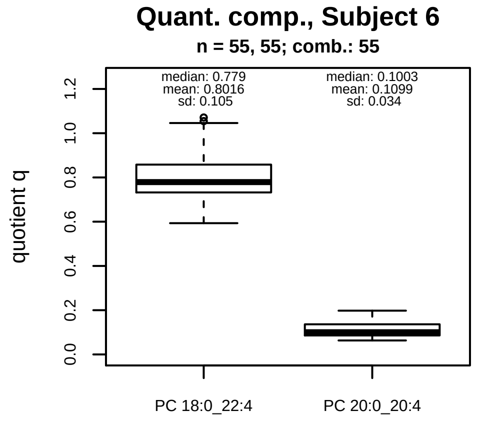


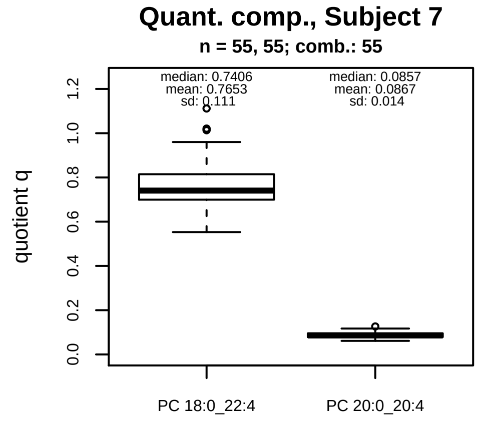


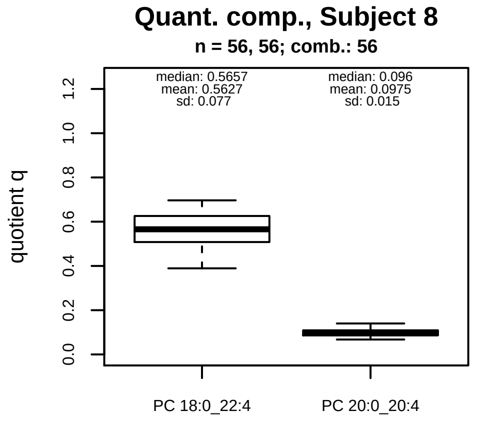


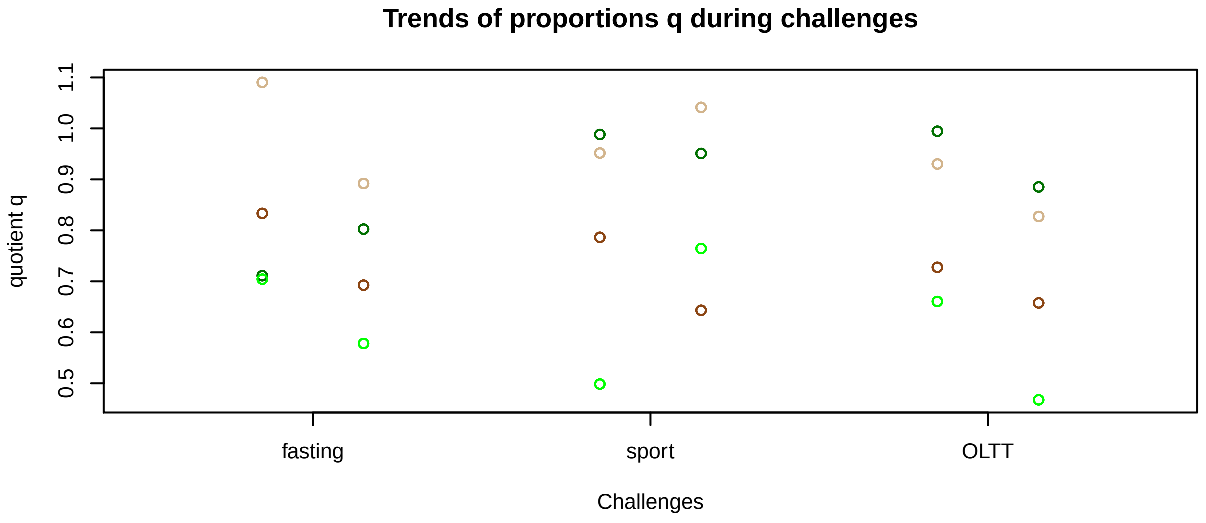


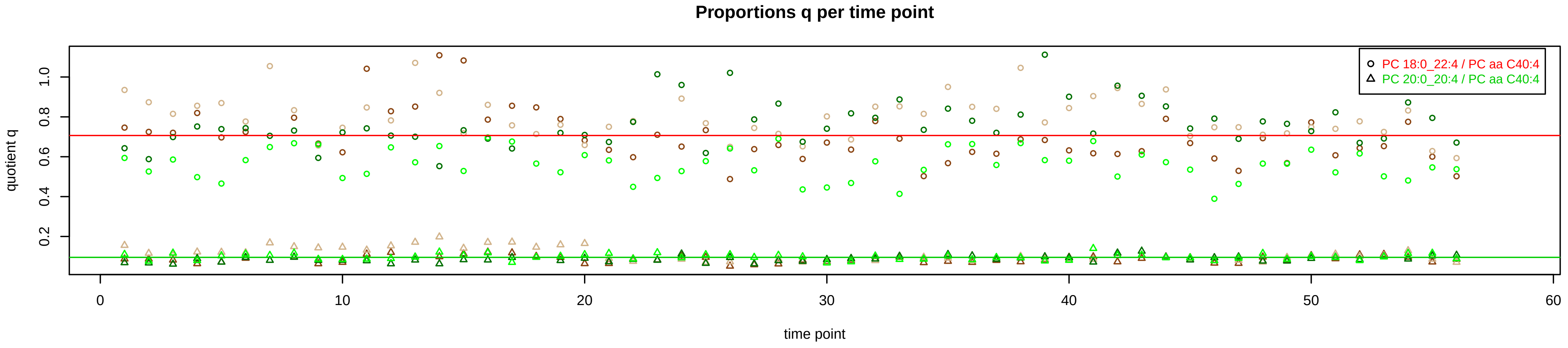

# PC aa C40:5 = PC 18:0_22:5 + PC 18:1_22:4 + R

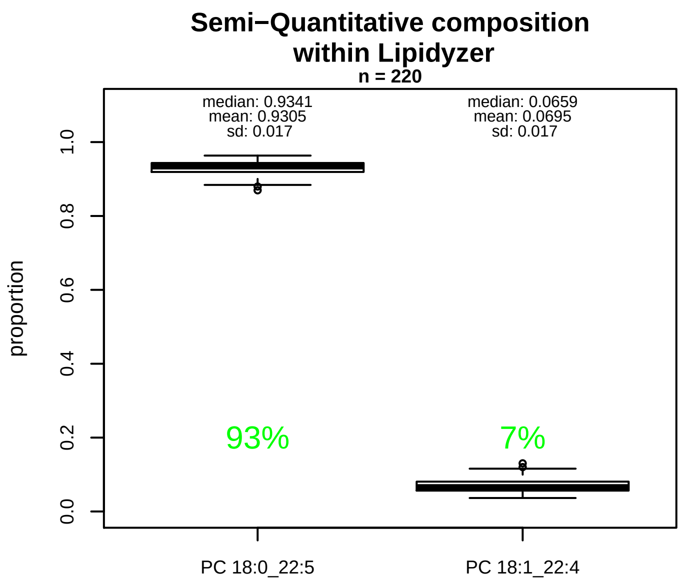


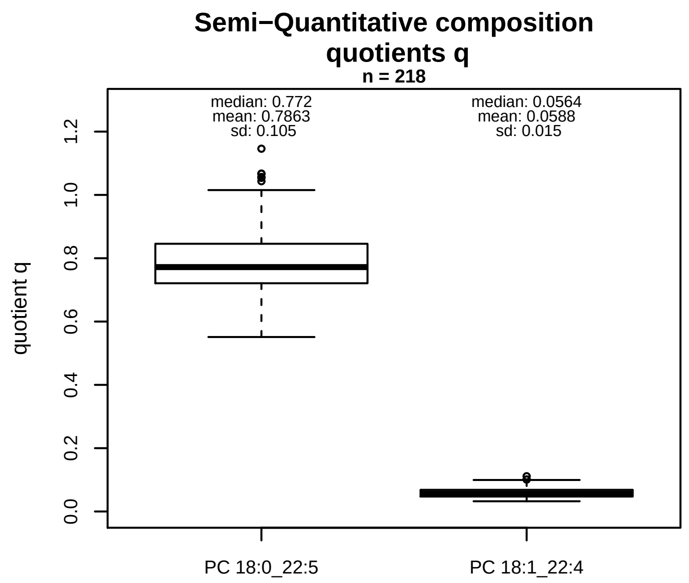


### Semi−Quantitative composition: Lipidyzer

PC aa C40:5 consists of:
PC 18:0_22:5 93%
PC 18:1_22:4 7%
and of (not quantified):
PC 18:3_22:2, PC 18:4_22:1, PC 20:0_20:5, PC 20:1_20:4,
PC 20:2_20:3
and further compounds

### Composition: mean of proportions q

conc(PC aa C40:5) * 0.7863 = conc(PC 18:0_22:5) [var(q)=0.7863]
Percentiles: 5%−>0.6343, 25%−>0.721, 75%−>0.8457, 95%−>0.9819
conc(PC aa C40:5) * 0.0588 = conc(PC 18:1_22:4) [var(q)=0.0588]
Percentiles: 5%−>0.0373, 25%−>0.0473, 75%−>0.0684, 95%−>0.084

### Linear model

```
PC aa C40:5 ~ b * ( PC 18:0_22:5
   + PC 18:1_22:4 )
```

b = 1.00995

$R^2$ = 0.76522

### Ranges

| Measure | AbsoluteIDQ | sum(Lipidyzer) | delta |
|---|---|---|---|
| Min | 5.45 | 4.55 | 0.9 |
| Max | 16.1 | 16.02 | 0.08 |
| Mean | 10.39 | 8.74 | 1.65 |
| Median | 9.76 | 8.55 | 1.21 |
| SD | 2.72 | 2.34 | 0.37 |

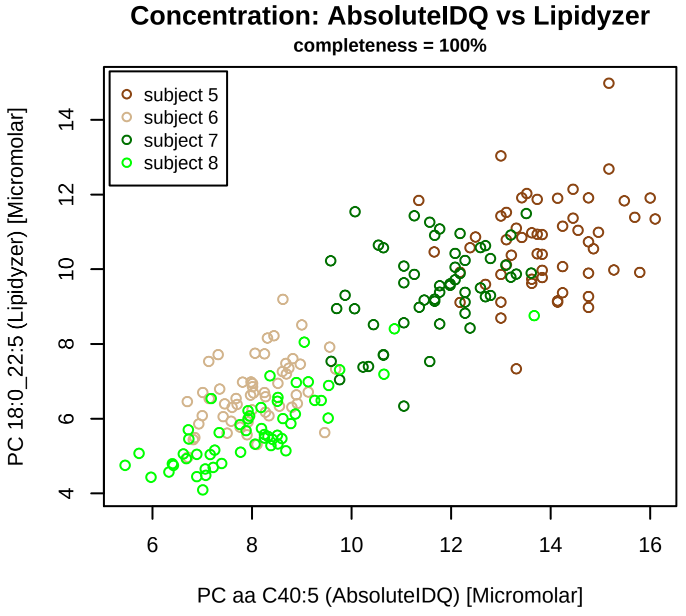


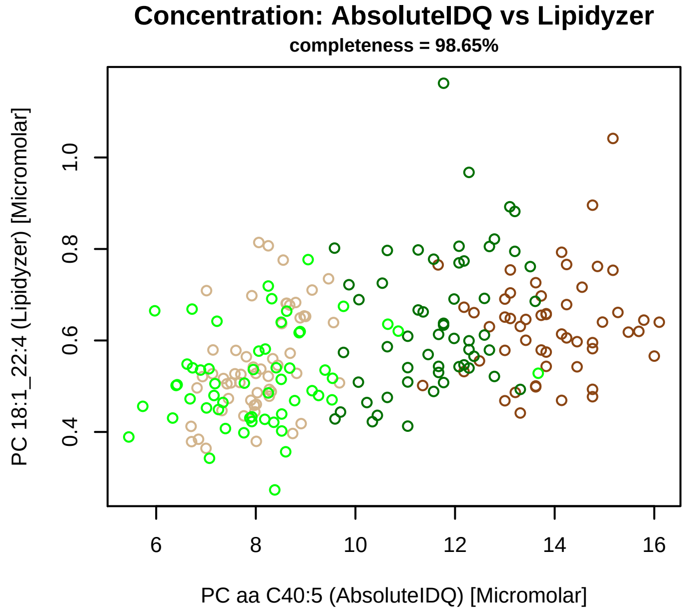


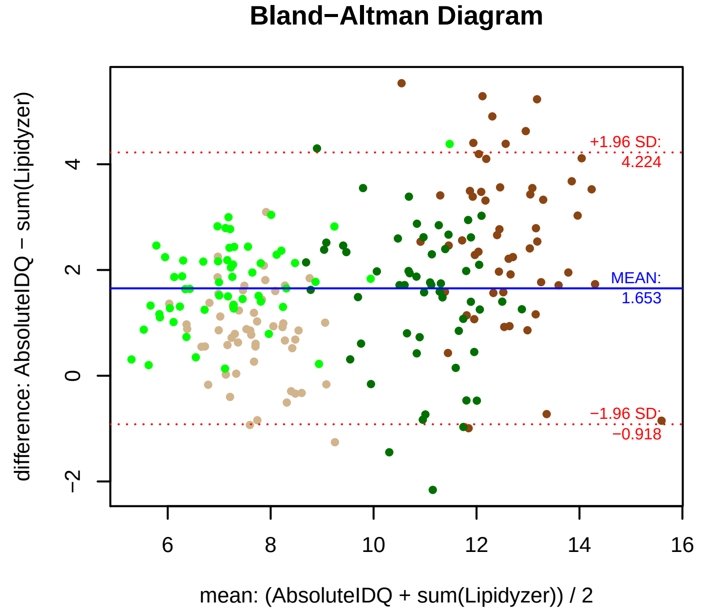


### Stability of composition

**(PC 18:0_22:5 + PC 18:1_22:4) / PC aa C40:5**
**Shapiro−Wilk Test of log quotients, pv: 0.55079; OK**
SW: ok; ANOVA: Comp. ~ Subject_ID −> pv = 0.06767; EQUAL
Wilcoxon: Challenge; fasting: 0.25; sport: 0.125; OLTT: 0.25

**PC 18:0_22:5 / PC aa C40:5**
**Shapiro−Wilk Test of log quotients, pv: 0.63789; OK**
SW: ok; ANOVA: Comp. ~ Subject_ID −> pv = 0.00534; DIFFERENCE
Wilcoxon: Challenge; fasting: 0.25; sport: 0.125; OLTT: 0.25

**PC 18:1_22:4 / PC aa C40:5**
**Shapiro−Wilk Test of log quotients, pv: 0.30263; OK**
SW: ok; ANOVA: Comp. ~ Subject_ID −> pv = 0; DIFFERENCE
Wilcoxon: Challenge; fasting: 0.125; sport: 0.625; OLTT: 1

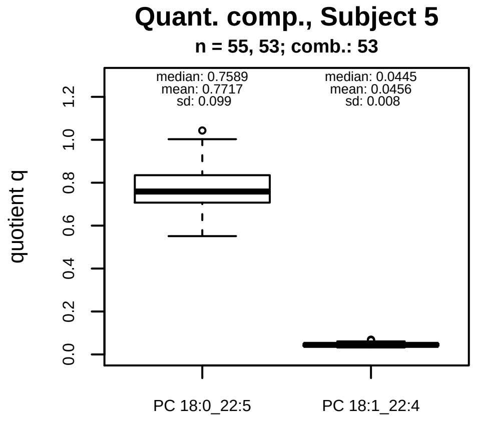


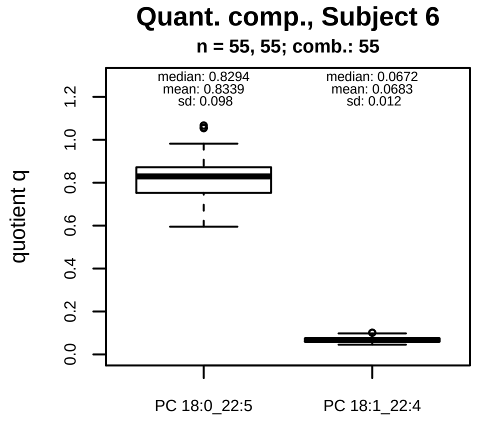


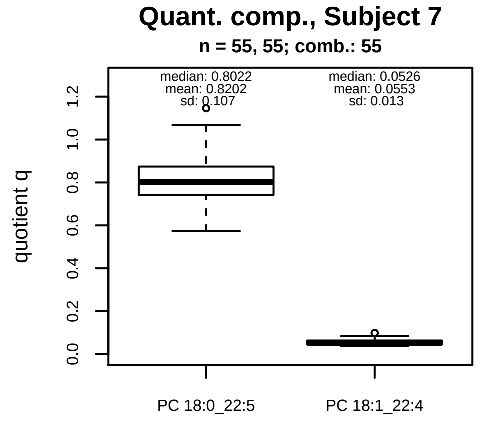


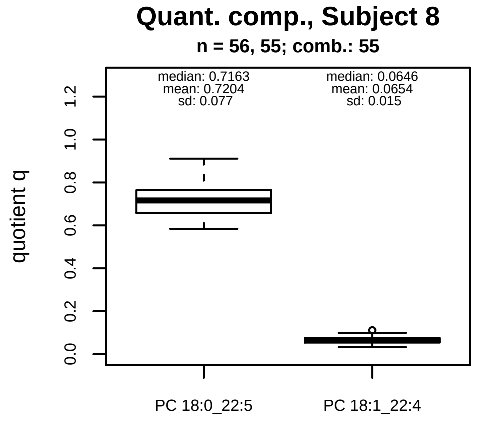


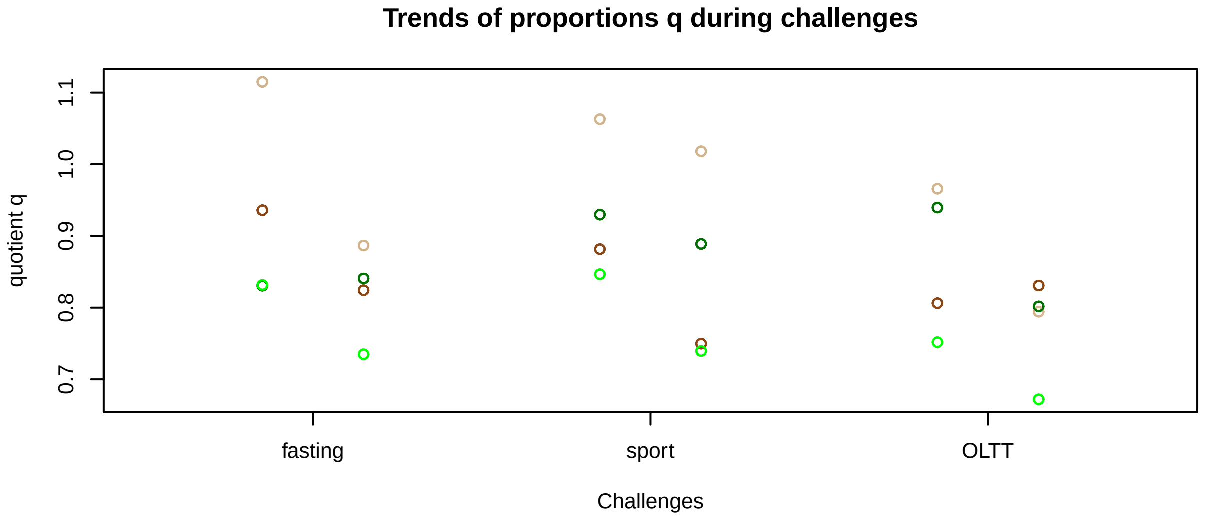


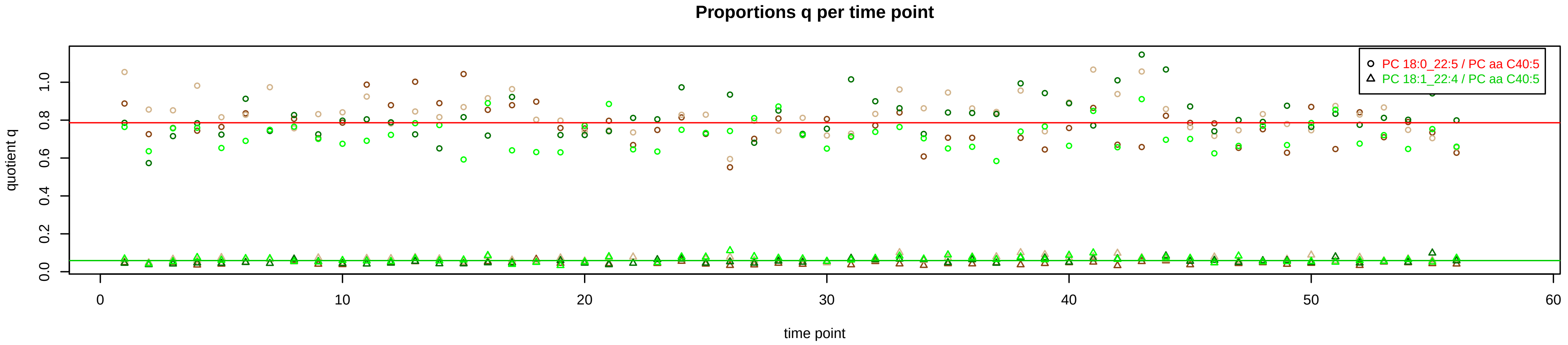

# PC aa C40:6 = PC 18:0_22:6 + PC 18:1_22:5 + PC 18:2_22:4 + R

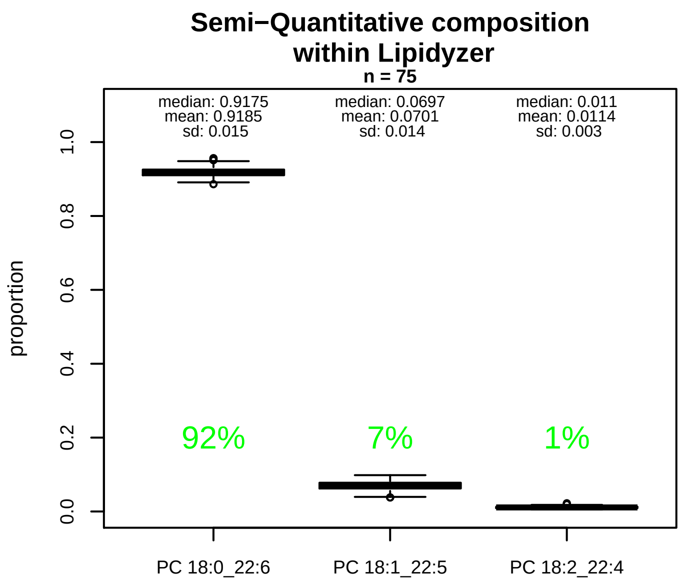


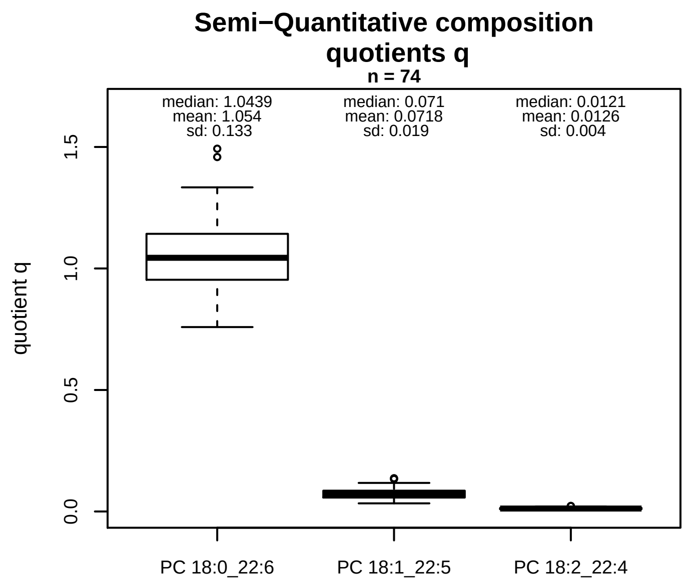


### Semi−Quantitative composition: Lipidyzer

PC aa C40:6 consists of:
PC 18:0_22:6 91.8%
PC 18:1_22:5 7%
PC 18:2_22:4 1.1%
and of (not quantified):
PC 18:4_22:2, PC 20:1_20:5, PC 20:2_20:4, PC 20:3_20:3,
and further compounds

### Linear model

```
PC aa C40:6 ~ b * ( PC 18:0_22:6
  + PC 18:1_22:5
  + PC 18:2_22:4 )
```

b = 0.91019

$R^2$ = 0.82665

### Ranges

| Measure | AbsoluteIDQ | sum(Lipidyzer) | delta |
|---|---|---|---|
| Min | 14.3 | 16.78 | 2.48 |
| Max | 43.5 | 43.62 | 0.12 |
| Mean | 25.21 | 24.91 | 0.3 |
| Median | 23.5 | 24.17 | 0.67 |
| SD | 7.46 | 5.35 | 2.11 |

### Composition: mean of proportions q

conc(PC aa C40:6) * 1.054 = conc(PC 18:0_22:6) [var(q)=1.054]
Percentiles: 5%−>0.8588, 25%−>0.9535, 75%−>1.1426, 95%−>1.2734
conc(PC aa C40:6) * 0.0718 = conc(PC 18:1_22:5) [var(q)=0.0718]
Percentiles: 5%−>0.0429, 25%−>0.0573, 75%−>0.0856, 95%−>0.1039
conc(PC aa C40:6) * 0.0126 = conc(PC 18:2_22:4) [var(q)=0.0126]
Percentiles: 5%−>0.0076, 25%−>0.0099, 75%−>0.0147, 95%−>0.0186

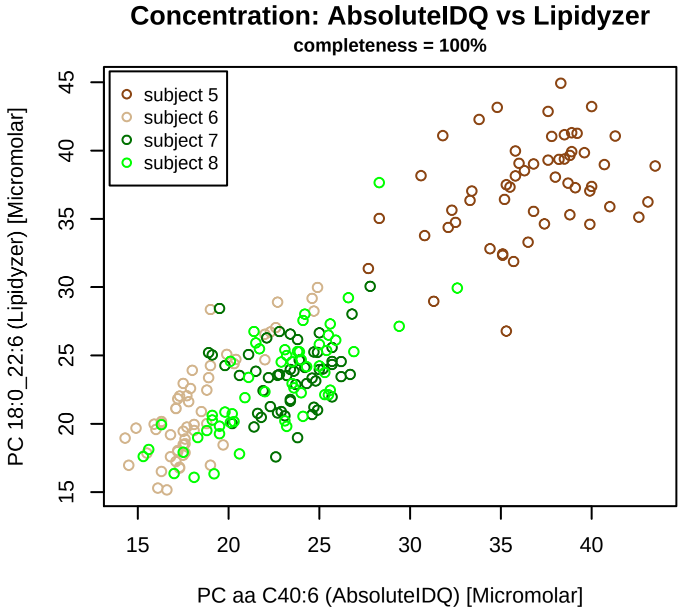


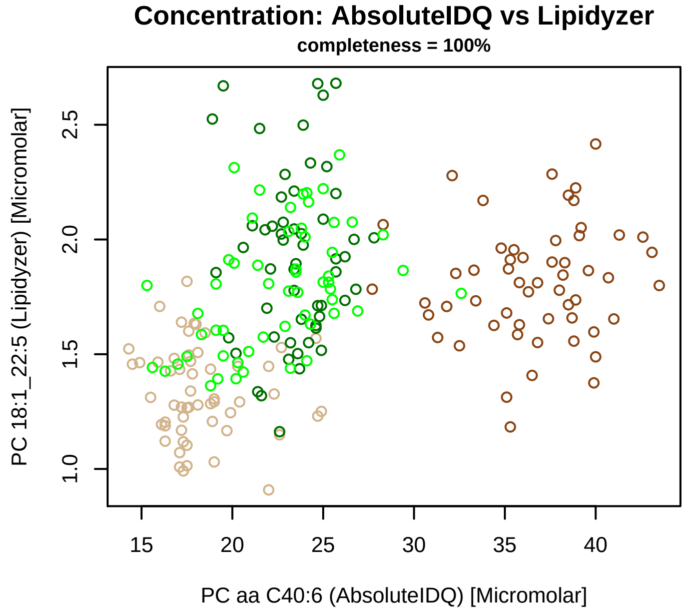


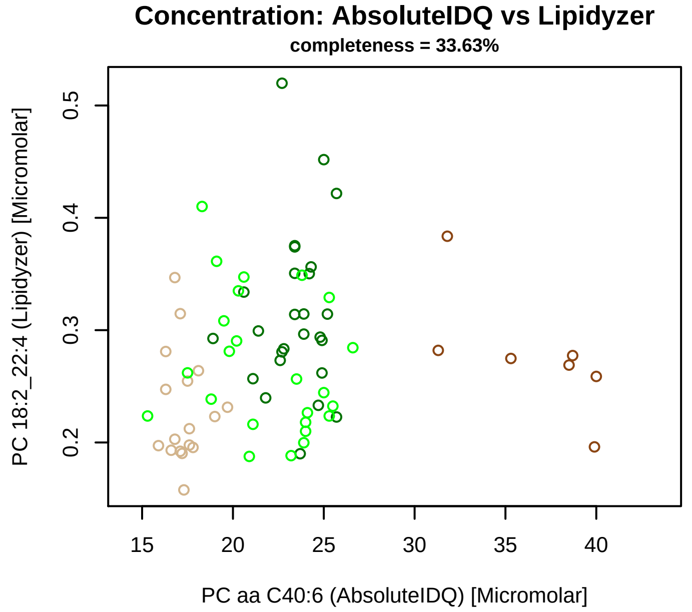


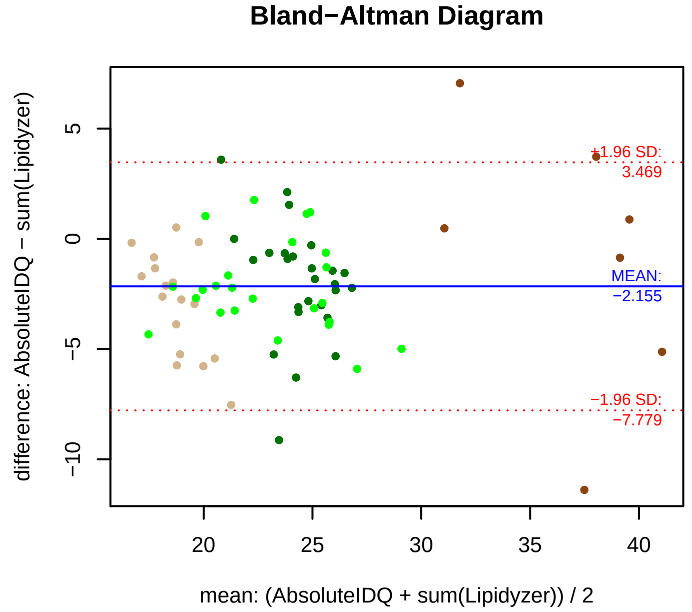


### Stability of composition

**(PC 18:0_22:6 + PC 18:1_22:5 + PC 18:2_22:4) / PC aa C40:6**
**Shapiro−Wilk Test of log quotients, pv: 0.76064; OK**
SW: ok; ANOVA: Comp. ~ Subject_ID −> pv = 0.82272; EQUAL
Wilcoxon: Challenge; sport: 1

**PC 18:0_22:6 / PC aa C40:6**
**Shapiro−Wilk Test of log quotients, pv: 0.25525; OK**
SW: ok; ANOVA: Comp. ~ Subject_ID −> pv = 0.08; EQUAL
Wilcoxon: Challenge; fasting: 0.125; sport: 0.875; OLTT: 0.375

**PC 18:1_22:5 / PC aa C40:6**
**Shapiro−Wilk Test of log quotients, pv: 0.04469; OK**
SW: ok; ANOVA: Comp. ~ Subject_ID −> pv = 0; DIFFERENCE
Wilcoxon: Challenge; fasting: 0.625; sport: 0.875; OLTT: 0.875

**PC 18:2_22:4 / PC aa C40:6**
**Shapiro−Wilk Test of log quotients, pv: 0.65312; OK**
SW: ok; ANOVA: Comp. ~ Subject_ID −> pv = 0.02697; EQUAL
Wilcoxon: Challenge; sport: 1

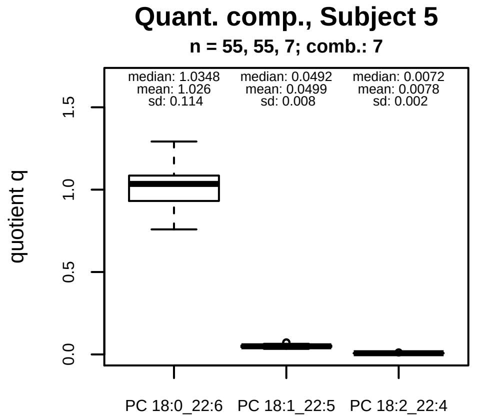


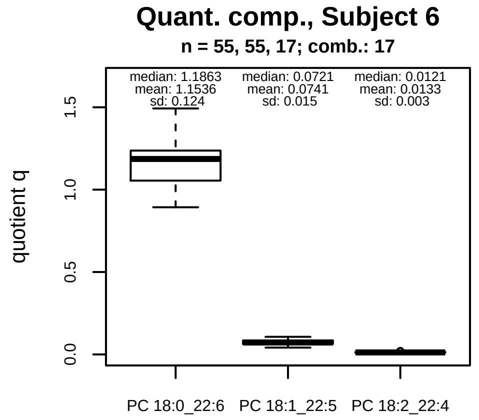


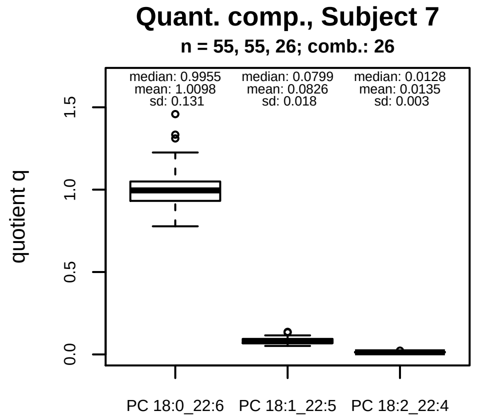


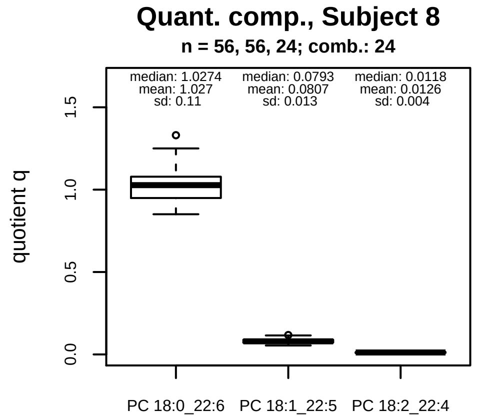


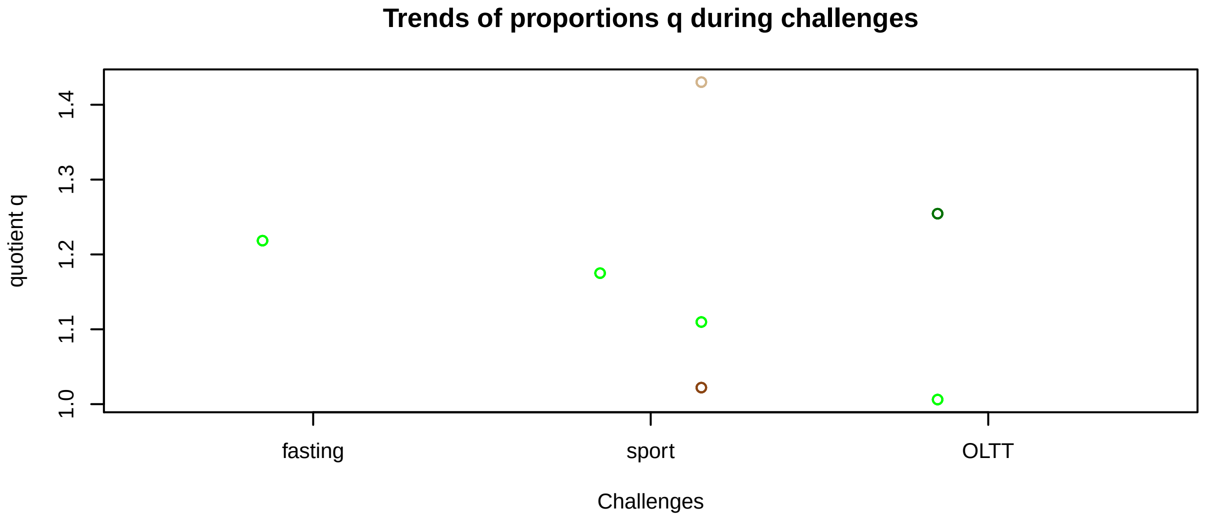


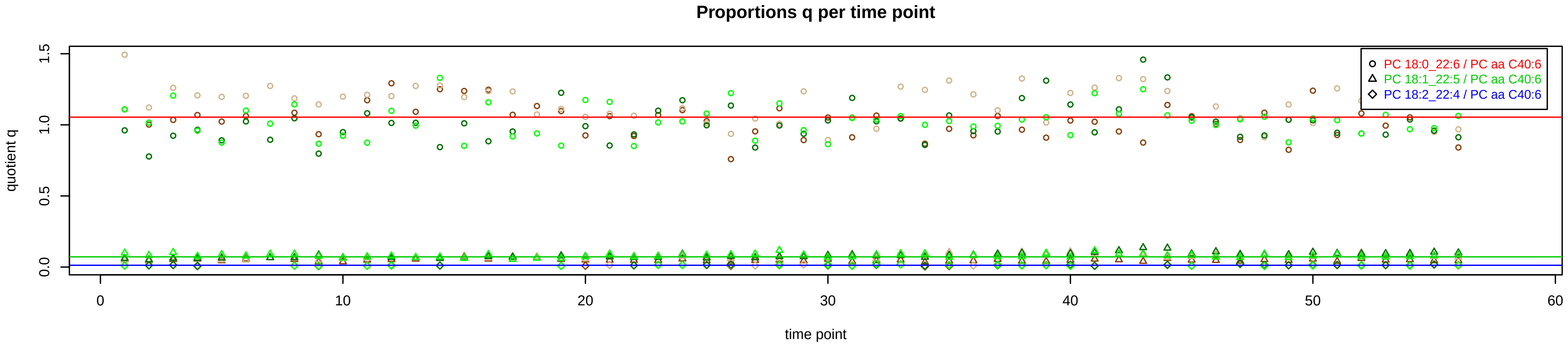

# PC aa C42:0 = R

No independent Variable measured.

### Qualitative composition

PC aa C42:0 consists of:
PC 16:0_26:0, PC 18:0_24:0, PC 20:0_22:0, PC 21:0_21:0,
and further compounds

# PC aa C42:1 = R

No independent Variable measured.

### Qualitative composition

PC aa C42:1 consists of:
PC 18:0_24:1, PC 18:1_24:0, PC 20:0_22:1, PC 20:1_22:0,
and further compounds

# PC aa C42:2 = R

No independent Variable measured.

### Qualitative composition

PC aa C42:2 consists of:
PC 16:0_26:2, PC 18:1_24:1, PC 18:2_24:0, PC 20:0_22:2,
PC 20:2_22:0
and further compounds

# PC aa C42:4 = R

**No independent Variable measured.**

### Qualitative composition

PC aa C42:4 consists of:
PC 18:3_24:1, PC 18:4_24:0, PC 20:0_22:4, PC 20:2_22:2,
PC 20:4_22:0
and further compounds

# PC aa C42:5 = PC 20:0_22:5 + R

PC 20:0_22:5 excluded because of missingness > 75%

**No independent Variable with coverage >0.25 out of PC 20:0_22:5**

### Qualitative composition

PC aa C42:5 consists of:
PC 20:0_22:5, PC 20:1_22:4, PC 20:3_22:2, PC 20:4_22:1, PC 20:5_22:0
and further compounds

# PC aa C42:6 = PC 20:0_22:6 + R

#### Semi−Quantitative composition within Lipidyzer

n = 145

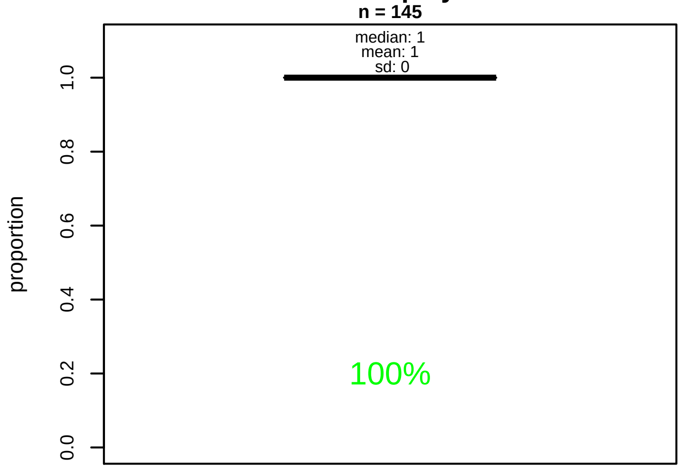


#### Semi−Quantitative composition quotients q

n = 144

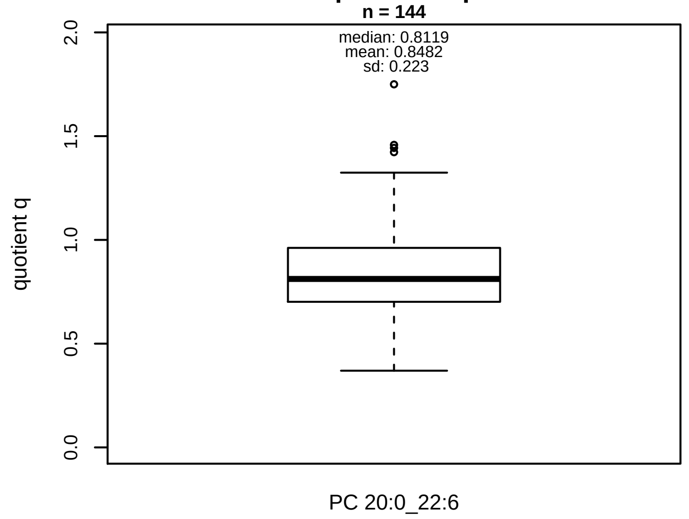


#### Semi−Quantitative composition: Lipidyzer

PC aa C42:6 consists of:
PC 20:0_22:6 100%
and of (not quantified):
PC 20:5_22:1
and further compounds

#### Composition: mean of proportions q

conc(PC aa C42:6) * 0.8482 = conc(PC 20:0_22:6) [var(q)=0.8482]
Percentiles: 5%−>0.5587, 25%−>0.703, 75%−>0.9614, 95%−>1.2468

#### Linear model

```
PC aa C42:6 ~ b * ( PC 20:0_22:6 )
```

b = 0.21904

$R^2$ = 0.05859

#### Ranges

| Measure | AbsoluteIDQ | sum(Lipidyzer) | delta |
|---|---|---|---|
| Min | 0.41 | 0.27 | 0.14 |
| Max | 0.97 | 0.86 | 0.11 |
| Mean | 0.69 | 0.58 | 0.11 |
| Median | 0.69 | 0.57 | 0.13 |
| SD | 0.13 | 0.13 | 0 |

#### Concentration: AbsoluteIDQ vs Lipidyzer

completeness = 65.02%

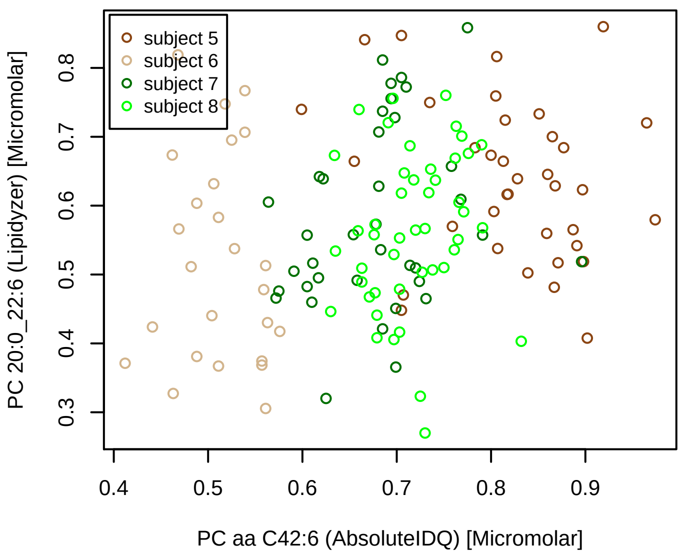


#### Bland−Altman Diagram

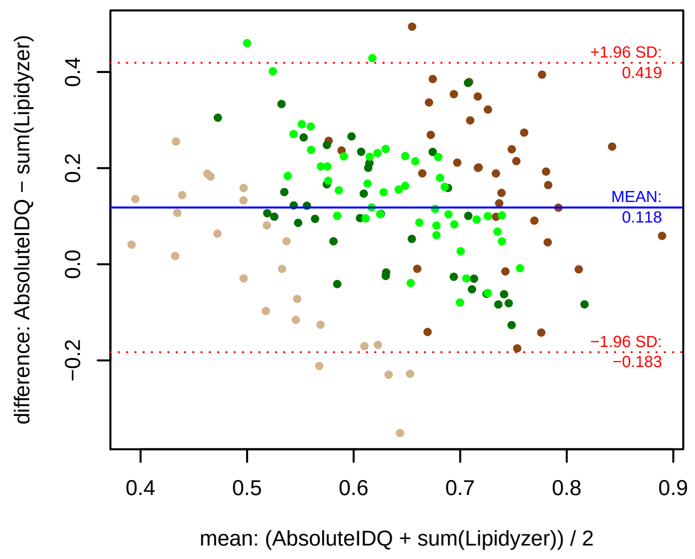


#### Stability of composition

**PC 20:0_22:6 / PC aa C42:6**
**Shapiro−Wilk Test of log quotients, pv: 0.79973; OK**
SW: no; Kruskal: Comp. ~ Subject_ID −> pv = 0.00367; DIFFERENCE
Wilcoxon: Challenge; fasting: 0.5; sport: 1; OLTT: 1

#### Quant. comp., Subject 5

n = 36; comb.: 36

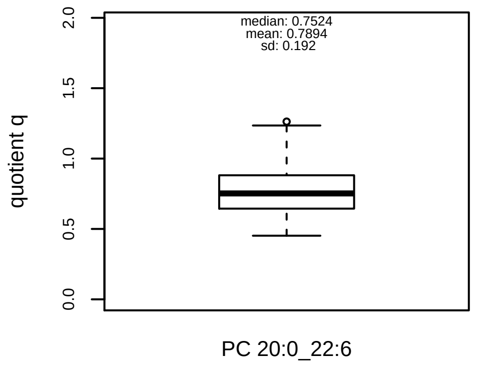


#### Quant. comp., Subject 6

n = 25; comb.: 25

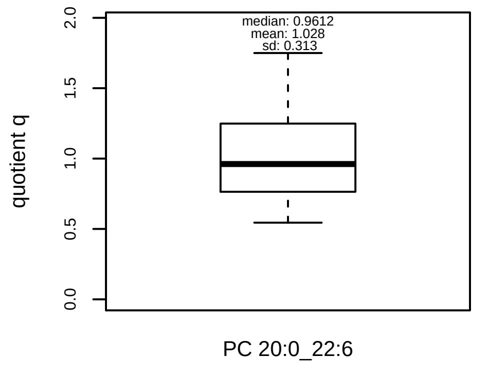


#### Quant. comp., Subject 7

n = 37; comb.: 37

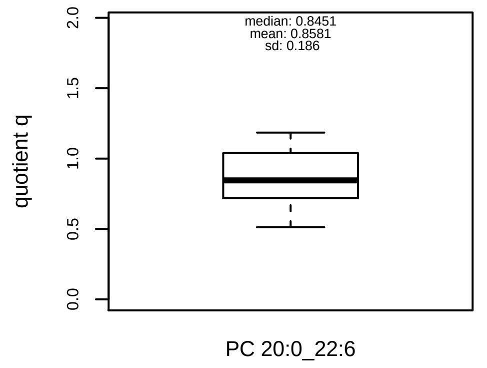


#### Quant. comp., Subject 8

n = 46; comb.: 46

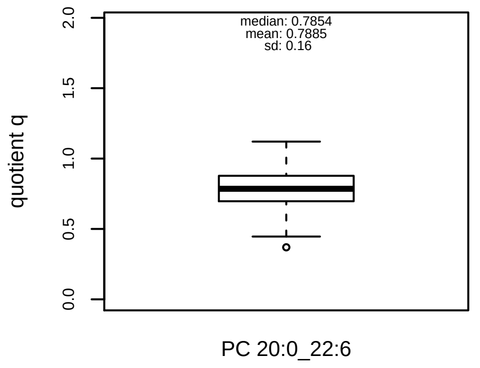


#### Trends of proportions q during challenges

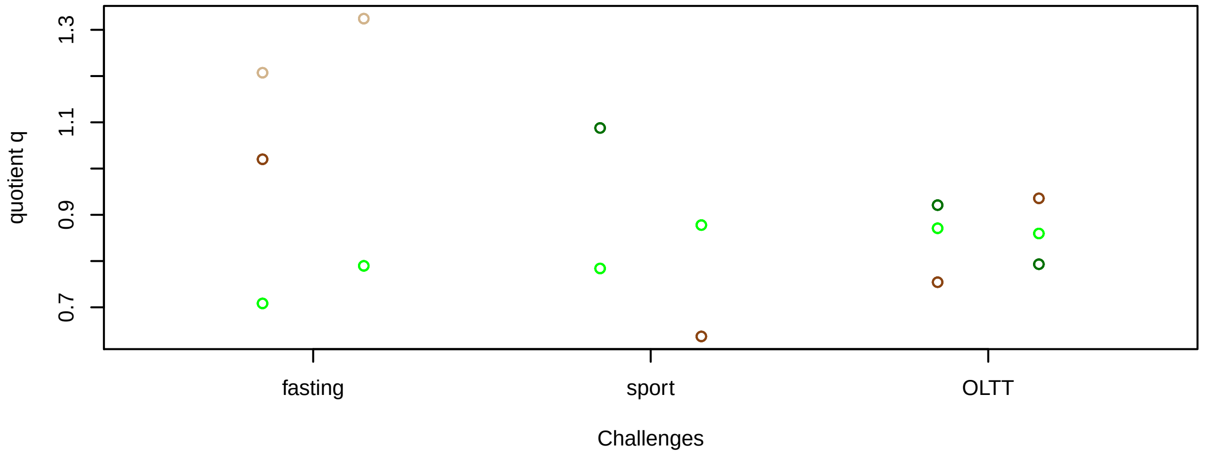


#### Proportions q per time point

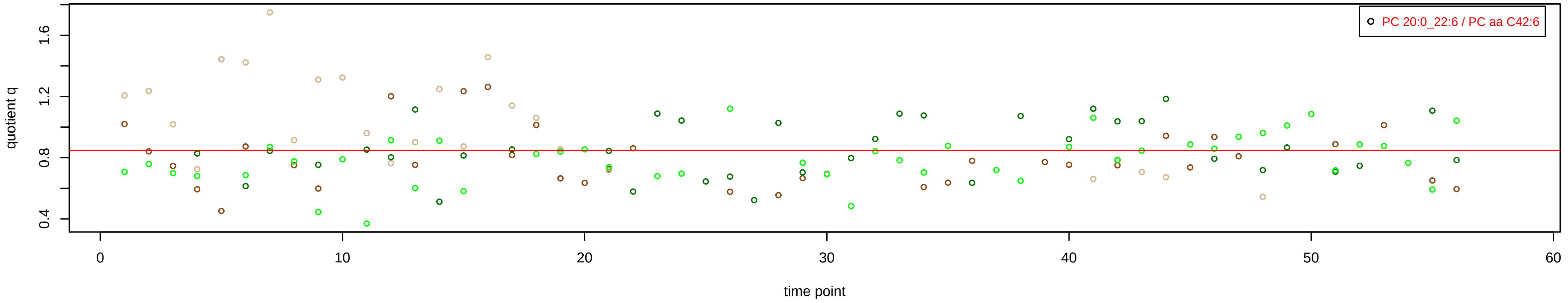

# PC ae C30:0 = R

### Qualitative composition

PC ae C30:0 consists of:
PC O−14:0_16:0, PC O−18:0_12:0, PC 10:0_19:0, PC 12:0_1
PC 13:0_16:0, PC 14:0_15:0, PC 20:0_9:0, PC 8:0_21:0,
[13C1]SM 33:0
and further compounds

**No independent Variable measured.**

# PC ae C32:1 = PC 15:0_16:1 + PC 17:0_14:1 + R

PC 15:0_16:1, PC 17:0_14:1 excluded because of missingness > 75%

**No independent Variable with coverage >0.25 out of PC 15:0_16:1, PC 17:0_14:1**

### Qualitative composition

PC ae C32:1 consists of:
PC 15:0_16:1, PC 17:0_14:1, PC O−14:0_18:1, PC O−16:0_[illegible]
PC O−18:0_14:1, PC O−20:1_12:0, PC 12:0_19:1, PC 13:0_[illegible]
PC 16:0_15:1
and further compounds

# PC ae C32:2 = R

No independent Variable measured.

### Qualitative composition

PC ae C32:2 consists of:
PC O−14:0_18:2, PC O−14:1_18:1, PC O−16:1_16:1, PC 13:
PC 15:1_16:1
and further compounds

# PC ae C34:0 = R

**No independent Variable measured.**

### Qualitative composition

PC ae C34:0 consists of:
PC O−16:0_18:0, PC O−17:0_17:0, PC O−20:0_14:0, PC 10:
PC 11:0_22:0, PC 13:0_20:0, PC 15:0_18:0, PC 16:0_17:0,
PC 21:0_12:0
and further compounds

# PC ae C34:1 = PC 15:0_18:1 + PC 17:0_16:1 + R

#### Semi−Quantitative composition within Lipidyzer
n = 122

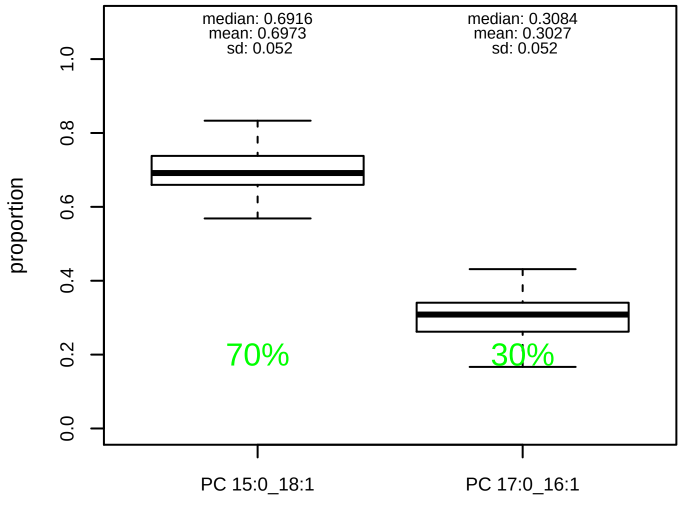


#### Semi−Quantitative composition quotients q
n = 121

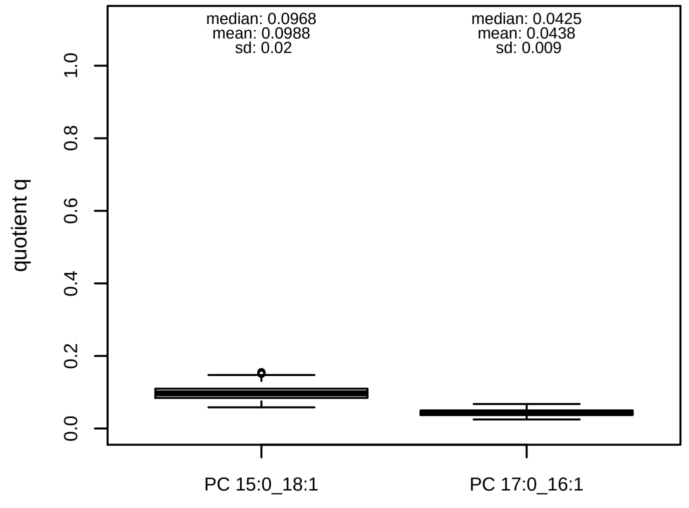


#### Semi−Quantitative composition: Lipidyzer

PC ae C34:1 consists of:
PC 15:0_18:1 69.7%
PC 17:0_16:1 30.3%
and of (not quantified):
PC O−16:0_18:1, PC O−18:0_16:1, PC O−20:0_14:1, PC O−PC 13:0_20:1, PC 14:0_19:1, PC 14:1_19:0, PC 15:1_18:0, [13C1]SM 37:1
and further compounds

#### Linear model

```
PC ae C34:1 ~ b * ( PC 15:0_18:1
   + PC 17:0_16:1 )
```

b = 4.14514

$R^2$ = 0.57752

#### Ranges

| Measure | AbsoluteIDQ | sum(Lipidyzer) | delta |
|---|---|---|---|
| Min | 5.38 | 0.78 | 4.6 |
| Max | 15.34 | 2.26 | 13.08 |
| Mean | 9.27 | 1.45 | 7.82 |
| Median | 8.81 | 1.51 | 7.3 |
| SD | 2.43 | 0.35 | 2.08 |

#### Composition: mean of proportions q

conc(PC ae C34:1) * 0.0988 = conc(PC 15:0_18:1) [var(q)=0.0988]
Percentiles: 5%−>0.0705, 25%−>0.0843, 75%−>0.1097, 95%−>0.1373
conc(PC ae C34:1) * 0.0438 = conc(PC 17:0_16:1) [var(q)=0.0438]
Percentiles: 5%−>0.0307, 25%−>0.0371, 75%−>0.0499, 95%−>0.0601

#### Concentration: AbsoluteIDQ vs Lipidyzer
completeness = 100%

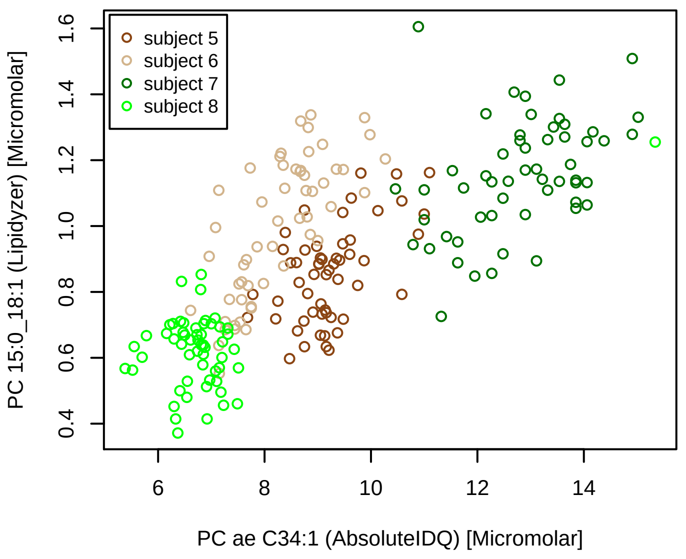


#### Concentration: AbsoluteIDQ vs Lipidyzer
completeness = 54.71%

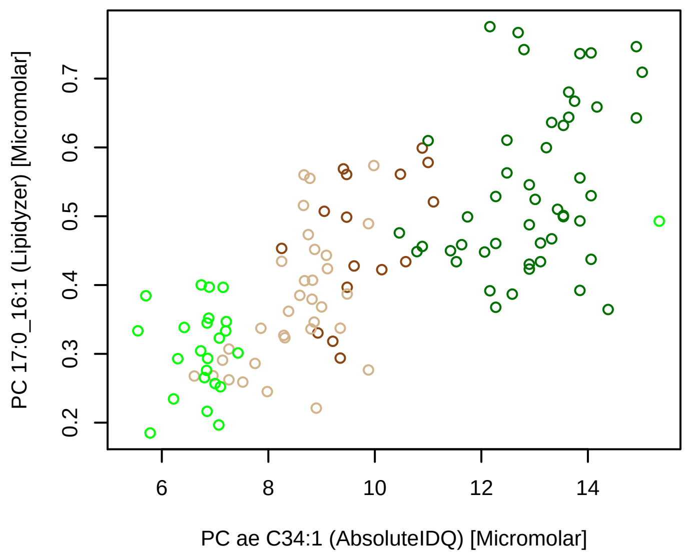


#### Bland−Altman Diagram

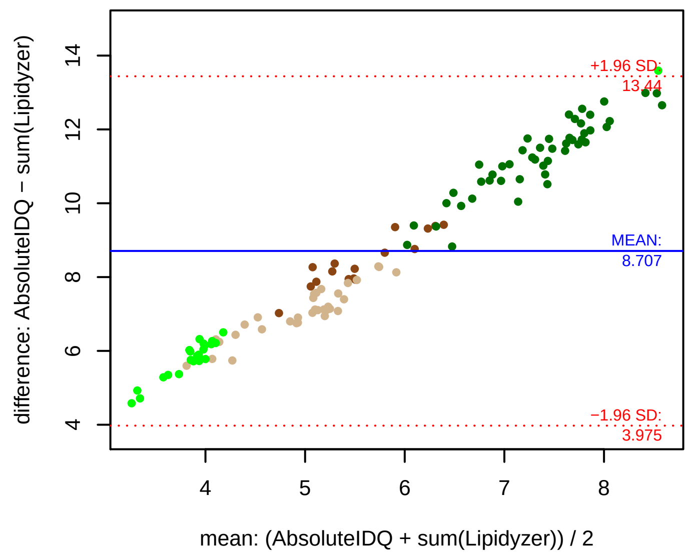


#### Stability of composition

**(PC 15:0_18:1 + PC 17:0_16:1) / PC ae C34:1**
**Shapiro−Wilk Test of log quotients, pv: 0.31322; OK**
SW: ok; ANOVA: Comp. ~ Subject_ID −> pv = 0.0045; DIFFERENCE
Wilcoxon: Challenge; sport: 0.5; OLTT: 1

**PC 15:0_18:1 / PC ae C34:1**
**Shapiro−Wilk Test of log quotients, pv: 0.37336; OK**
SW: ok; ANOVA: Comp. ~ Subject_ID −> pv = 0.01068; EQUAL
Wilcoxon: Challenge; fasting: 0.25; sport: 0.625; OLTT: 0.875

**PC 17:0_16:1 / PC ae C34:1**
**Shapiro−Wilk Test of log quotients, pv: 0.65757; OK**
SW: ok; ANOVA: Comp. ~ Subject_ID −> pv = 0.322; EQUAL
Wilcoxon: Challenge; sport: 0.75; OLTT: 1

#### Quant. comp., Subject 5
n = 55, 16; comb.: 16

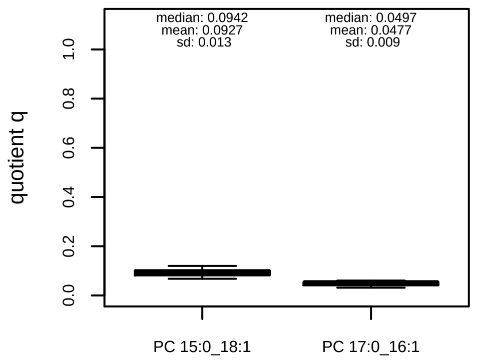


#### Quant. comp., Subject 6
n = 55, 33; comb.: 33

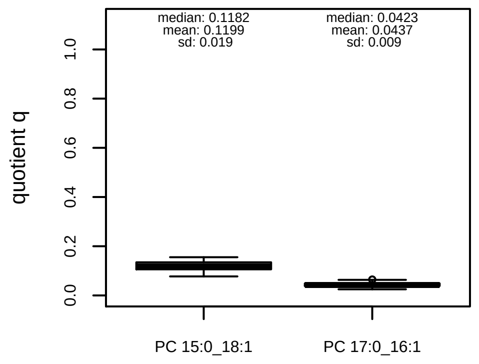


#### Quant. comp., Subject 7
n = 55, 48; comb.: 48

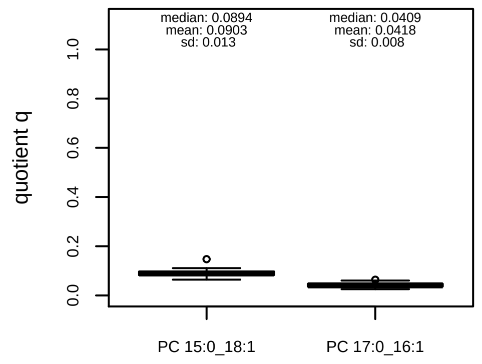


#### Quant. comp., Subject 8
n = 56, 24; comb.: 24

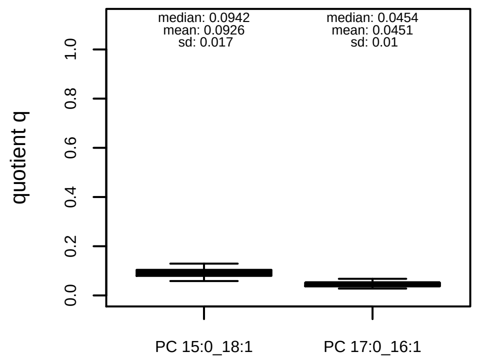


#### Trends of proportions q during challenges

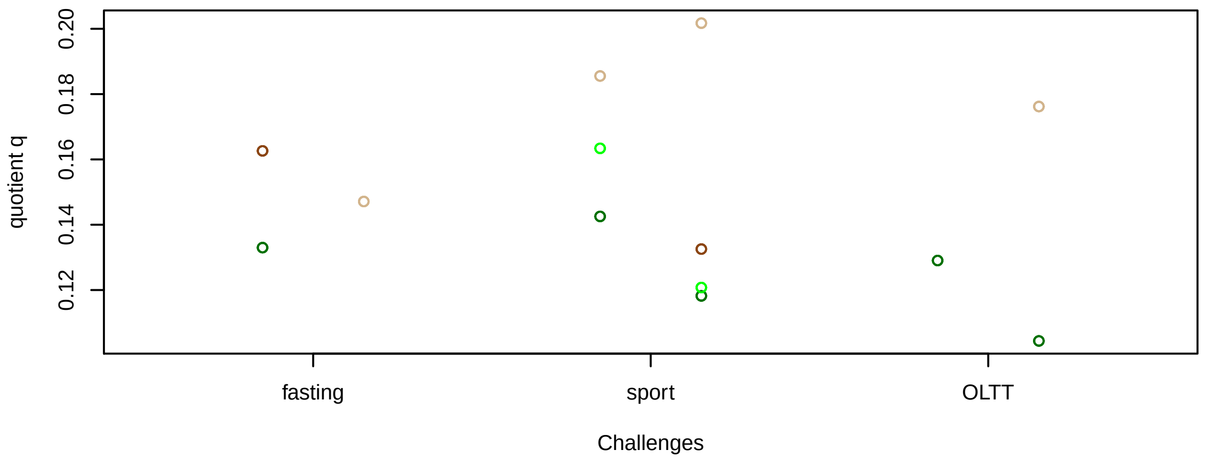


#### Proportions q per time point

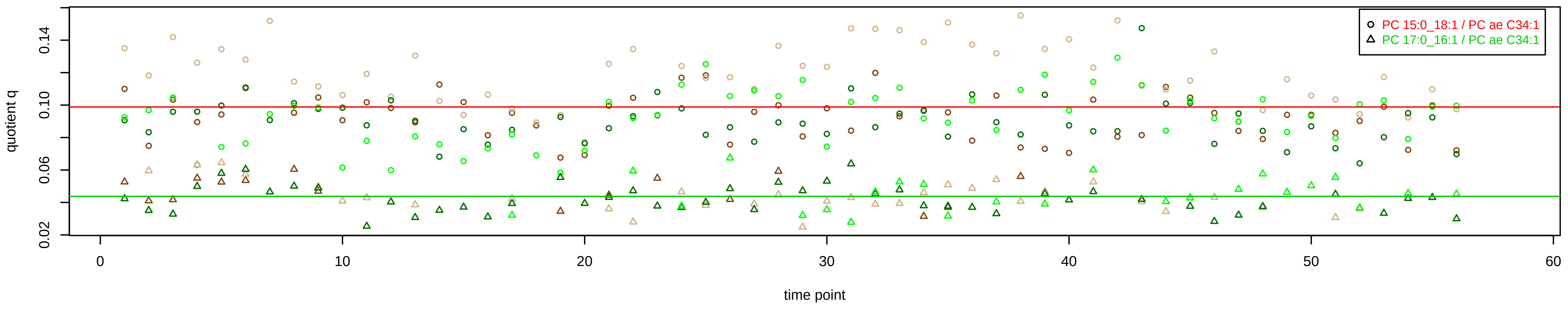

# PC ae C34:2 = PC 15:0_18:2 + R

**Semi−Quantitative composition within Lipidyzer**

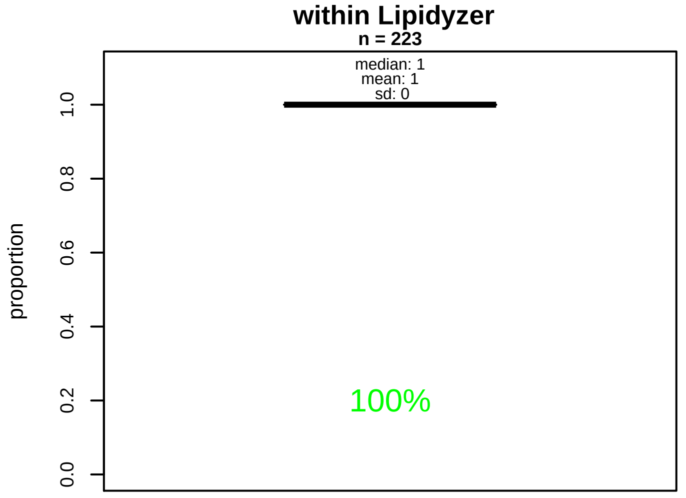


**Semi−Quantitative composition quotients q**

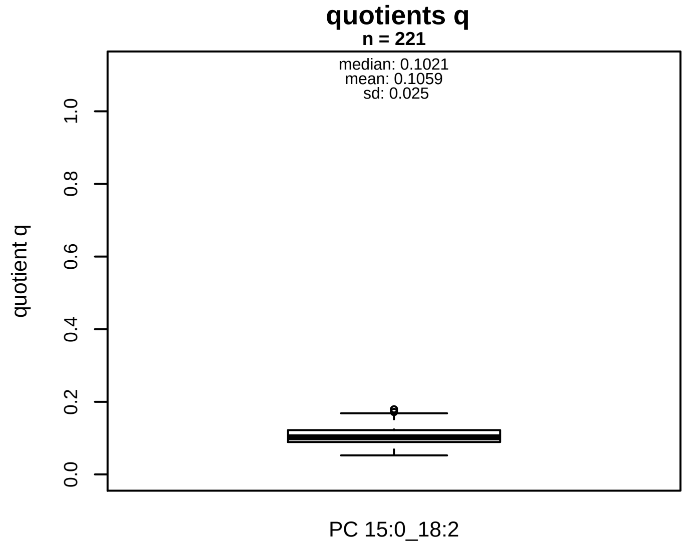


#### Semi−Quantitative composition: Lipidyzer

PC ae C34:2 consists of:
PC 15:0_18:2 100%
and of (not quantified):
PC O−16:0_18:2, PC O−16:1_18:1, PC O−20:1_14:1, PC 13:
PC 14:1_19:1, PC 15:1_18:1, PC 16:0_17:2, PC 16:1_17:1,
and further compounds

#### Composition: mean of proportions q

conc(PC ae C34:2) * 0.1059 = conc(PC 15:0_18:2) [var(q)=0.1059]
Percentiles: 5%−>0.0698, 25%−>0.0892, 75%−>0.1219, 95%−>0.1541

#### Linear model

```
PC ae C34:2 ~ b * ( PC 15:0_18:2 )
```

b = 3.24787

$R^2$ = 0.07136

#### Ranges

| Measure | AbsoluteIDQ | sum(Lipidyzer) | delta |
|---|---|---|---|
| Min | 6.73 | 0.63 | 6.1 |
| Max | 16.96 | 1.65 | 15.31 |
| Mean | 10.48 | 1.07 | 9.41 |
| Median | 10.19 | 1.07 | 9.12 |
| SD | 2.28 | 0.19 | 2.1 |

**Concentration: AbsoluteIDQ vs Lipidyzer**

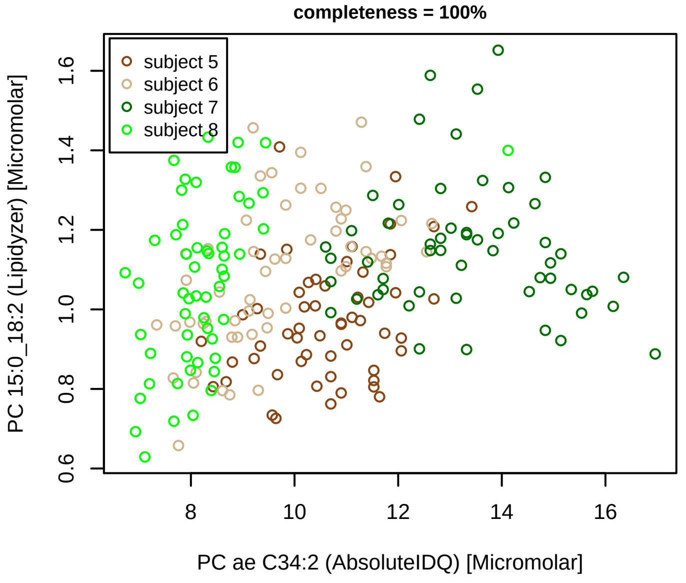


**Bland−Altman Diagram**

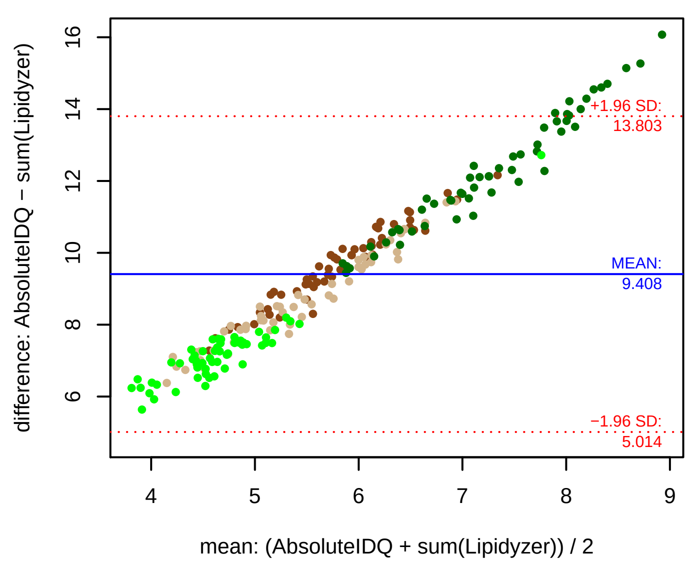


#### Stability of composition

**PC 15:0_18:2 / PC ae C34:2**
**Shapiro−Wilk Test of log quotients, pv: 0.62993; OK**
SW: ok; ANOVA: Comp. ~ Subject_ID −> pv = 0; DIFFERENCE
Wilcoxon: Challenge; fasting: 0.625; sport: 0.375; OLTT: 1

**Quant. comp., Subject 5**

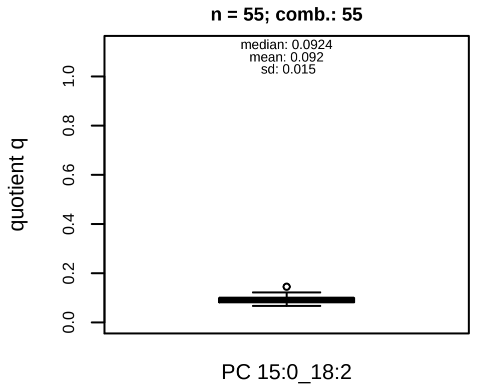


**Quant. comp., Subject 6**

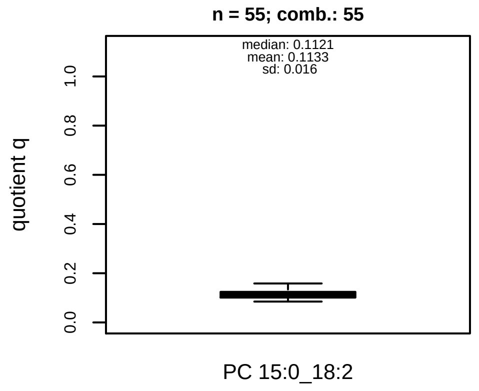


**Quant. comp., Subject 7**

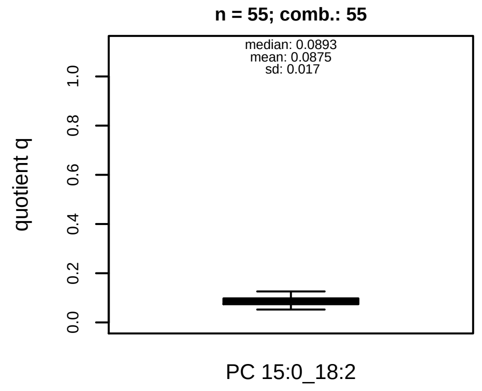


**Quant. comp., Subject 8**

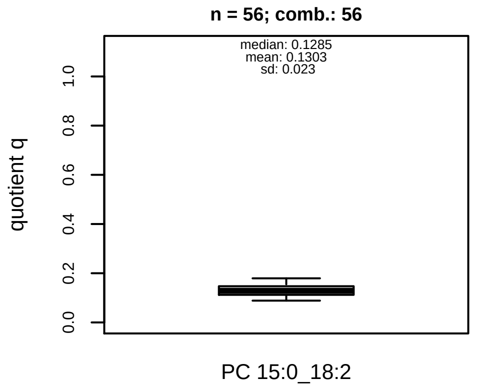


**Trends of proportions q during challenges**

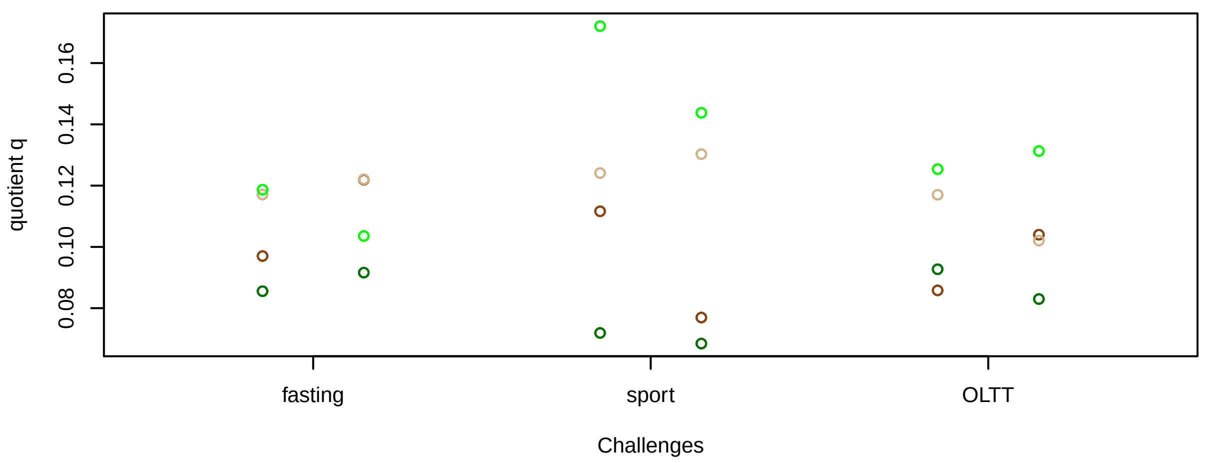


**Proportions q per time point**

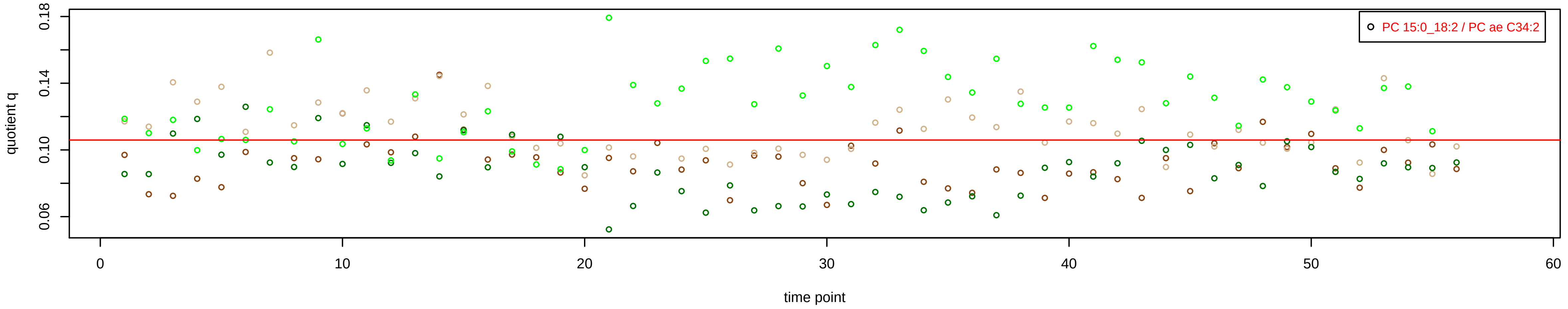

# PC ae C34:3 = PC 15:0_18:3 + R

PC 15:0_18:3 excluded because of missingness > 75%

**No independent Variable with coverage >0.25 out of PC 15:0_18:3**

### Qualitative composition

PC ae C34:3 consists of:
PC 15:0_18:3, PC O−16:0_18:3, PC O−16:1_18:2, PC 13:0_2
PC 16:1_17:2
and further compounds

# PC ae C36:1 = PC 17:0_18:1 + R

#### Semi−Quantitative composition within Lipidyzer
n = 223

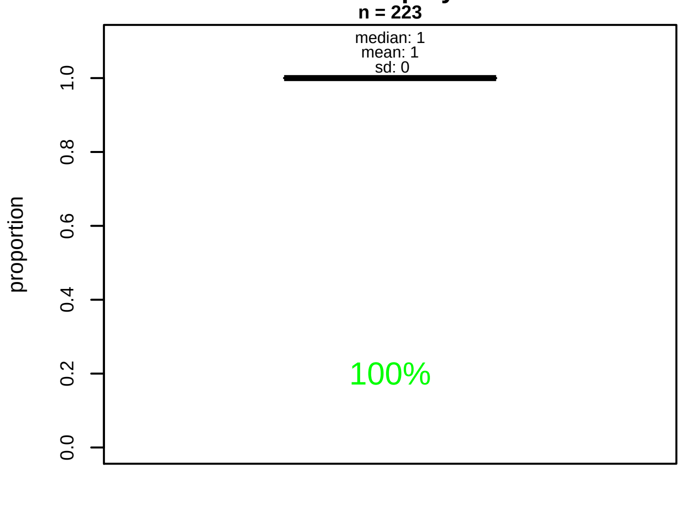


#### Semi−Quantitative composition quotients q
n = 221

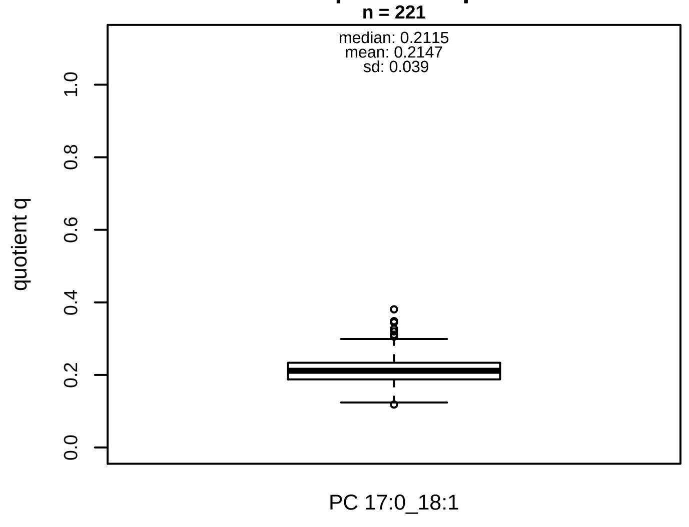


#### Semi−Quantitative composition: Lipidyzer

PC ae C36:1 consists of:
PC 17:0_18:1 100%
and of (not quantified):
PC O−16:0_20:1, PC O−18:0_18:1, PC O−20:0_16:1, PC O−
PC 13:0_22:1, PC 14:1_21:0, PC 15:0_20:1, PC 15:1_20:0,
PC 16:0_19:1, PC 16:1_19:0, PC 17:1_18:0, PC 18:4_18:4,
[13C1]SM 39:1
and further compounds

#### Composition: mean of proportions q

conc(PC ae C36:1) * 0.2147 = conc(PC 17:0_18:1) [var(q)=0.2147]
Percentiles: 5%−>0.1601, 25%−>0.1877, 75%−>0.2335, 95%−>0.2878

#### Linear model

```
PC ae C36:1 ~ b * ( PC 17:0_18:1 )
```

b = 4.23026

$R^2$ = 0.64372

#### Ranges

| Measure | AbsoluteIDQ | sum(Lipidyzer) | delta |
|---|---|---|---|
| Min | 3.28 | 0.65 | 2.63 |
| Max | 14.62 | 2.52 | 12.1 |
| Mean | 7.04 | 1.48 | 5.56 |
| Median | 6.97 | 1.5 | 5.47 |
| SD | 2.25 | 0.42 | 1.82 |

#### Concentration: AbsoluteIDQ vs Lipidyzer
completeness = 100%

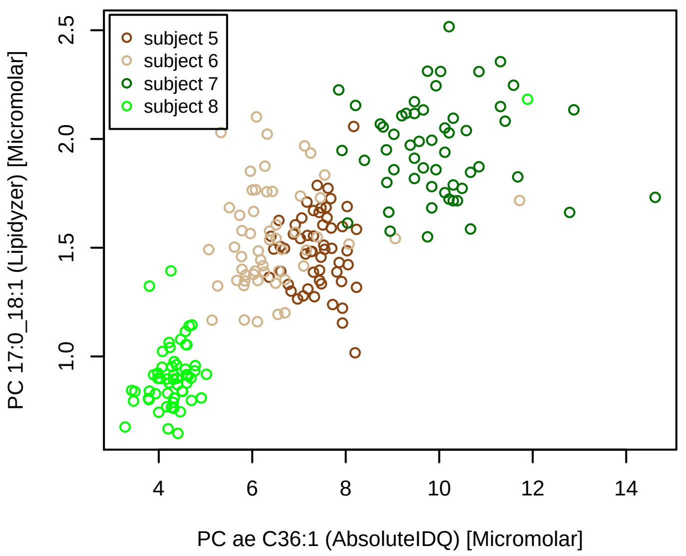


#### Bland−Altman Diagram

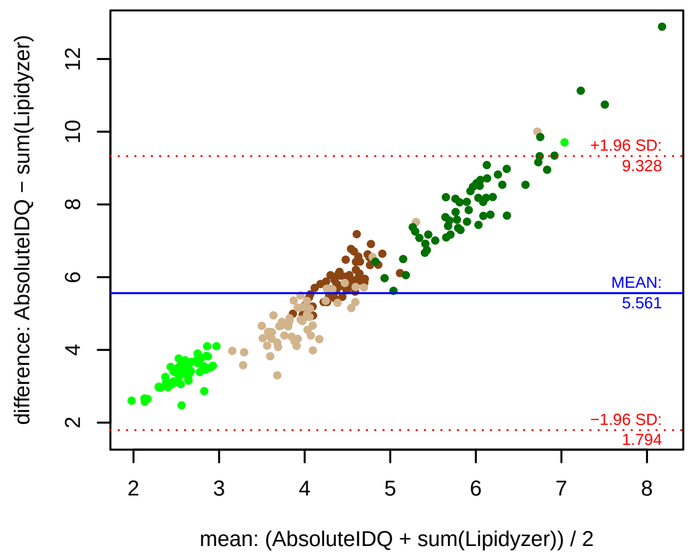


#### Stability of composition

**PC 17:0_18:1 / PC ae C36:1**
**Shapiro−Wilk Test of log quotients, pv: 0.04571; OK**
SW: ok; ANOVA: Comp. ~ Subject_ID −> pv = 0.5455; EQUAL
Wilcoxon: Challenge; fasting: 0.25; sport: 0.375; OLTT: 0.875

#### Quant. comp., Subject 5
n = 55; comb.: 55

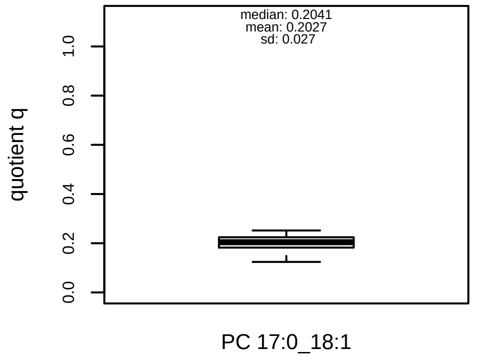


#### Quant. comp., Subject 6
n = 55; comb.: 55

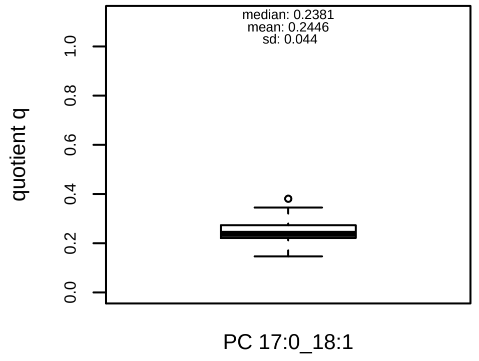


#### Quant. comp., Subject 7
n = 55; comb.: 55

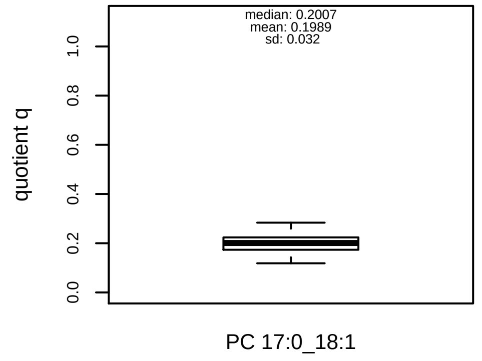


#### Quant. comp., Subject 8
n = 56; comb.: 56

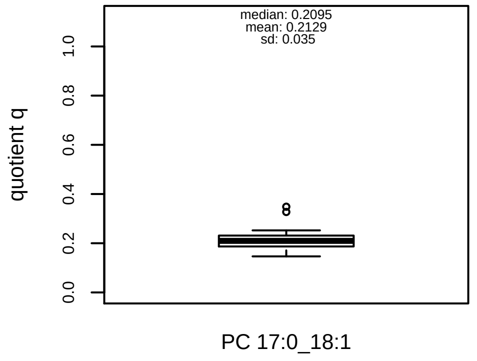


#### Trends of proportions q during challenges

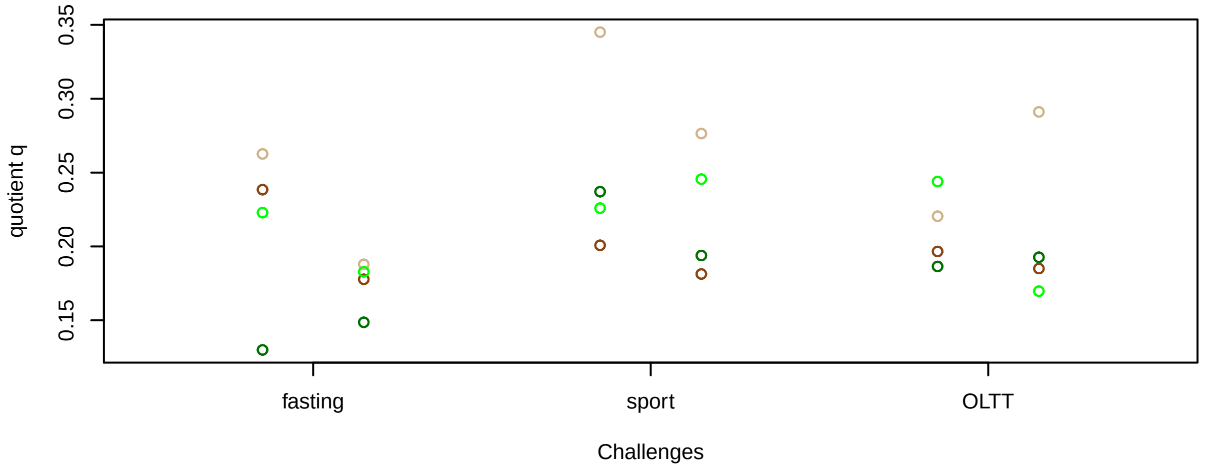


#### Proportions q per time point

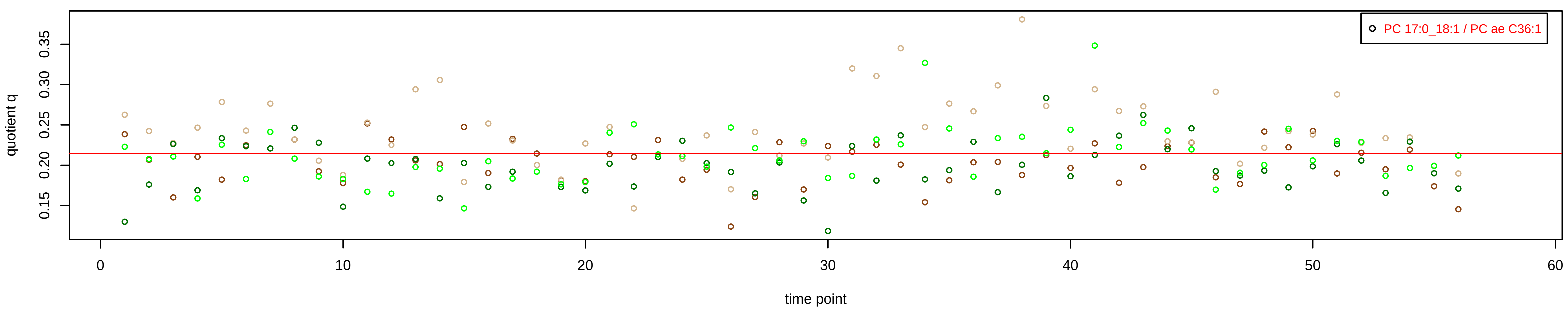

# PC ae C36:2 = PC 17:0_18:2 + R

#### Semi−Quantitative composition within Lipidyzer

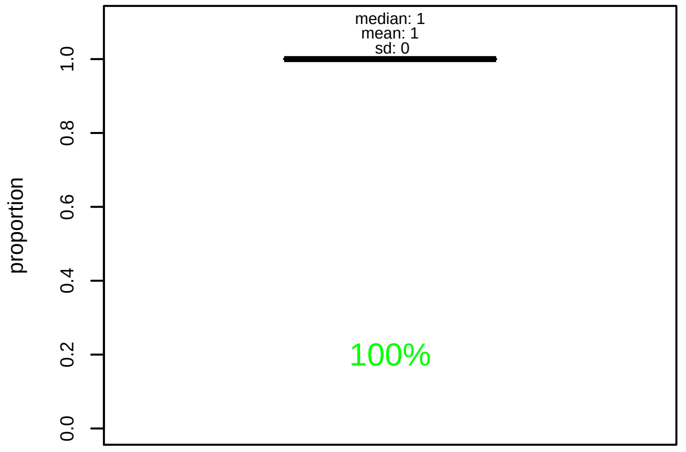


#### Semi−Quantitative composition quotients q

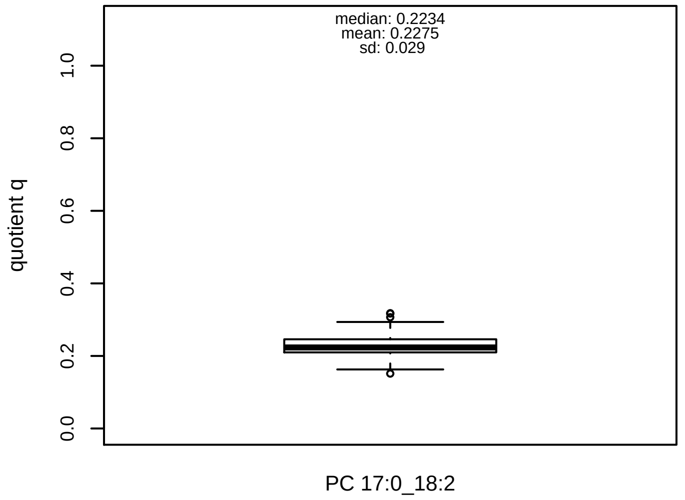


#### Semi−Quantitative composition: Lipidyzer

PC ae C36:2 consists of:
PC 17:0_18:2 100%
and of (not quantified):
PC O−16:0_20:2, PC O−18:0_18:2, PC O−18:1_18:1, PC O−
PC 13:0_22:2, PC 15:0_20:2, PC 15:1_20:1, PC 16:1_19:1,
[13C1]SM 39:2
and further compounds

#### Composition: mean of proportions q

conc(PC ae C36:2) * 0.2275 = conc(PC 17:0_18:2) [var(q)=0.2275]
Percentiles: 5%−>0.1837, 25%−>0.2098, 75%−>0.2459, 95%−>0.2799

#### Linear model

```
PC ae C36:2 ~ b * ( PC 17:0_18:2 )
```

b = 3.27954

$R^2$ = 0.52816

#### Ranges

| Measure | AbsoluteIDQ | sum(Lipidyzer) | delta |
|---|---|---|---|
| Min | 9.2 | 1.76 | 7.44 |
| Max | 19.24 | 4.41 | 14.83 |
| Mean | 13.38 | 3.02 | 10.36 |
| Median | 13.03 | 2.96 | 10.07 |
| SD | 2.27 | 0.51 | 1.76 |

#### Concentration: AbsoluteIDQ vs Lipidyzer

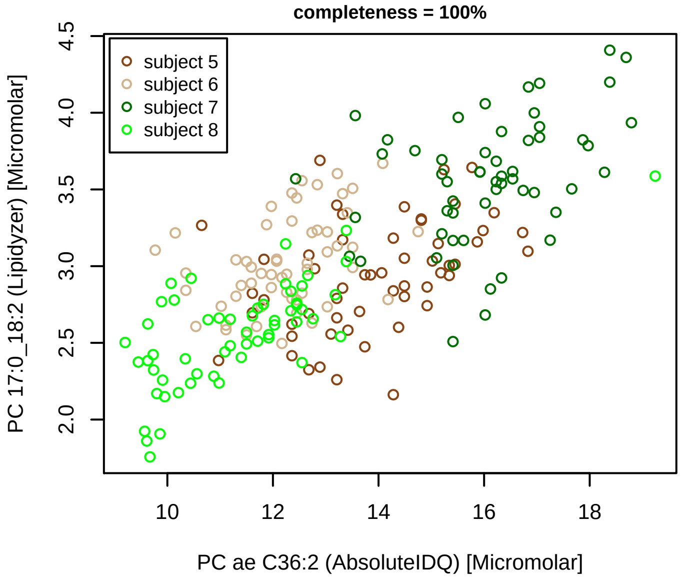


#### Bland−Altman Diagram

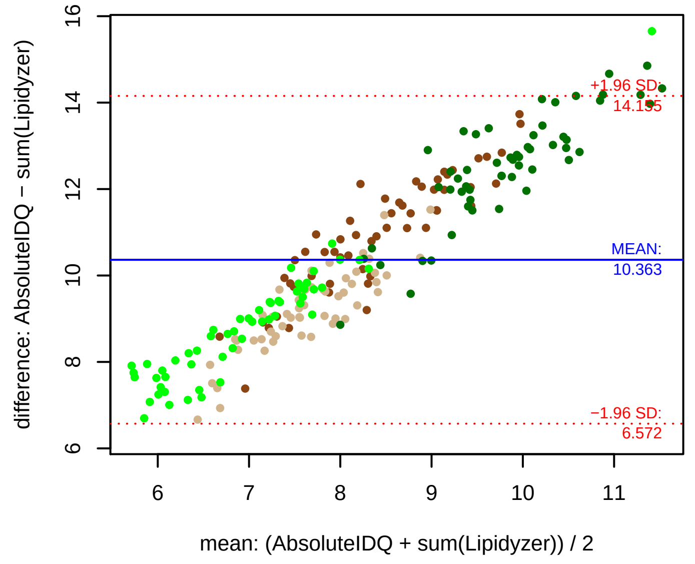


#### Stability of composition

**PC 17:0_18:2 / PC ae C36:2**
**Shapiro−Wilk Test of log quotients, pv: 0.867; OK**
SW: ok; ANOVA: Comp. ~ Subject_ID −> pv = 0.19747; EQUAL
Wilcoxon: Challenge; fasting: 0.625; sport: 0.375; OLTT: 0.125

#### Quant. comp., Subject 5

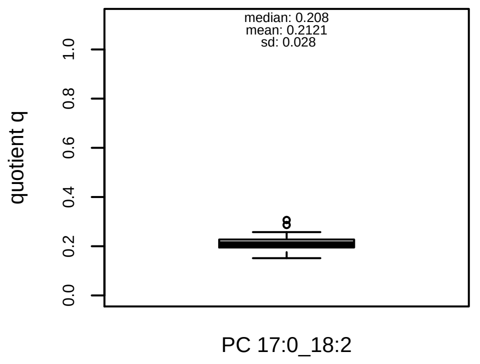


#### Quant. comp., Subject 6

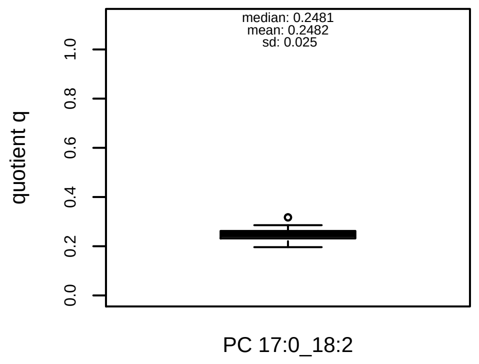


#### Quant. comp., Subject 7

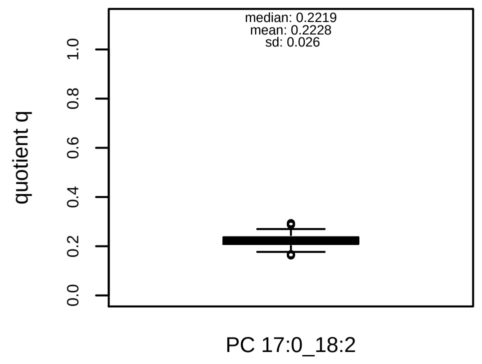


#### Quant. comp., Subject 8

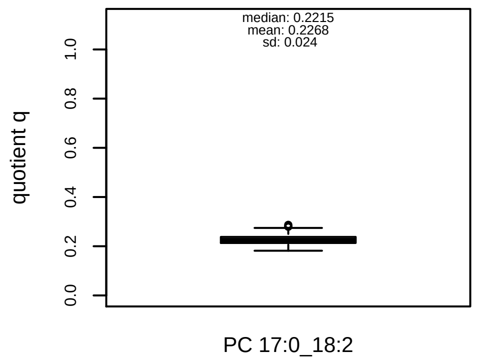


#### Trends of proportions q during challenges

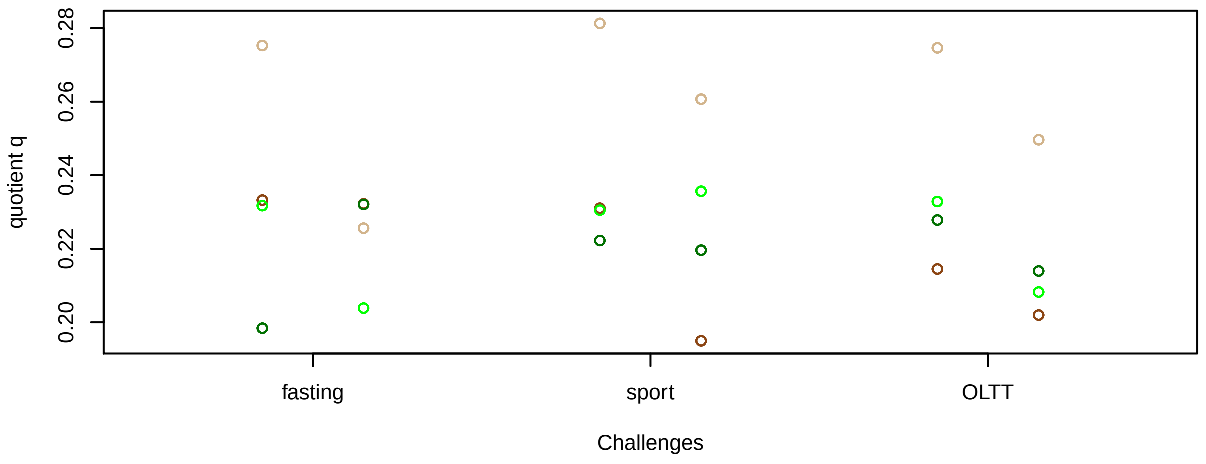


#### Proportions q per time point

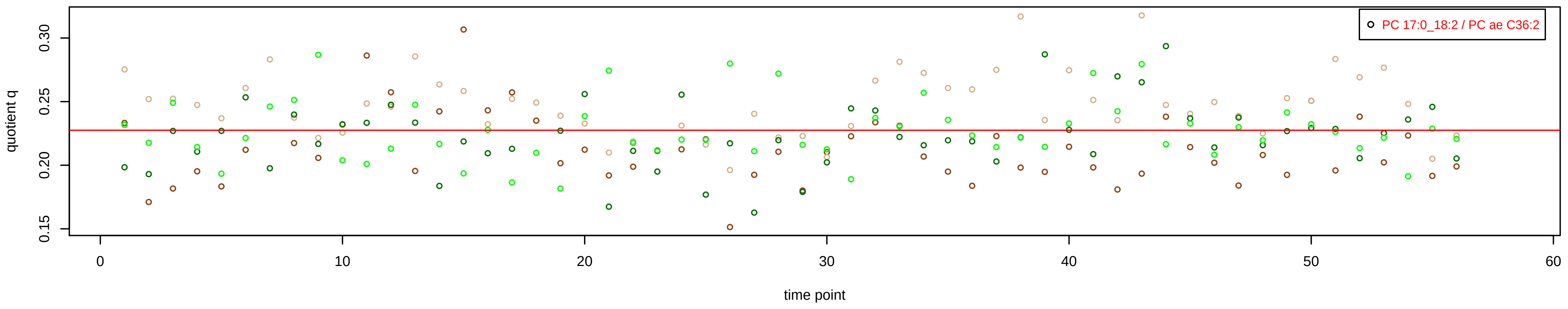

# PC ae C36:3 = PC 15:0_20:3 + PC 17:0_18:3 + R

PC 17:0_18:3 excluded because of missingness > 75%

#### Semi−Quantitative composition within Lipidyzer

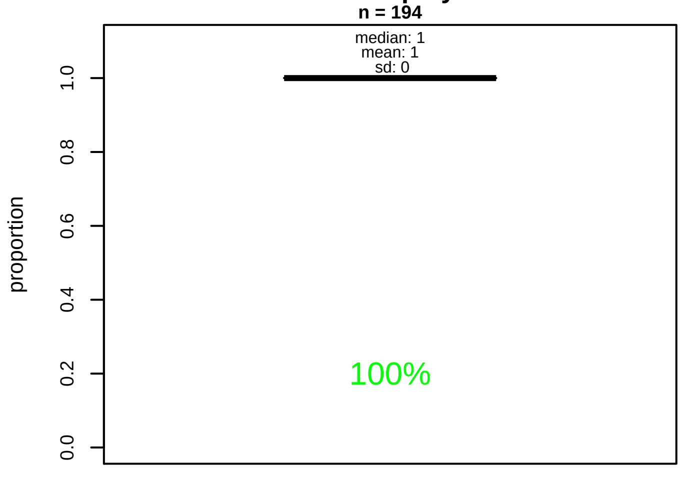


#### Semi−Quantitative composition quotients q

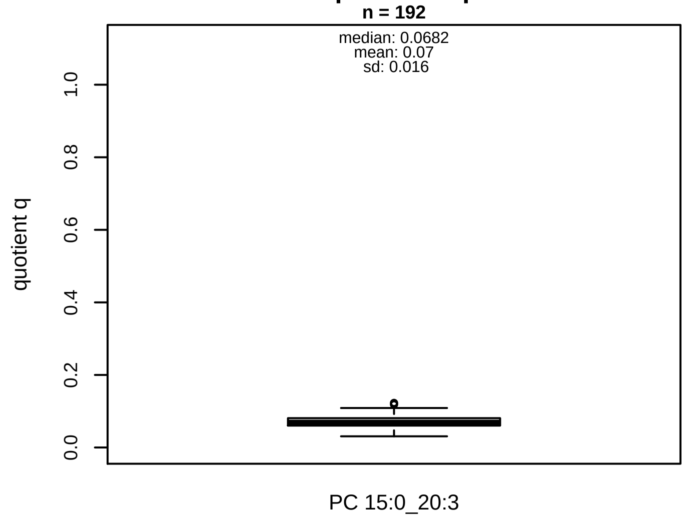


#### Semi−Quantitative composition: Lipidyzer

PC ae C36:3 consists of:
PC 15:0_20:3 100%
and of (not quantified):
PC 17:0_18:3, PC O−16:0_20:3, PC O−18:0_18:3, PC O−18:
PC O−16:1_20:2, PC 15:1_20:2, PC 17:1_18:2, PC 17:2_18:
and further compounds

#### Composition: mean of proportions q

conc(PC ae C36:3) * 0.07 = conc(PC 15:0_20:3) [var(q)=0.07]
Percentiles: 5%−>0.0458, 25%−>0.0603, 75%−>0.0809, 95%−>0.095

#### Linear model

```
PC ae C36:3 ~ b * ( PC 15:0_20:3 )
```

b = 8.36338

$R^2$ = 0.49946

#### Ranges

| Measure | AbsoluteIDQ | sum(Lipidyzer) | delta |
|---|---|---|---|
| Min | 4.88 | 0.19 | 4.69 |
| Max | 13.58 | 1.29 | 12.29 |
| Mean | 7.83 | 0.55 | 7.28 |
| Median | 7.65 | 0.54 | 7.11 |
| SD | 1.97 | 0.17 | 1.8 |

#### Concentration: AbsoluteIDQ vs Lipidyzer

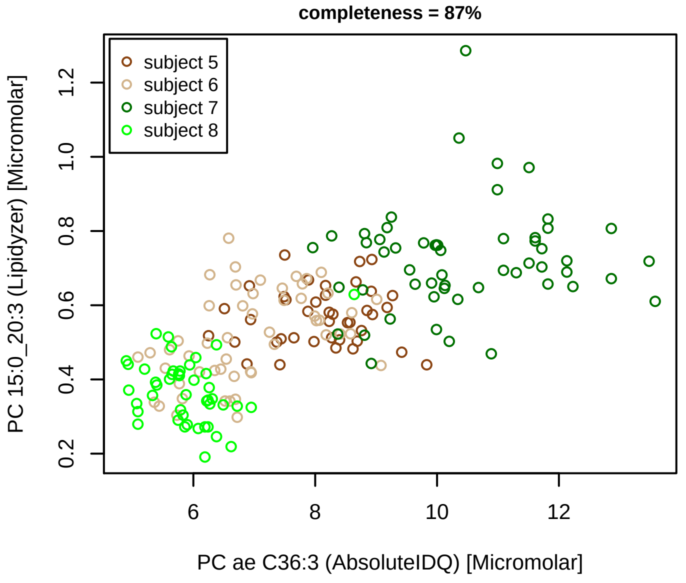


#### Bland−Altman Diagram

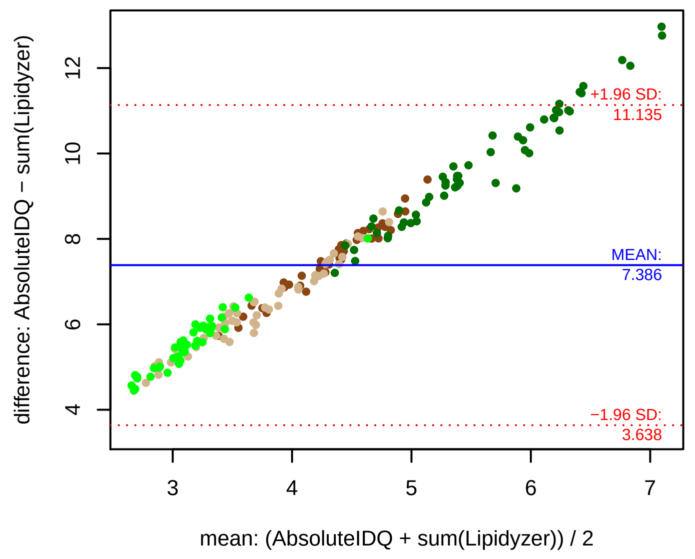


#### Stability of composition

**PC 15:0_20:3 / PC ae C36:3**
**Shapiro−Wilk Test of log quotients, pv: 0.11199; OK**
SW: ok; ANOVA: Comp. ~ Subject_ID −> pv = 0.00182; DIFFERENCE
Wilcoxon: Challenge; fasting: 0.25; sport: 0.75; OLTT: 0.625

#### Quant. comp., Subject 5

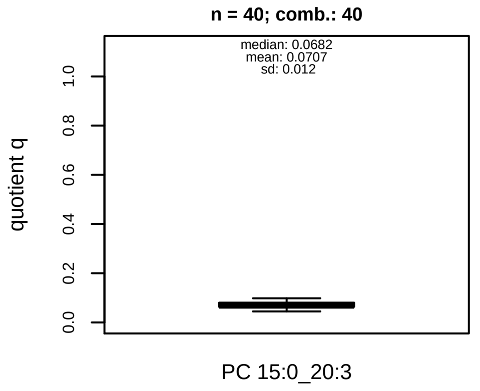


#### Quant. comp., Subject 6

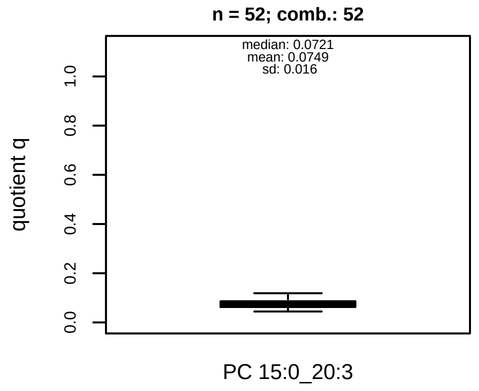


#### Quant. comp., Subject 7

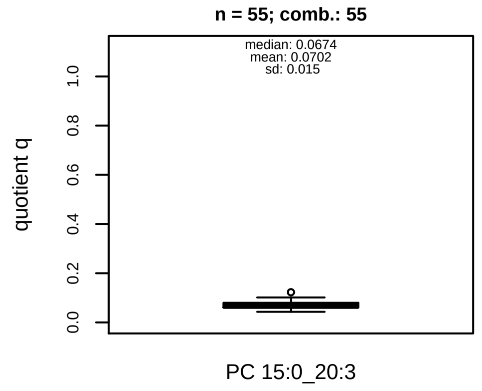


#### Quant. comp., Subject 8

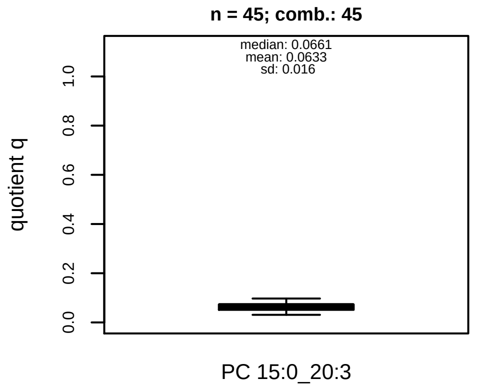


#### Trends of proportions q during challenges

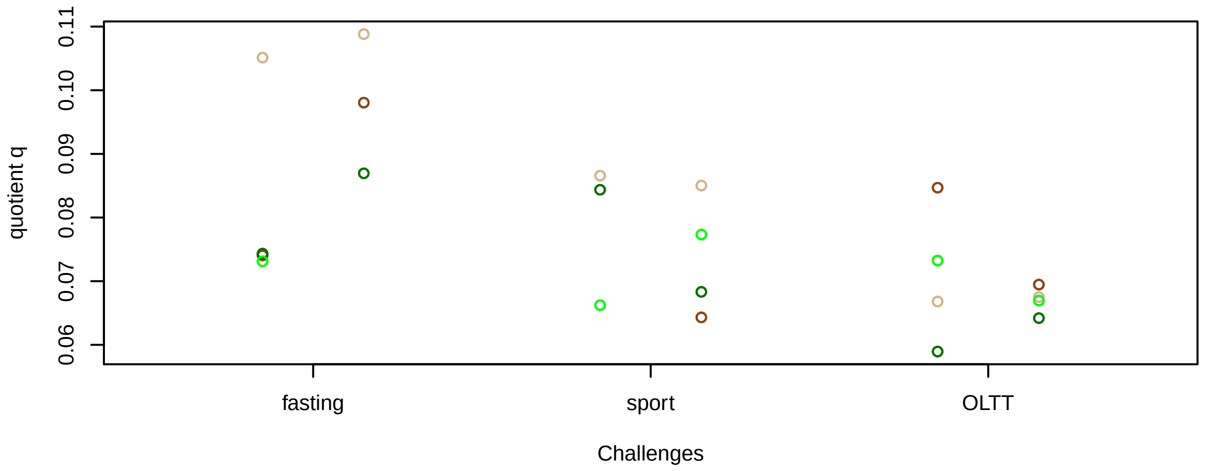


#### Proportions q per time point

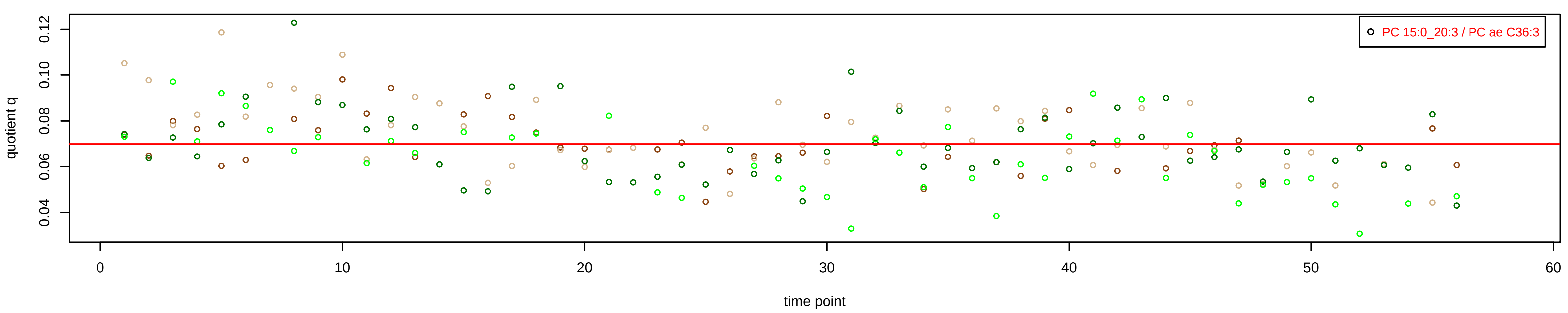

# PC ae C36:4 = PC 15:0_20:4 + R

#### Semi−Quantitative composition within Lipidyzer

n = 223

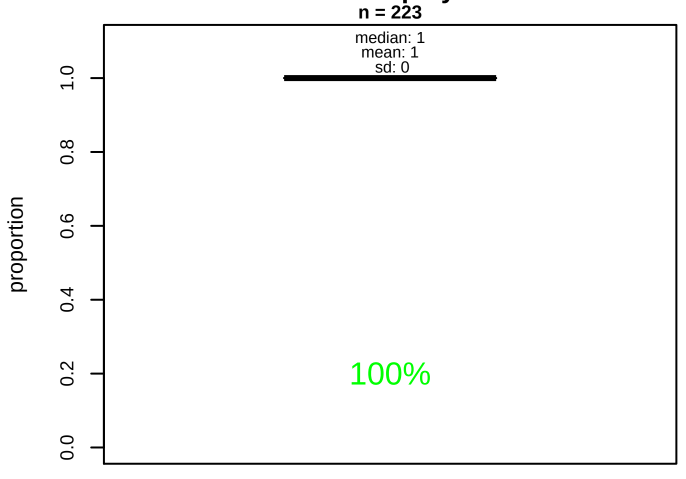


#### Semi−Quantitative composition quotients q

n = 221

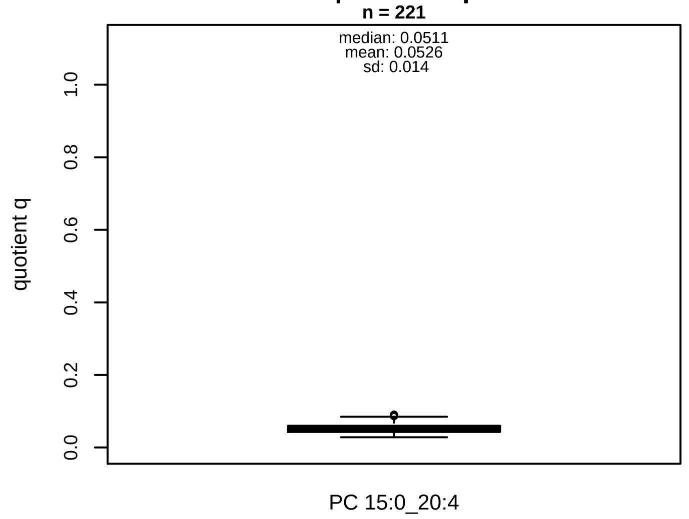


#### Semi−Quantitative composition: Lipidyzer

PC ae C36:4 consists of:
PC 15:0_20:4 100%
and of (not quantified):
PC O−16:0_20:4, PC O−16:1_20:3, PC O−18:0_18:4, PC O−
PC O−18:2_18:2, PC 13:0_22:4, PC 15:1_20:3, PC 17:0_18:4
PC 17:2_18:2
and further compounds

#### Composition: mean of proportions q

conc(PC ae C36:4) * 0.0526 = conc(PC 15:0_20:4) [var(q)=0.0526]
Percentiles: 5%−>0.0339, 25%−>0.0429, 75%−>0.0599, 95%−>0.0793

#### Linear model

```
PC ae C36:4 ~ b * ( PC 15:0_20:4 )
```

b = 11.15708

$R^2$ = 0.25993

#### Ranges

| Measure | AbsoluteIDQ | sum(Lipidyzer) | delta |
|---|---|---|---|
| Min | 8.59 | 0.45 | 8.14 |
| Max | 26.01 | 1.8 | 24.21 |
| Mean | 17.54 | 0.88 | 16.66 |
| Median | 18.55 | 0.86 | 17.69 |
| SD | 4.63 | 0.21 | 4.42 |

#### Concentration: AbsoluteIDQ vs Lipidyzer

completeness = 100%

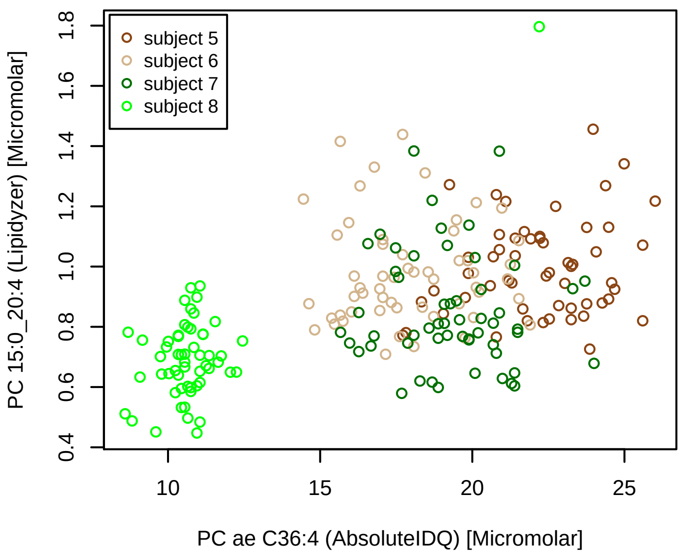


#### Bland−Altman Diagram

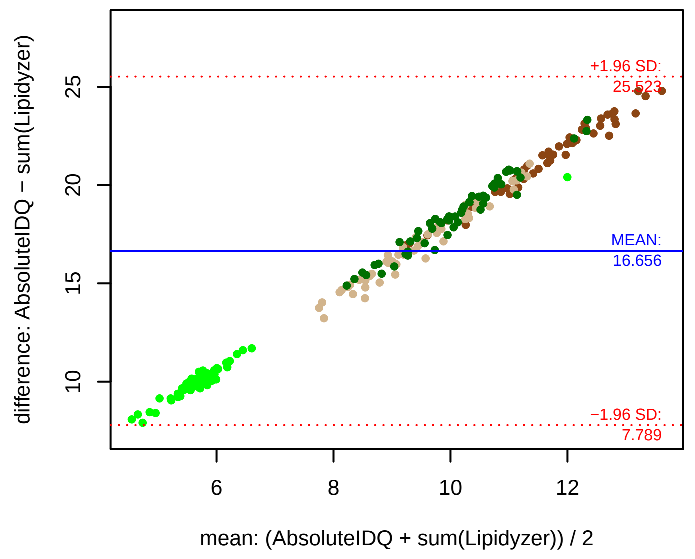


#### Stability of composition

**PC 15:0_20:4 / PC ae C36:4**
**Shapiro−Wilk Test of log quotients, pv: 0.32439; OK**
SW: ok; ANOVA: Comp. ~ Subject_ID −> pv = 0; DIFFERENCE
Wilcoxon: Challenge; fasting: 1; sport: 0.875; OLTT: 0.25

#### Quant. comp., Subject 5

n = 55; comb.: 55

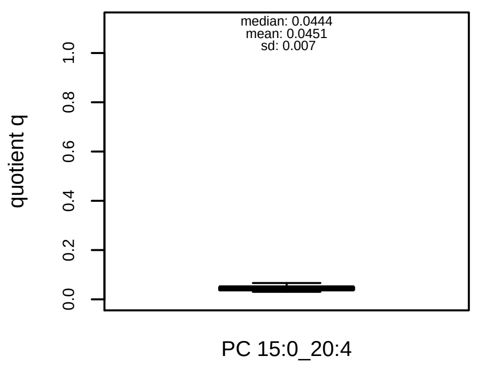


#### Quant. comp., Subject 6

n = 55; comb.: 55

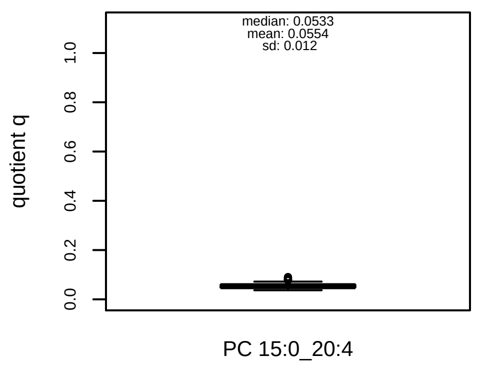


#### Quant. comp., Subject 7

n = 55; comb.: 55

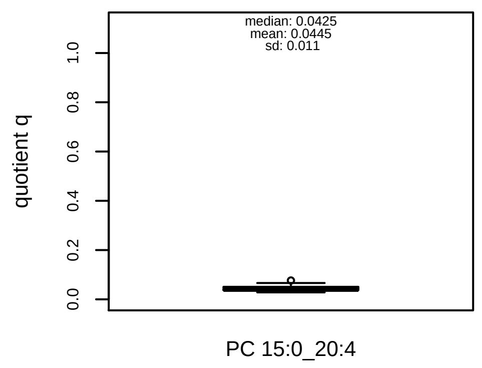


#### Quant. comp., Subject 8

n = 56; comb.: 56

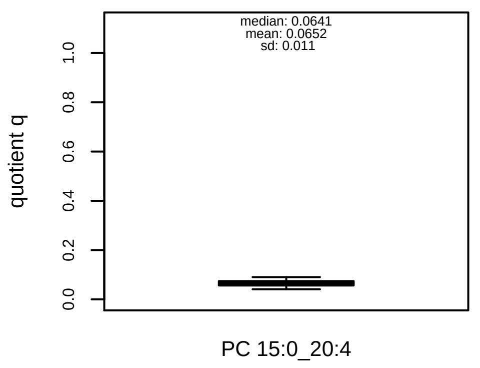


#### Trends of proportions q during challenges

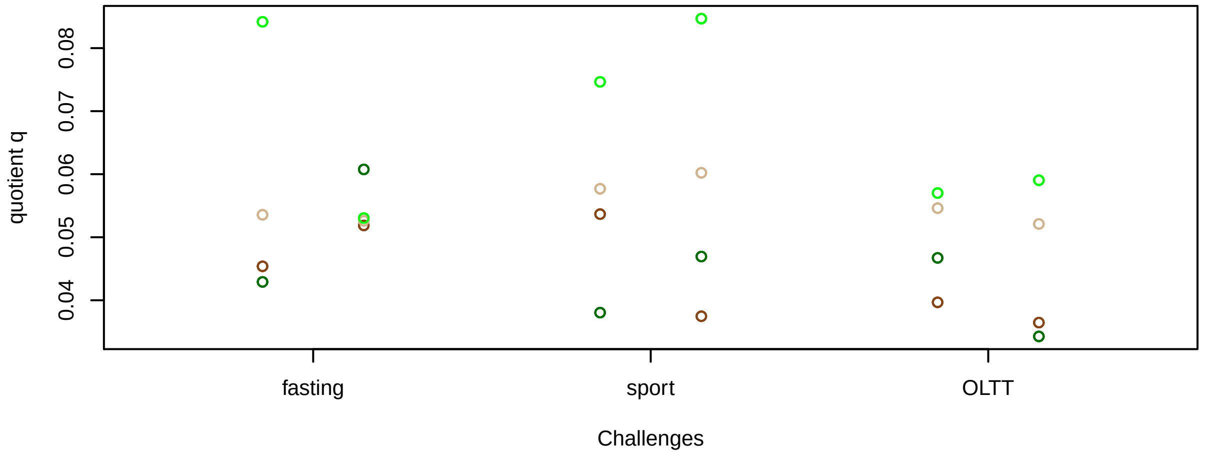


#### Proportions q per time point

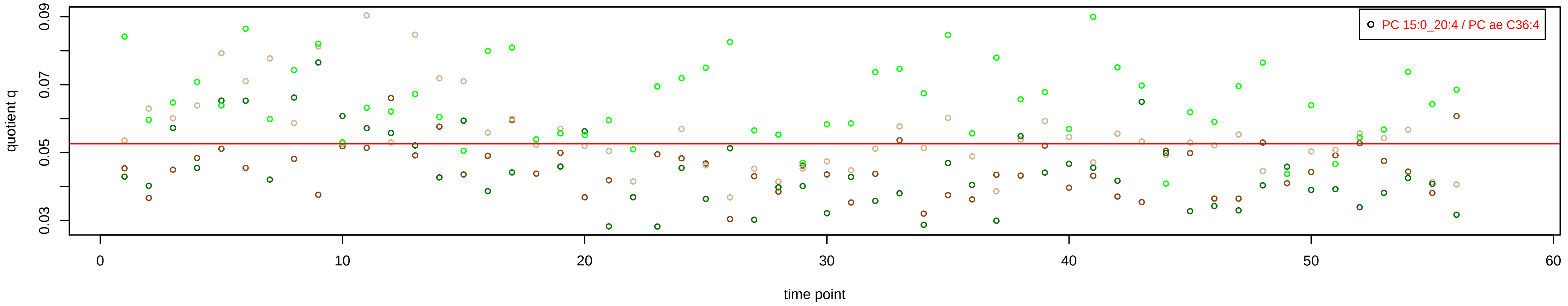

# PC ae C36:5 = PC 15:0_20:5 + R

PC 15:0_20:5 excluded because of missingness > 75%

**No independent Variable with coverage >0.25 out of PC 15:0_20:5**

### Qualitative composition

PC ae C36:5 consists of:
PC 15:0_20:5, PC O−16:0_20:5, PC O−16:1_20:4, PC O−18:
PC O−18:2_18:3, PC 15:1_20:4, PC 17:1_18:4, PC 17:2_18:3
and further compounds

# PC ae C38:0 = PC 18:2_20:5 + R

PC 18:2_20:5 excluded because of missingness > 75%

**No independent Variable with coverage >0.25 out of PC 18:2_20:5**

### Qualitative composition

PC ae C38:0 consists of:
PC 18:2_20:5, PC O−16:0_22:0, PC O−18:0_20:0, PC 15:0_2
PC 16:0_21:0, PC 17:0_20:0, PC 18:0_19:0, PC 16:1_22:6,
PC 18:4_20:3
and further compounds

# PC ae C38:2 = R

**No independent Variable measured.**

### Qualitative composition

PC ae C38:2 consists of:
PC O−16:0_22:2, PC O−18:0_20:2, PC O−18:1_20:1, PC O−
PC O−16:1_22:1, PC 15:0_22:2, PC 15:1_22:1, PC 17:0_20:2
PC 17:1_20:1, PC 17:2_20:0, PC 18:1_19:1, PC 18:2_19:0,
PC 18:4_20:5
and further compounds

# PC ae C38:3 = PC 17:0_20:3 + R

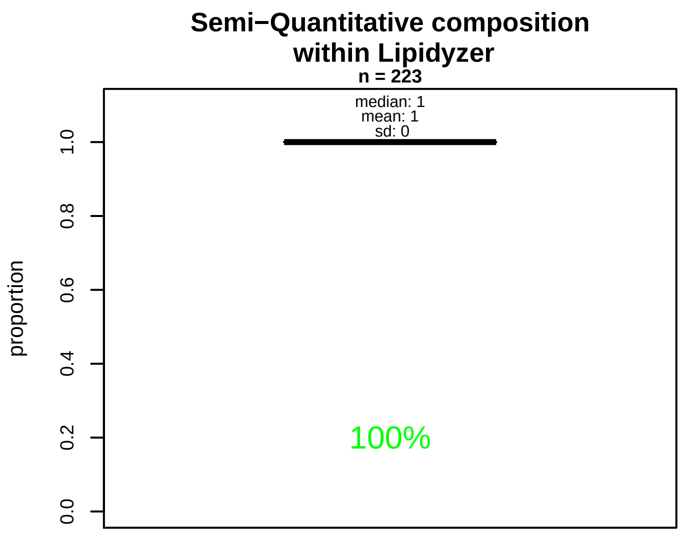


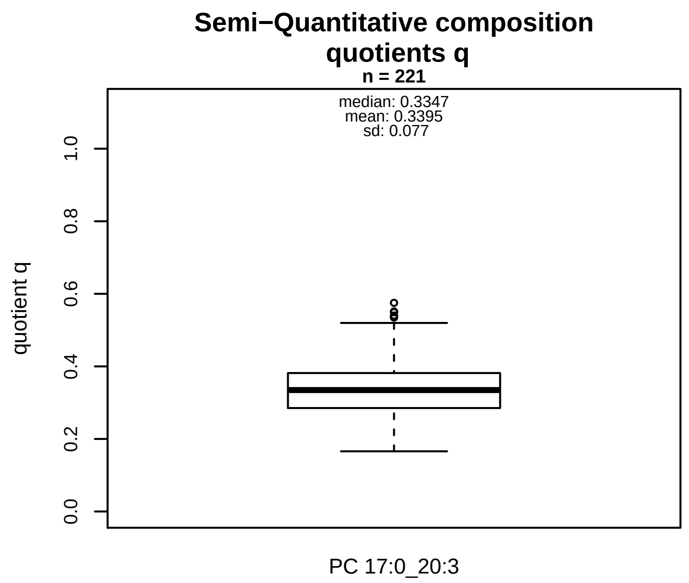


### Semi−Quantitative composition: Lipidyzer

PC ae C38:3 consists of:
PC 17:0_20:3 100%
and of (not quantified):
PC O−18:0_20:3, PC O−16:1_22:2, PC O−18:1_20:2, PC O−
PC O−18:3_20:0, PC 15:1_22:2, PC 17:1_20:2, PC 17:2_20::
PC 18:3_19:0
and further compounds

### Composition: mean of proportions q

conc(PC ae C38:3) * 0.3395 = conc(PC 17:0_20:3) [var(q)=0.3395]
Percentiles: 5%−>0.2248, 25%−>0.2852, 75%−>0.3816, 95%−>0.4767

### Linear model

```
PC ae C38:3 ~ b * ( PC 17:0_20:3 )
```

b = 1.66896

$R^2$ = 0.64485

### Ranges

| Measure | AbsoluteIDQ | sum(Lipidyzer) | delta |
|---|---|---|---|
| Min | 2 | 0.43 | 1.57 |
| Max | 6.15 | 2.75 | 3.4 |
| Mean | 3.68 | 1.27 | 2.41 |
| Median | 3.73 | 1.27 | 2.46 |
| SD | 0.97 | 0.47 | 0.5 |

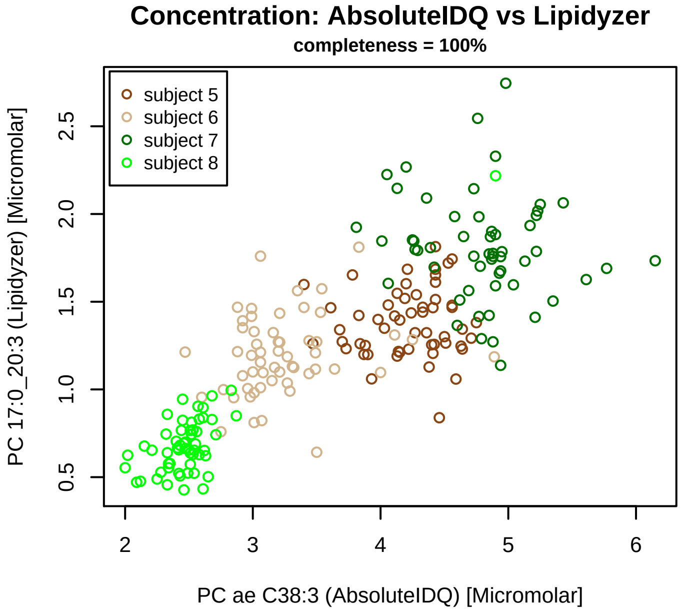


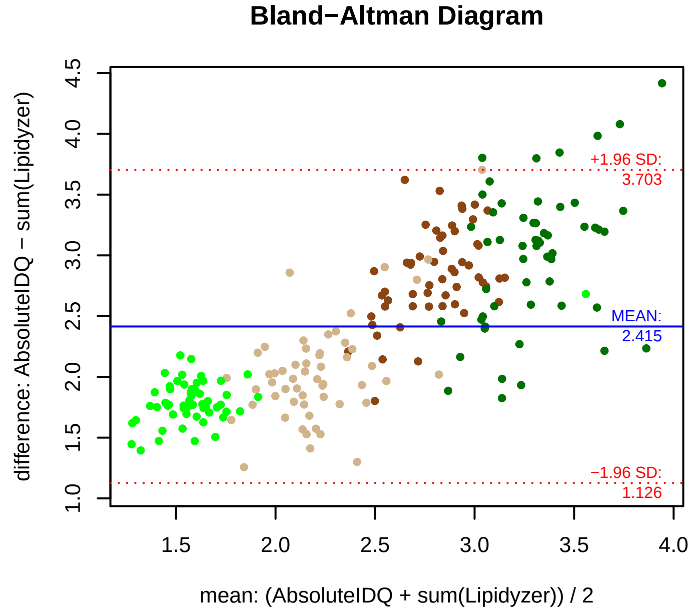


### Stability of composition

**PC 17:0_20:3 / PC ae C38:3**
**Shapiro−Wilk Test of log quotients, pv: 0.30664; OK**
SW: ok; ANOVA: Comp. ~ Subject_ID −> pv = 0.00011; DIFFERENCE
Wilcoxon: Challenge; fasting: 0.875; sport: 0.875; OLTT: 0.125

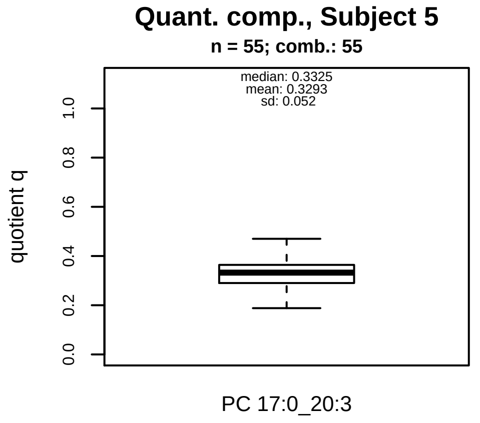


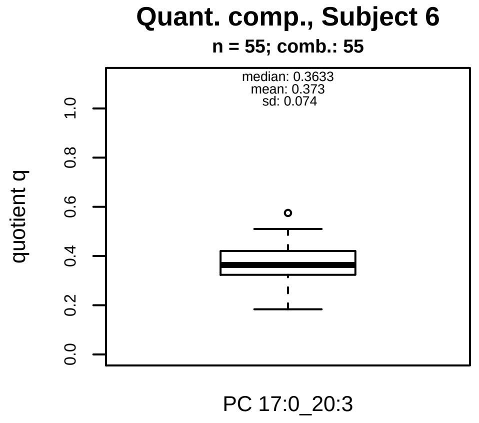


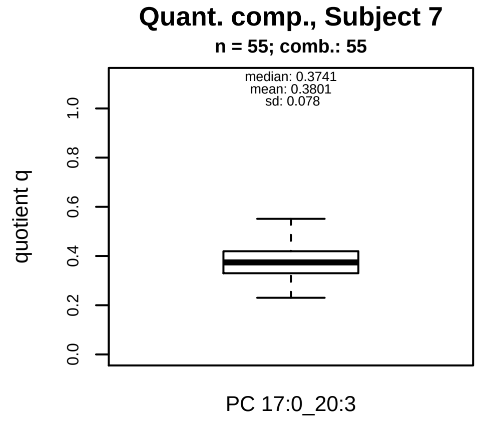


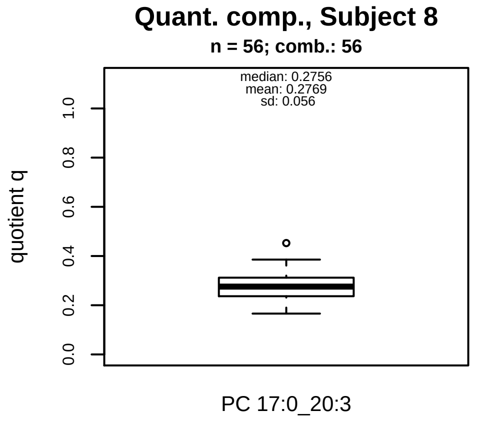


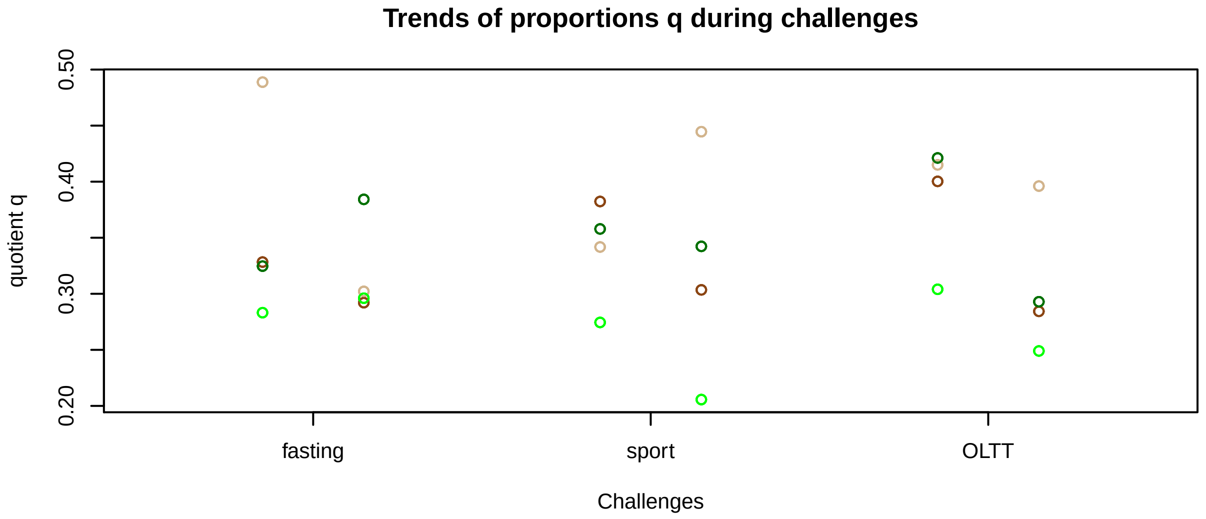


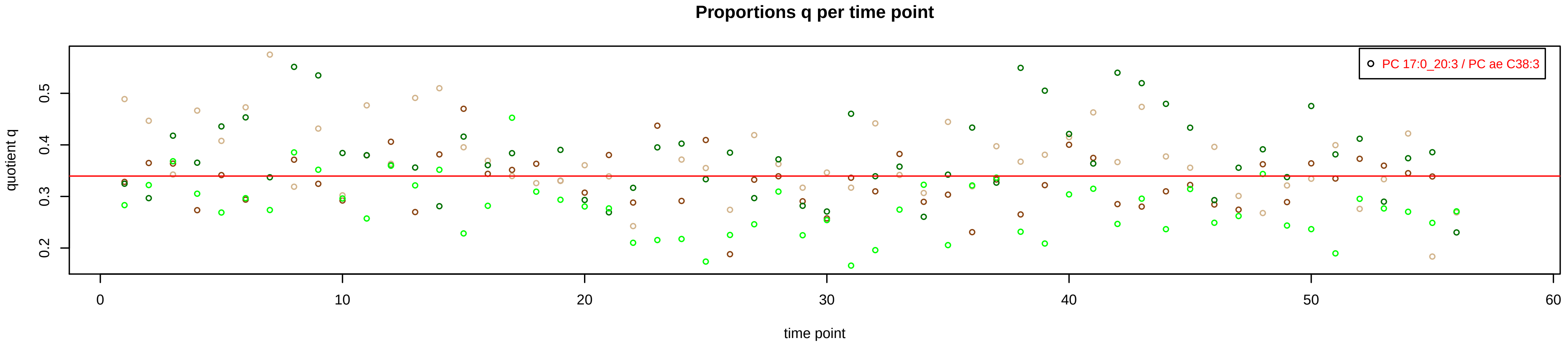

# PC ae C38:4 = PC 17:0_20:4 + R

#### Semi−Quantitative composition within Lipidyzer

n = 223

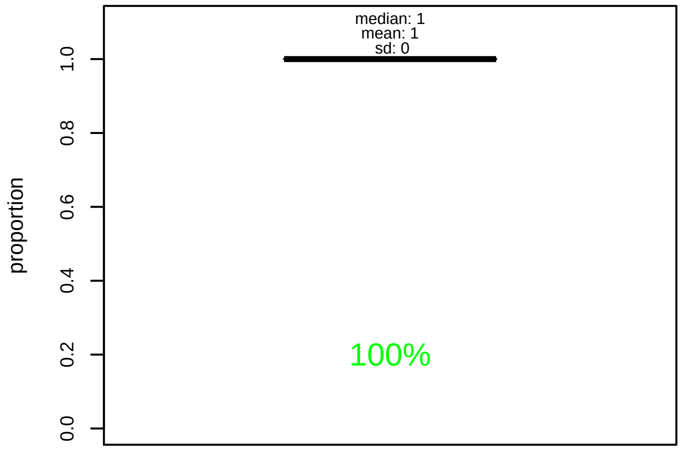


#### Semi−Quantitative composition quotients q

n = 221

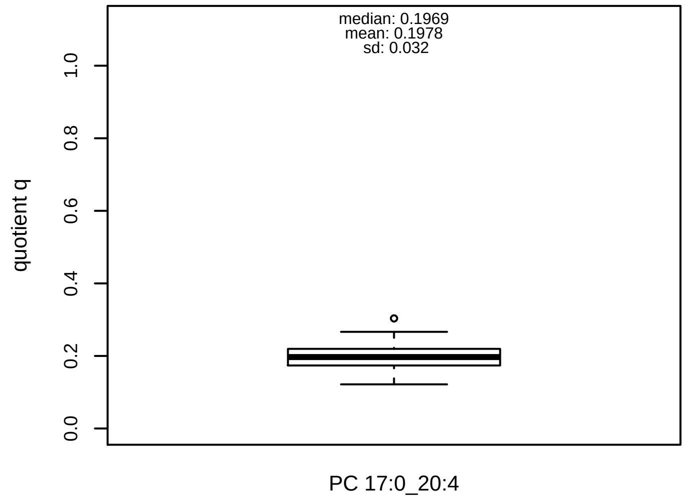


#### Semi−Quantitative composition: Lipidyzer

PC ae C38:4 consists of:
PC 17:0_20:4 100%
and of (not quantified):
PC O−16:0_22:4, PC O−18:0_20:4, PC O−20:0_18:4, PC O−
PC O−18:2_20:2, PC O−20:1_18:3, PC 15:0_22:4, PC 17:1_
PC 18:4_19:0
and further compounds

#### Composition: mean of proportions q

conc(PC ae C38:4) * 0.1978 = conc(PC 17:0_20:4) [var(q)=0.1978]
Percentiles: 5%−>0.1495, 25%−>0.1737, 75%−>0.2195, 95%−>0.2547

#### Linear model

```
PC ae C38:4 ~ b * ( PC 17:0_20:4 )
```

b = 3.91691

$R^2$ = 0.64147

#### Ranges

| Measure | AbsoluteIDQ | sum(Lipidyzer) | delta |
|---|---|---|---|
| Min | 6.25 | 1.05 | 5.2 |
| Max | 21 | 4.38 | 16.62 |
| Mean | 13.93 | 2.73 | 11.2 |
| Median | 14.55 | 2.81 | 11.74 |
| SD | 3.65 | 0.75 | 2.91 |

#### Concentration: AbsoluteIDQ vs Lipidyzer

completeness = 100%

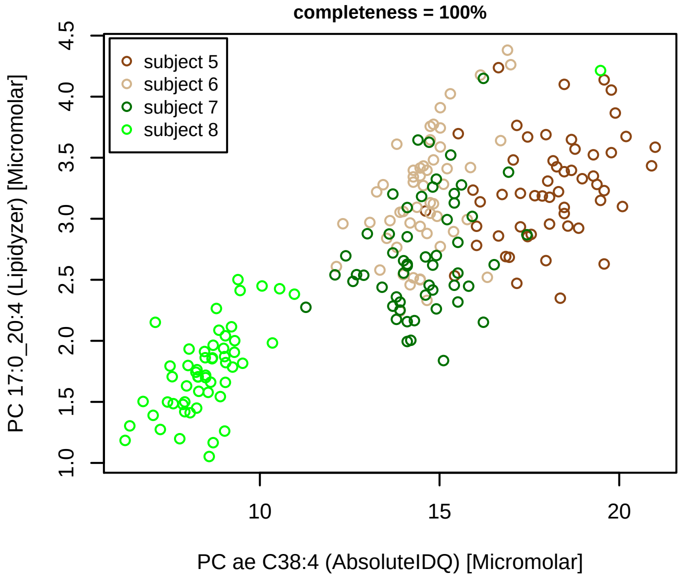


#### Bland−Altman Diagram

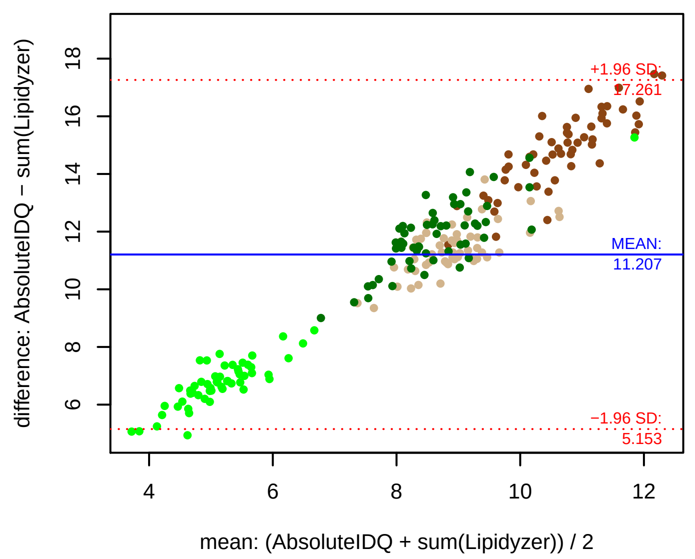


#### Stability of composition

**PC 17:0_20:4 / PC ae C38:4**
**Shapiro−Wilk Test of log quotients, pv: 0.19617; OK**
SW: ok; ANOVA: Comp. ~ Subject_ID −> pv = 0.01837; EQUAL
Wilcoxon: Challenge; fasting: 0.875; sport: 0.375; OLTT: 0.125

#### Quant. comp., Subject 5

n = 55; comb.: 55

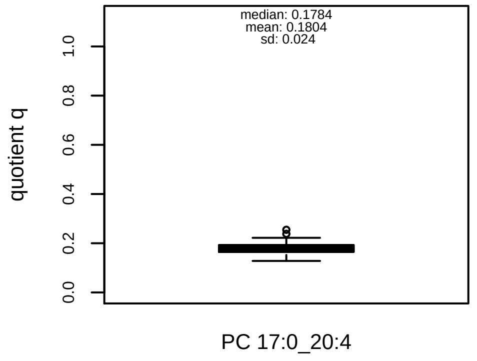


#### Quant. comp., Subject 6

n = 55; comb.: 55

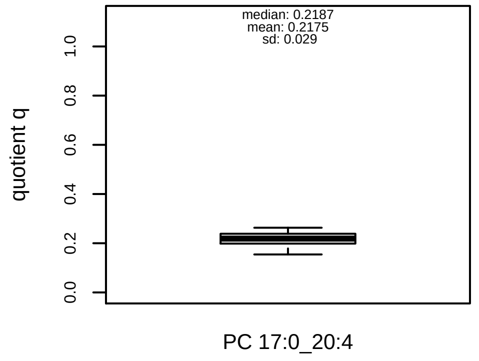


#### Quant. comp., Subject 7

n = 55; comb.: 55

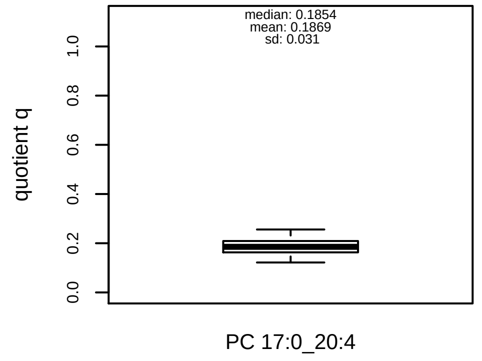


#### Quant. comp., Subject 8

n = 56; comb.: 56

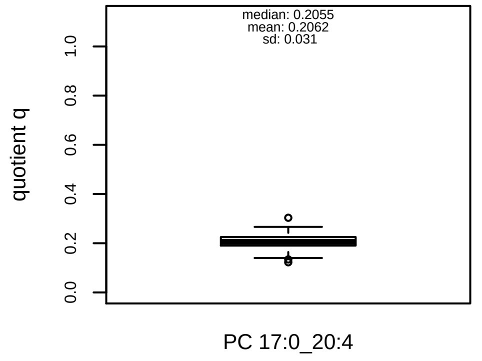


#### Trends of proportions q during challenges

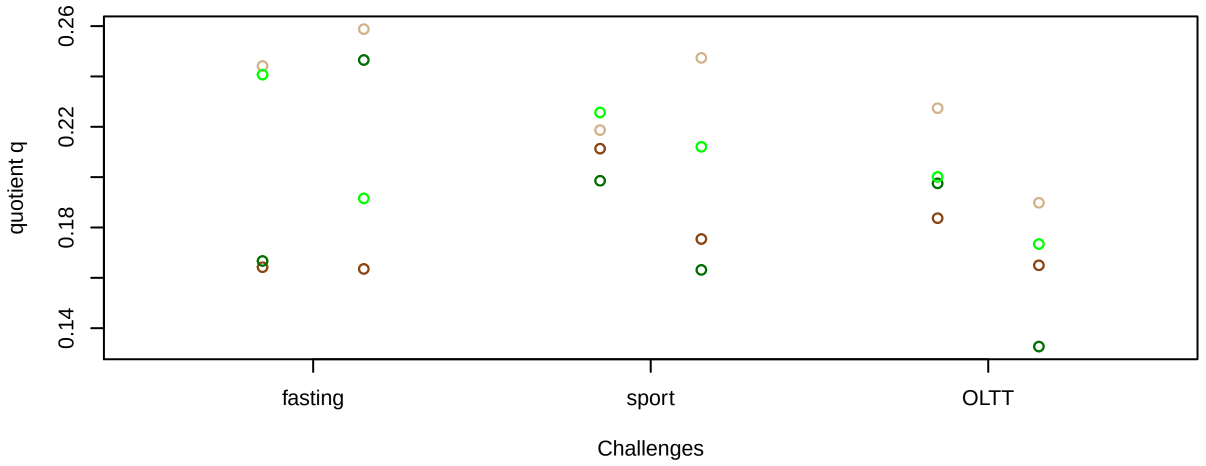


#### Proportions q per time point

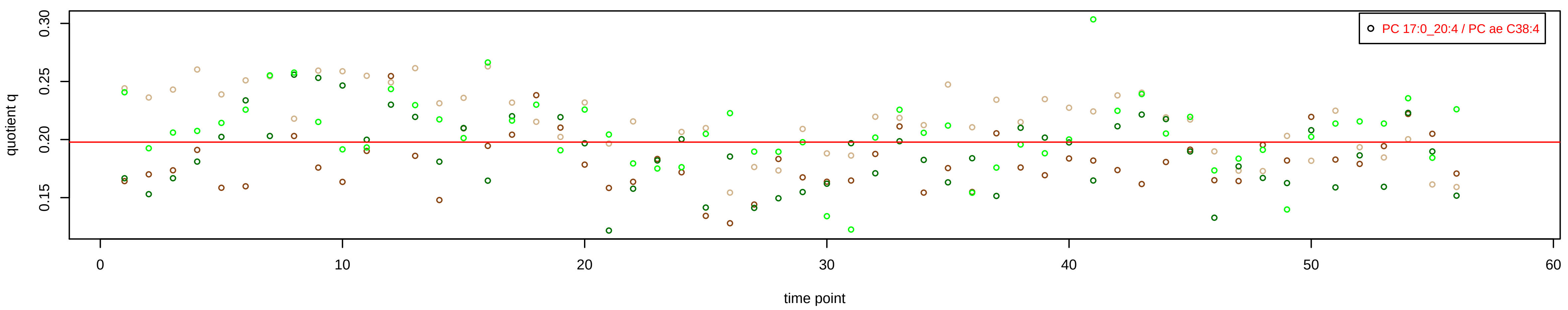

# PC ae C38:5 = PC 15:0_22:5 + PC 17:0_20:5 + R

PC 15:0_22:5 excluded because of missingness > 75%

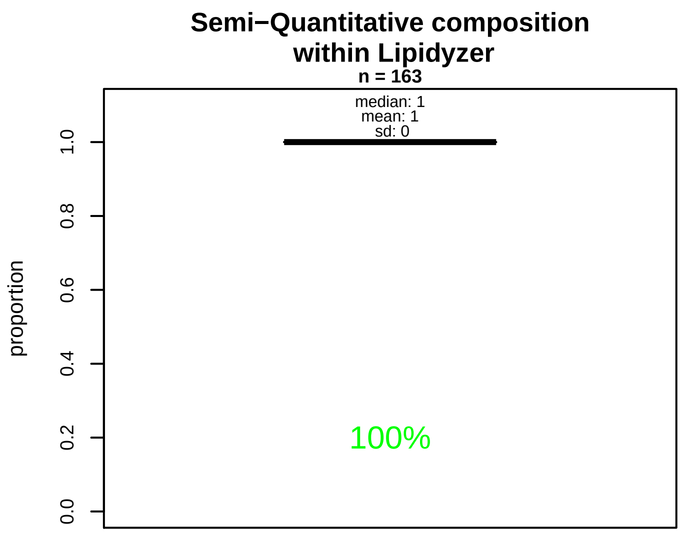


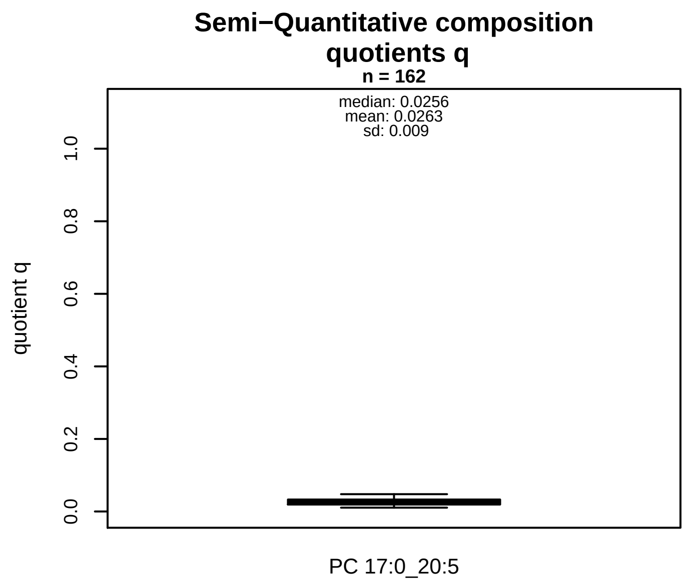


### Semi−Quantitative composition: Lipidyzer

PC ae C38:5 consists of:
  PC 17:0_20:5  100%
and of (not quantified):
  PC 15:0_22:5,    PC O−16:0_22:5,    PC O−16:1_22:4,    PC O−18:
  PC O−18:1_20:4,    PC O−18:2_20:3,    PC O−18:4_20:1,    PC 15:
  PC 18:4_19:1
  and further compounds

### Composition: mean of proportions q

conc(PC ae C38:5) * 0.0263 = conc(PC 17:0_20:5)   [var(q)=0.0263]
Percentiles: 5%−>0.0145, 25%−>0.0191, 75%−>0.0328, 95%−>0.0408

### Linear model

```
PC ae C38:5 ~ b * ( PC 17:0_20:5 )
```

b = 3.44394

$R^2$ = 0.02174

### Ranges

| Measure | AbsoluteIDQ | sum(Lipidyzer) | delta |
|---|---|---|---|
| Min | 12.5 | 0.2 | 12.3 |
| Max | 27.9 | 0.99 | 26.91 |
| Mean | 19.34 | 0.5 | 18.84 |
| Median | 19 | 0.5 | 18.5 |
| SD | 3.67 | 0.15 | 3.52 |

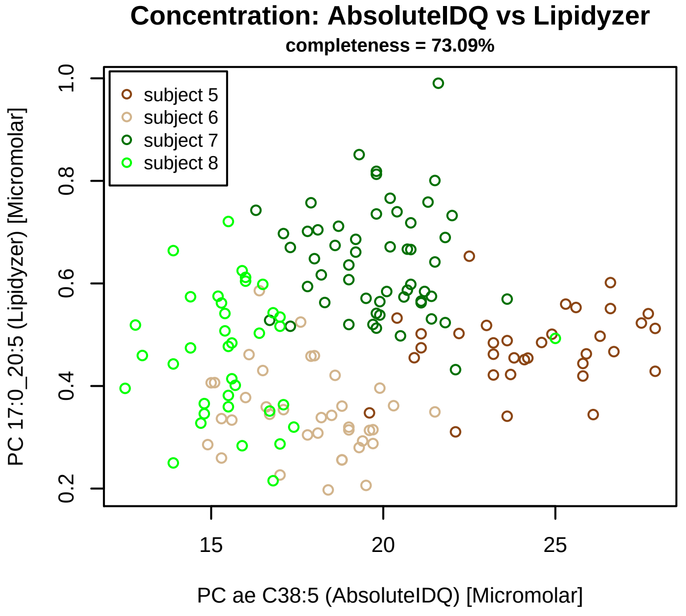


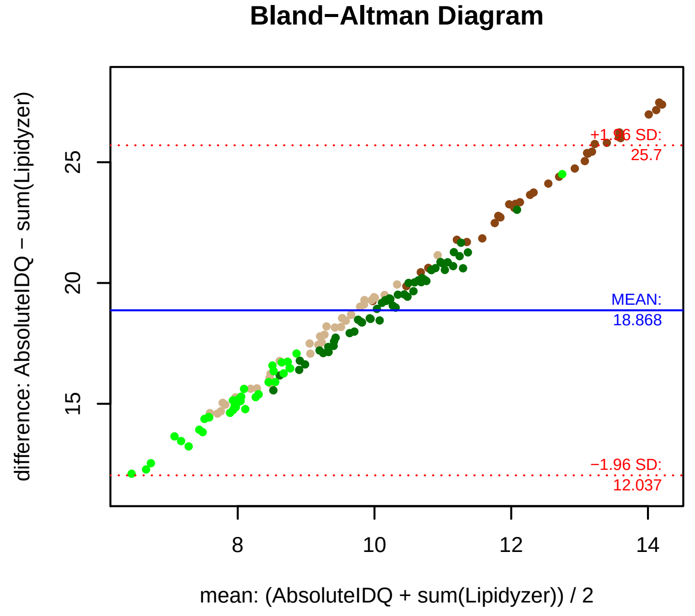


### Stability of composition

**PC 17:0_20:5 / PC ae C38:5**
**Shapiro−Wilk Test of log quotients, pv: 0.02234; OK**
SW: ok; ANOVA: Comp. ~ Subject_ID −> pv = 0; DIFFERENCE
Wilcoxon: Challenge;  fasting: 0.5;  sport: 0.875;  OLTT: 0.125

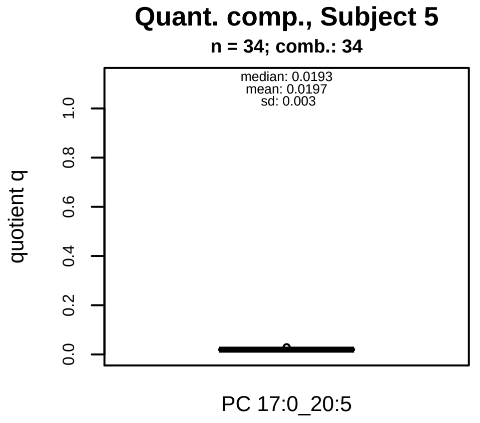


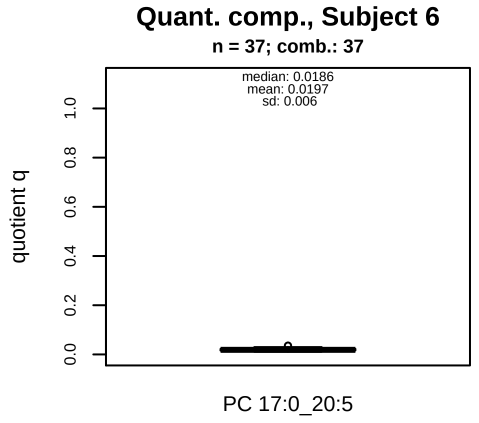


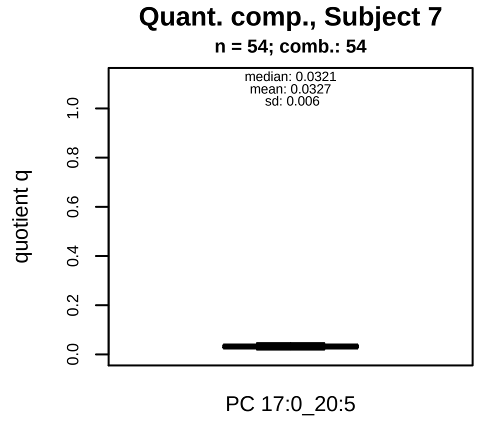


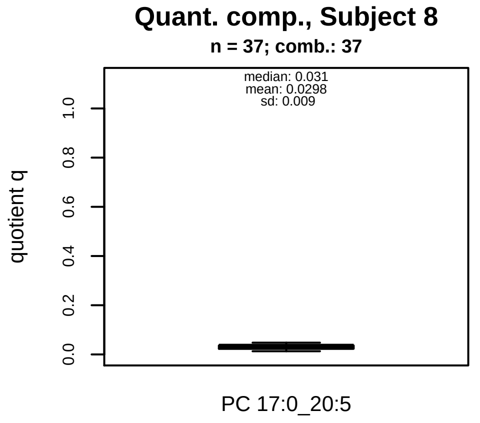


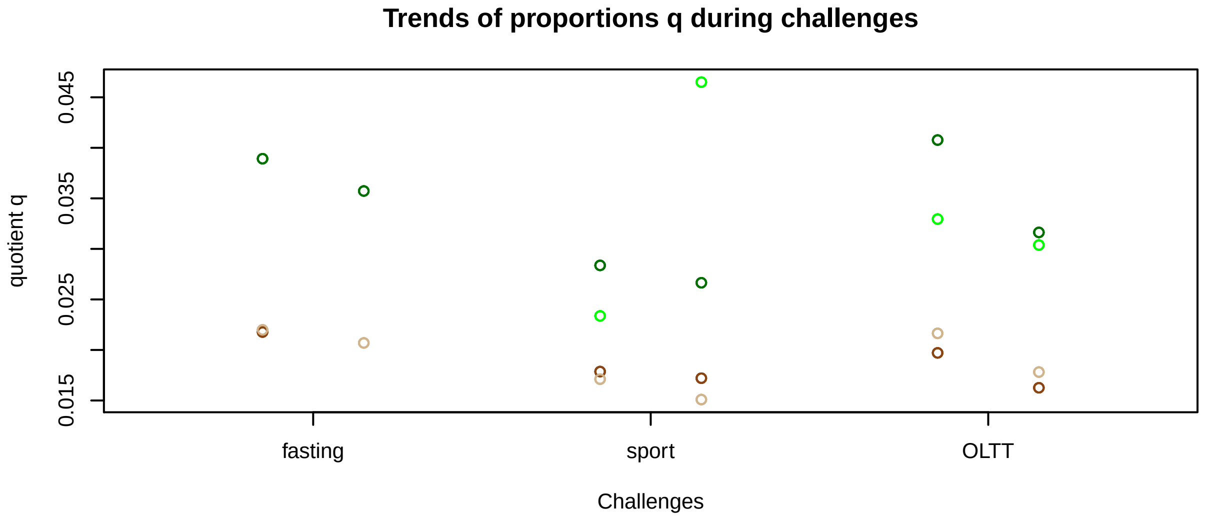


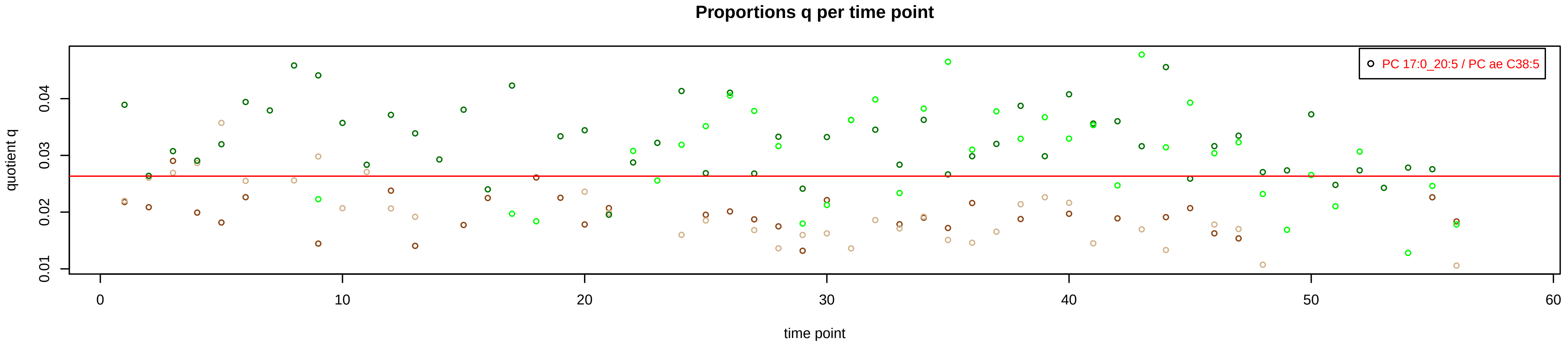

# PC ae C38:6 = PC 15:0_22:6 + R

#### Semi−Quantitative composition within Lipidyzer

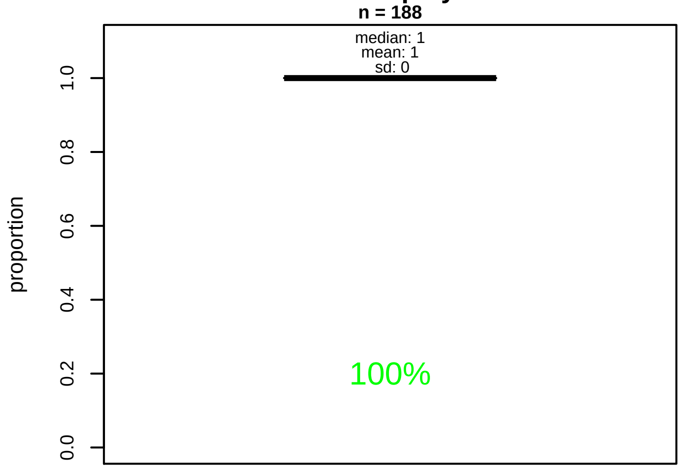


#### Semi−Quantitative composition quotients q

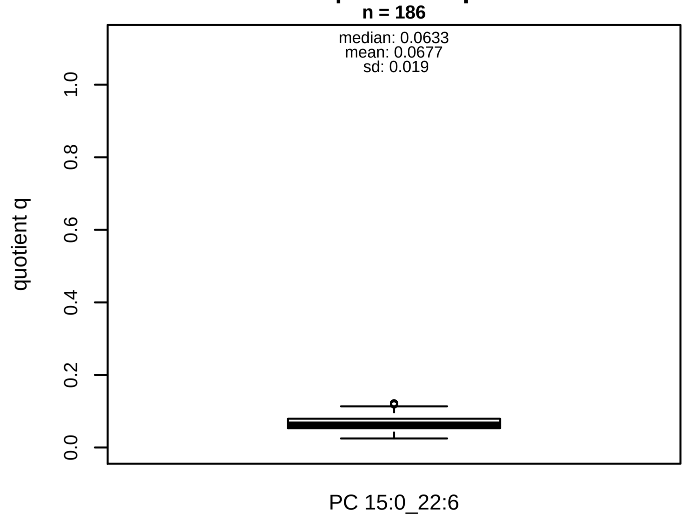


#### Semi−Quantitative composition: Lipidyzer

PC ae C38:6 consists of:
PC 15:0_22:6 100%
and of (not quantified):
PC O−16:0_22:6, PC O−18:1_20:5, PC O−18:2_20:5, PC 17:
PC 17:2_20:4
and further compounds

#### Composition: mean of proportions q

conc(PC ae C38:6) * 0.0677 = conc(PC 15:0_22:6) [var(q)=0.0677]
Percentiles: 5%−>0.0433, 25%−>0.0532, 75%−>0.0786, 95%−>0.1084

#### Linear model

```
PC ae C38:6 ~ b * ( PC 15:0_22:6 )
```

b = 5.55836

$R^2$ = 0.12603

#### Ranges

| Measure | AbsoluteIDQ | sum(Lipidyzer) | delta |
|---|---|---|---|
| Min | 5.29 | 0.17 | 5.12 |
| Max | 12.77 | 0.96 | 11.81 |
| Mean | 7.85 | 0.51 | 7.34 |
| Median | 7.32 | 0.51 | 6.81 |
| SD | 1.87 | 0.12 | 1.75 |

#### Concentration: AbsoluteIDQ vs Lipidyzer

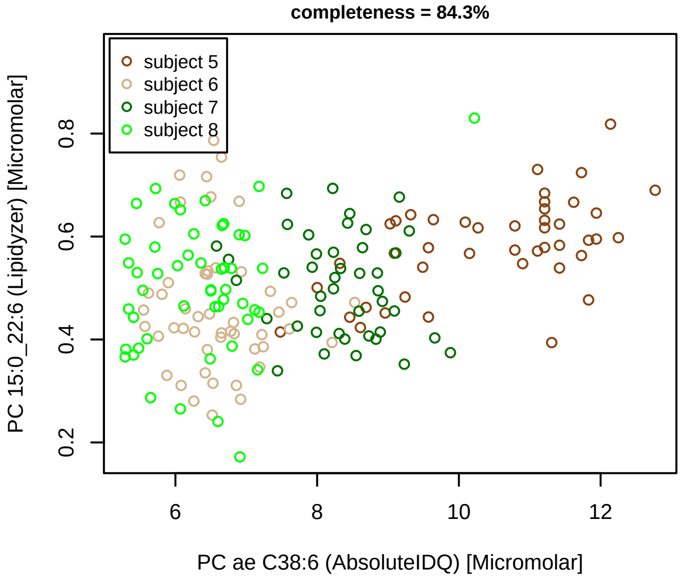


#### Bland−Altman Diagram

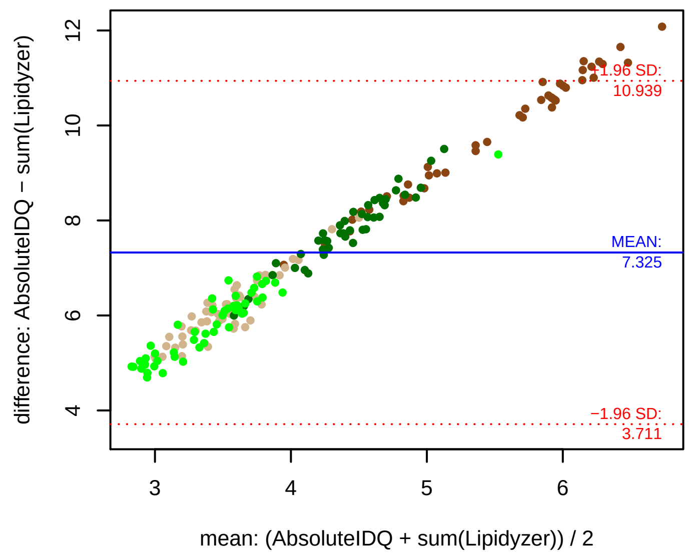


#### Stability of composition

**PC 15:0_22:6 / PC ae C38:6**
**Shapiro−Wilk Test of log quotients, pv: 0.22571; OK**
SW: no; Kruskal: Comp. ~ Subject_ID −> pv = 0; DIFFERENCE
Wilcoxon: Challenge; fasting: 0.75; sport: 0.625; OLTT: 0.5

#### Quant. comp., Subject 5

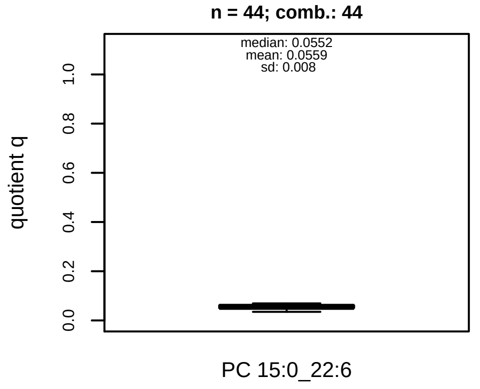


#### Quant. comp., Subject 6

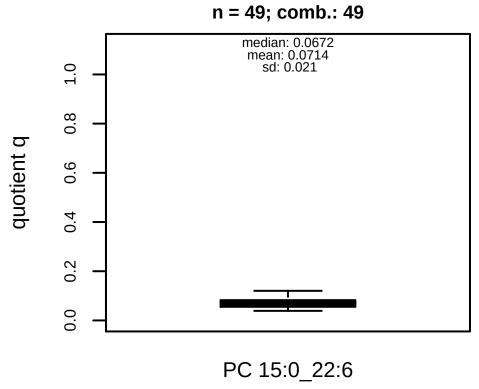


#### Quant. comp., Subject 7

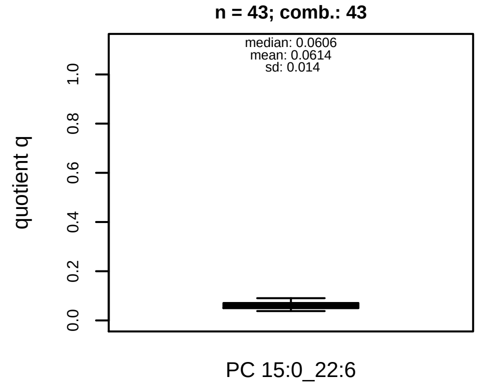


#### Quant. comp., Subject 8

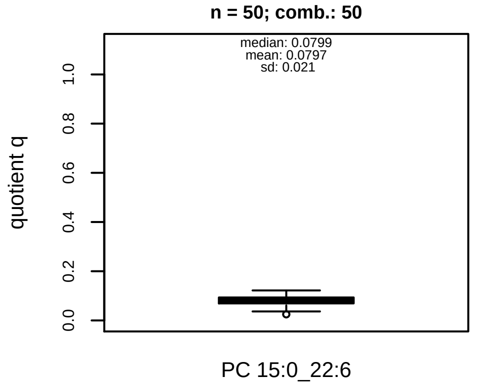


#### Trends of proportions q during challenges

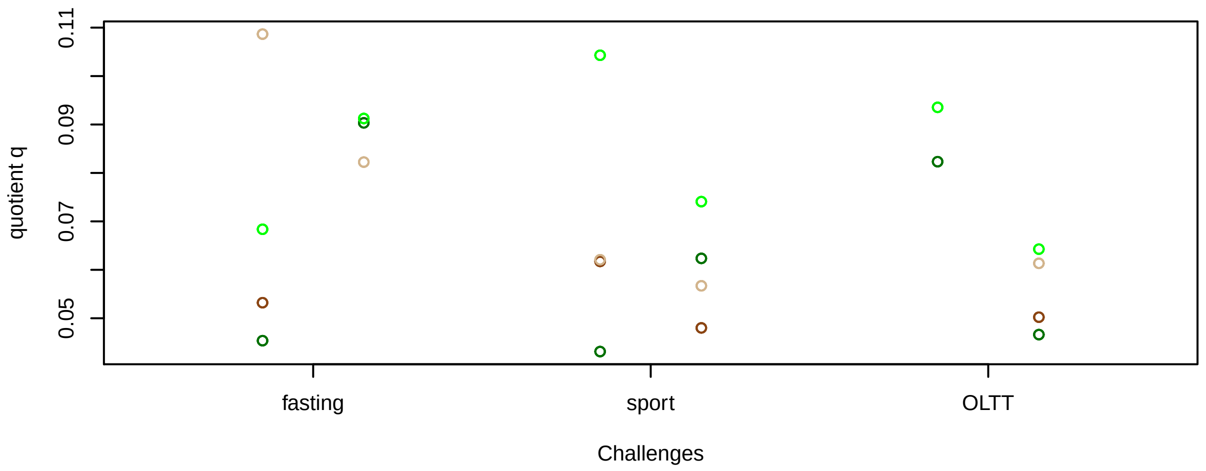


#### Proportions q per time point

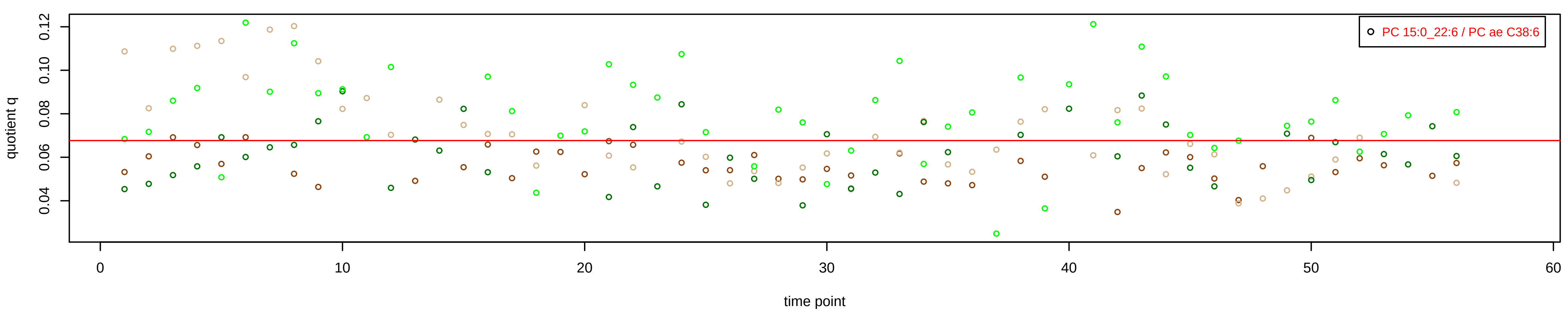

# PC ae C40:1 = PC 18:2_22:6 + R

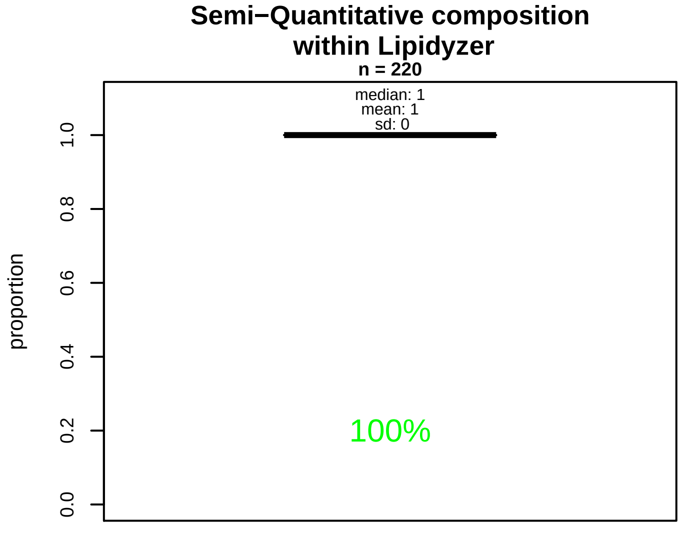


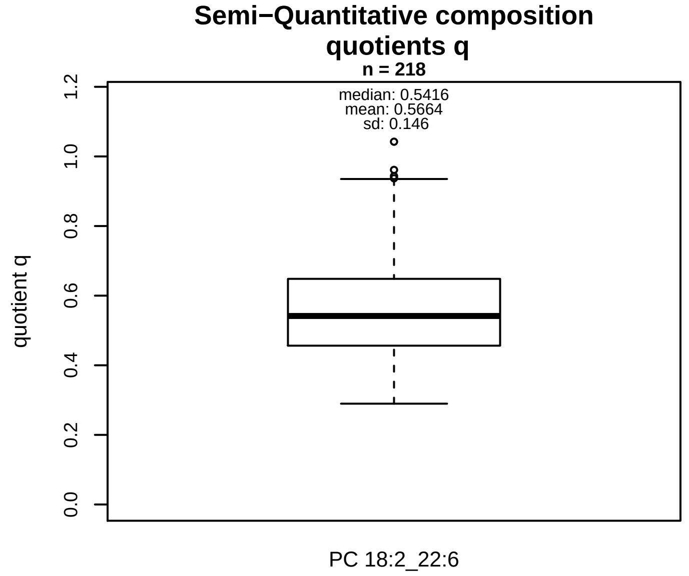


### Semi−Quantitative composition: Lipidyzer

PC ae C40:1 consists of:
PC 18:2_22:6 100%
and of (not quantified):
PC O−18:0_22:1, PC O−18:1_22:0, PC O−20:0_20:1, PC 17:
PC 17:1_22:0, PC 18:1_21:0, PC 19:0_20:1, PC 19:1_20:0,
PC 20:4_20:4
and further compounds

### Composition: mean of proportions q

conc(PC ae C40:1) * 0.5664 = conc(PC 18:2_22:6) [var(q)=0.5664]
Percentiles: 5%−>0.3782, 25%−>0.4566, 75%−>0.648, 95%−>0.8648

### Linear model

```
PC ae C40:1 ~ b * ( PC 18:2_22:6 )
```

b = 0.44624

$R^2$ = 0.20278

### Ranges

| Measure | AbsoluteIDQ | sum(Lipidyzer) | delta |
|---|---|---|---|
| Min | 1.27 | 0.5 | 0.77 |
| Max | 2.64 | 2.11 | 0.53 |
| Mean | 1.82 | 1.03 | 0.79 |
| Median | 1.8 | 0.96 | 0.84 |
| SD | 0.31 | 0.31 | 0 |

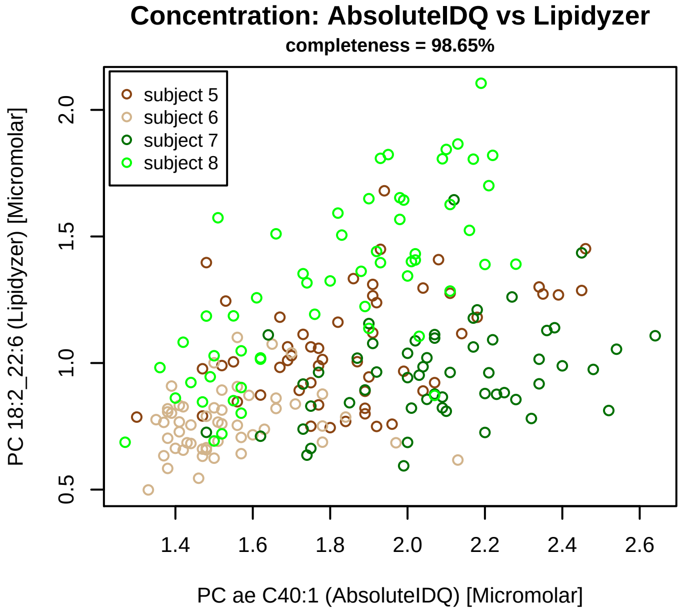


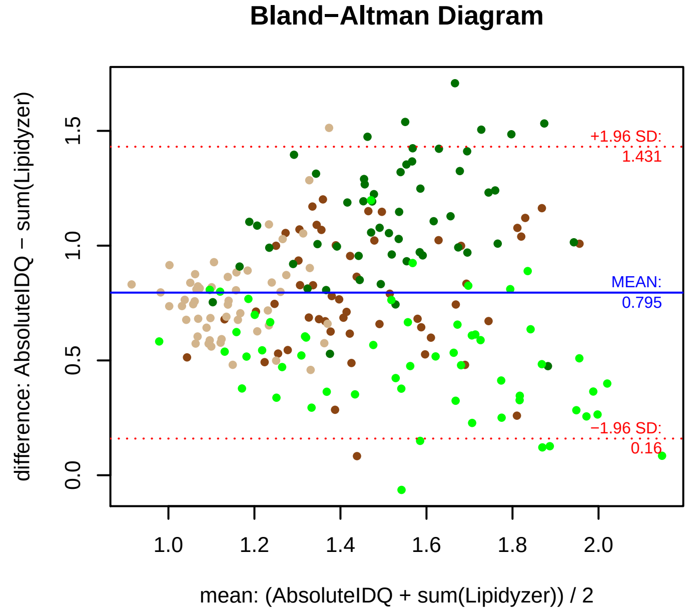


### Stability of composition

**PC 18:2_22:6 / PC ae C40:1**
**Shapiro−Wilk Test of log quotients, pv: 0.32761; OK**
SW: ok; ANOVA: Comp. ~ Subject_ID −> pv = 6e−05; DIFFERENCE
Wilcoxon: Challenge; fasting: 1; sport: 1; OLTT: 0.875

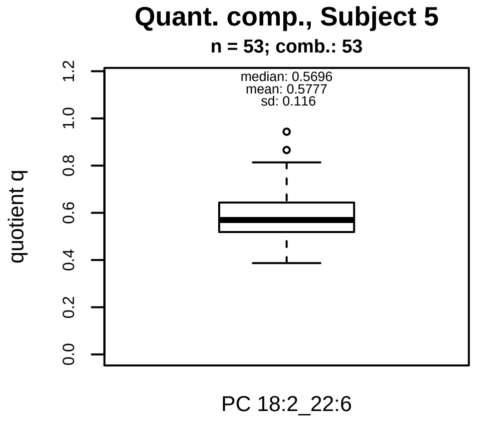


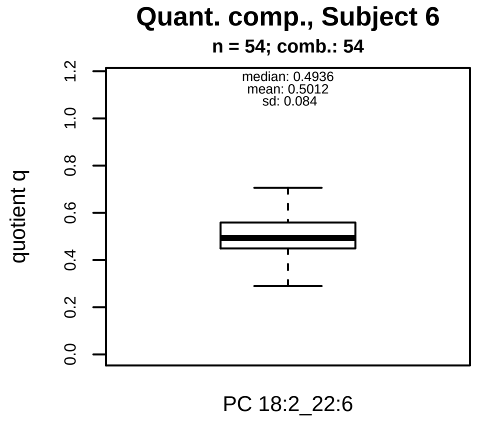


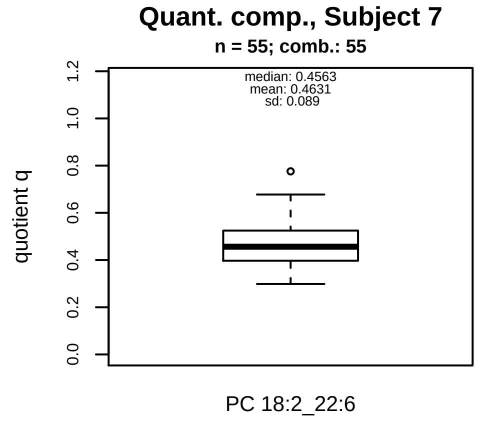


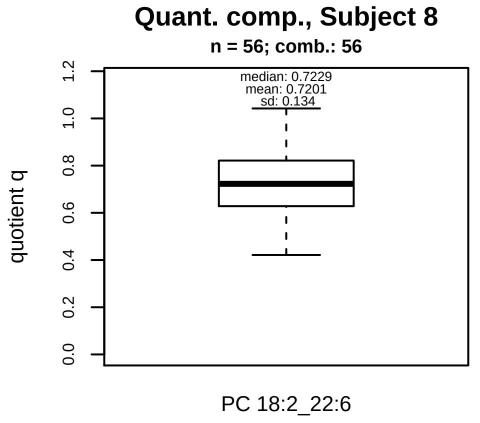


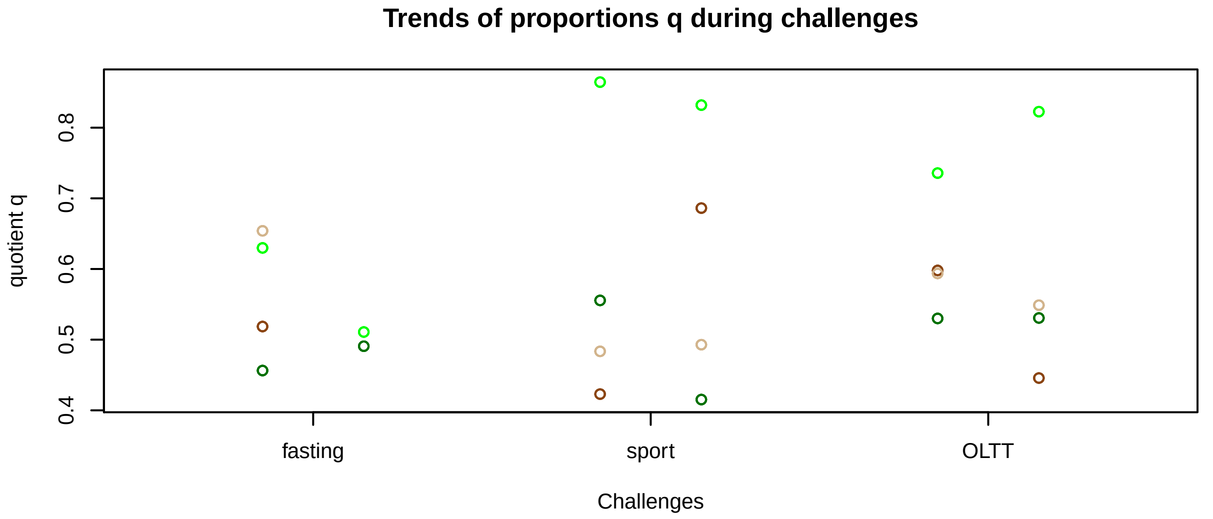


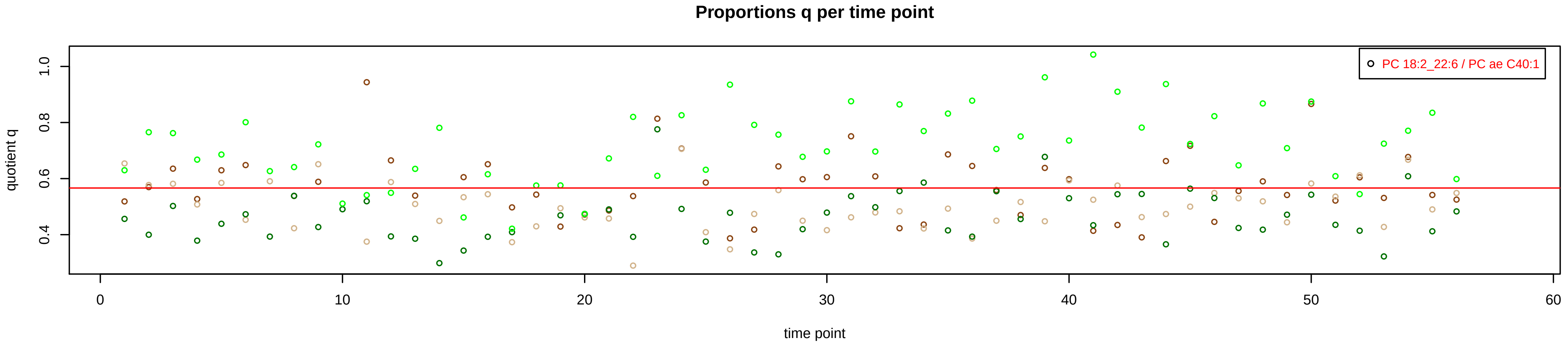

# PC ae C40:2 = R

**No independent Variable measured.**

### Qualitative composition

PC ae C40:2 consists of:
PC O−18:0_22:2, PC O−18:1_22:1, PC O−18:2_22:0, PC O−[illegible]
PC O−20:1_20:1, PC 17:0_22:2, PC 17:1_22:1, PC 17:2_22:[illegible]
PC 18:2_21:0, PC 19:0_20:2, PC 19:1_20:1, PC 18:3_22:6,
[13C1]SM 43:2
and further compounds

# PC ae C40:3 = R

No independent Variable measured.

### Qualitative composition

PC ae C40:3 consists of:
PC O−18:1_22:2, PC O−18:2_22:1, PC O−20:0_20:3, PC O−[illegible]
PC O−22:0_18:3, PC 17:1_22:2, PC 17:2_22:1, PC 18:3_21:[illegible]
PC 19:0_20:3, PC 19:1_20:2, PC 18:4_22:6, PC 20:5_20:5,
and further compounds

# PC ae C40:4 = R

**No independent Variable measured.**

### Qualitative composition

PC ae C40:4 consists of:
PC O−18:0_22:4, PC O−18:2_22:2, PC O−20:0_20:4, PC O−
PC 17:0_22:4, PC 17:2_22:2, PC 18:4_21:0, PC 19:0_20:4,
PC 19:1_20:3
and further compounds

# PC ae C40:5 = PC 17:0_22:5 + R

#### Semi−Quantitative composition within Lipidyzer

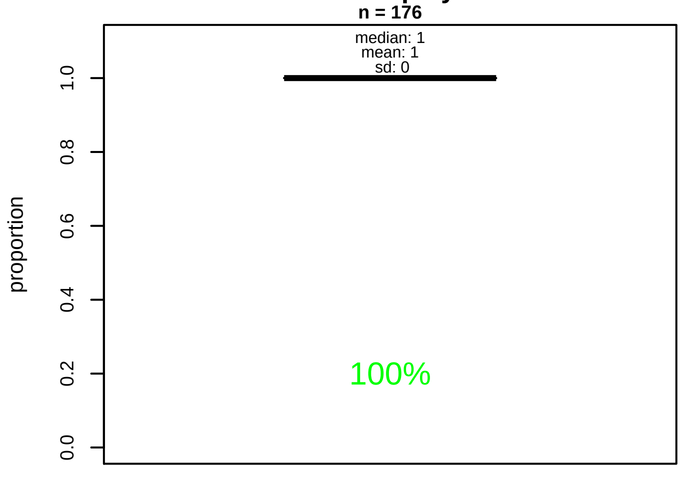


#### Semi−Quantitative composition quotients q

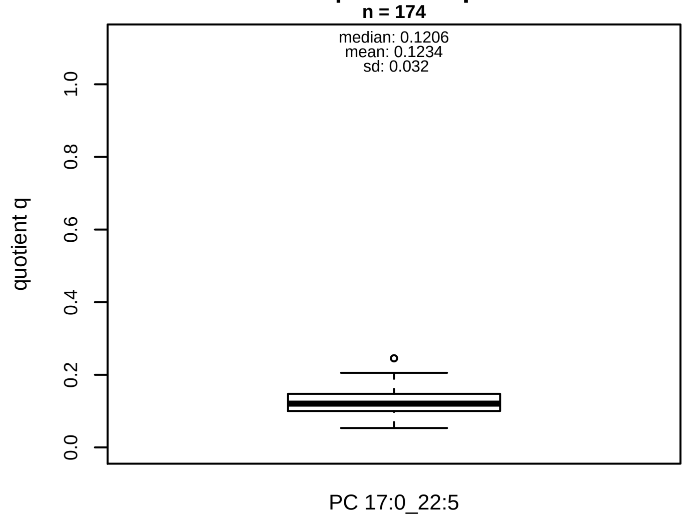


#### Semi−Quantitative composition: Lipidyzer

PC ae C40:5 consists of:
PC 17:0_22:5 100%
and of (not quantified):
PC O−18:0_22:5, PC O−18:1_22:4, PC O−20:0_20:5, PC O−
PC 16:0_23:5, PC 17:1_22:4, PC 19:0_20:5, PC 19:1_20:4,
and further compounds

#### Composition: mean of proportions q

conc(PC ae C40:5) * 0.1234 = conc(PC 17:0_22:5) [var(q)=0.1234]
Percentiles: 5%−>0.0708, 25%−>0.1003, 75%−>0.1471, 95%−>0.1735

#### Linear model

```
PC ae C40:5 ~ b * ( PC 17:0_22:5 )
```

b = 2.47007

$R^2$ = 0.19655

#### Ranges

| Measure | AbsoluteIDQ | sum(Lipidyzer) | delta |
|---|---|---|---|
| Min | 2.22 | 0.15 | 2.07 |
| Max | 5.64 | 0.85 | 4.79 |
| Mean | 3.66 | 0.45 | 3.22 |
| Median | 3.53 | 0.44 | 3.09 |
| SD | 0.73 | 0.13 | 0.6 |

#### Concentration: AbsoluteIDQ vs Lipidyzer

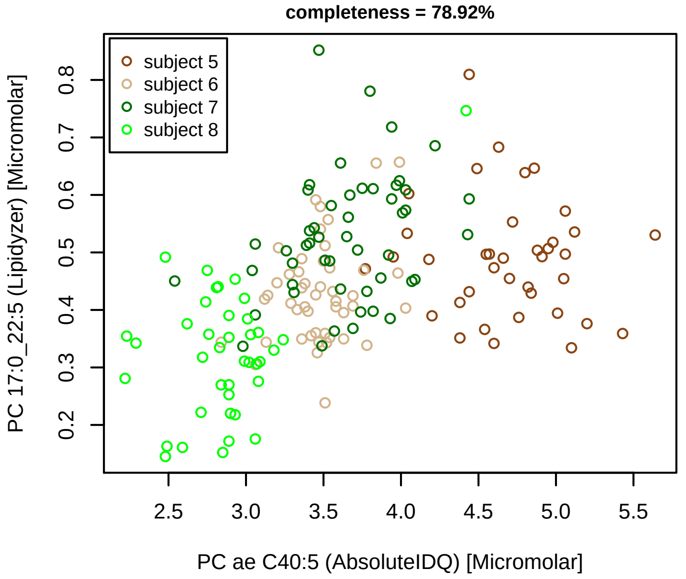


#### Bland−Altman Diagram

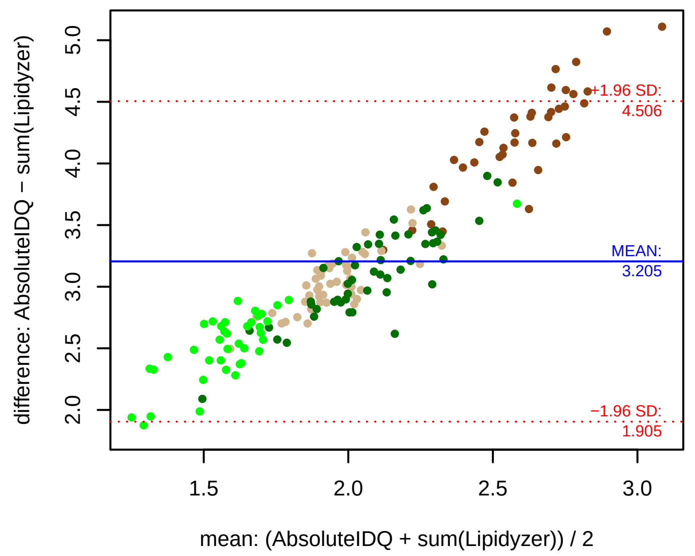


#### Stability of composition

**PC 17:0_22:5 / PC ae C40:5**
**Shapiro−Wilk Test of log quotients, pv: 0.00815; NO**
SW: ok; ANOVA: Comp. ~ Subject_ID −> pv = 0.09316; EQUAL
Wilcoxon: Challenge; fasting: 1; sport: 0.375; OLTT: 0.5

#### Quant. comp., Subject 5

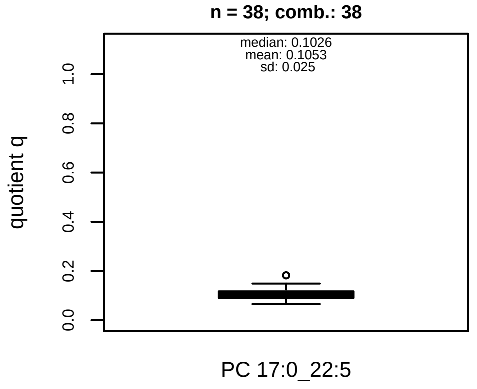


#### Quant. comp., Subject 6

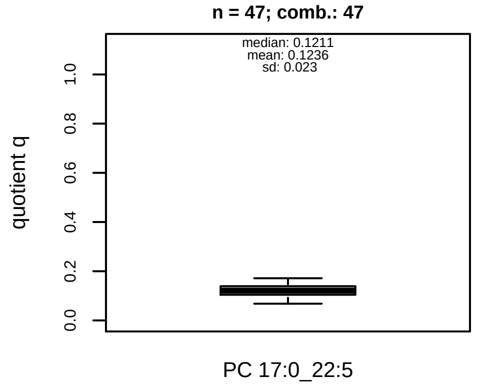


#### Quant. comp., Subject 7

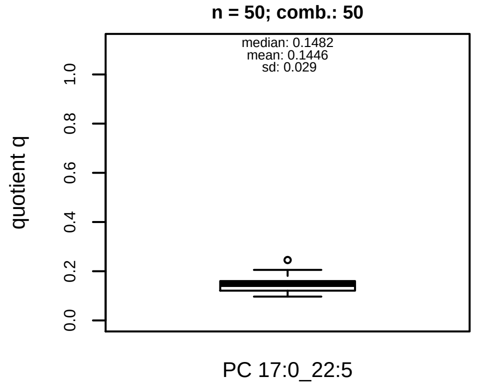


#### Quant. comp., Subject 8

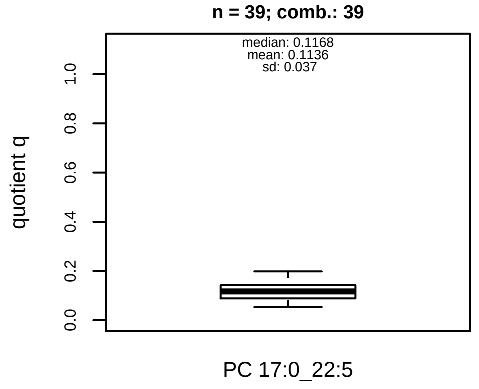


#### Trends of proportions q during challenges

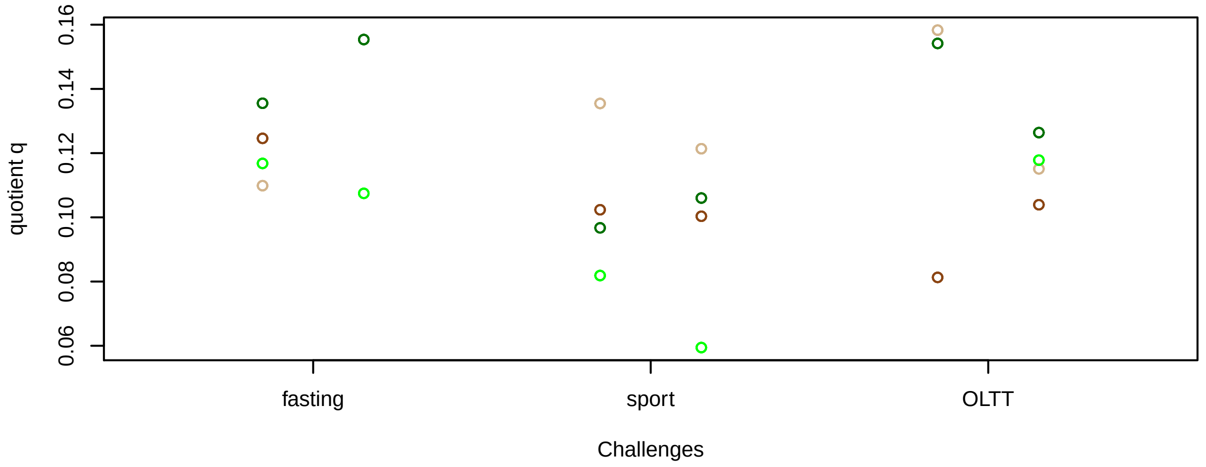


#### Proportions q per time point

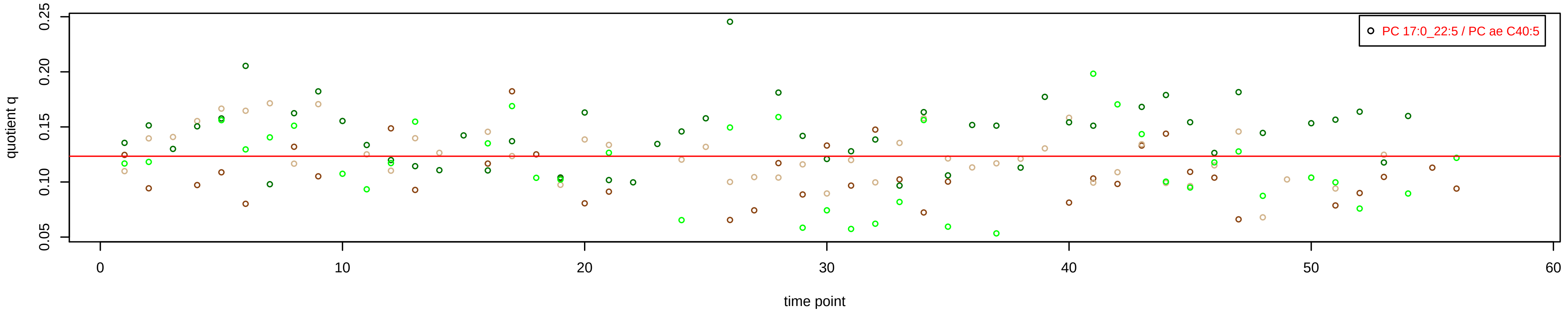

# PC ae C40:6 = PC 17:0_22:6 + R

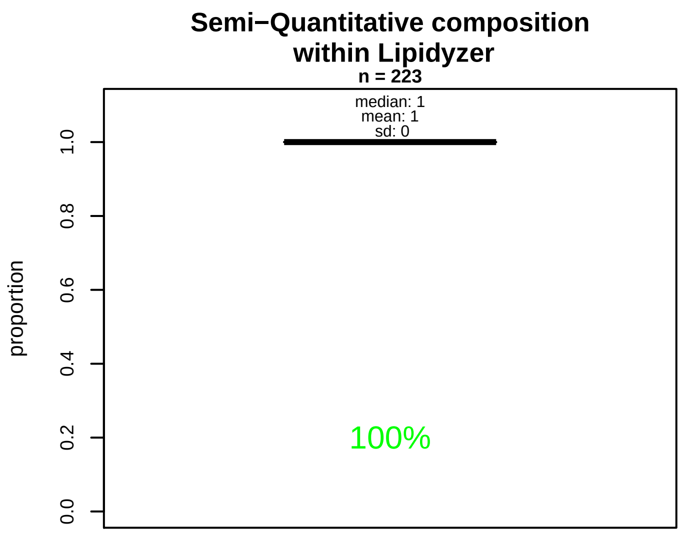


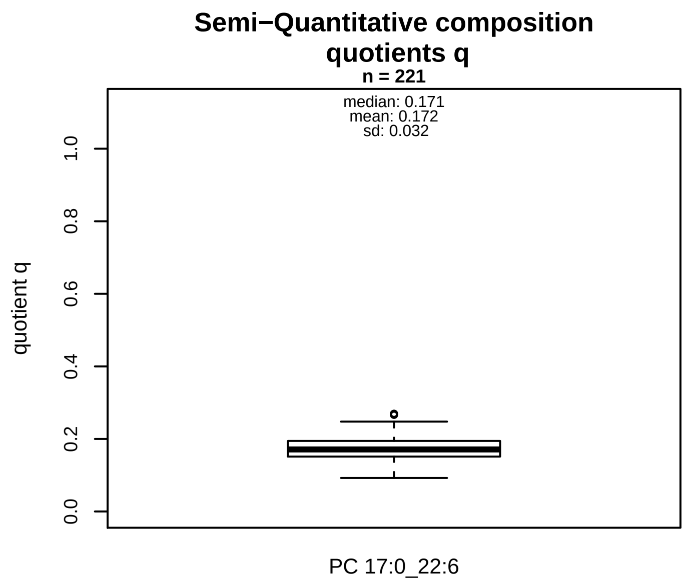


### Semi−Quantitative composition: Lipidyzer

PC ae C40:6 consists of:
PC 17:0_22:6 100%
and of (not quantified):
PC O−18:0_22:6, PC O−18:2_22:4, PC O−20:1_20:5, PC 17:
PC 19:1_20:5
and further compounds

### Composition: mean of proportions q

conc(PC ae C40:6) * 0.172 = conc(PC 17:0_22:6) [var(q)=0.172]
Percentiles: 5%−>0.1216, 25%−>0.1513, 75%−>0.1945, 95%−>0.2262

### Linear model

```
PC ae C40:6 ~ b * ( PC 17:0_22:6 )
```

b = 3.28445

$R^2$ = 0.56301

### Ranges

| Measure | AbsoluteIDQ | sum(Lipidyzer) | delta |
|---|---|---|---|
| Min | 3.54 | 0.43 | 3.11 |
| Max | 8.53 | 1.63 | 6.9 |
| Mean | 5.34 | 0.92 | 4.42 |
| Median | 5.01 | 0.87 | 4.14 |
| SD | 1.12 | 0.26 | 0.86 |

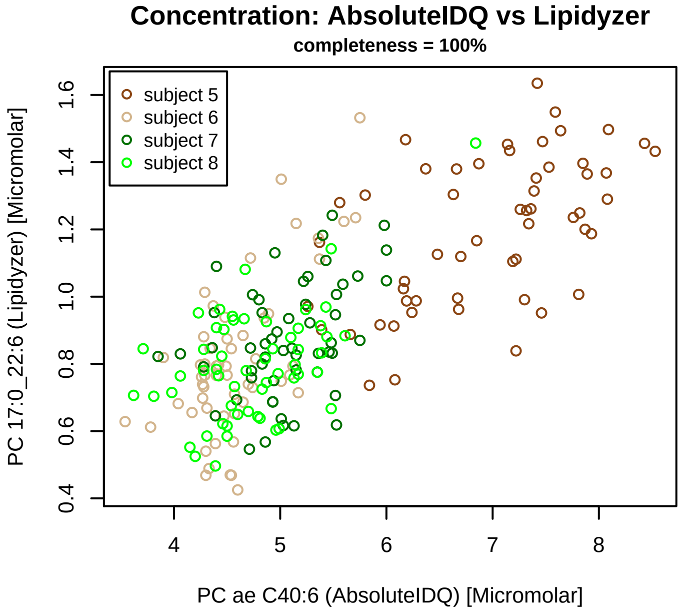


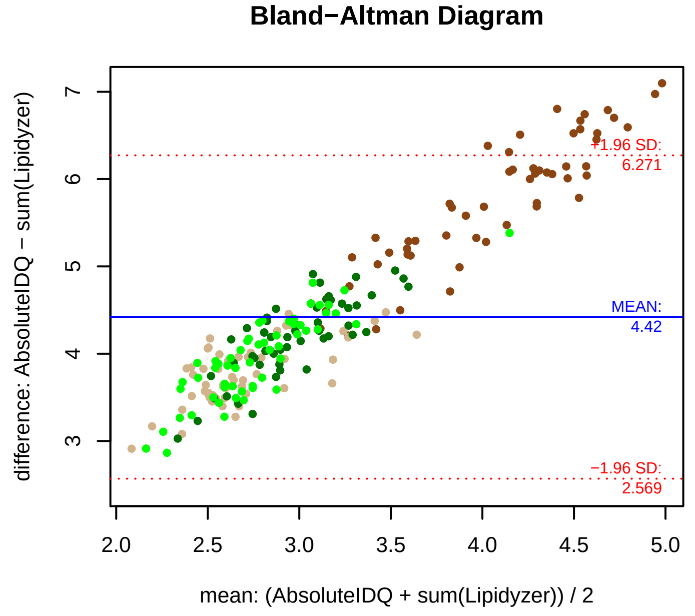


### Stability of composition

**PC 17:0_22:6 / PC ae C40:6**
**Shapiro−Wilk Test of log quotients, pv: 0.15729; OK**
SW: ok; ANOVA: Comp. ~ Subject_ID −> pv = 0.41419; EQUAL
Wilcoxon: Challenge; fasting: 0.875; sport: 0.625; OLTT: 0.25

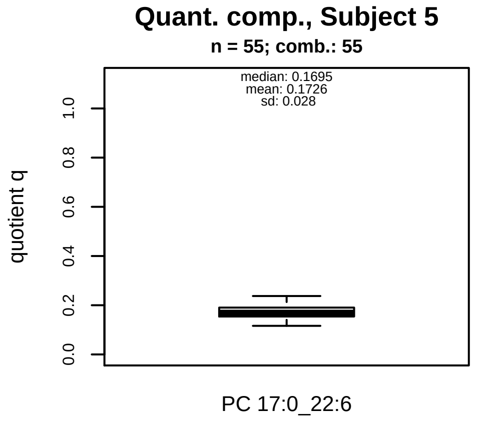


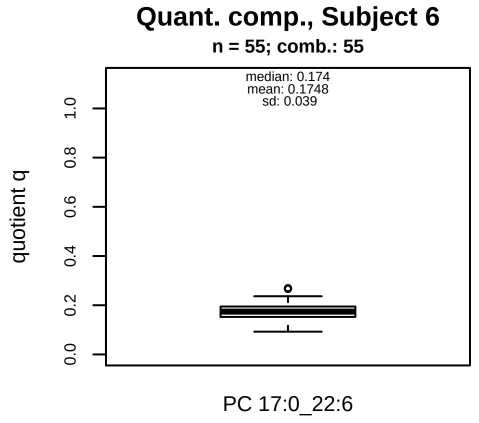


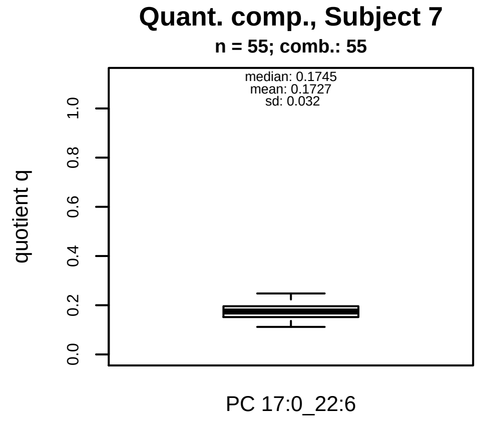


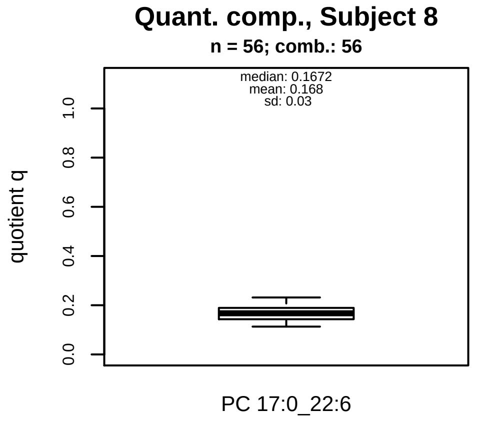


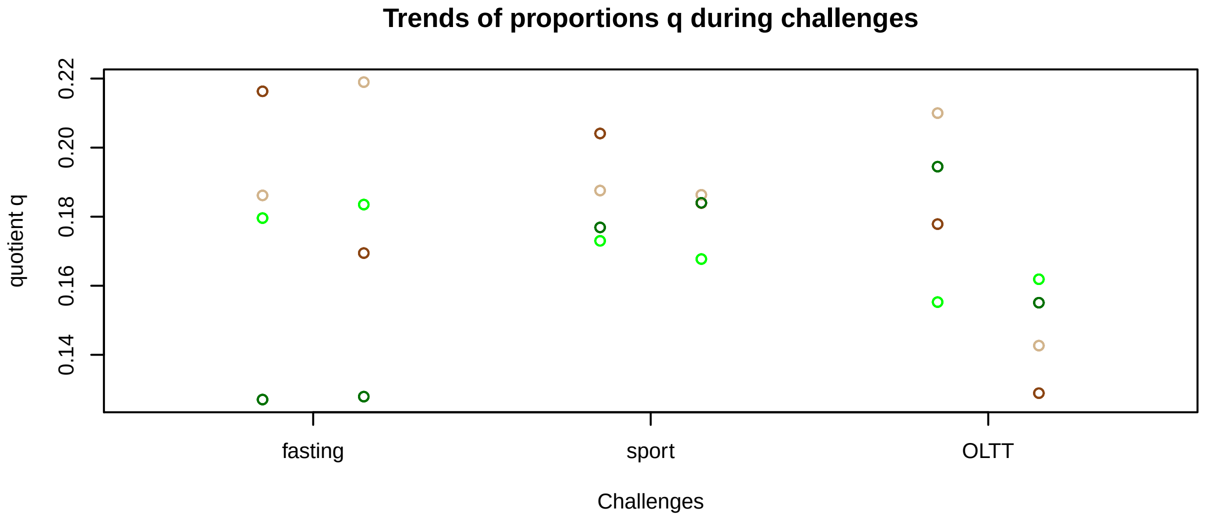


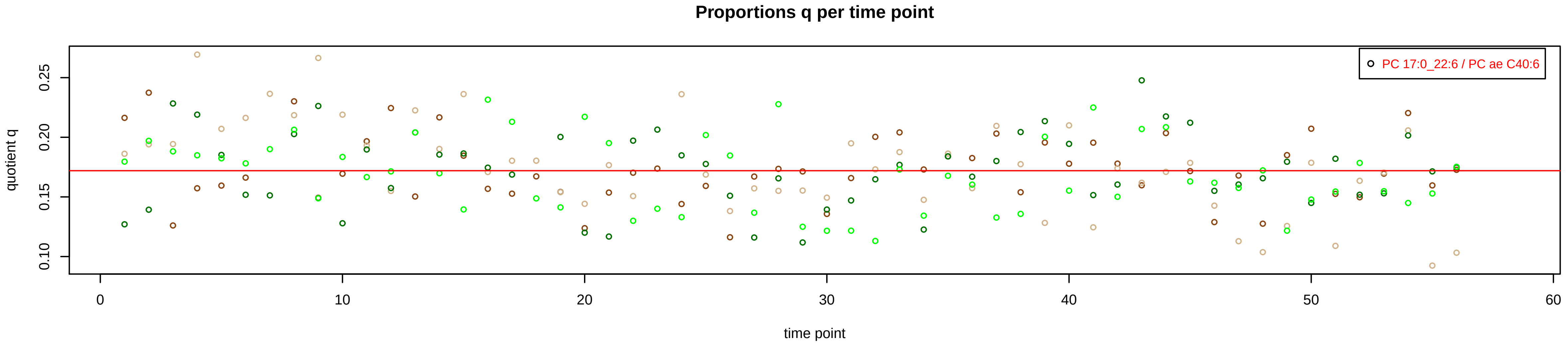

# PC ae C42:0 = R

No independent Variable measured.

### Qualitative composition

PC ae C42:0 consists of:
PC O−20:0_22:0, PC 19:0_22:0, PC 20:0_21:0, PC 23:0_18:(
PC 20:5_22:2
and further compounds

# PC ae C42:1 = R

No independent Variable measured.

### Qualitative composition

PC ae C42:1 consists of:
PC O−20:0_22:1, PC O−18:1_24:0, PC O−20:1_22:0, PC 19:
PC 19:1_22:0, PC 20:1_21:0, PC 20:2_22:6, PC 20:4_22:4,
and further compounds

# PC ae C42:2 = R

**No independent Variable measured.**

### Qualitative composition

PC ae C42:2 consists of:
PC O−20:0_22:2, PC O−18:2_24:0, PC O−20:1_22:1, PC O−
PC 19:0_22:2, PC 19:1_22:1, PC 20:2_21:0, PC 20:3_22:6,
PC 20:5_22:4
and further compounds

# PC ae C42:3 = R

**No independent Variable measured.**

### Qualitative composition

PC ae C42:3 consists of:
PC O−24:0_18:3, PC O−20:1_22:2, PC O−18:2_24:1, PC 21:
PC 19:1_22:2, PC 20:3_21:0, PC 20:5_22:5, PC 20:4_22:6,
and further compounds

# PC ae C42:4 = R

### Qualitative composition

PC ae C42:4 consists of:
PC O−20:0_22:4, PC 19:0_22:4, PC 20:4_21:0, PC 20:5_22:[illegible]
and further compounds

**No independent Variable measured.**

# PC ae C42:5 = R

No independent Variable measured.

### Qualitative composition

PC ae C42:5 consists of:
PC 20:5_21:0
and further compounds

# PC ae C44:3 = R

No independent Variable measured.

### Qualitative composition

PC ae C44:3 consists of:
PC 22:4_22:6
and further compounds

# PC ae C44:4 = R

### Qualitative composition

PC ae C44:4 consists of:
PC O−24:0_20:4, PC O−22:1_22:3, PC O−22:2_22:2, PC 21:
and further compounds

**No independent Variable measured.**

# PC ae C44:5 = R

### Qualitative composition

PC ae C44:5 consists of:
PC 22:6_22:6
and further compounds

**No independent Variable measured.**

# PC ae C44:6 = R

### Qualitative composition

PC ae C44:6 consists of:
PC 21:0_22:6
and further compounds

**No independent Variable measured.**
