## Supplemental Figure S2 for "Characterization of bulk phosphatidylcholine compositions in human plasma using side-chain resolving lipidomics"

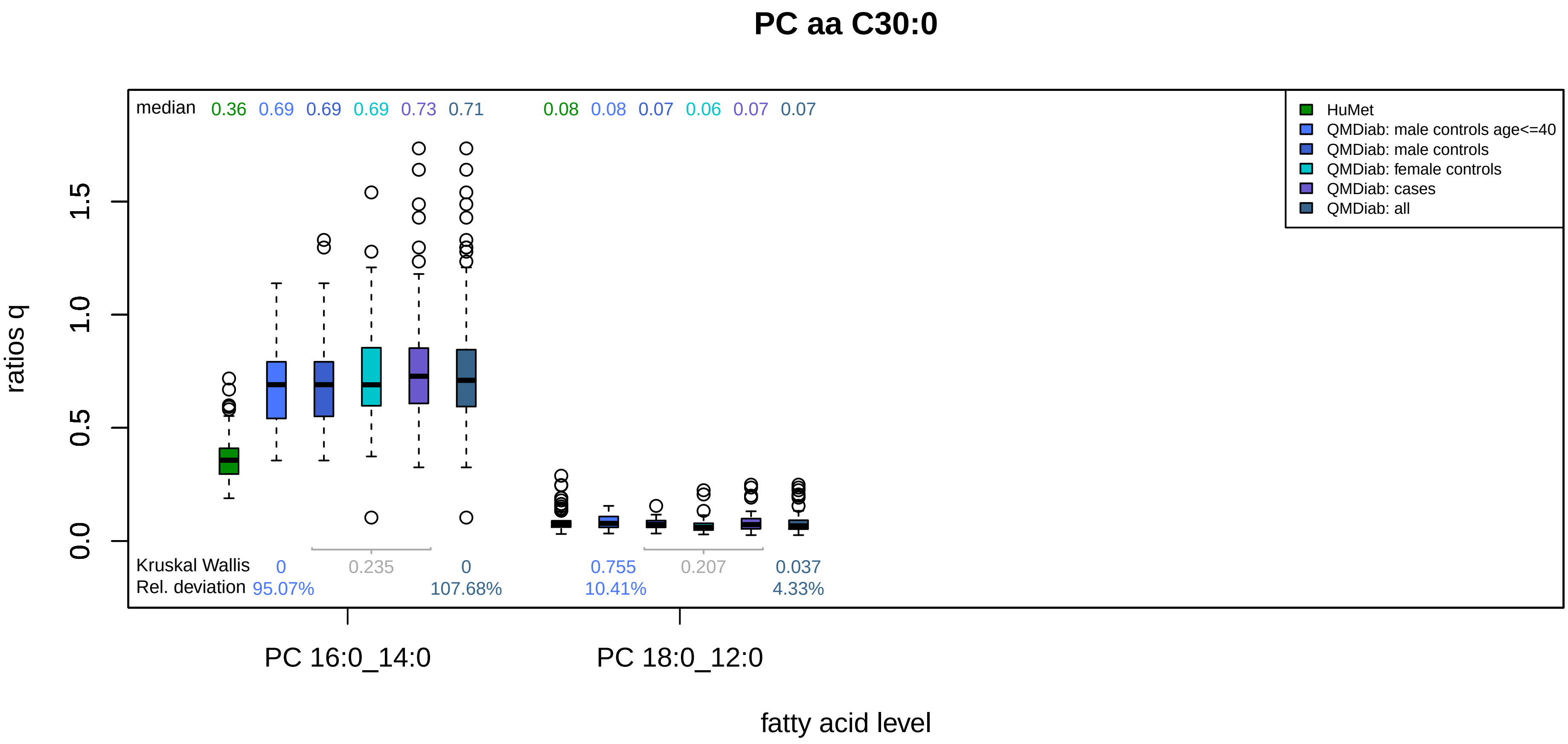

PC aa C30:0
median 0.36 0.69 0.69 0.69 0.73 0.71 0.08 0.08 0.07 0.06 0.07 0.07
HuMet
QMDiab: male controls age<=40
QMDiab: male controls
QMDiab: female controls
QMDiab: cases
QMDiab: all
ratios q
0.0
0.5
1.0
1.5
Kruskal Wallis 0 0.235 0 0.755 0.207 0.037
Rel. deviation 95.07% 107.68% 10.41% 4.33%
PC 16:0_14:0
PC 18:0_12:0
fatty acid level

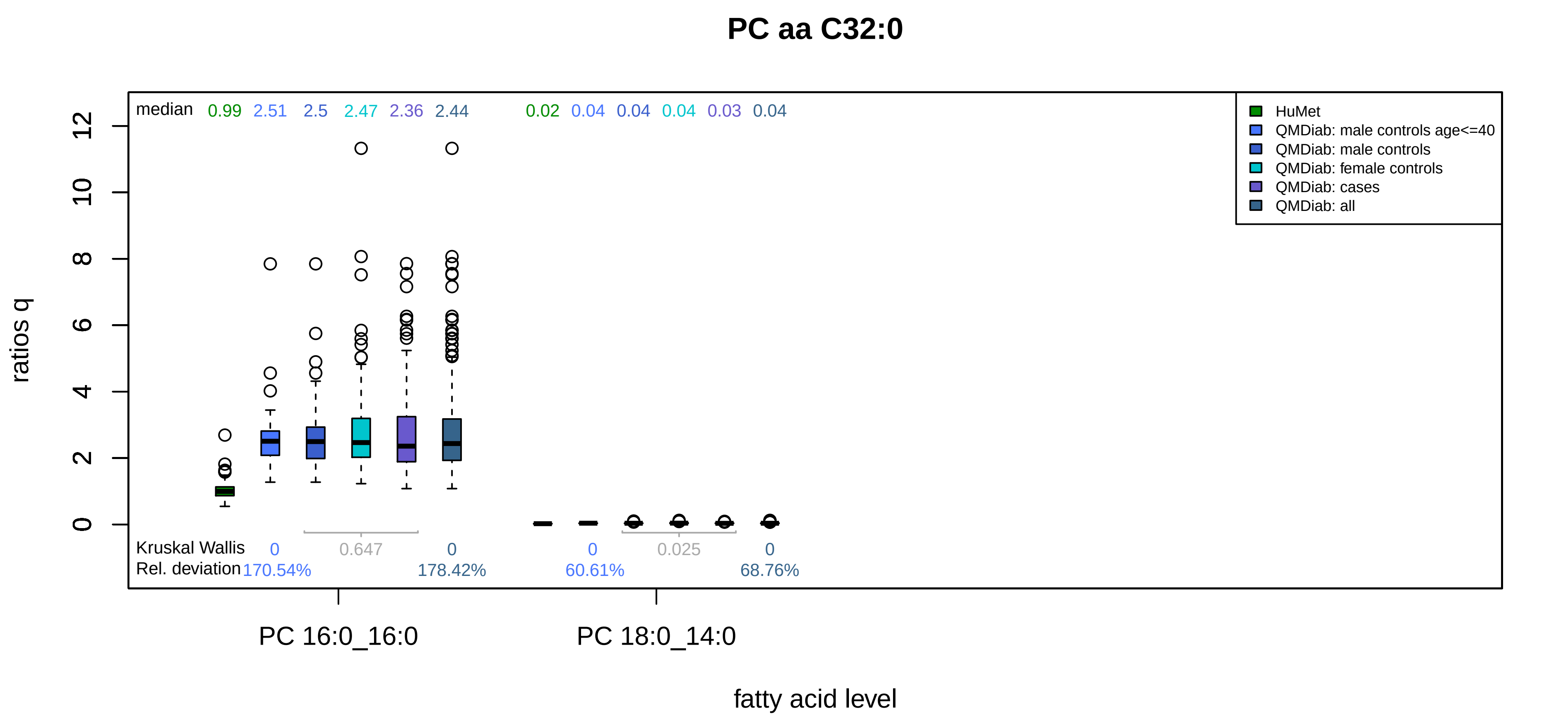

PC aa C32:0
median
0.99
2.51
2.5
2.47
2.36
2.44
0.02
0.04
0.04
0.04
0.03
0.04
HuMet
QMDiab: male controls age<=40
QMDiab: male controls
QMDiab: female controls
QMDiab: cases
QMDiab: all
ratios q
0
2
4
6
8
10
12
Kruskal Wallis
0
0.647
0
0
0.025
0
Rel. deviation
170.54%
178.42%
60.61%
68.76%
PC 16:0_16:0
PC 18:0_14:0
fatty acid level

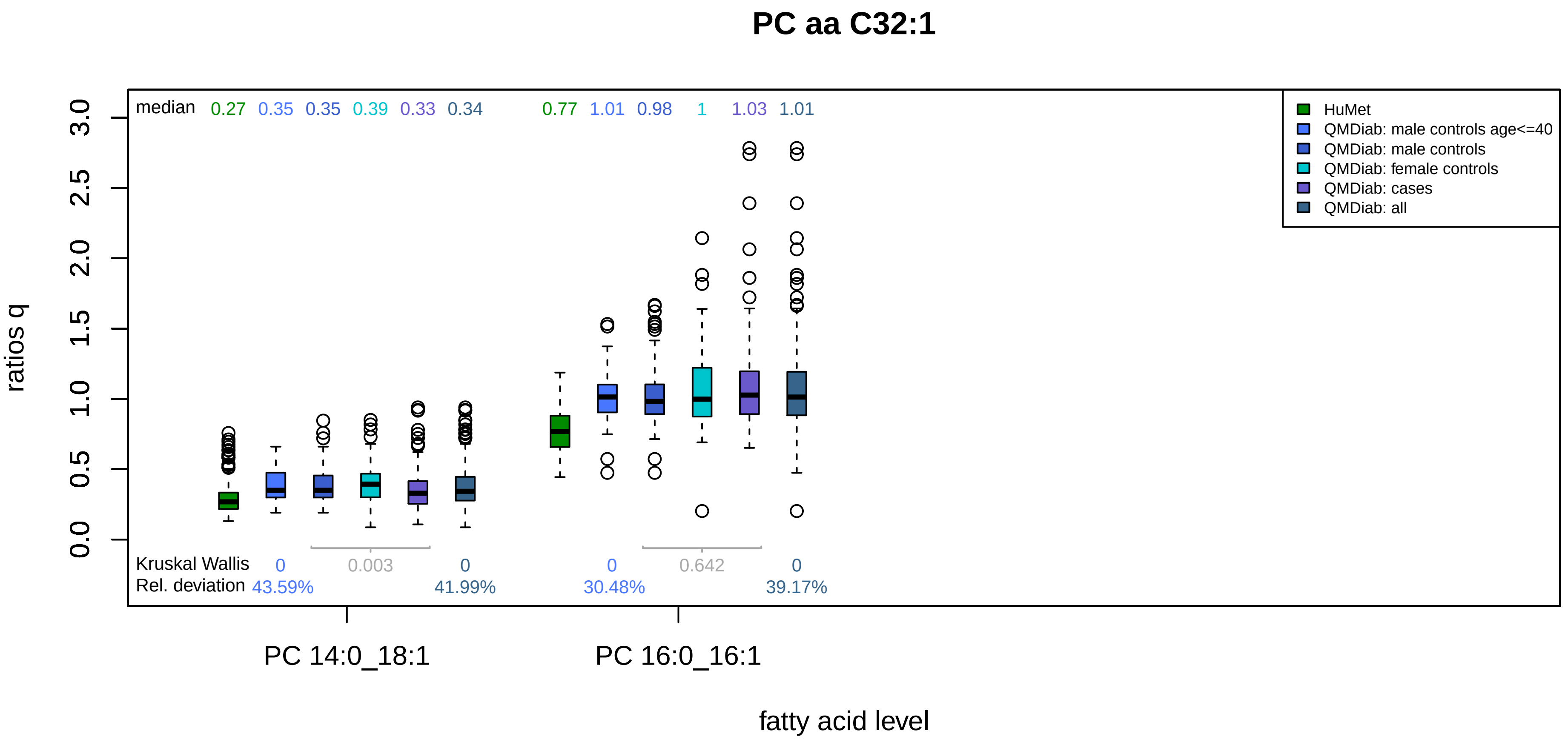

PC aa C32:1
median 0.27 0.35 0.35 0.39 0.33 0.34 0.77 1.01 0.98 1 1.03 1.01
HuMet
QMDiab: male controls age<=40
QMDiab: male controls
QMDiab: female controls
QMDiab: cases
QMDiab: all
ratios q
0.0 0.5 1.0 1.5 2.0 2.5 3.0
Kruskal Wallis 0 0.003 0 0 0.642 0
Rel. deviation 43.59% 41.99% 30.48% 39.17%
PC 14:0_18:1
PC 16:0_16:1
fatty acid level

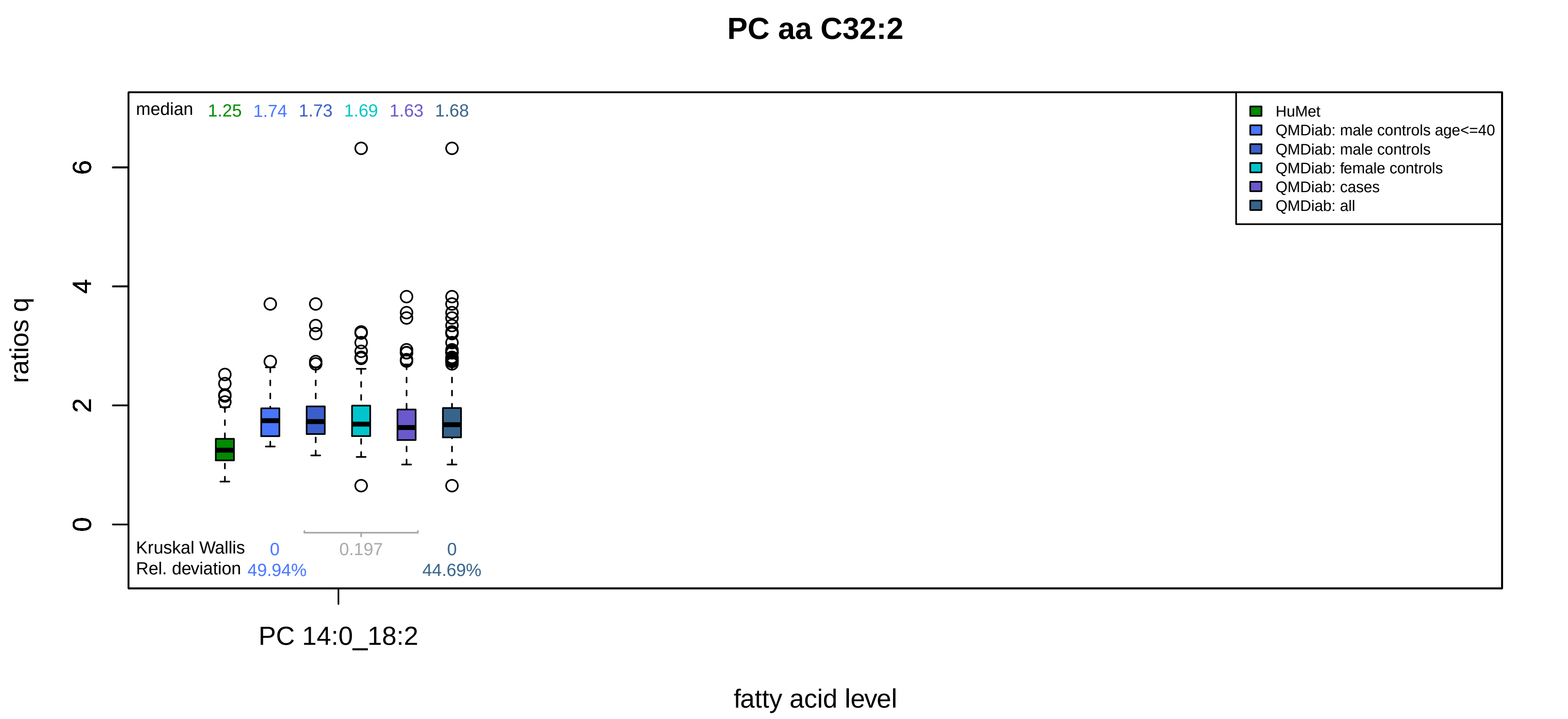
PC aa C32:2
median 1.25 1.74 1.73 1.69 1.63 1.68
HuMet
QMDiab: male controls age<=40
QMDiab: male controls
QMDiab: female controls
QMDiab: cases
QMDiab: all
ratios q
0
2
4
6
Kruskal Wallis 0 0.197 0
Rel. deviation 49.94% 44.69%
PC 14:0_18:2
fatty acid level

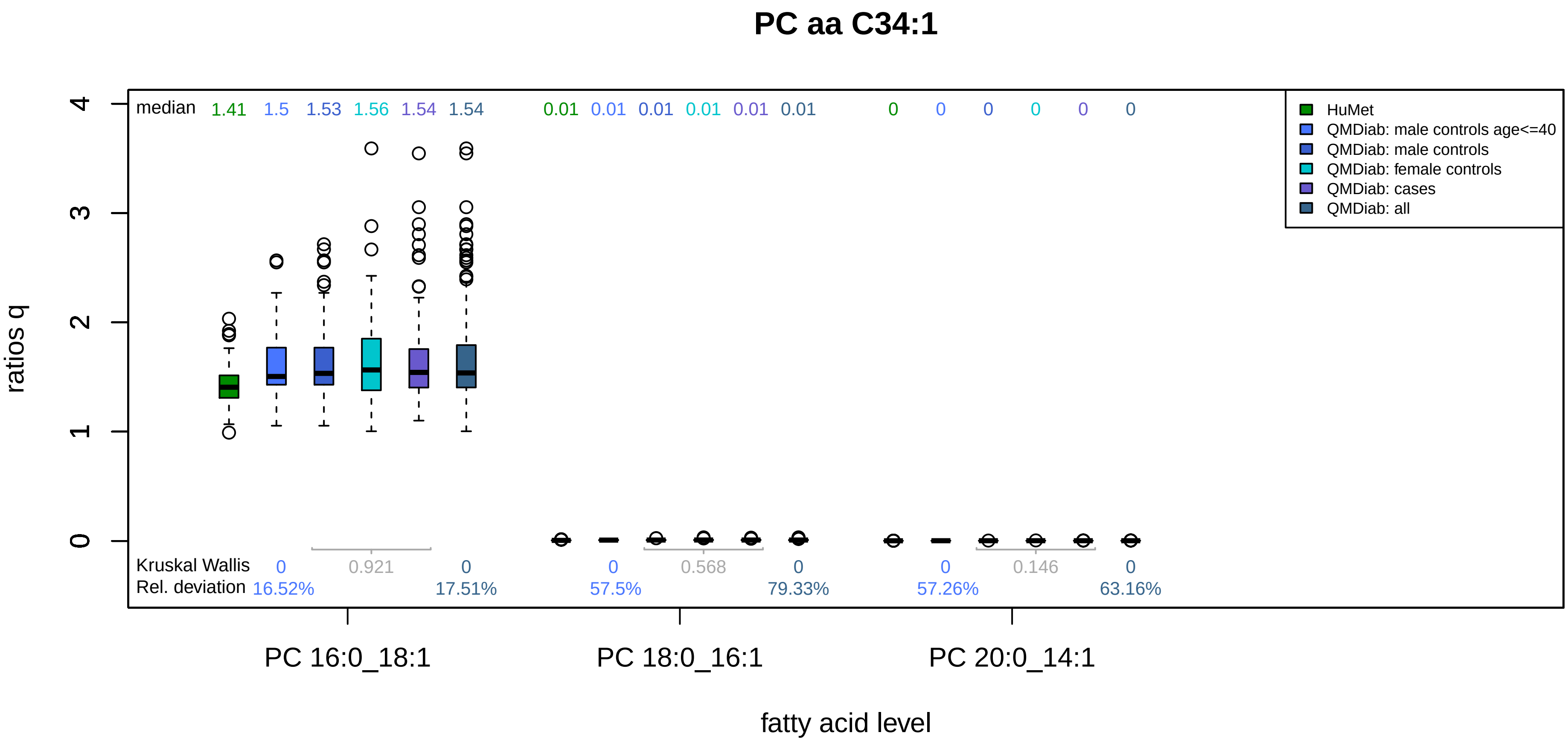
PC aa C34:1
median 1.41 1.5 1.53 1.56 1.54 1.54 0.01 0.01 0.01 0.01 0.01 0.01 0 0 0 0 0 0
HuMet
QMDiab: male controls age<=40
QMDiab: male controls
QMDiab: female controls
QMDiab: cases
QMDiab: all
ratios q
0
1
2
3
4
Kruskal Wallis 0 0.921 0 0 0.568 0 0 0.146 0
Rel. deviation 16.52% 17.51% 57.5% 79.33% 57.26% 63.16%
PC 16:0_18:1
PC 18:0_16:1
PC 20:0_14:1
fatty acid level

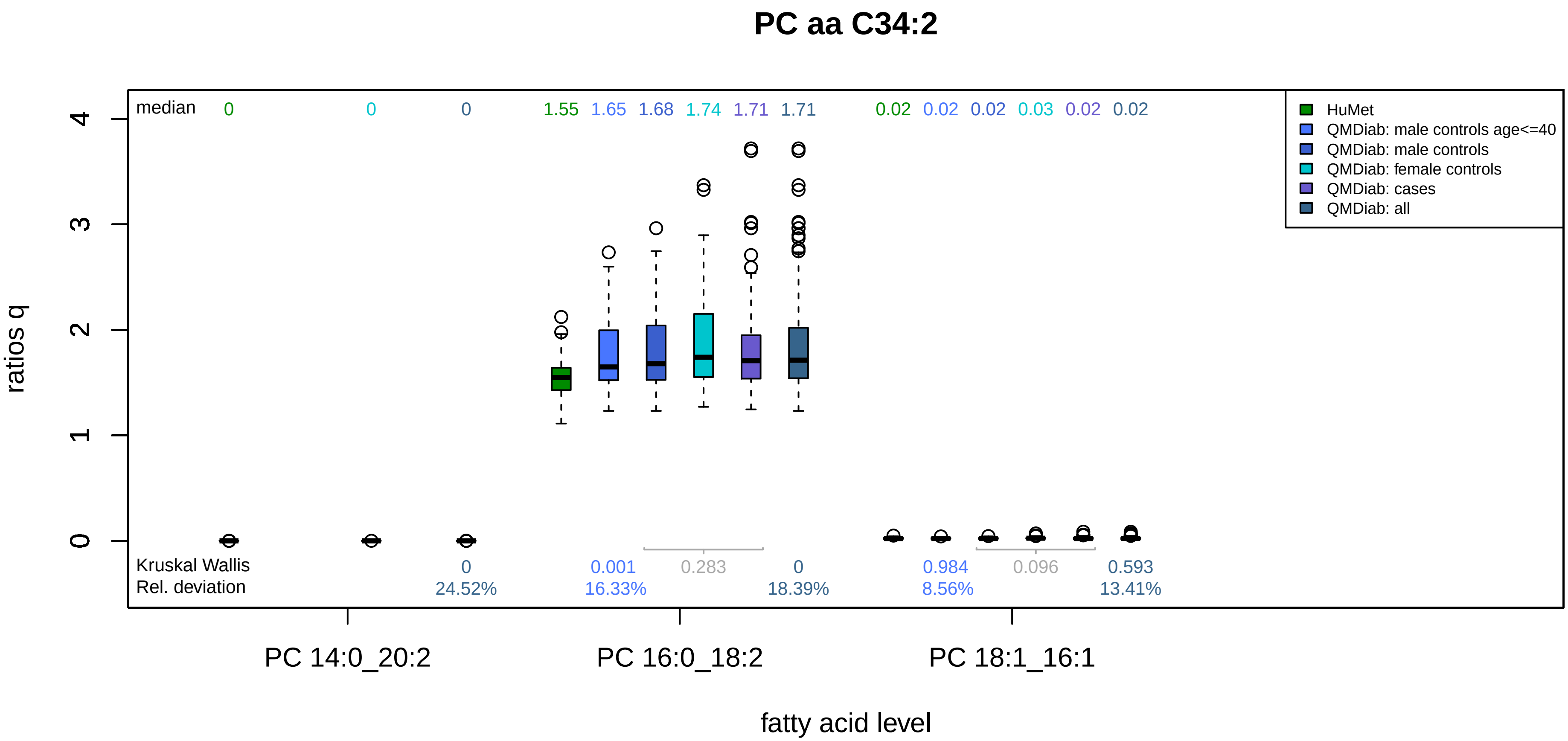
PC aa C34:2
median
0
0
0
1.55
1.65
1.68
1.74
1.71
1.71
0.02
0.02
0.02
0.03
0.02
0.02
HuMet
QMDiab: male controls age<=40
QMDiab: male controls
QMDiab: female controls
QMDiab: cases
QMDiab: all
ratios q
0
1
2
3
4
Kruskal Wallis
Rel. deviation
0
24.52%
0.001
16.33%
0.283
0
18.39%
0.984
8.56%
0.096
0.593
13.41%
PC 14:0_20:2
PC 16:0_18:2
PC 18:1_16:1
fatty acid level

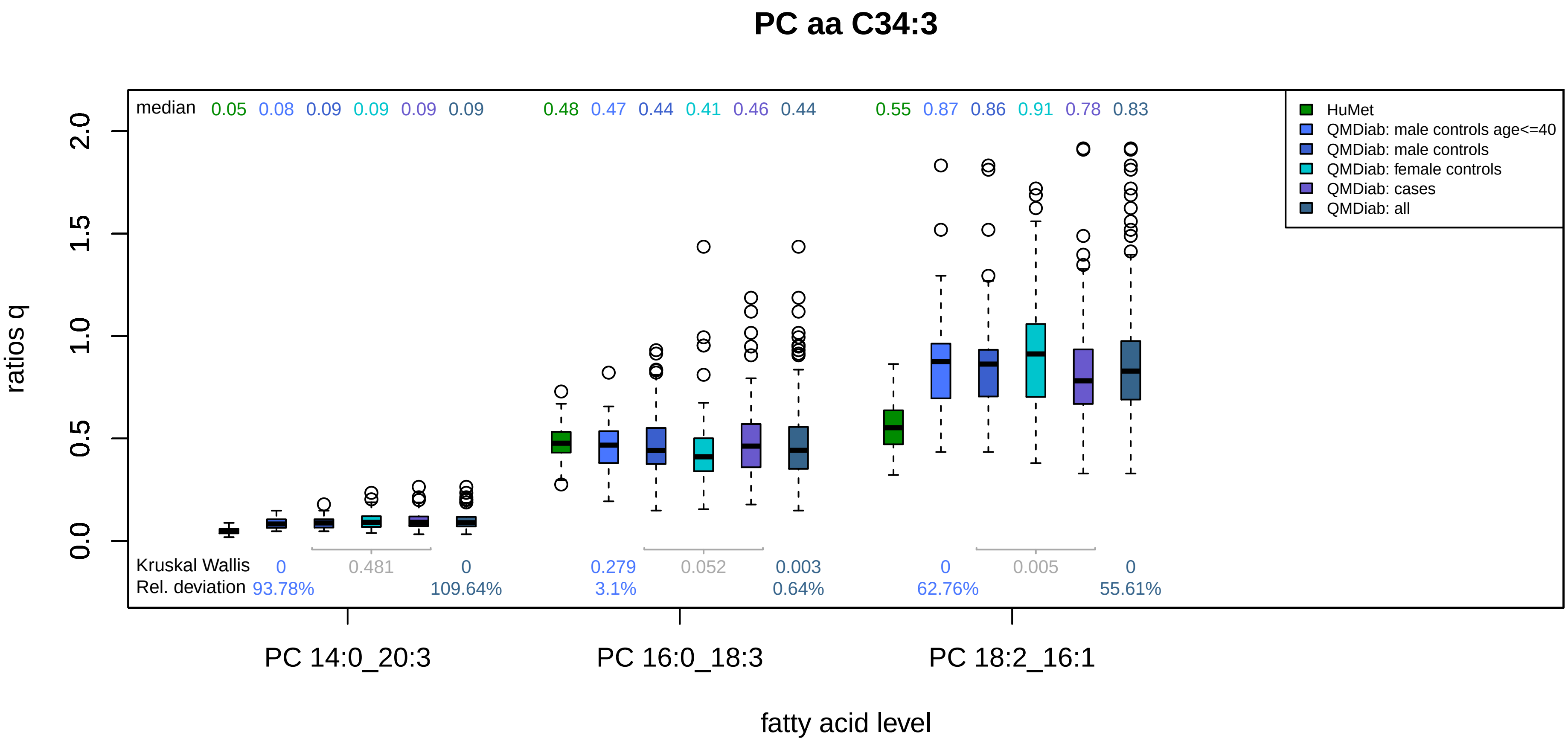

PC aa C34:3
median 0.05 0.08 0.09 0.09 0.09 0.09 0.48 0.47 0.44 0.41 0.46 0.44 0.55 0.87 0.86 0.91 0.78 0.83
HuMet
QMDiab: male controls age<=40
QMDiab: male controls
QMDiab: female controls
QMDiab: cases
QMDiab: all
ratios q
0.0
0.5
1.0
1.5
2.0
Kruskal Wallis 0 0.481 0 0.279 0.052 0.003 0 0.005 0
Rel. deviation 93.78% 109.64% 3.1% 0.64% 62.76% 55.61%
PC 14:0_20:3
PC 16:0_18:3
PC 18:2_16:1
fatty acid level

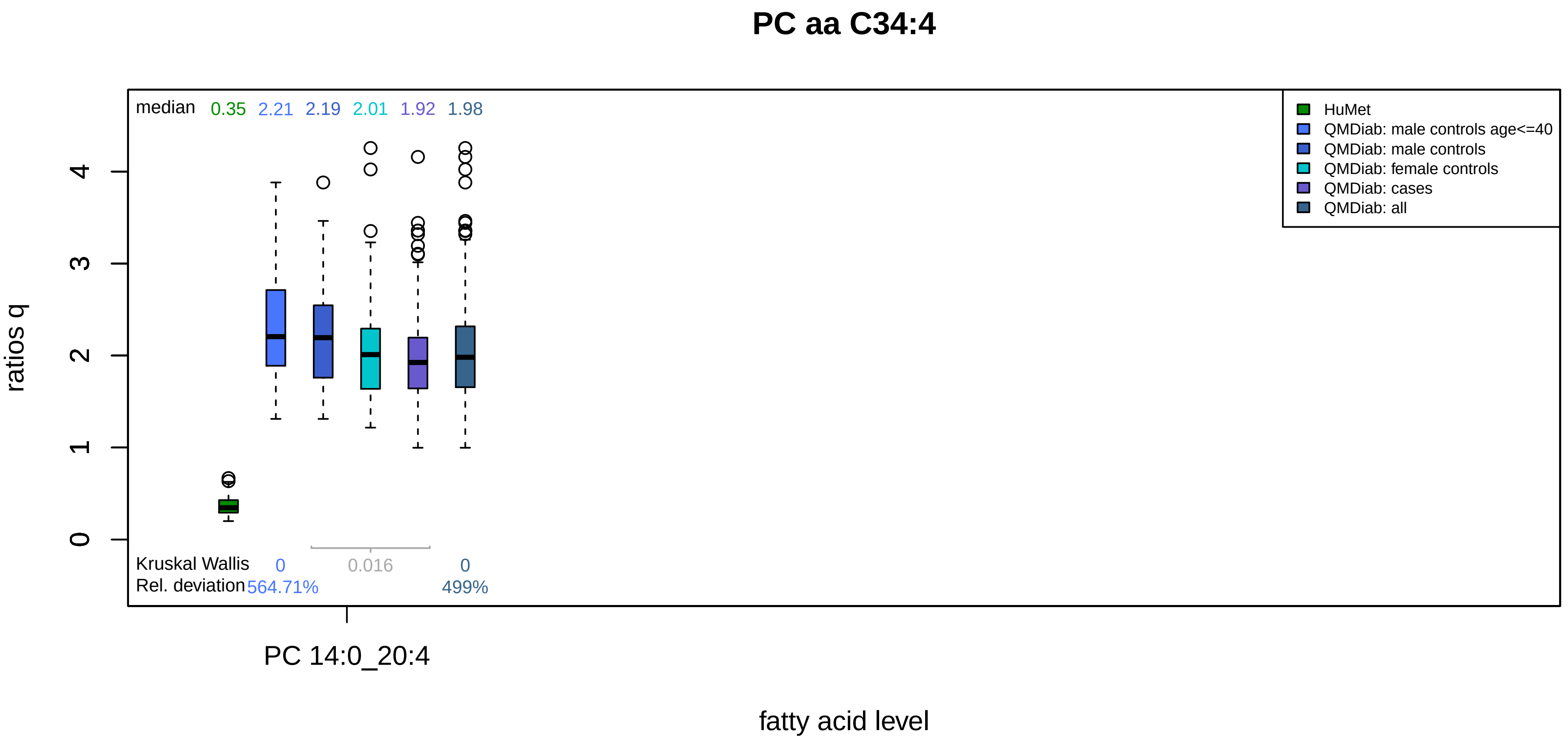
PC aa C34:4
median 0.35 2.21 2.19 2.01 1.92 1.98
HuMet
QMDiab: male controls age<=40
QMDiab: male controls
QMDiab: female controls
QMDiab: cases
QMDiab: all
ratios q
0
1
2
3
4
Kruskal Wallis 0 0.016 0
Rel. deviation 564.71% 499%
PC 14:0_20:4
fatty acid level

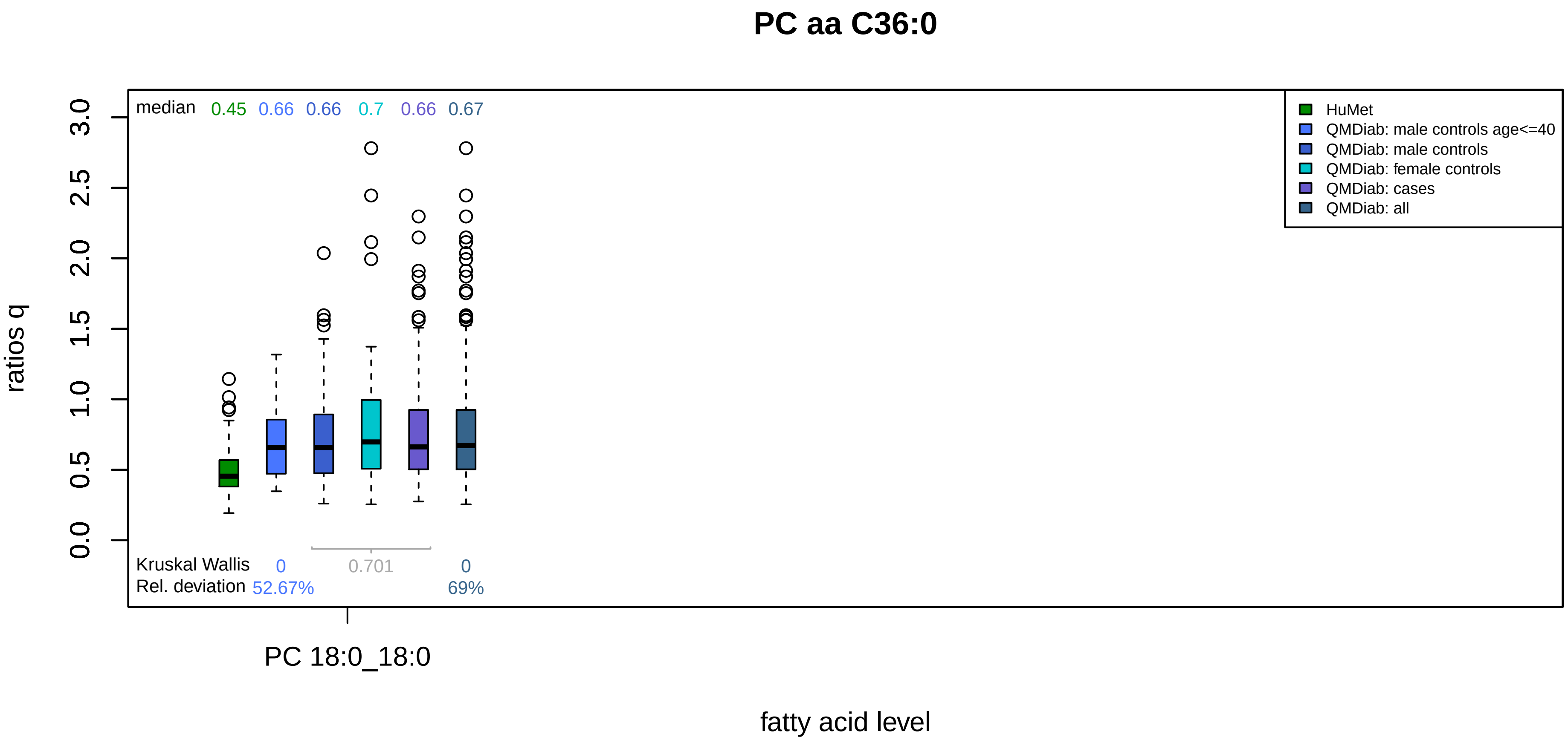

PC aa C36:0
median 0.45 0.66 0.66 0.7 0.66 0.67
HuMet
QMDiab: male controls age<=40
QMDiab: male controls
QMDiab: female controls
QMDiab: cases
QMDiab: all
ratios q
0.0
0.5
1.0
1.5
2.0
2.5
3.0
Kruskal Wallis 0 0.701 0
Rel. deviation 52.67% 69%
PC 18:0_18:0
fatty acid level

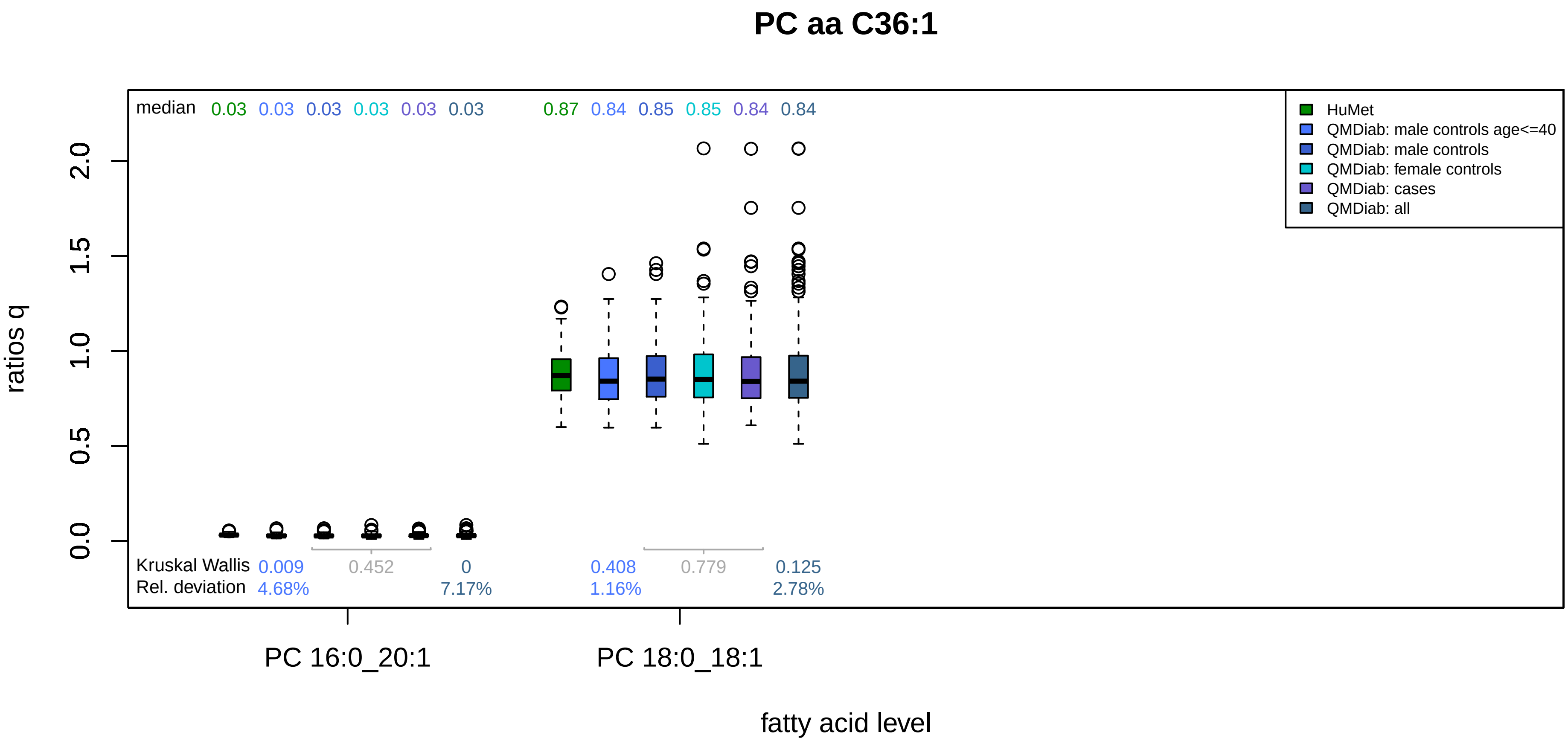

PC aa C36:1
median 0.03 0.03 0.03 0.03 0.03 0.03 0.87 0.84 0.85 0.85 0.84 0.84
HuMet
QMDiab: male controls age<=40
QMDiab: male controls
QMDiab: female controls
QMDiab: cases
QMDiab: all
ratios q
0.0
0.5
1.0
1.5
2.0
Kruskal Wallis 0.009 0.452 0 0.408 0.779 0.125
Rel. deviation 4.68% 7.17% 1.16% 2.78%
PC 16:0_20:1
PC 18:0_18:1
fatty acid level

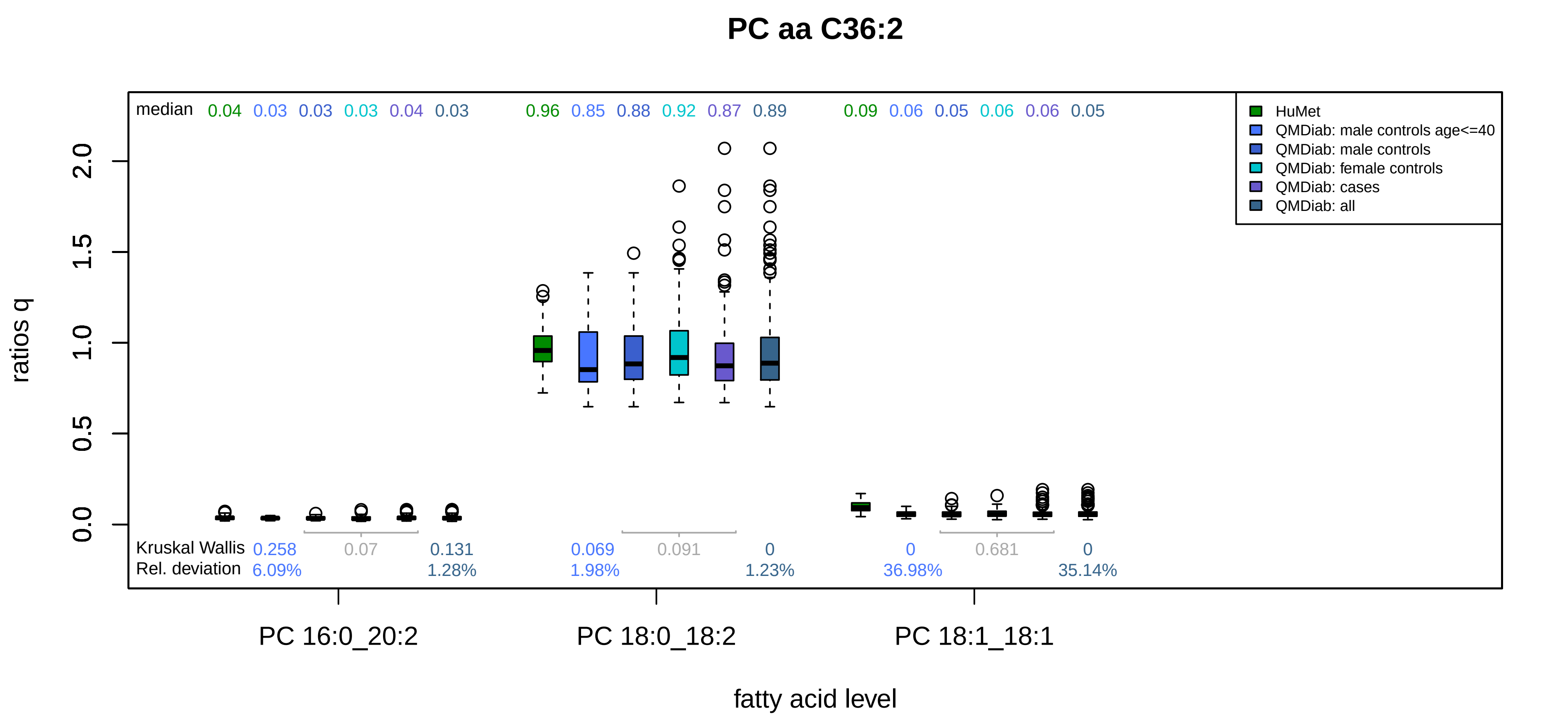

PC aa C36:2
median 0.04 0.03 0.03 0.03 0.04 0.03 0.96 0.85 0.88 0.92 0.87 0.89 0.09 0.06 0.05 0.06 0.06 0.05
HuMet
QMDiab: male controls age<=40
QMDiab: male controls
QMDiab: female controls
QMDiab: cases
QMDiab: all
ratios q
0.0
0.5
1.0
1.5
2.0
Kruskal Wallis 0.258 0.07 0.131 0.069 0.091 0 0 0.681 0
Rel. deviation 6.09% 1.28% 1.98% 1.23% 36.98% 35.14%
PC 16:0_20:2
PC 18:0_18:2
PC 18:1_18:1
fatty acid level

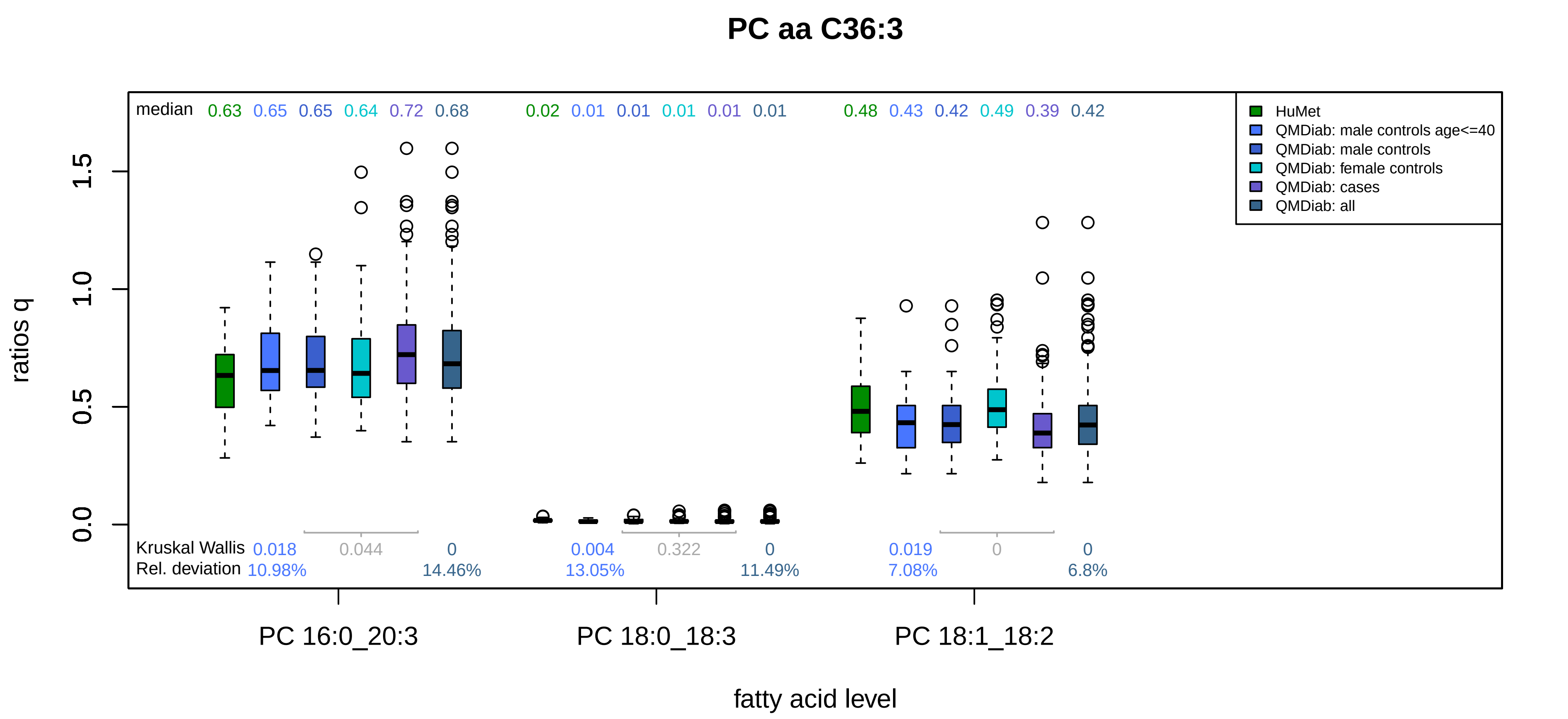
PC aa C36:3
median 0.63 0.65 0.65 0.64 0.72 0.68 0.02 0.01 0.01 0.01 0.01 0.01 0.48 0.43 0.42 0.49 0.39 0.42
HuMet
QMDiab: male controls age<=40
QMDiab: male controls
QMDiab: female controls
QMDiab: cases
QMDiab: all
ratios q
0.0
0.5
1.0
1.5
Kruskal Wallis 0.018 0.044 0 0.004 0.322 0 0.019 0 0
Rel. deviation 10.98% 14.46% 13.05% 11.49% 7.08% 6.8%
PC 16:0_20:3
PC 18:0_18:3
PC 18:1_18:2
fatty acid level

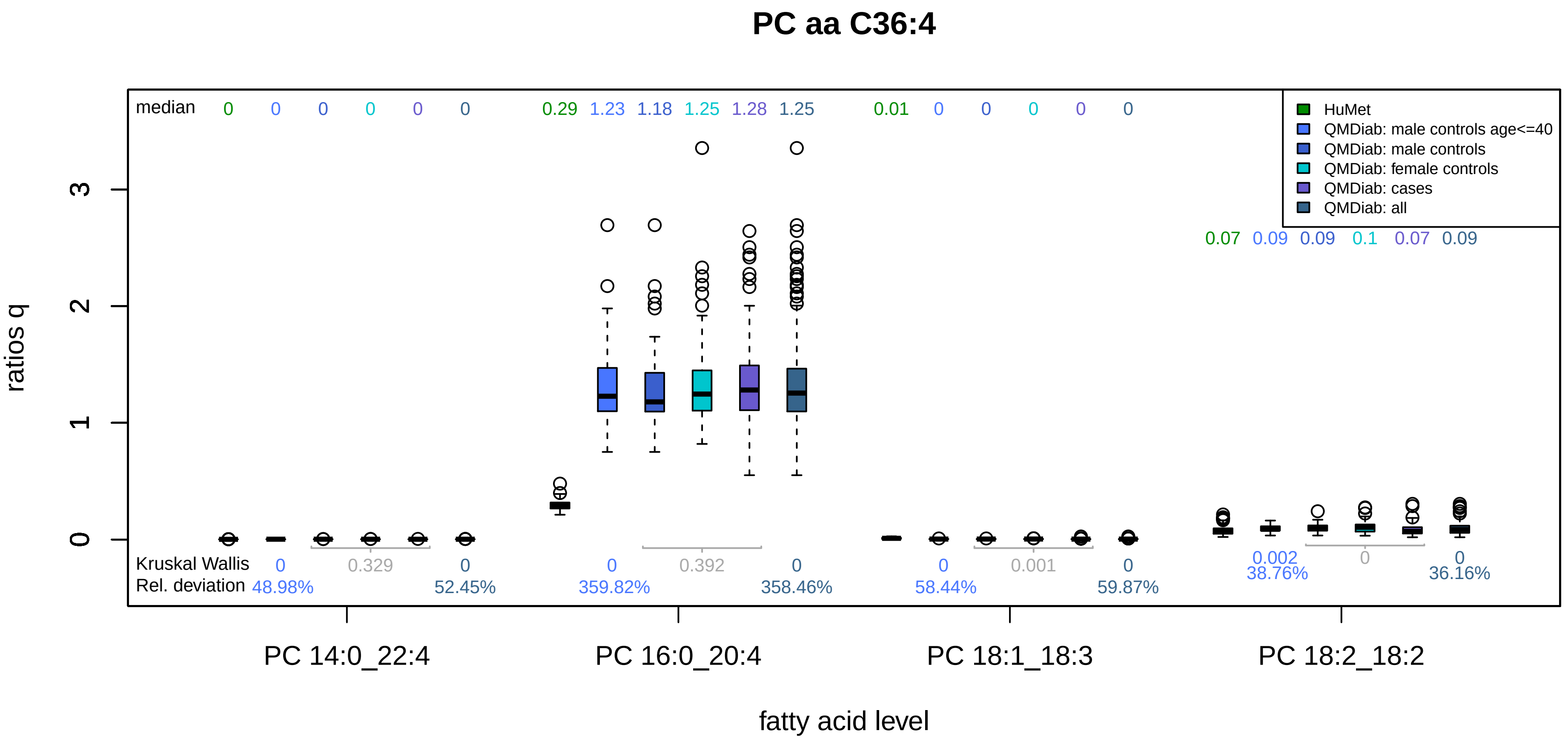
PC aa C36:4
median
0 0 0 0 0 0
0.29 1.23 1.18 1.25 1.28 1.25
0.01 0 0 0 0 0
0.07 0.09 0.09 0.1 0.07 0.09
HuMet
QMDiab: male controls age<=40
QMDiab: male controls
QMDiab: female controls
QMDiab: cases
QMDiab: all
ratios q
0
1
2
3
Kruskal Wallis
Rel. deviation
0 0.329 0
48.98% 52.45%
0 0.392 0
359.82% 358.46%
0 0.001 0
58.44% 59.87%
0.002 0 0
38.76% 36.16%
PC 14:0_22:4
PC 16:0_20:4
PC 18:1_18:3
PC 18:2_18:2
fatty acid level

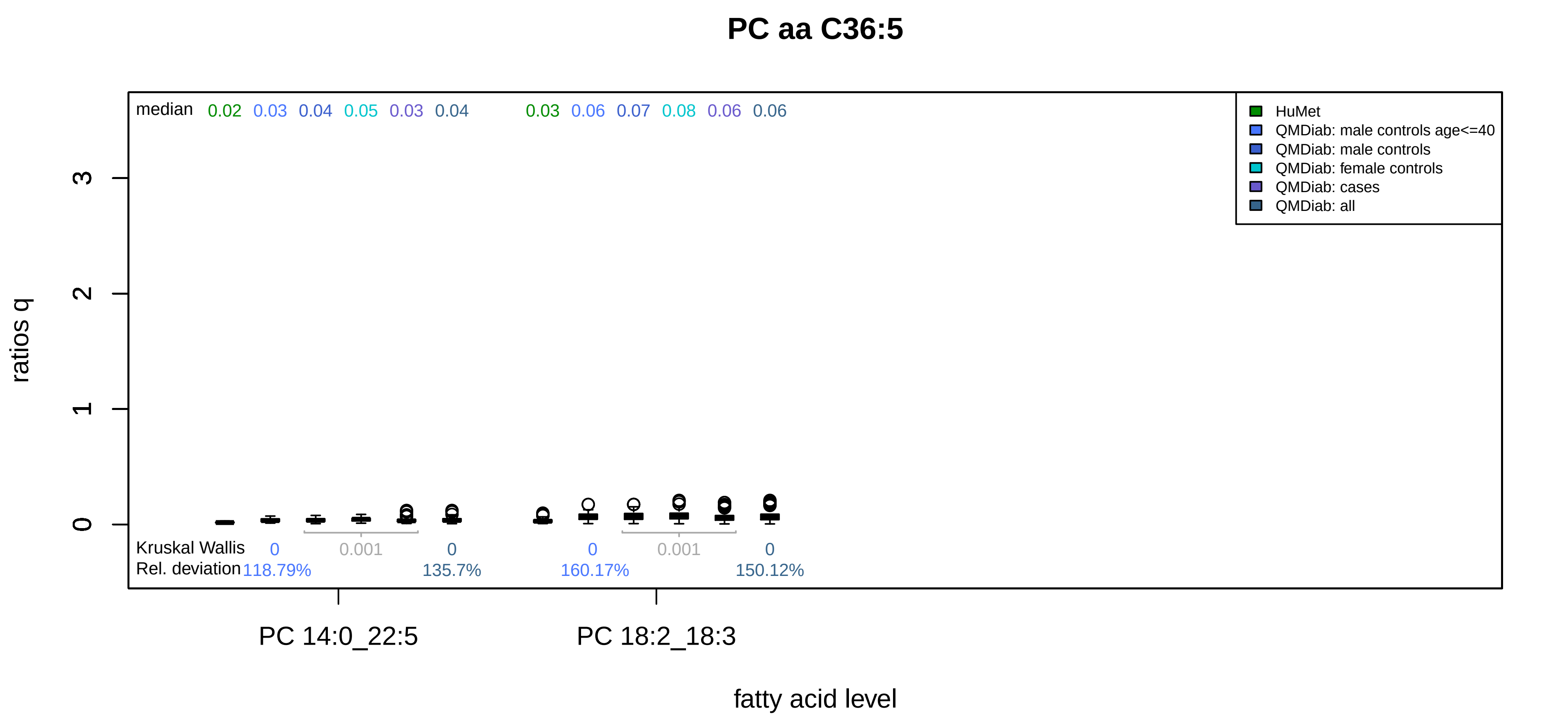

PC aa C36:5
median 0.02 0.03 0.04 0.05 0.03 0.04 0.03 0.06 0.07 0.08 0.06 0.06
HuMet
QMDiab: male controls age<=40
QMDiab: male controls
QMDiab: female controls
QMDiab: cases
QMDiab: all
ratios q
0
1
2
3
Kruskal Wallis 0 0.001 0 0 0.001 0
Rel. deviation 118.79% 135.7% 160.17% 150.12%
PC 14:0_22:5
PC 18:2_18:3
fatty acid level

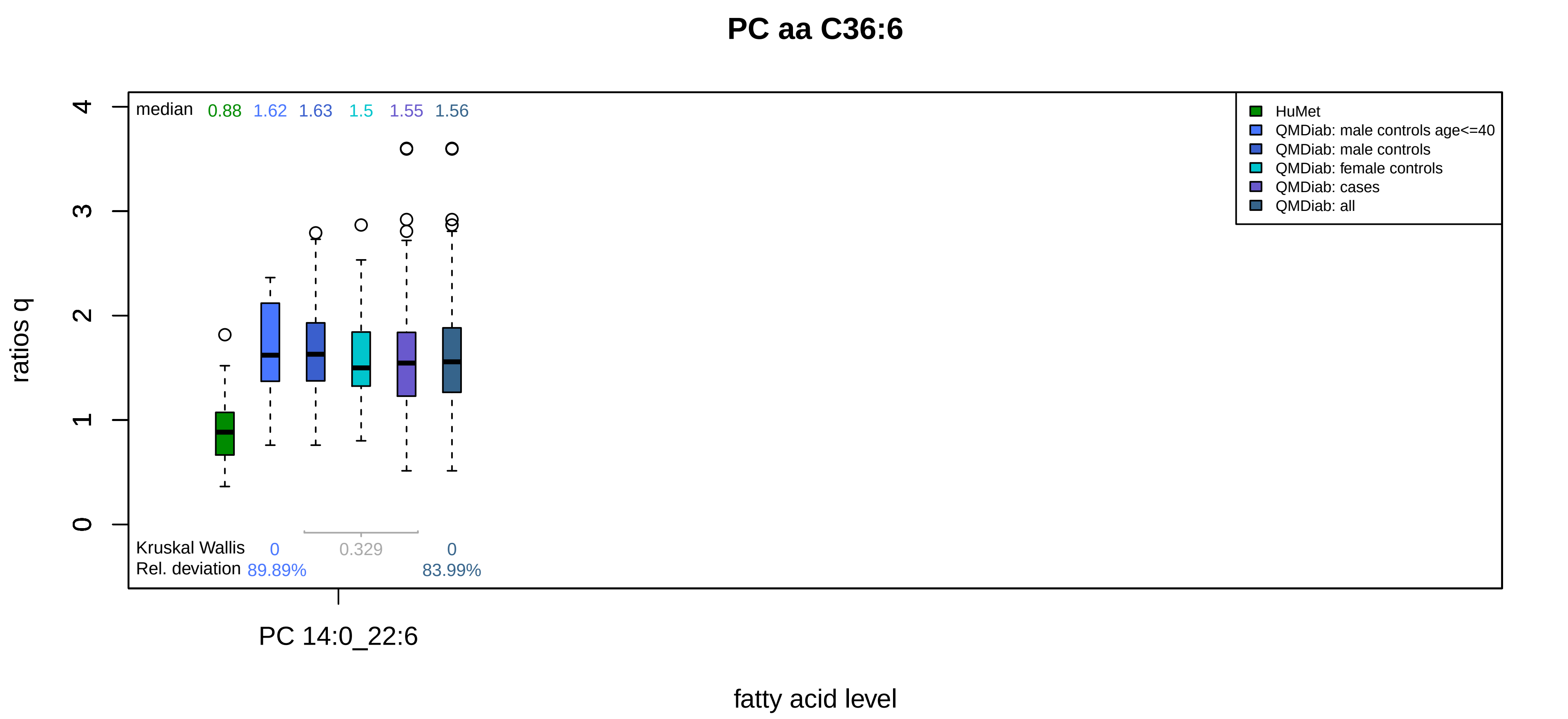
PC aa C36:6
median 0.88 1.62 1.63 1.5 1.55 1.56
HuMet
QMDiab: male controls age<=40
QMDiab: male controls
QMDiab: female controls
QMDiab: cases
QMDiab: all
ratios q
0
1
2
3
4
Kruskal Wallis 0 0.329 0
Rel. deviation 89.89% 83.99%
PC 14:0_22:6
fatty acid level

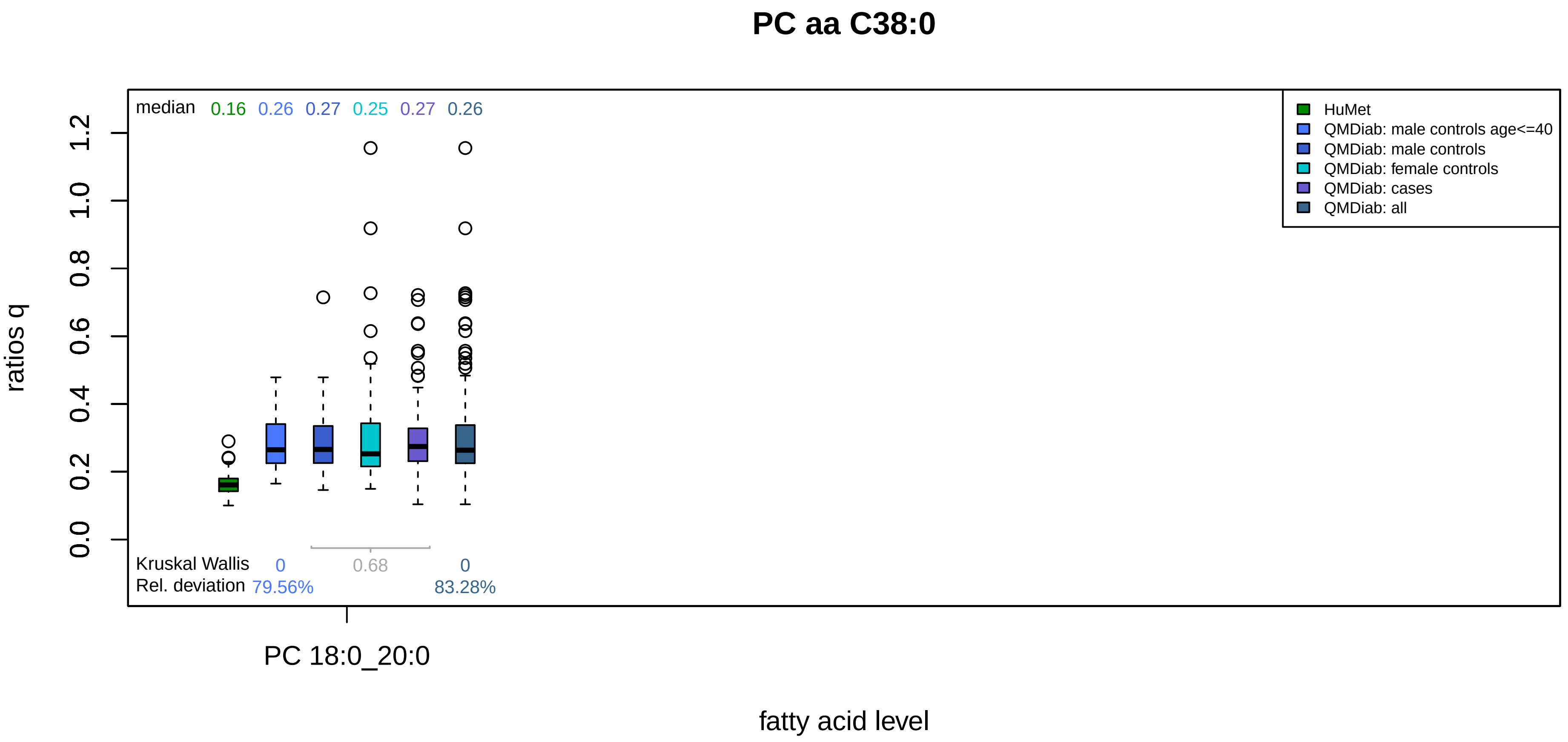

PC aa C38:0
median 0.16 0.26 0.27 0.25 0.27 0.26
HuMet
QMDiab: male controls age<=40
QMDiab: male controls
QMDiab: female controls
QMDiab: cases
QMDiab: all
ratios q
0.0
0.2
0.4
0.6
0.8
1.0
1.2
Kruskal Wallis 0 0.68 0
Rel. deviation 79.56% 83.28%
PC 18:0_20:0
fatty acid level

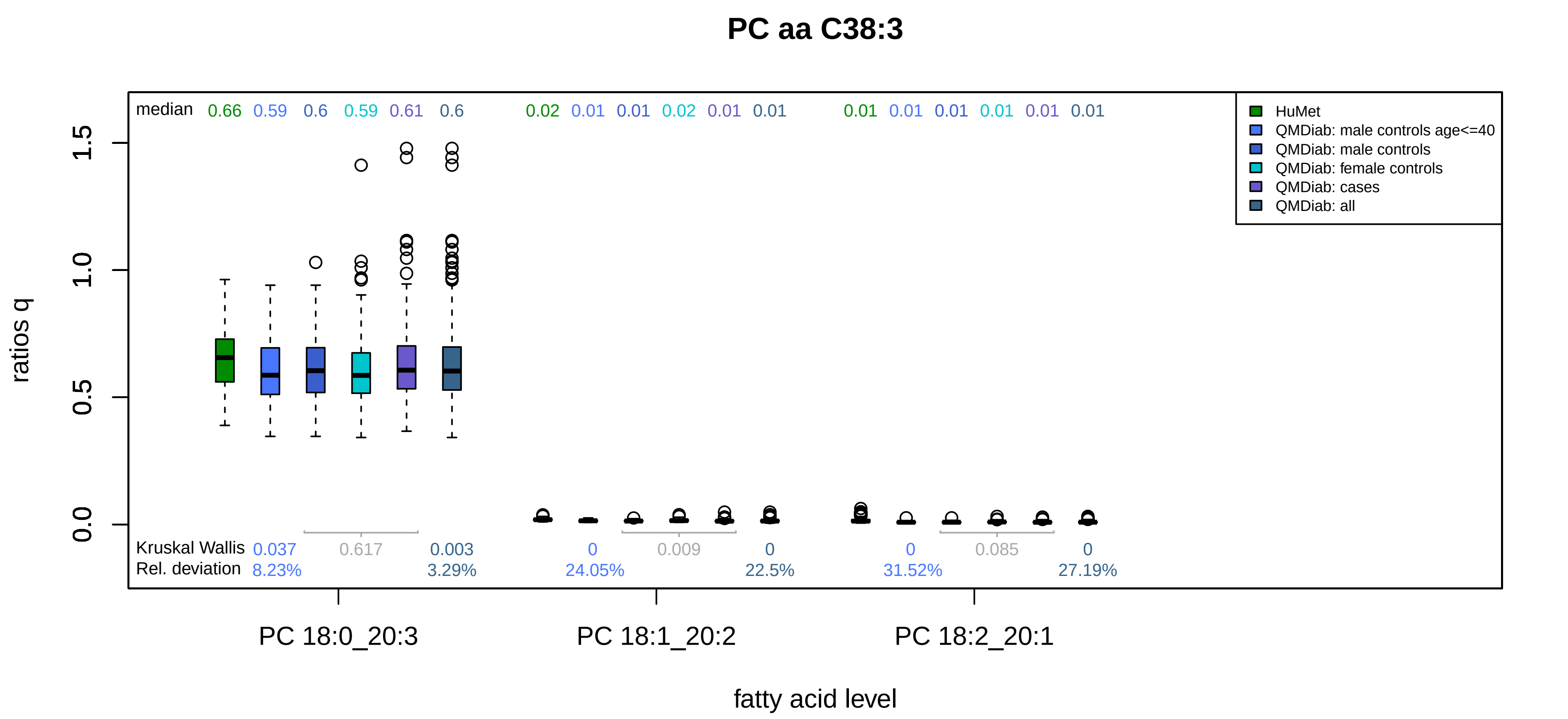

PC aa C38:3
median 0.66 0.59 0.6 0.59 0.61 0.6 0.02 0.01 0.01 0.02 0.01 0.01 0.01 0.01 0.01 0.01 0.01 0.01
HuMet
QMDiab: male controls age<=40
QMDiab: male controls
QMDiab: female controls
QMDiab: cases
QMDiab: all
ratios q
0.0
0.5
1.0
1.5
Kruskal Wallis 0.037 0.617 0.003 0 0.009 0 0 0.085 0
Rel. deviation 8.23% 3.29% 24.05% 22.5% 31.52% 27.19%
PC 18:0_20:3
PC 18:1_20:2
PC 18:2_20:1
fatty acid level

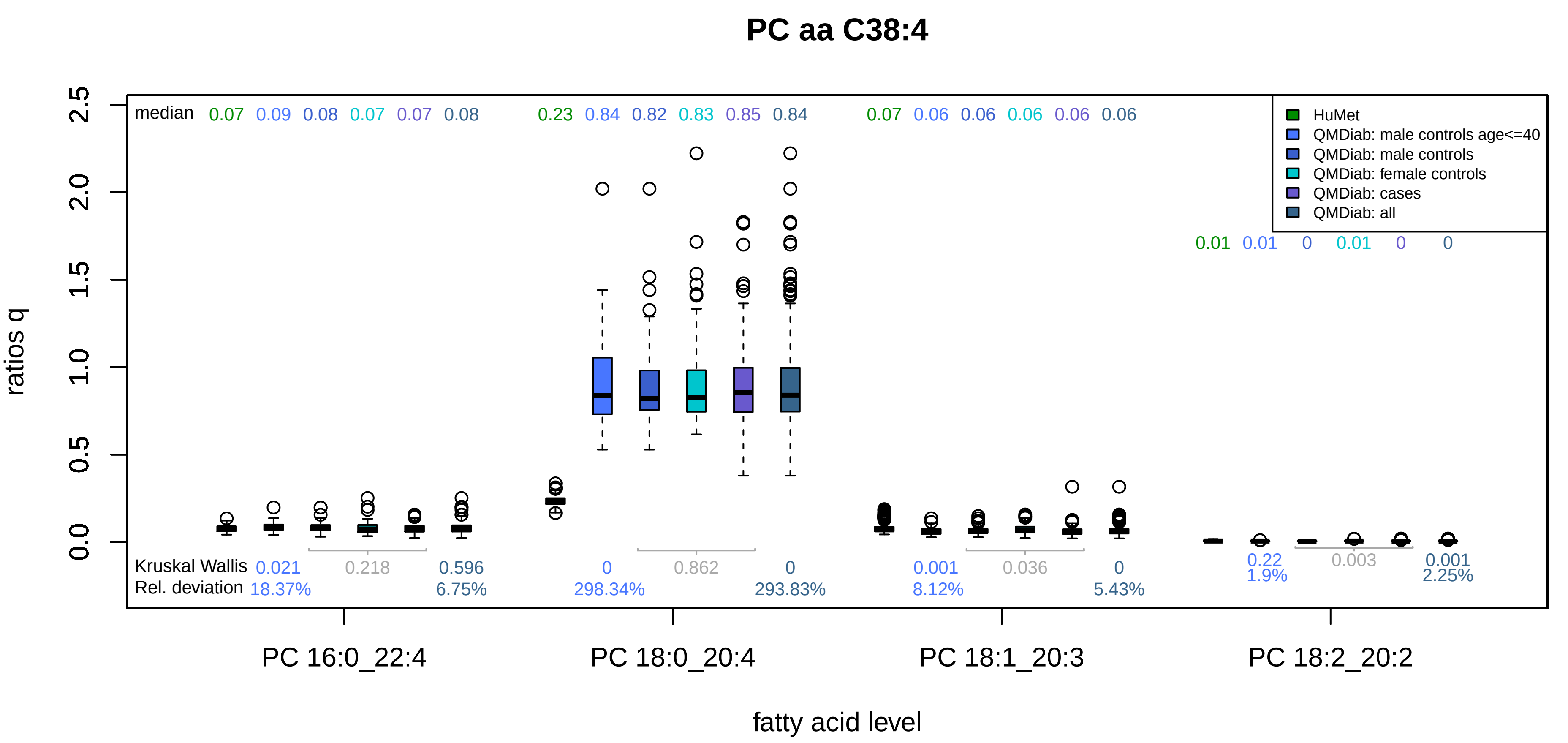

PC aa C38:4
median 0.07 0.09 0.08 0.07 0.07 0.08 0.23 0.84 0.82 0.83 0.85 0.84 0.07 0.06 0.06 0.06 0.06 0.06
0.01 0.01 0 0.01 0 0
HuMet
QMDiab: male controls age<=40
QMDiab: male controls
QMDiab: female controls
QMDiab: cases
QMDiab: all
ratios q
0.0
0.5
1.0
1.5
2.0
2.5
Kruskal Wallis 0.021 0.218 0.596 0 0.862 0 0.001 0.036 0 0.22 0.003 0.001
Rel. deviation 18.37% 6.75% 298.34% 293.83% 8.12% 5.43% 1.9% 2.25%
PC 16:0_22:4
PC 18:0_20:4
PC 18:1_20:3
PC 18:2_20:2
fatty acid level

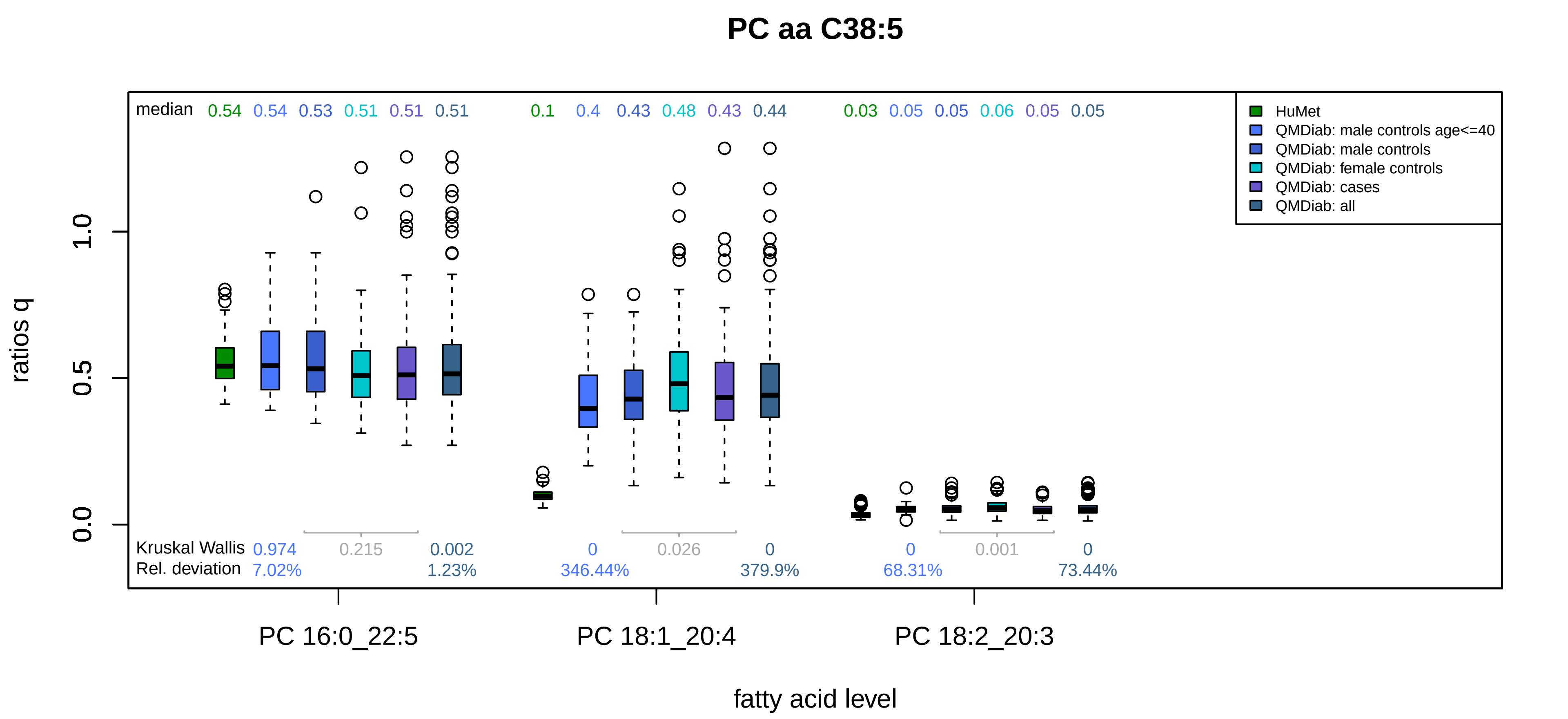

PC aa C38:5
median
0.54 0.54 0.53 0.51 0.51 0.51
0.1 0.4 0.43 0.48 0.43 0.44
0.03 0.05 0.05 0.06 0.05 0.05
HuMet
QMDiab: male controls age<=40
QMDiab: male controls
QMDiab: female controls
QMDiab: cases
QMDiab: all
ratios q
0.0
0.5
1.0
Kruskal Wallis 0.974 0.215 0.002
0 0.026 0
0 0.001 0
Rel. deviation 7.02% 1.23%
346.44% 379.9%
68.31% 73.44%
PC 16:0_22:5
PC 18:1_20:4
PC 18:2_20:3
fatty acid level

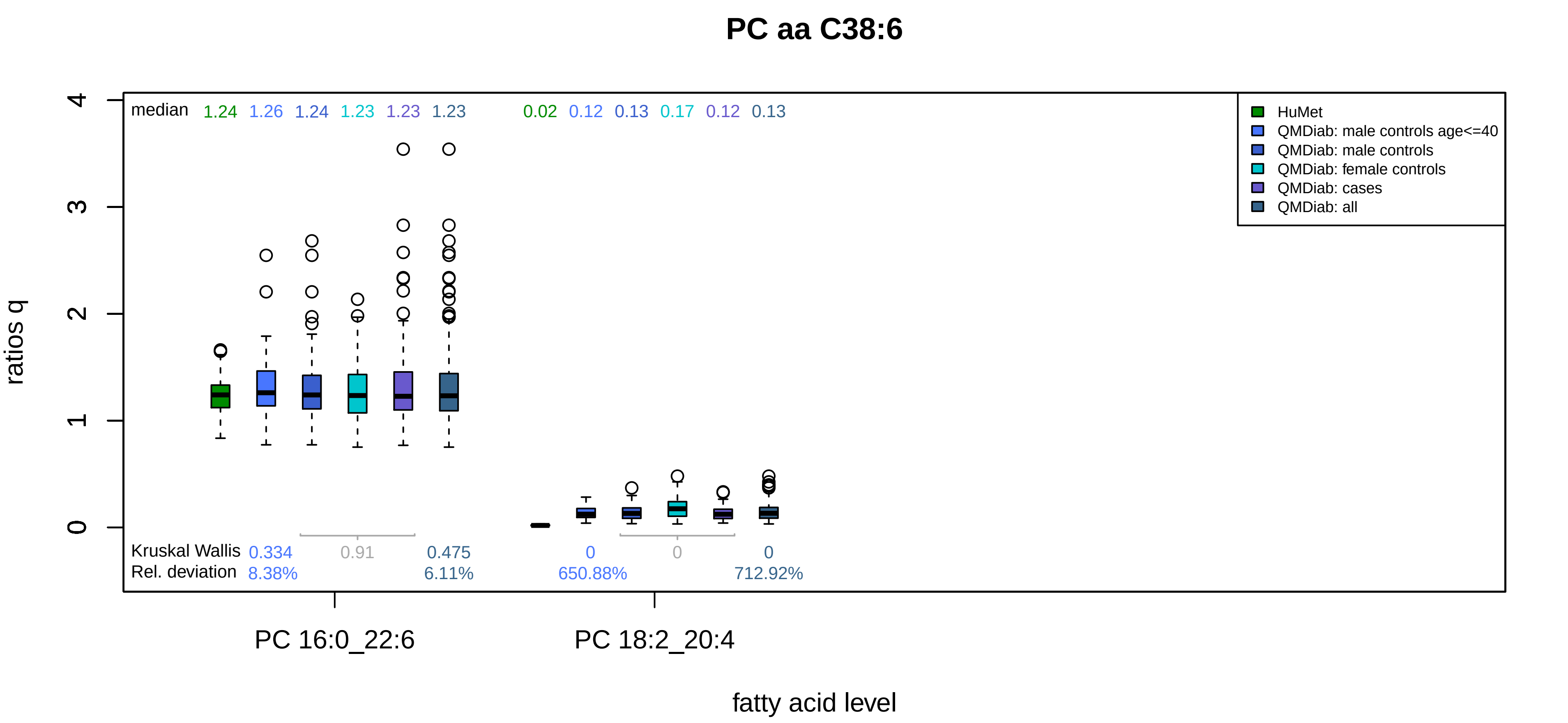
PC aa C38:6
median
1.24
1.26
1.24
1.23
1.23
1.23
0.02
0.12
0.13
0.17
0.12
0.13
HuMet
QMDiab: male controls age<=40
QMDiab: male controls
QMDiab: female controls
QMDiab: cases
QMDiab: all
ratios q
0
1
2
3
4
Kruskal Wallis
0.334
0.91
0.475
0
0
0
Rel. deviation
8.38%
6.11%
650.88%
712.92%
PC 16:0_22:6
PC 18:2_20:4
fatty acid level

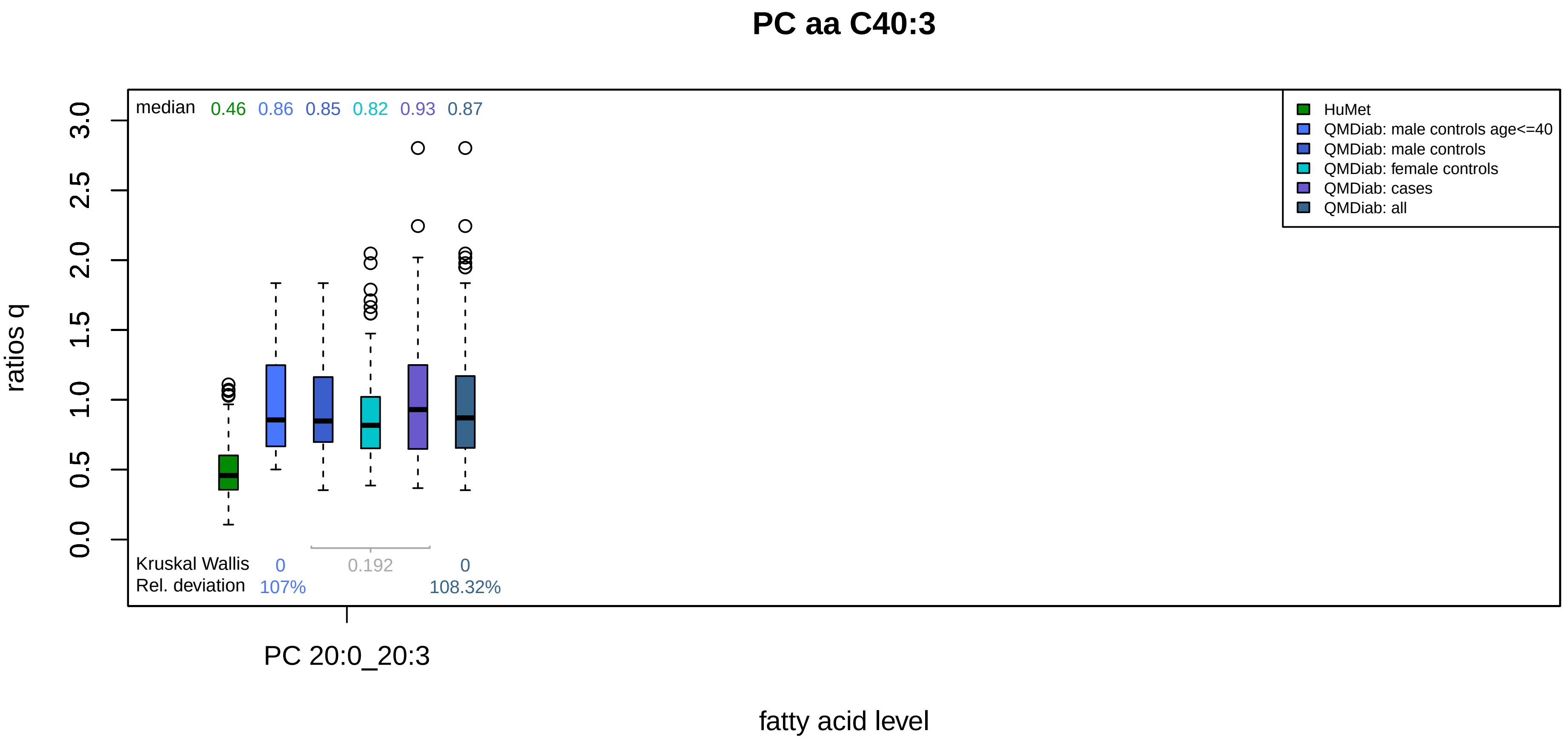
PC aa C40:3
median 0.46 0.86 0.85 0.82 0.93 0.87
HuMet
QMDiab: male controls age<=40
QMDiab: male controls
QMDiab: female controls
QMDiab: cases
QMDiab: all
ratios q
0.0
0.5
1.0
1.5
2.0
2.5
3.0
Kruskal Wallis 0 0.192 0
Rel. deviation 107% 108.32%
PC 20:0_20:3
fatty acid level

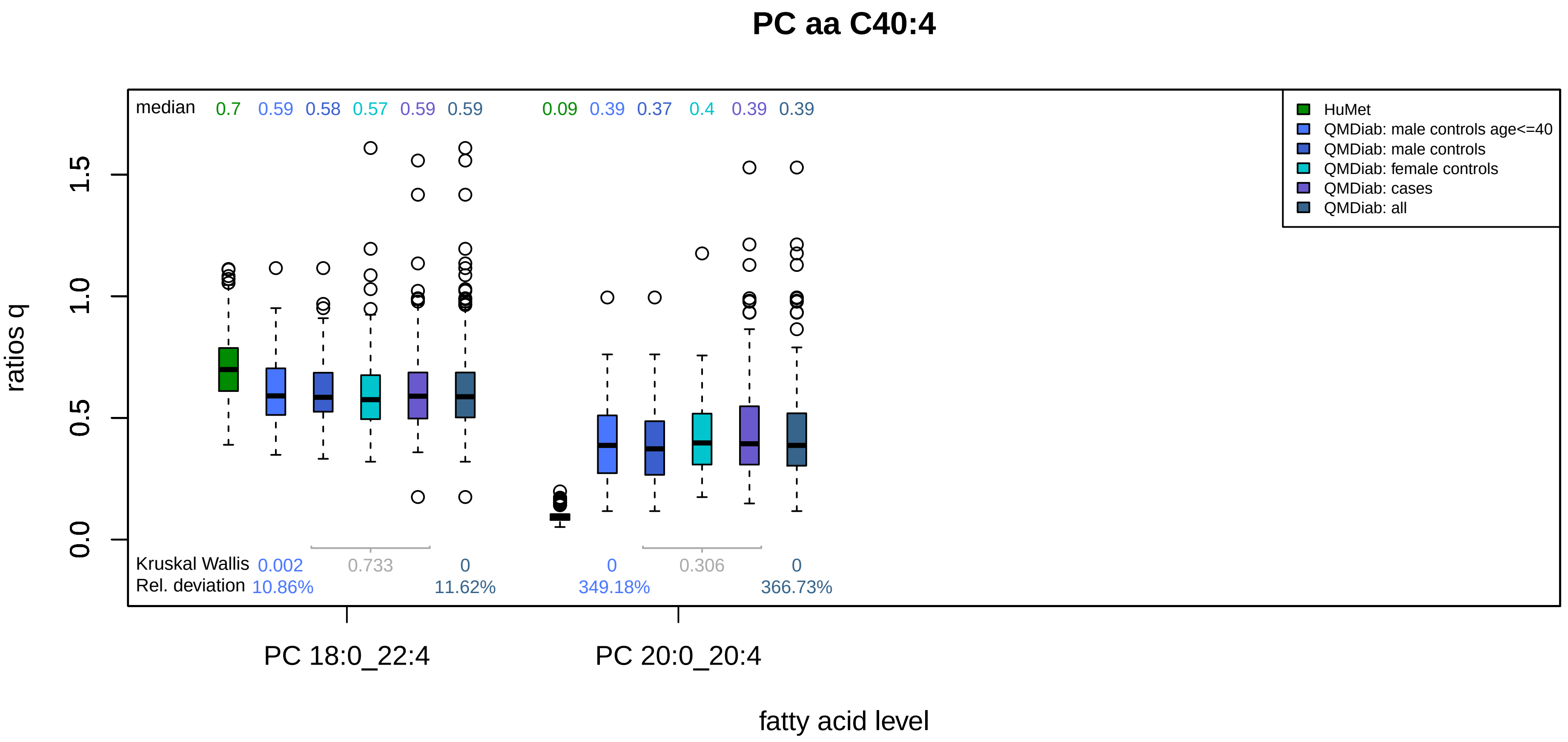
PC aa C40:4
median
0.7
0.59
0.58
0.57
0.59
0.59
0.09
0.39
0.37
0.4
0.39
0.39
HuMet
QMDiab: male controls age<=40
QMDiab: male controls
QMDiab: female controls
QMDiab: cases
QMDiab: all
ratios q
0.0
0.5
1.0
1.5
Kruskal Wallis
0.002
0.733
0
0
0.306
0
Rel. deviation
10.86%
11.62%
349.18%
366.73%
PC 18:0_22:4
PC 20:0_20:4
fatty acid level

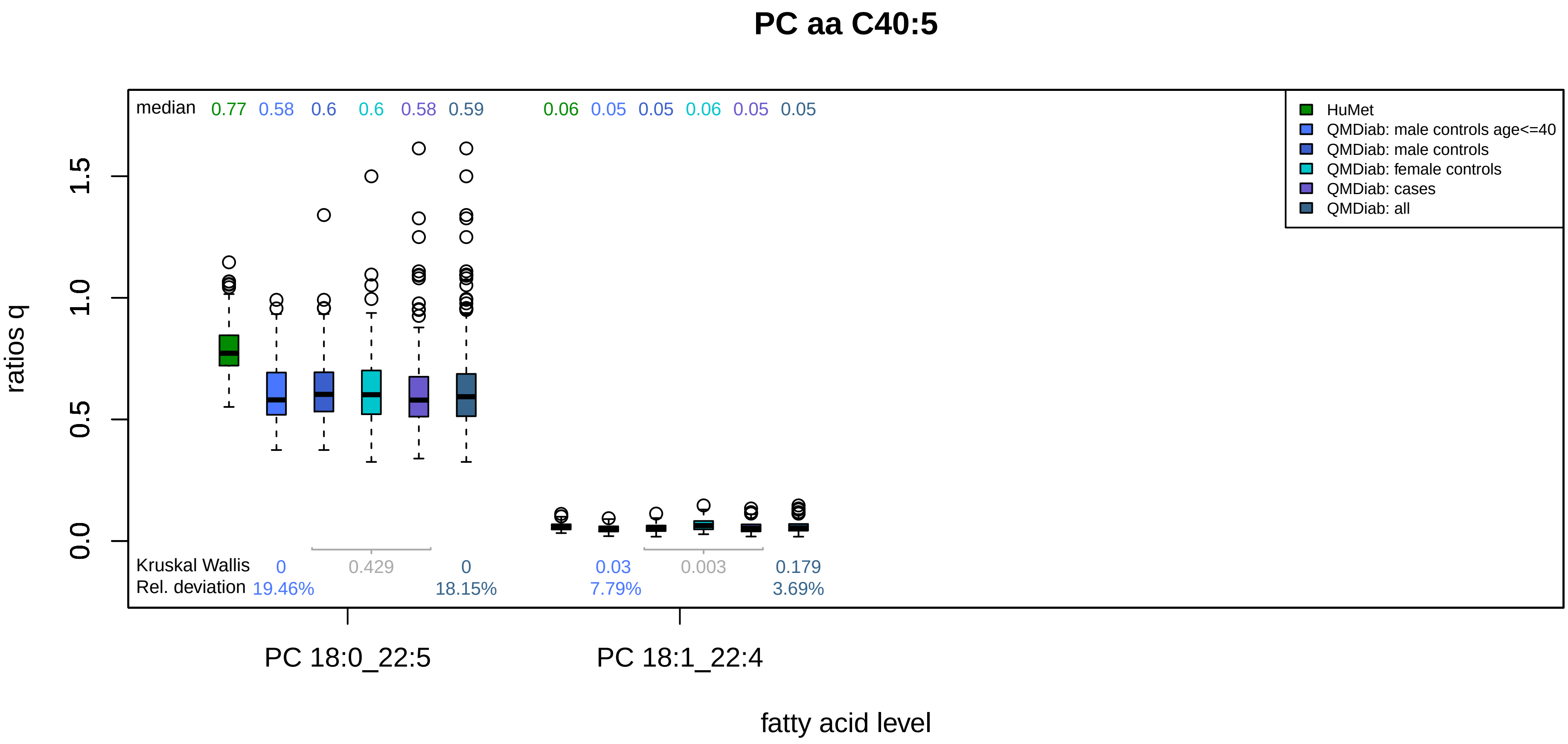

PC aa C40:5
median 0.77 0.58 0.6 0.6 0.58 0.59 0.06 0.05 0.05 0.06 0.05 0.05
HuMet
QMDiab: male controls age<=40
QMDiab: male controls
QMDiab: female controls
QMDiab: cases
QMDiab: all
ratios q
0.0
0.5
1.0
1.5
Kruskal Wallis 0 0.429 0 0.03 0.003 0.179
Rel. deviation 19.46% 18.15% 7.79% 3.69%
PC 18:0_22:5
PC 18:1_22:4
fatty acid level

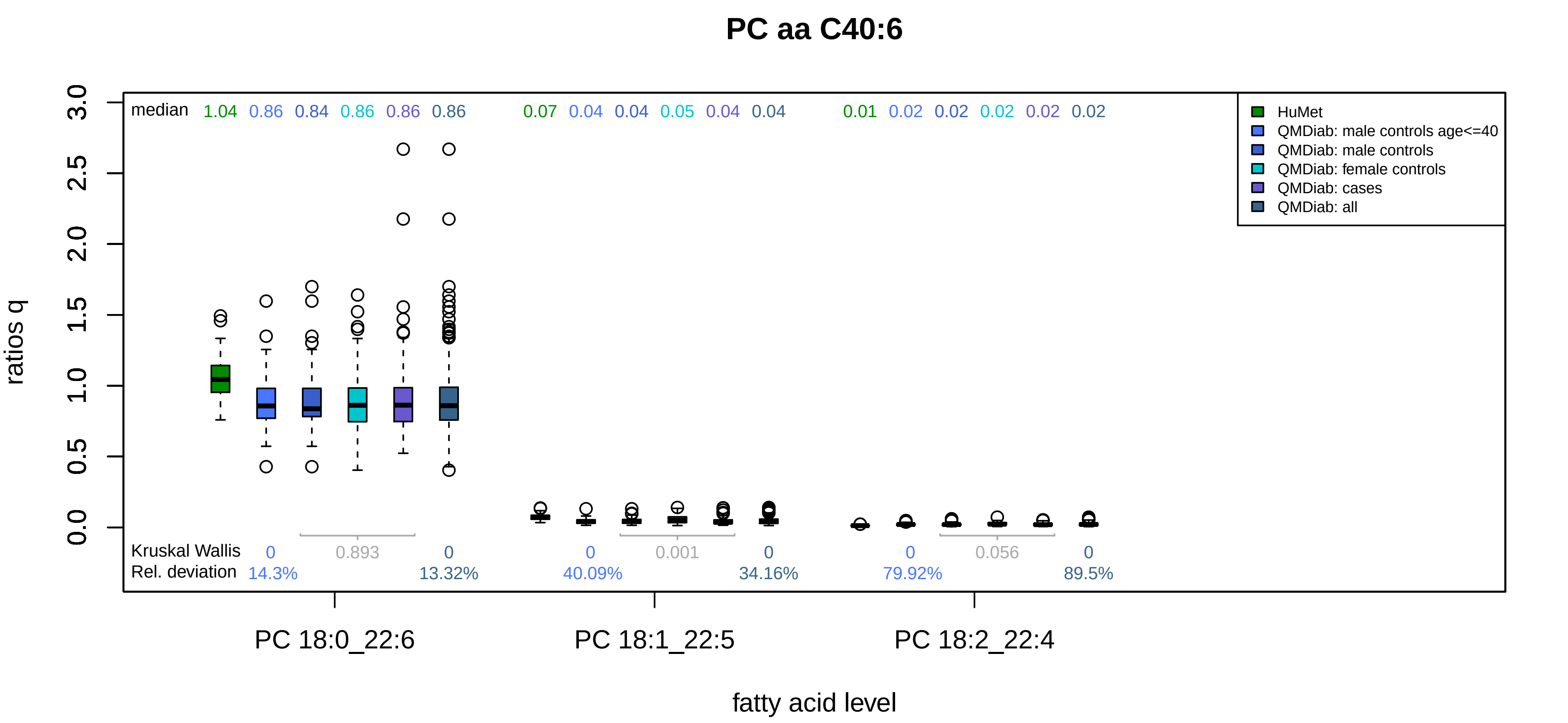
PC aa C40:6
median 1.04 0.86 0.84 0.86 0.86 0.86 0.07 0.04 0.04 0.05 0.04 0.04 0.01 0.02 0.02 0.02 0.02 0.02
HuMet
QMDiab: male controls age<=40
QMDiab: male controls
QMDiab: female controls
QMDiab: cases
QMDiab: all
ratios q
0.0
0.5
1.0
1.5
2.0
2.5
3.0
Kruskal Wallis 0 0.893 0 0 0.001 0 0 0.056 0
Rel. deviation 14.3% 13.32% 40.09% 34.16% 79.92% 89.5%
PC 18:0_22:6
PC 18:1_22:5
PC 18:2_22:4
fatty acid level

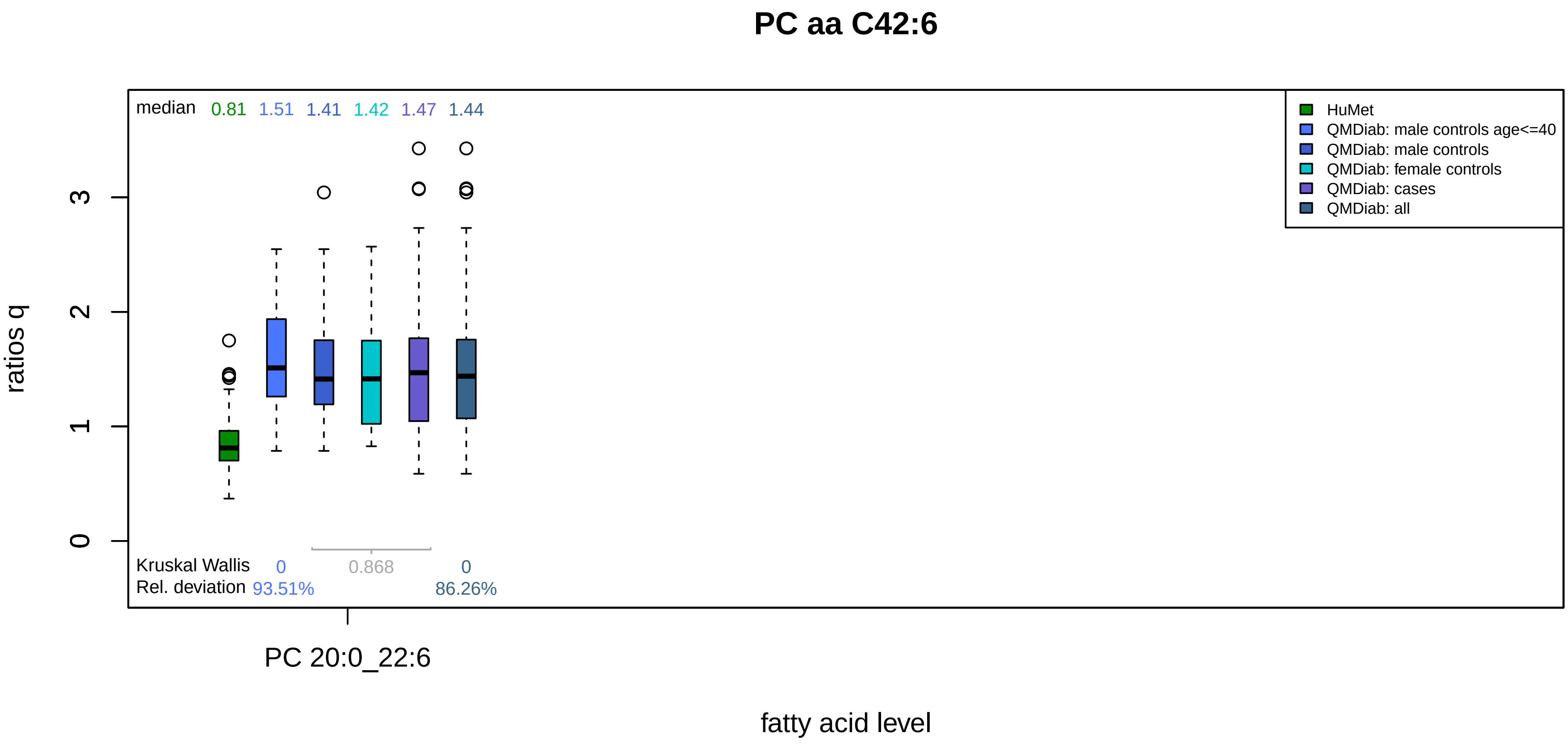

PC aa C42:6
median 0.81 1.51 1.41 1.42 1.47 1.44
HuMet
QMDiab: male controls age<=40
QMDiab: male controls
QMDiab: female controls
QMDiab: cases
QMDiab: all
ratios q
0
1
2
3
Kruskal Wallis 0 0.868 0
Rel. deviation 93.51% 86.26%
PC 20:0_22:6
fatty acid level

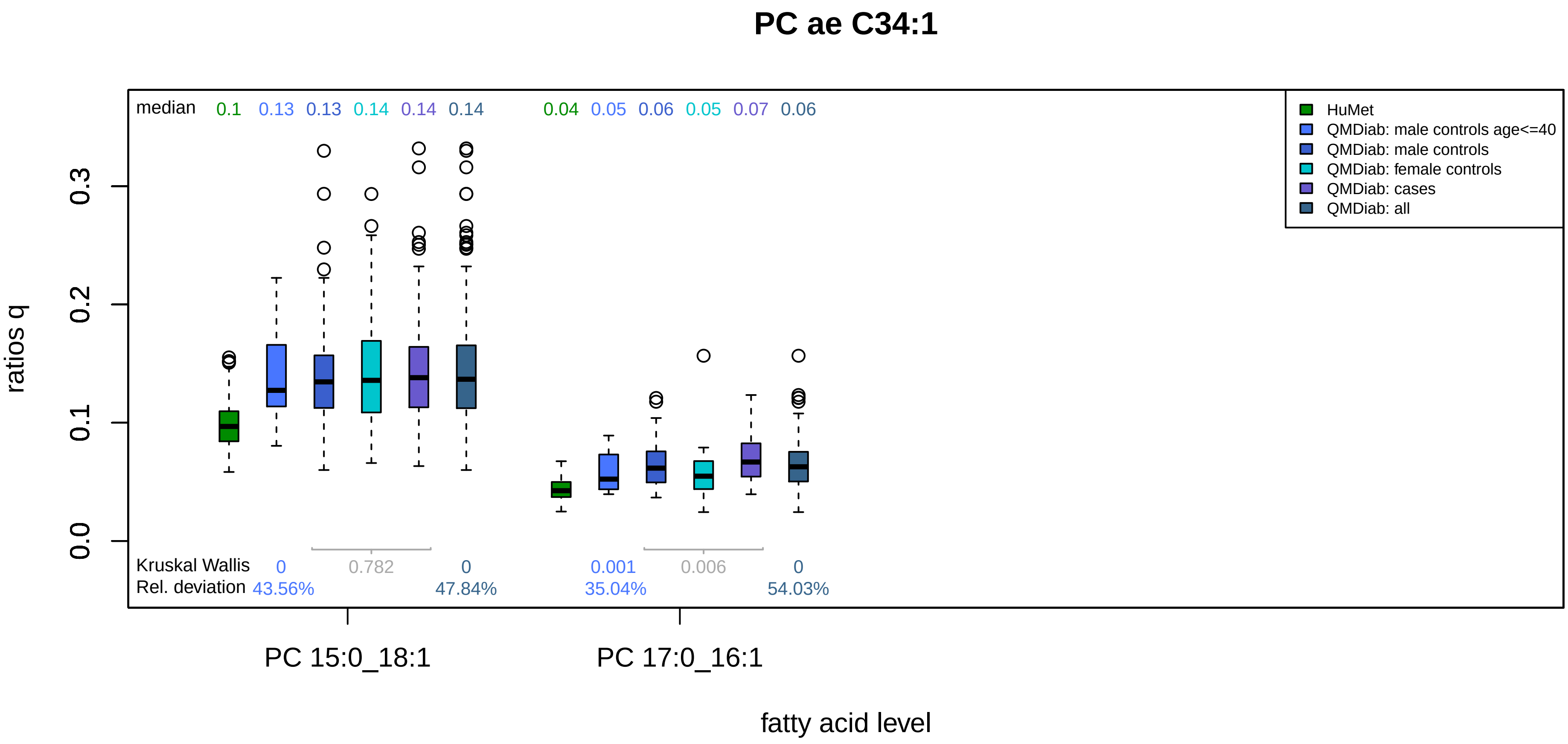

PC ae C34:1
median
0.1
0.13
0.13
0.14
0.14
0.14
0.04
0.05
0.06
0.05
0.07
0.06
HuMet
QMDiab: male controls age<=40
QMDiab: male controls
QMDiab: female controls
QMDiab: cases
QMDiab: all
ratios q
0.0
0.1
0.2
0.3
Kruskal Wallis
0
0.782
0
0.001
0.006
0
Rel. deviation
43.56%
47.84%
35.04%
54.03%
PC 15:0_18:1
PC 17:0_16:1
fatty acid level

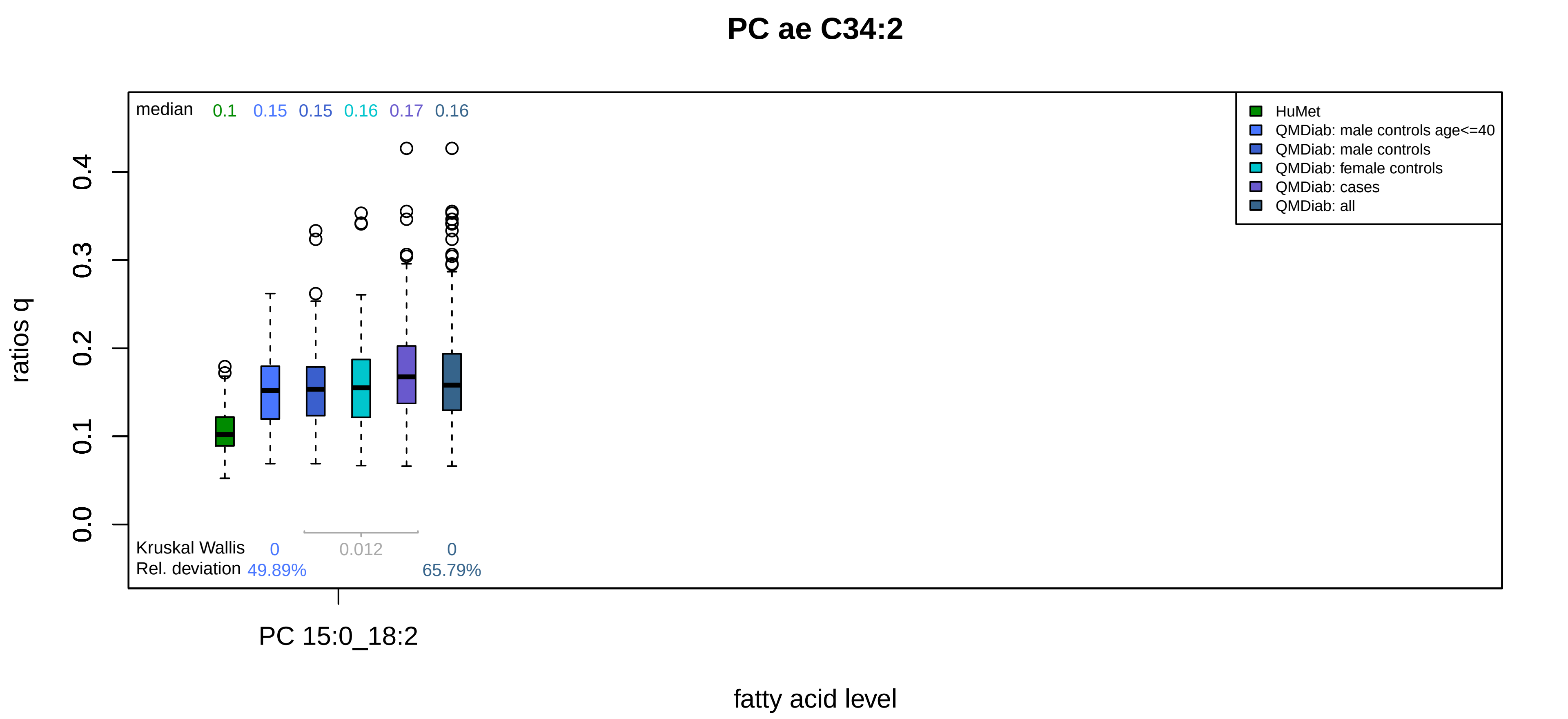

PC ae C34:2
median 0.1 0.15 0.15 0.16 0.17 0.16
HuMet
QMDiab: male controls age<=40
QMDiab: male controls
QMDiab: female controls
QMDiab: cases
QMDiab: all
ratios q
0.0
0.1
0.2
0.3
0.4
Kruskal Wallis 0 0.012 0
Rel. deviation 49.89% 65.79%
PC 15:0_18:2
fatty acid level

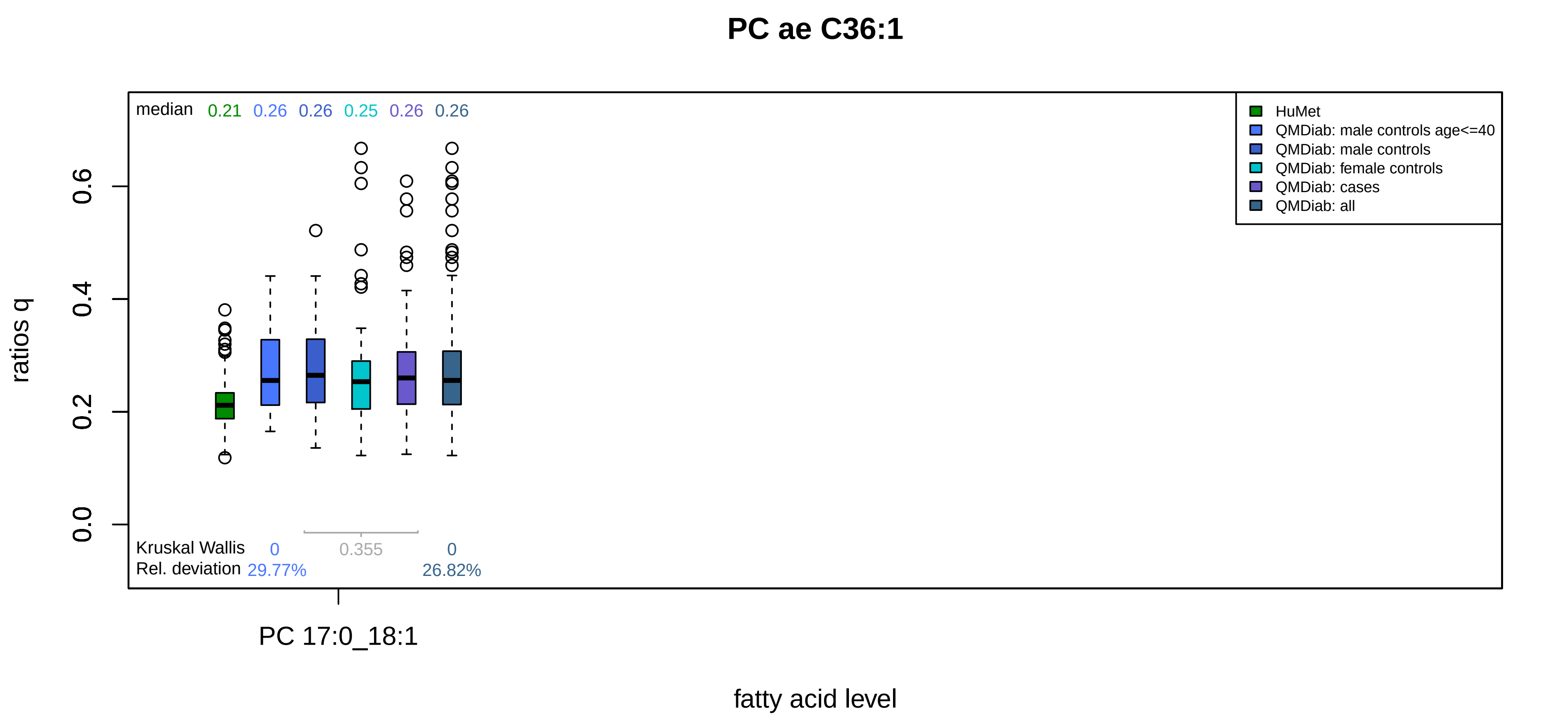

PC ae C36:1
median 0.21 0.26 0.26 0.25 0.26 0.26
HuMet
QMDiab: male controls age<=40
QMDiab: male controls
QMDiab: female controls
QMDiab: cases
QMDiab: all
ratios q
0.0
0.2
0.4
0.6
Kruskal Wallis 0 0.355 0
Rel. deviation 29.77% 26.82%
PC 17:0_18:1
fatty acid level

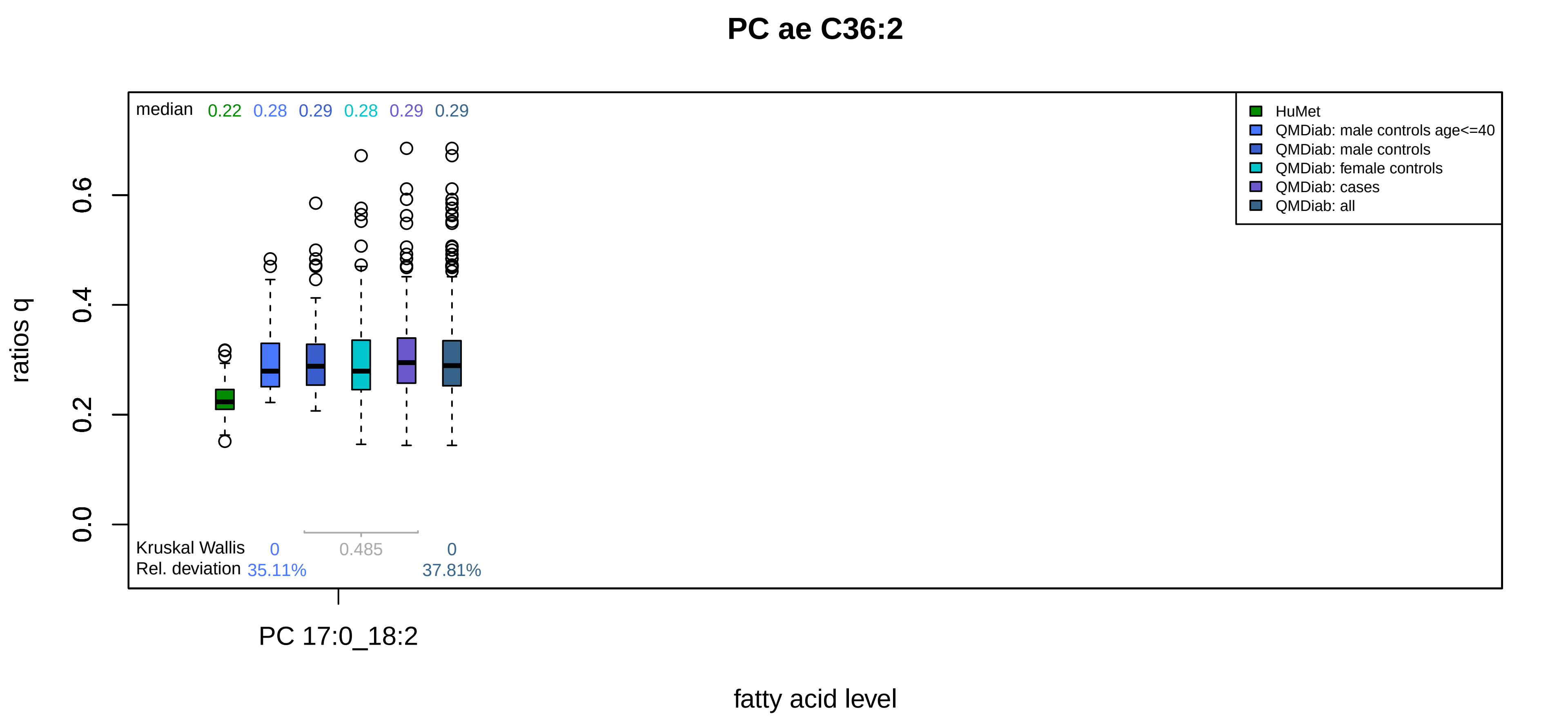

PC ae C36:2
median 0.22 0.28 0.29 0.28 0.29 0.29
HuMet
QMDiab: male controls age<=40
QMDiab: male controls
QMDiab: female controls
QMDiab: cases
QMDiab: all
ratios q
0.0
0.2
0.4
0.6
Kruskal Wallis 0 0.485 0
Rel. deviation 35.11% 37.81%
PC 17:0_18:2
fatty acid level

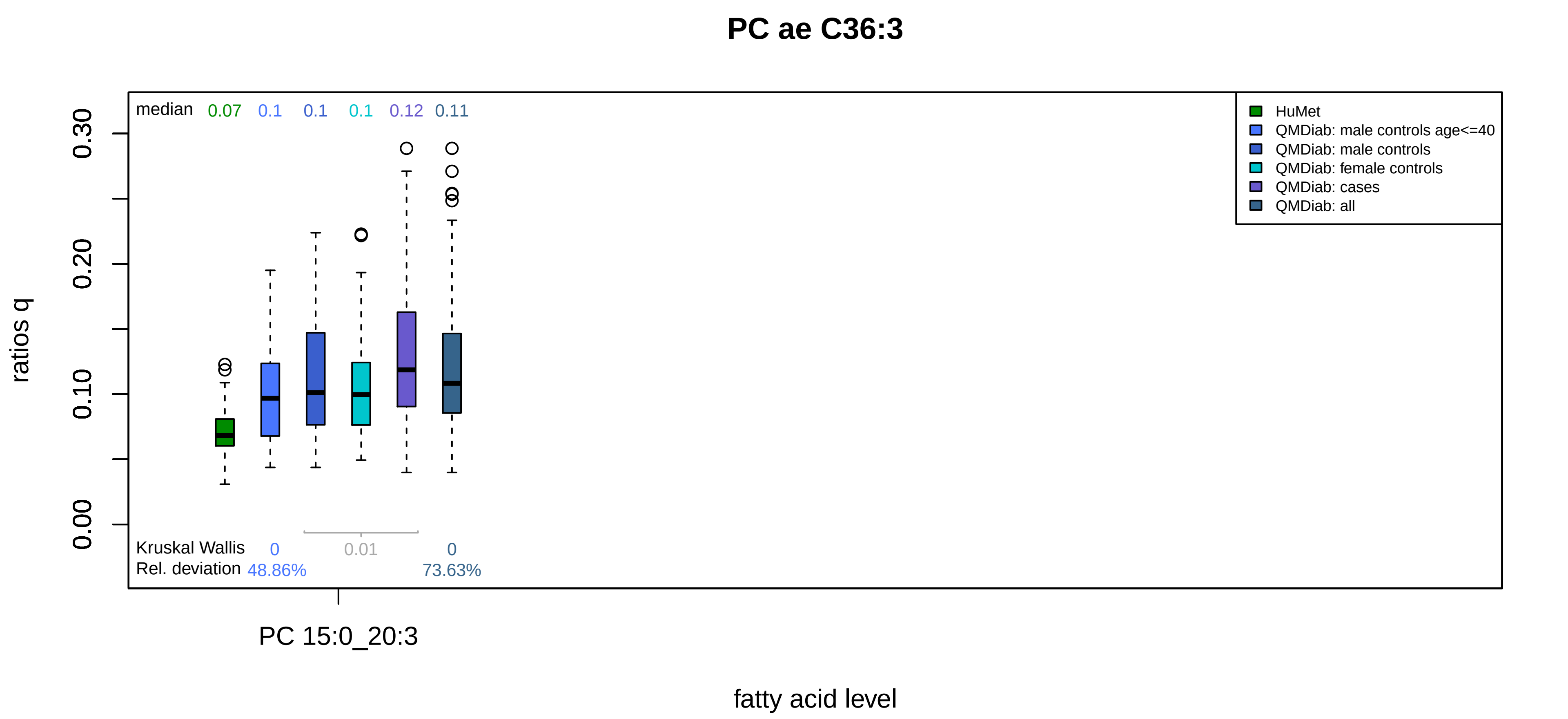
PC ae C36:3
median 0.07 0.1 0.1 0.1 0.12 0.11
HuMet
QMDiab: male controls age<=40
QMDiab: male controls
QMDiab: female controls
QMDiab: cases
QMDiab: all
ratios q
0.00
0.10
0.20
0.30
Kruskal Wallis 0 0.01 0
Rel. deviation 48.86% 73.63%
PC 15:0_20:3
fatty acid level

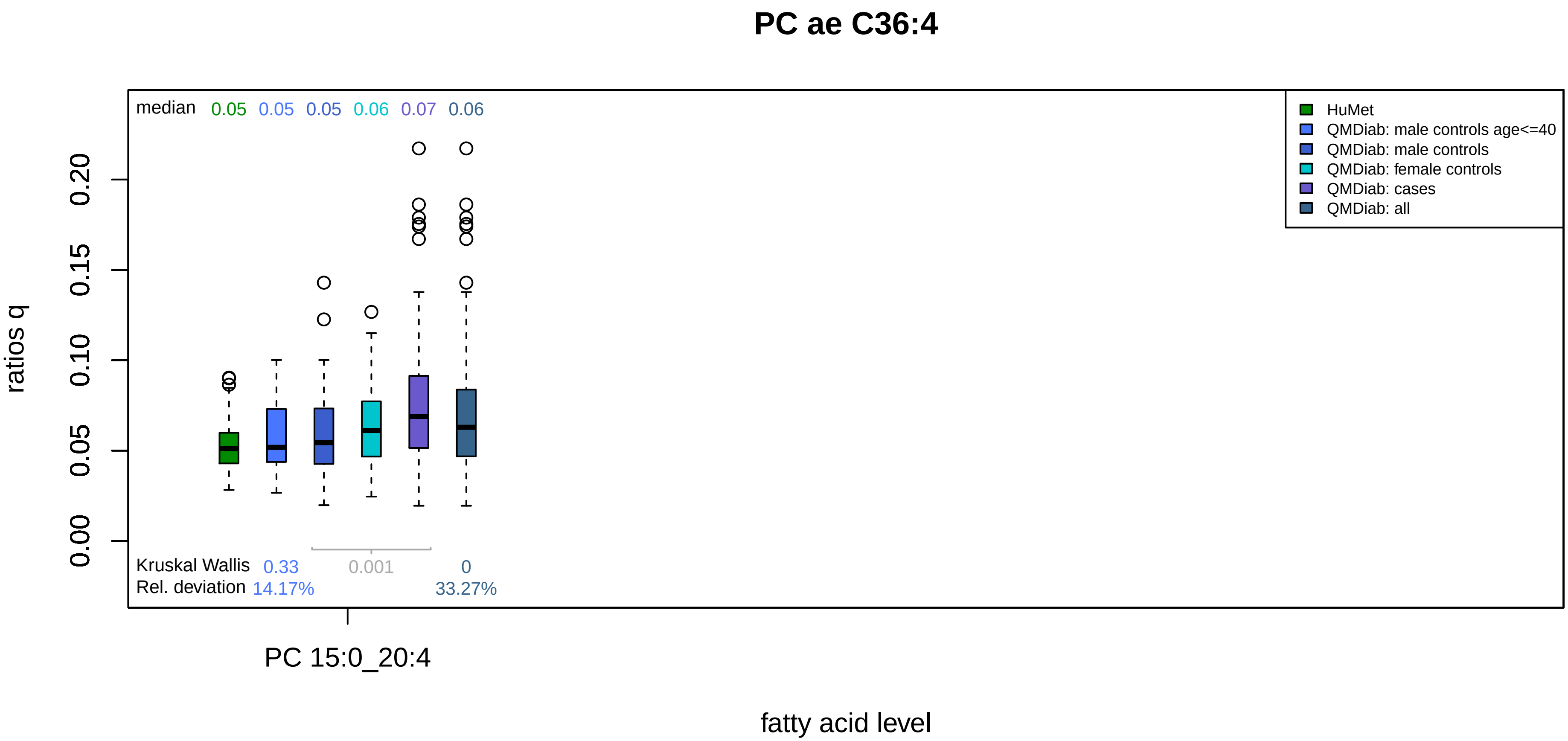
PC ae C36:4
median 0.05 0.05 0.05 0.06 0.07 0.06
HuMet
QMDiab: male controls age<=40
QMDiab: male controls
QMDiab: female controls
QMDiab: cases
QMDiab: all
ratios q
0.00
0.05
0.10
0.15
0.20
Kruskal Wallis 0.33 0.001 0
Rel. deviation 14.17% 33.27%
PC 15:0_20:4
fatty acid level

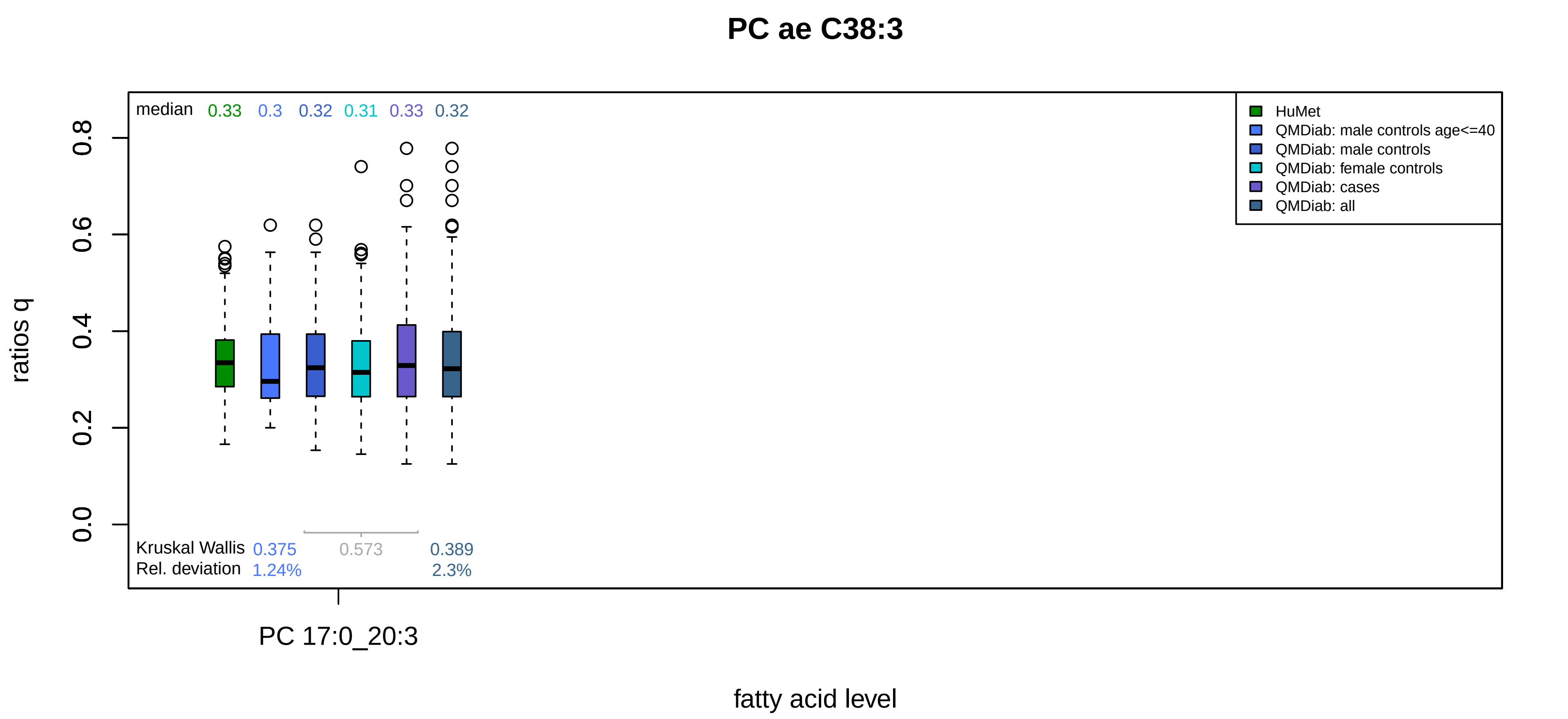

PC ae C38:3
median 0.33 0.3 0.32 0.31 0.33 0.32
HuMet
QMDiab: male controls age<=40
QMDiab: male controls
QMDiab: female controls
QMDiab: cases
QMDiab: all
ratios q
0.0
0.2
0.4
0.6
0.8
Kruskal Wallis 0.375 0.573 0.389
Rel. deviation 1.24% 2.3%
PC 17:0_20:3
fatty acid level

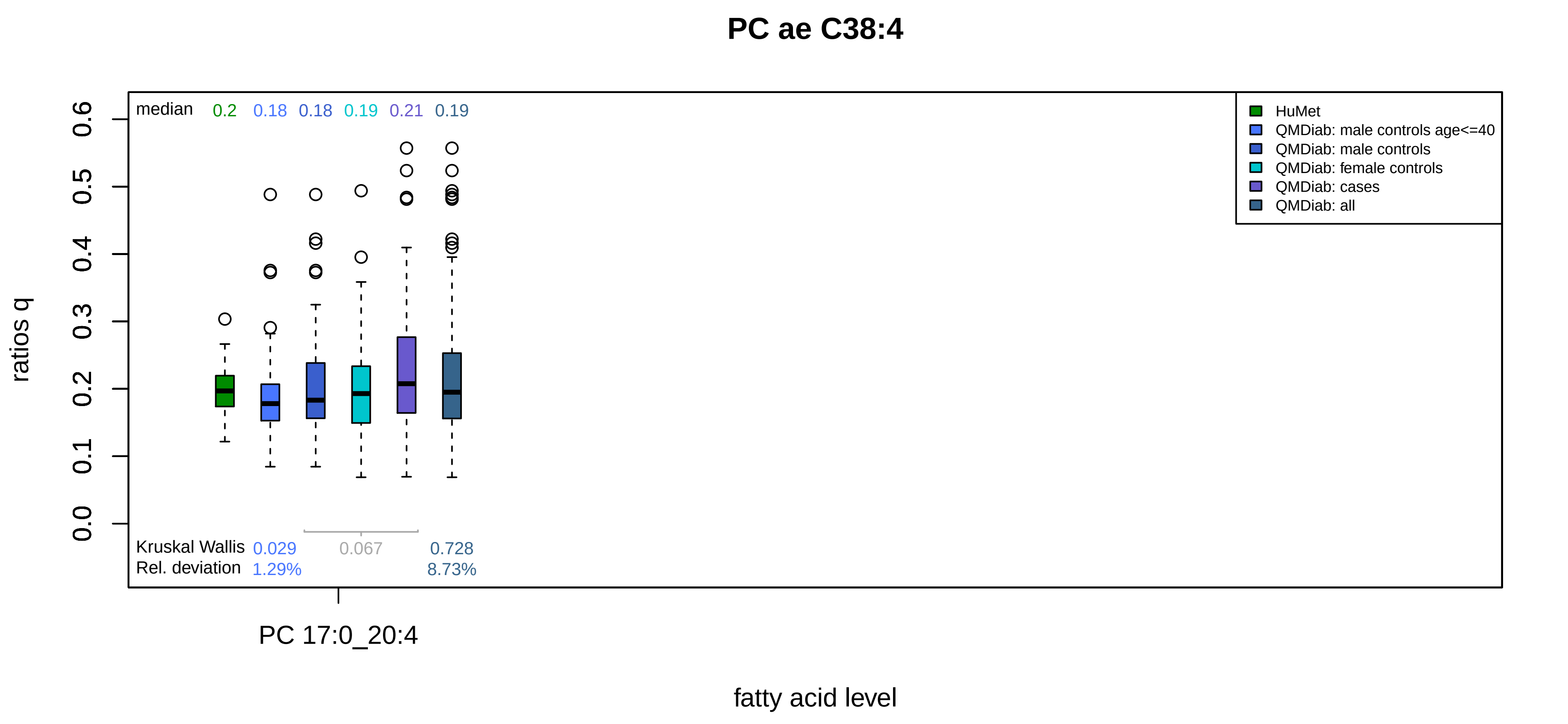
PC ae C38:4
median 0.2 0.18 0.18 0.19 0.21 0.19
HuMet
QMDiab: male controls age<=40
QMDiab: male controls
QMDiab: female controls
QMDiab: cases
QMDiab: all
ratios q
0.0
0.1
0.2
0.3
0.4
0.5
0.6
Kruskal Wallis 0.029 0.067 0.728
Rel. deviation 1.29% 8.73%
PC 17:0_20:4
fatty acid level

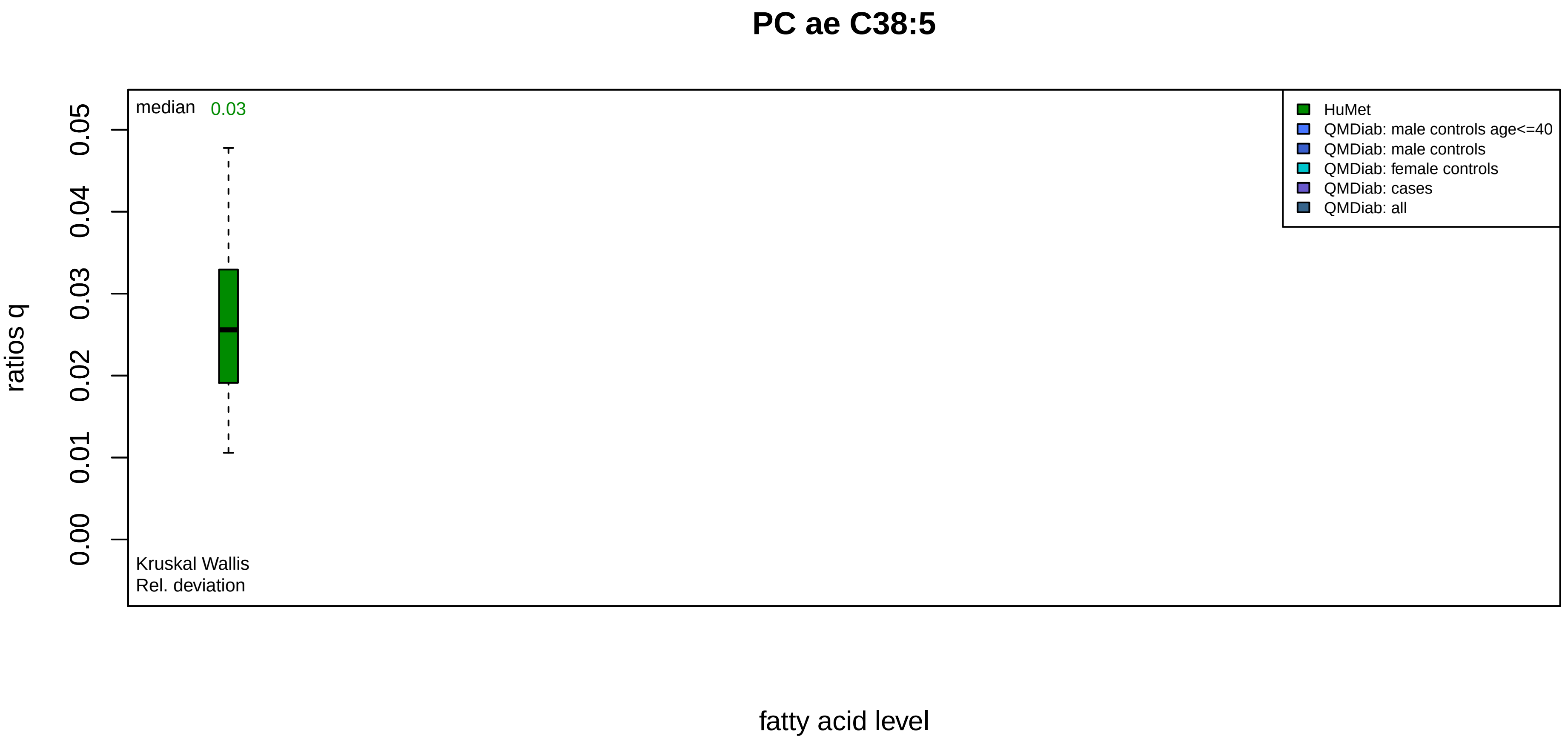

PC ae C38:5
median
0.03
HuMet
QMDiab: male controls age<=40
QMDiab: male controls
QMDiab: female controls
QMDiab: cases
QMDiab: all
ratios q
0.00
0.01
0.02
0.03
0.04
0.05
Kruskal Wallis
Rel. deviation
fatty acid level

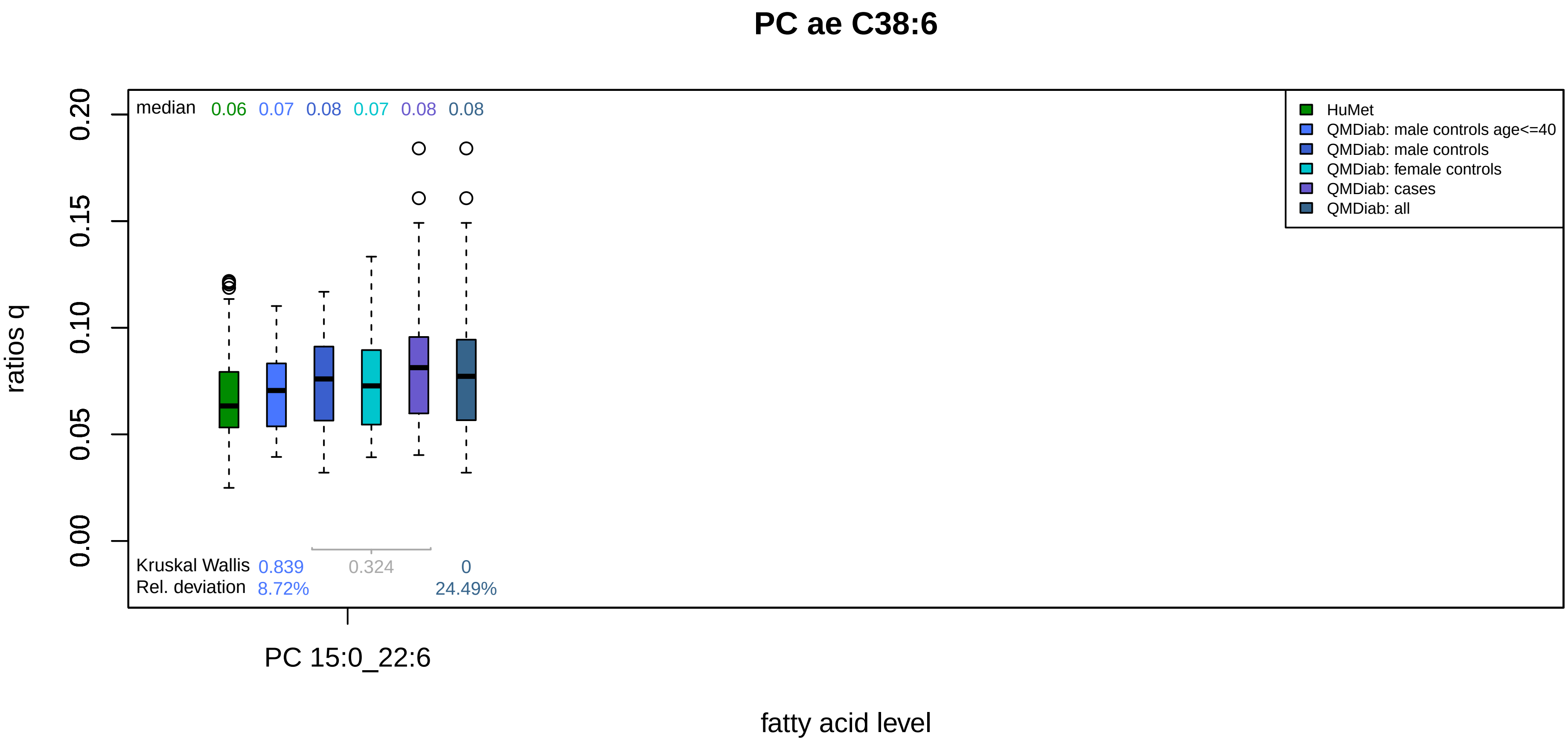

PC ae C38:6
median 0.06 0.07 0.08 0.07 0.08 0.08
HuMet
QMDiab: male controls age<=40
QMDiab: male controls
QMDiab: female controls
QMDiab: cases
QMDiab: all
ratios q
0.00
0.05
0.10
0.15
0.20
Kruskal Wallis 0.839 0.324 0
Rel. deviation 8.72% 24.49%
PC 15:0_22:6
fatty acid level

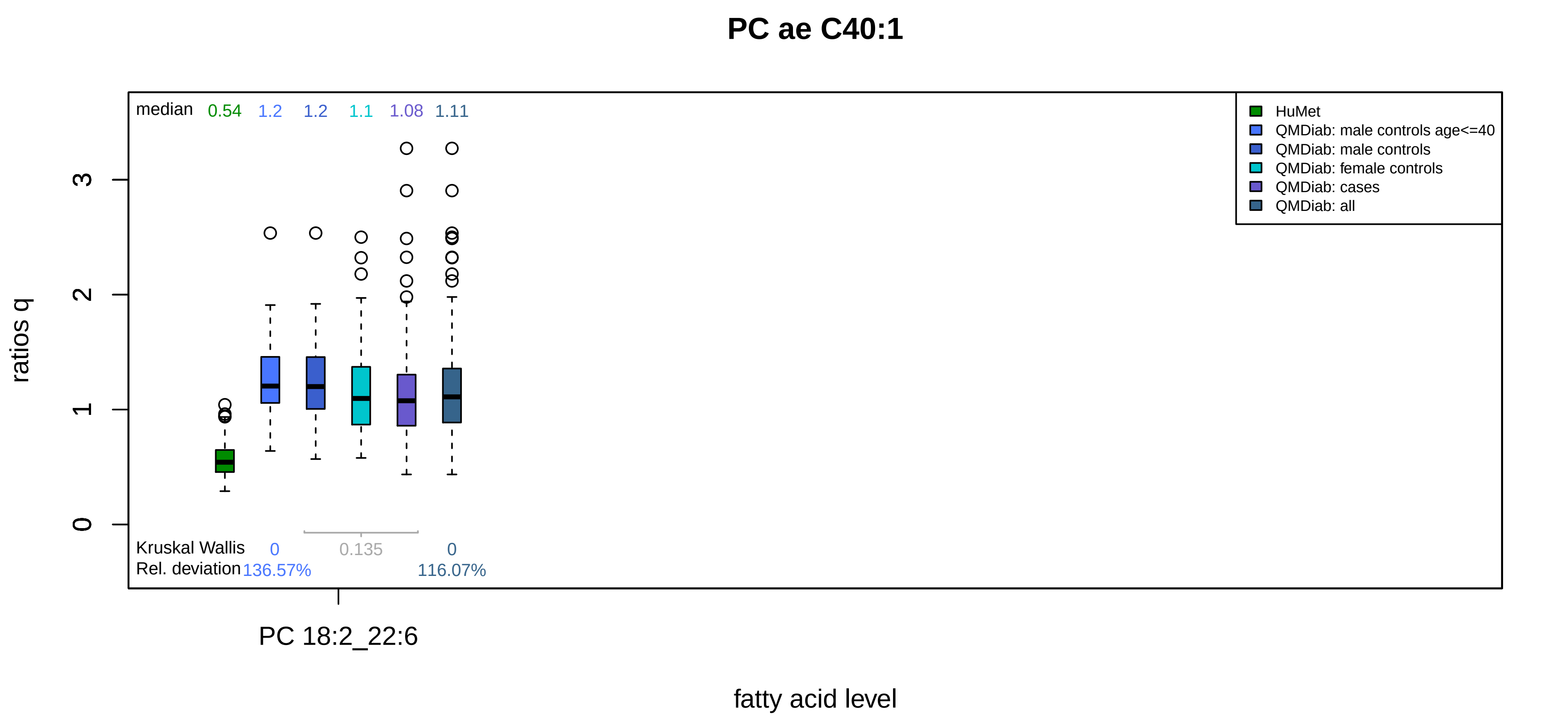
PC ae C40:1
median 0.54 1.2 1.2 1.1 1.08 1.11
HuMet
QMDiab: male controls age<=40
QMDiab: male controls
QMDiab: female controls
QMDiab: cases
QMDiab: all
ratios q
0
1
2
3
Kruskal Wallis 0 0.135 0
Rel. deviation 136.57% 116.07%
PC 18:2_22:6
fatty acid level

# PC ae C40:5

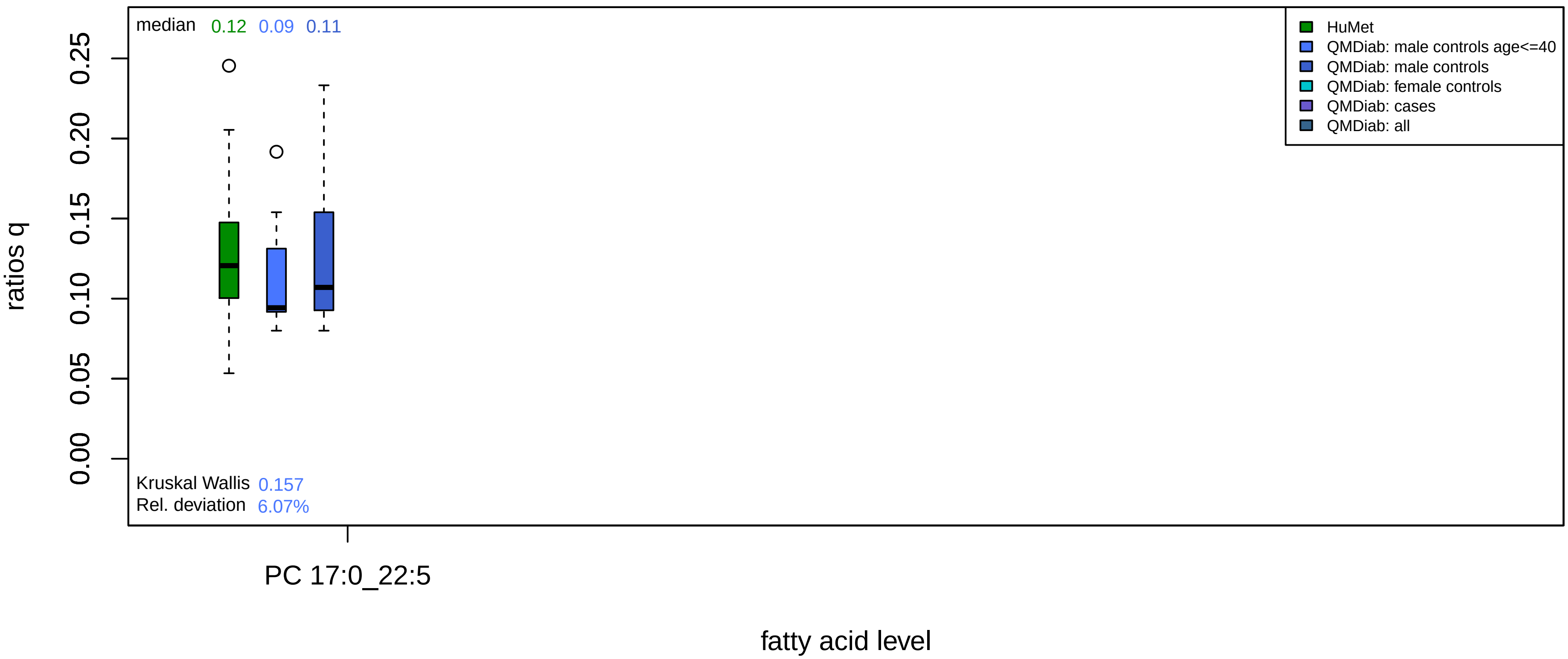

median 0.12 0.09 0.11
HuMet
QMDiab: male controls age<=40
QMDiab: male controls
QMDiab: female controls
QMDiab: cases
QMDiab: all
ratios q
0.00
0.05
0.10
0.15
0.20
0.25
Kruskal Wallis 0.157
Rel. deviation 6.07%
PC 17:0_22:5
fatty acid level

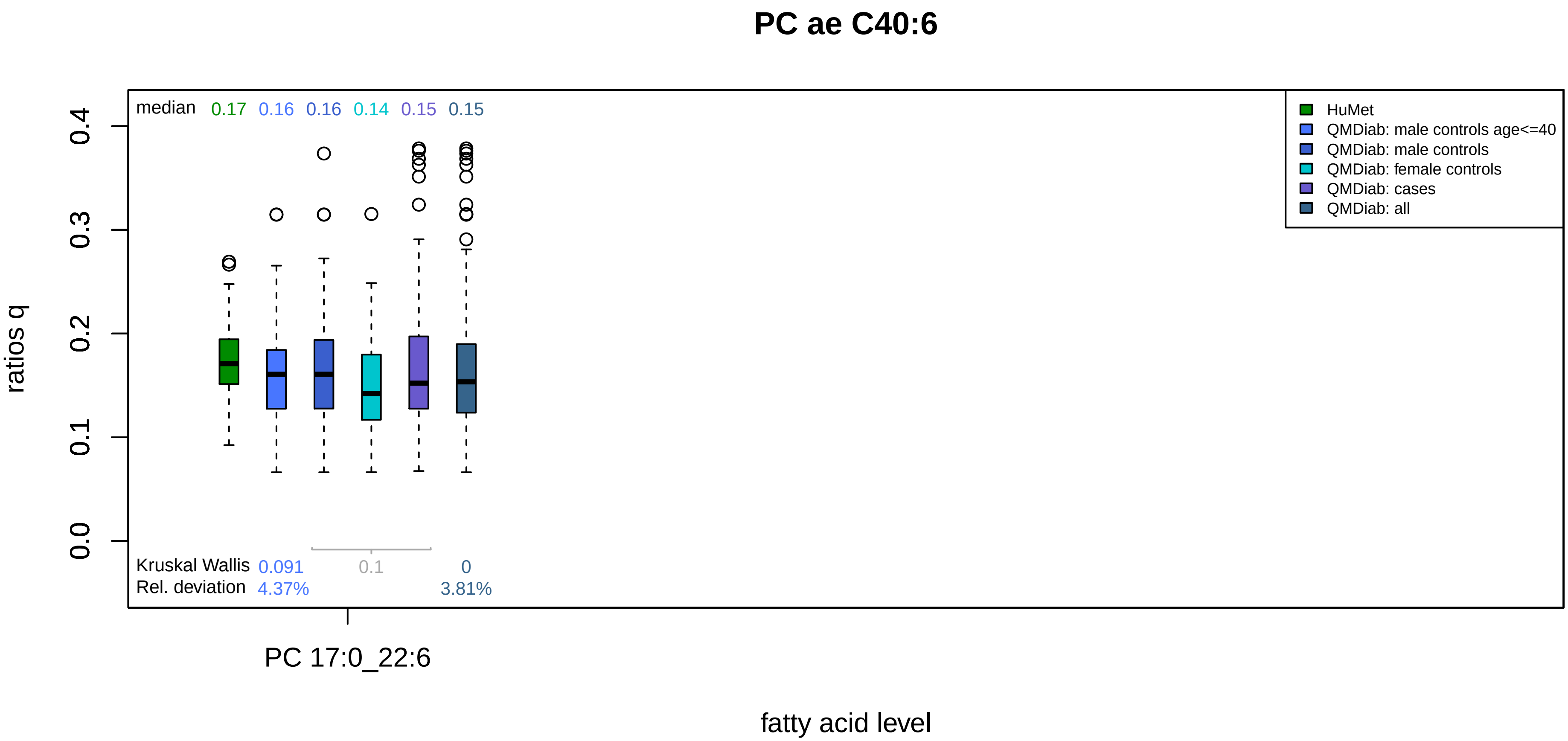
PC ae C40:6
median 0.17 0.16 0.16 0.14 0.15 0.15
HuMet
QMDiab: male controls age<=40
QMDiab: male controls
QMDiab: female controls
QMDiab: cases
QMDiab: all
ratios q
0.0
0.1
0.2
0.3
0.4
Kruskal Wallis 0.091 0.1 0
Rel. deviation 4.37% 3.81%
PC 17:0_22:6
fatty acid level
